# An acetyl-click screening platform identifies a small molecule inhibitor of Histone Acetyltransferase 1 (HAT1) with anti-tumor activity

**DOI:** 10.1101/2021.06.25.449993

**Authors:** Joshua J. Gruber, Amithvikram Rangarajan, Tristan Chou, Benjamin S. Geller, Selene Banuelos, Robert Greenhouse, Michael P. Snyder, Andrew M. Lipchik

## Abstract

HAT1 is a central regulator of chromatin synthesis that acetylates nascent histone H3:H4 tetramers in the cytoplasm. It may have a role in cancer metabolism by linking cytoplasmic production of acetyl-CoA to nuclear acetyl flux. This is because the HAT1 di-acetylation mark is not propagated in chromatin and instead is de-acetylated after nascent histone insertion into chromatin. Thus, HAT1 likely provides a nuclear source of free acetate that may be recycled to acetyl-CoA for nuclear acetylation reactions. Correspondingly, suppression of HAT1 protein expression impairs tumor growth. To ascertain whether targeting HAT1 is a viable anti-cancer treatment strategy we sought to identify small molecule inhibitors of HAT1. We developed a high-throughput HAT1 acetyl-click assay to facilitate drug discovery and enzymology. Screening of small molecules computationally predicted to bind the active site led to the discovery of multiple riboflavin analogs that inhibited HAT1 enzymatic activity by competing with acetyl-CoA binding. These hits were refined by synthesis and testing over 70 analogs, which yielded structure-activity relationships. The isoalloxazine core was required for enzymatic inhibition, whereas modifications of the ribityl sidechain improved enzymatic potency and cellular growth suppression. These efforts resulted in a lead compound (JG-2016) that suppressed growth of human cancer cells lines *in vitro* and impaired tumor growth *in vivo*. This is the first report of a small molecule inhibitor of the HAT1 enzyme complex and represents a step towards targeting this pathway for cancer therapy.

## INTRODUCTION

Nutrient metabolism and epigenetic reactions are integrated and sensed by cells to ensure that adequate substrates are available to meet the demands of transcriptional programs that spur cell division.^1^ By studying genes induced by epidermal growth factor (EGF) we identified HAT1 as the human acetyltransferase most highly induced by EGF stimulation in mammary cells^2^. HAT1 was also required for rapid cell proliferation and tumor formation *in vivo*.^2-6^ These data indicate that HAT1 plays a critical role in the coordinating anabolic and epigenetic processes for cell division that drives tumor growth.

HAT1 was the first histone acetyltransferase gene isolated,^7-9^ and subsequent work has established that it plays a critical role in chromatin replication, the process of making new nucleosomes during S-phase.^10^ In the cytosol, HAT1 di-acetylates histone H4 on lysines 5 and 12 of the amino-terminal histone tail. It then transits to the nucleus together with the histone tetramer and other histone chaperones^11^ to deposit nascent histones at the replication fork, or other sites of nucleosome insertion. Then HAT1 is released from chromatin^12^ and the HAT1 di-acetylation mark on histone H4 is quickly removed within a span of 15-30 minutes by the action of histone deacetylases.^13-15^ Thus, HAT1 does not directly acetylate chromatin, and the di-acetylation mark placed by HAT1 is not propagated to mature chromatin.

Our prior work suggested a model whereby free acetate derived from de-acetylation of nascent histones was recycled to acetyl-CoA via acetyl-CoA-synthetases to provide substrate for nascent chromatin acetylation.^2^ As each nascent nucleosome of newly replicated chromatin contributes 4 HAT1-dependent acetyl groups, this should provide adequate acetyl-CoA to allow for histone acetylation of the much sparser promoter and enhancer sites. Indeed, histone H3 acetylation marks are reduced in cells depleted from HAT1, as expected from this model.^2^ In addition, CBP auto-acetylation is strongly dependent on HAT1^16^ which also suggests a role for HAT1 in governing nuclear acetyl flux. Other links between mitochondrial processes and HAT1 function have also been reported.^5,17^

Although these genetic studies have shed light on HAT1 function, further development of chemical probes should prove useful to distinguish the importance of the enzyme’s catalytic activity from its structural role in protein:protein or protein:DNA complexes. Recently, a HAT1 bisubstrate inhibitor was designed by chemically ligating coenzyme A to the ζ-amine of lysine 12 in the histone H4 N-terminal 20-mer peptide, yielding a K_i_ of ∼ 1 nM towards bacterially-expressed recombinant HAT1.^18^ Although, useful for enzymatic assays, this probe is not cell permeable and therefore of limited utility to study cellular processes dependent on HAT1. Therefore, we sought to identify and design small molecule inhibitors of HAT1 to probe the effects of HAT1 activity in cells and validate its role as a pro-tumorigenic factor.

The design of small molecule acetyltransferase inhibitors has been hampered by non-specific and low-throughput assays, which have tended to yield bio-reactive molecules.^19,20^ However, specific, potent, small molecule acetyltransferase inhibitors targeting CBP/p300^21^and KAT6A/B^22,23^ have recently been described. We previously pioneered peptide-based sensors of non-receptor tyrosine kinases for cellular detection and monitoring of post-translational modification events including small molecule kinase inhibitor treatments.^24^ We then built a generalizable in silico pipeline to design, optimize and screen kinase-specific peptide substrates for drug discovery screens.^25^ The use of lanthanide coordination by peptide substrates allowed for the development of screening assays without the need for post-translational modification-specific antibodies.^26,27^ Therefore, these approaches were adapted with recent advances in acetylation monitoring^28^ to build a high-throughput, peptide-based sensor assay for HAT1 acetylation activity to facilitate drug discovery.

## RESULTS

### Design and validation of a HAT1 high-throughput enzymatic assay

HAT1 chemical probe screens or high-throughput enzymatic assays have yet to be described. Therefore, we designed a HAT1 enzymatic assay to specifically and rapidly measure the HAT1 di-acetylation product using a click-chemistry approach (Fig. 1A). The click-chemistry acetyl-CoA analog 4-pentynoyl-CoA allows for enzymatic transfer of an alkyne handle via an acylation reaction. First, the HAT1 enzyme complex (HAT1 + Rbap46) purified from human cells is co-incubated with 4-pentynoyl-CoA co-factor and a biotinylated H4 N-terminal peptide to allow for HAT1-dependent pentoylation of the peptide substate at lysines 5 and 12. Then, reaction products are bound to a neutravidin capture plate, followed by Cu(I)-catalyzed alkyne-azide cycloaddition with biotin-azide. After streptavidin-HRP binding, fluorescence signal can be detected by peroxidation of amplex red.

**Figure 1:**
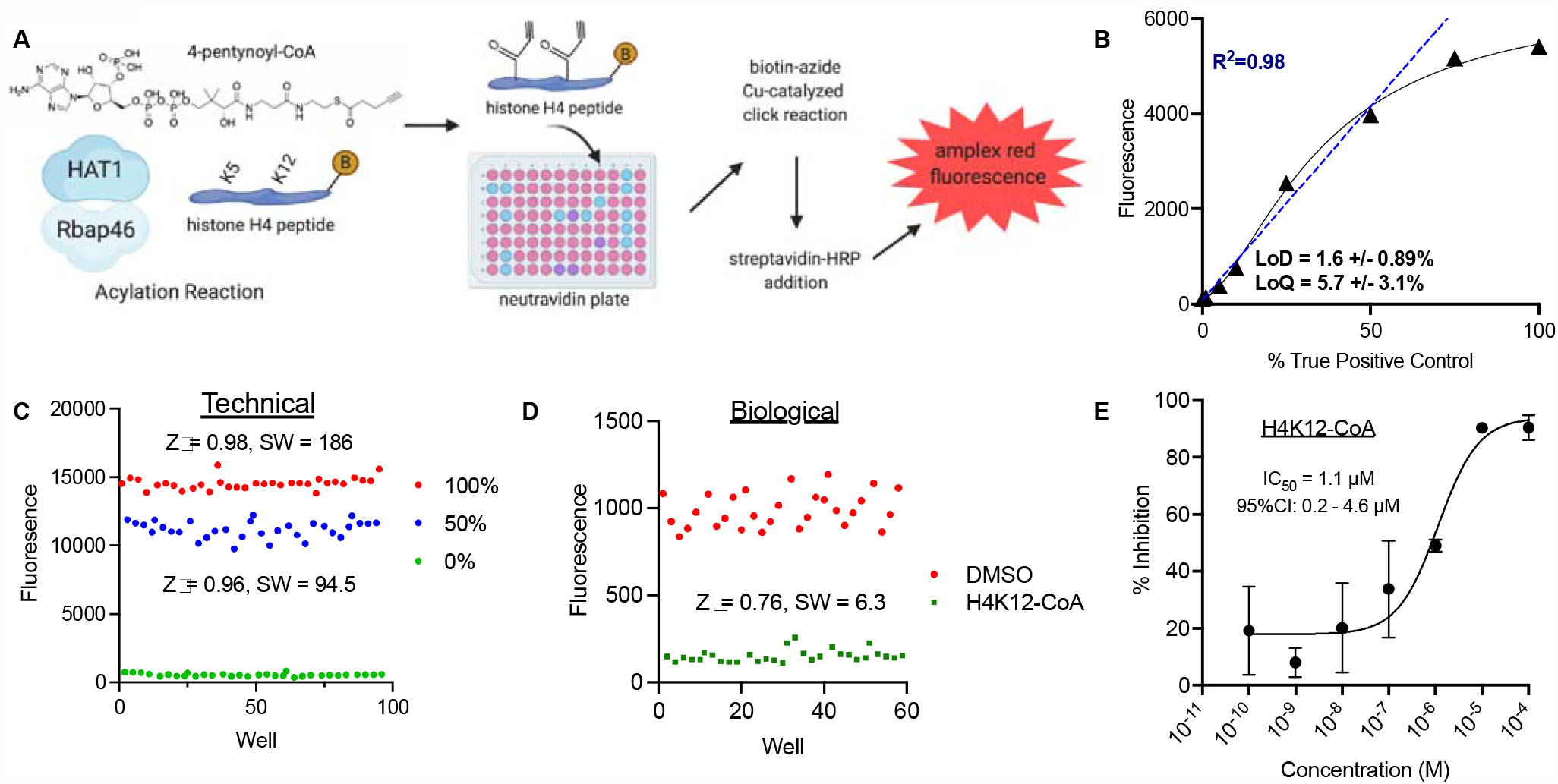
Design and validation of a HAT1 high-throughput enzymatic assay. A. Schematic of HAT1 acetylation assay. HAT1/Rbap46 purified enzyme complex is incubated with 4-pentynoyl-CoA and histone H4 N-terminal biotinylated peptide in the presence or absence of test compounds. Then reaction products are bound to neutravidin 96-well plate, washed, functionalized by copper(I)-catalyzed cycloaddition click chemistry with biotin-azide. Then, streptavidin-HRP is bound and detected by amplex red fluorescence reaction. B. Standard curve generated by mixing biotinylated positive control peptide (H4 N-terminal peptide with propargylglycine residues substituted for lysines 5 and 12) with the unmodified histone H4 N-terminal peptide of equal length, followed by binding products to neutravidin plate, click chemistry addition of biotin-azide, streptavidin-HRP binding and amplex red detection. Black line indicates nonlinear curve fit through all data. Blue dashed line is a linear fit for data from 0-50% positive control (R^2^=0.98). Limit of detection (LoD) and limit of quantitation (LoQ) are mean ± SD for three independent replicates. C. High-throughput characterization of technical replicates of 100%, 50% or 0% propargylglycine containing H4 peptides mixed with unmodified peptides spotted in checkboard pattern in 96-well neutravidin-plates. Z□ score and signal window are calculated for the difference between 0-100% (top) and 0-50% (below) peptide mixtures. D. High-throughput characterization of biological replicates of HAT1 acetylation reactions treated with either DMSO control or positive control bisubstrate inhibitor H4K12-CoA (10 μM). E. Dose-response study of HAT1 acetylation assay treated with H4K12-CoA. IC_50_ was calculated by 3 parameter least-squares regression assuming Hill slope -1.0.

This assay was dependent on exogenous expression and co-purification of both HAT1 and Rbap46 from a human cell line (Supp Fig. 1A). In contrast, utilization of HAT1 alone was less active, as previously described.^29^ Next, assay performance was optimized (Supp. Fig. 1B) and quantified. A standard curve was generated using a histone H4 peptide manufactured to have terminal alkynes at the 5^th^ and 12^th^ positions of the H4 N-terminal peptide (Fig. 1B) allowing for quantification of reaction products. The assay demonstrated linear performance (R^2^ = 0.98) when up to 50% true positive control peptides were used. Therefore, assay conditions were calibrated to produce signal within this upper threshold. The limit of detection (LoD) was 1.6 0.89% defined as the percentage of positive control peptide that gave a fluorescence reading corresponding to 3x the standard deviation of the unacetylated negative control peptide above baseline. The limit of quantitation (LoQ) was 5.7 ± 3.1% defined as the percentage of positive control peptide that gave fluorescence output corresponding to 10x the standard deviation greater than the baseline negative control.^30^ Thus the assay was designed to operate at fluorescent detection values greater than the LoQ.

Next, high-throughput assay performance was characterized using technical controls. A 96-well plate was arrayed in checkboard configuration consisting of H4 N-terminal peptides with 0%, 50% or 100% alkyne-containing positive control peptides (Fig. 1C). This allowed us to calculate the Z□ factor and the signal window (SW) for the maximal range of the assay (0-100%) and also for the linear range of the assay (0-50%). A Z□ factor between 0.5 and 1 indicates the assay performs adequately for high-throughput screening (HTS) as it indicates a robust dynamic range. Similarly, a SW > 2 also indicates wide separation between positive and negative controls suitable for HTS. Both the maximal range (0-100%) and linear range (0-50%) of the assay demonstrated excellent performance characteristics (Z□> 0.5, SW >2; Fig. 1C). Therefore, the assay is technically suitable for HTS.

To determine if the assay maintained appropriate HTS parameters under biological conditions the HAT1/Rbap46 complex was incubated with and without the H4K12-CoA bi-substrate inhibitor (positive control inhibitor), which caused robust inhibition with suitable high-throughput performance metrics Z□ and SW (Fig. 1D). To further demonstrate the biological characteristics of the assay we measured the IC_50_ for HAT1 inhibition by the bi-substrate inhibitor. H4K12-CoA inhibited HAT1/Rbap46 activity with an IC_50_ of ∼1 μM (Fig. 1E). As this bi-substrate inhibitor has been shown to be a specific inhibitor of HAT1, but not other acetyltransferases,^18^ this validates the specificity of our HAT1 high-throughput assay. Finally, although lysyl-CoA is a validated bi-substrate inhibitor for CBP/p300,^31^ it had no inhibitory activity towards HAT1/Rbap46 in our assay (Supp. Fig. 1C), in accordance with prior results from low-throughput assays,^18^ thereby demonstrating our assay specificity. Together these results demonstrate the ability of the HAT1/Rbap46 assay to detect biologically relevant acetylation.

### Structural modeling coupled with enzymatic screening to identify HAT1 inhibitors

Human HAT1 has been crystalized at high-resolution (1.9-Å) revealing residues that comprise binding surfaces for the histone H4 N-terminal substrate and the acetyl-CoA cofactor.^32^ The pantotheine moiety of acetyl-CoA resides in a canyon that orients the thioester bond in close proximity to the lysine 12 side-chain of H4, which is the preferred substrate site for acetyl transfer.^8,32^ We used this structural data to build a virtual docking workflow to identify small molecules with appropriate physiochemical properties to occupy the cofactor binding site (Fig. 2A). The NCI open collection of 265,242 molecules was screened against the HAT1 co-factor binding sites using Schrodinger Glide-based virtual screening with increasing precision cutoffs (see Methods). We focused on the top 0.001% of compounds yielding 274 hits from the starting collection. Of these, 39 were obtained and screened with our HAT1 acetylation assay at a single dosage (100 μM, equivalent to the co-factor concentration). The best compound (NSC-42186) was a natural-product derivative of riboflavin that displayed 31% inhibition at 100 μM (Fig. 2B). NSC-42186 showed appropriate dose-response activity with an enzymatic IC_50_ of 60.5 μM (95% CI: 20 – 115 μM; Fig. 2C). Thus, this combined virtual and enzymatic screening approach identified small molecule candidate HAT1 inhibitors.

**Figure 2:**
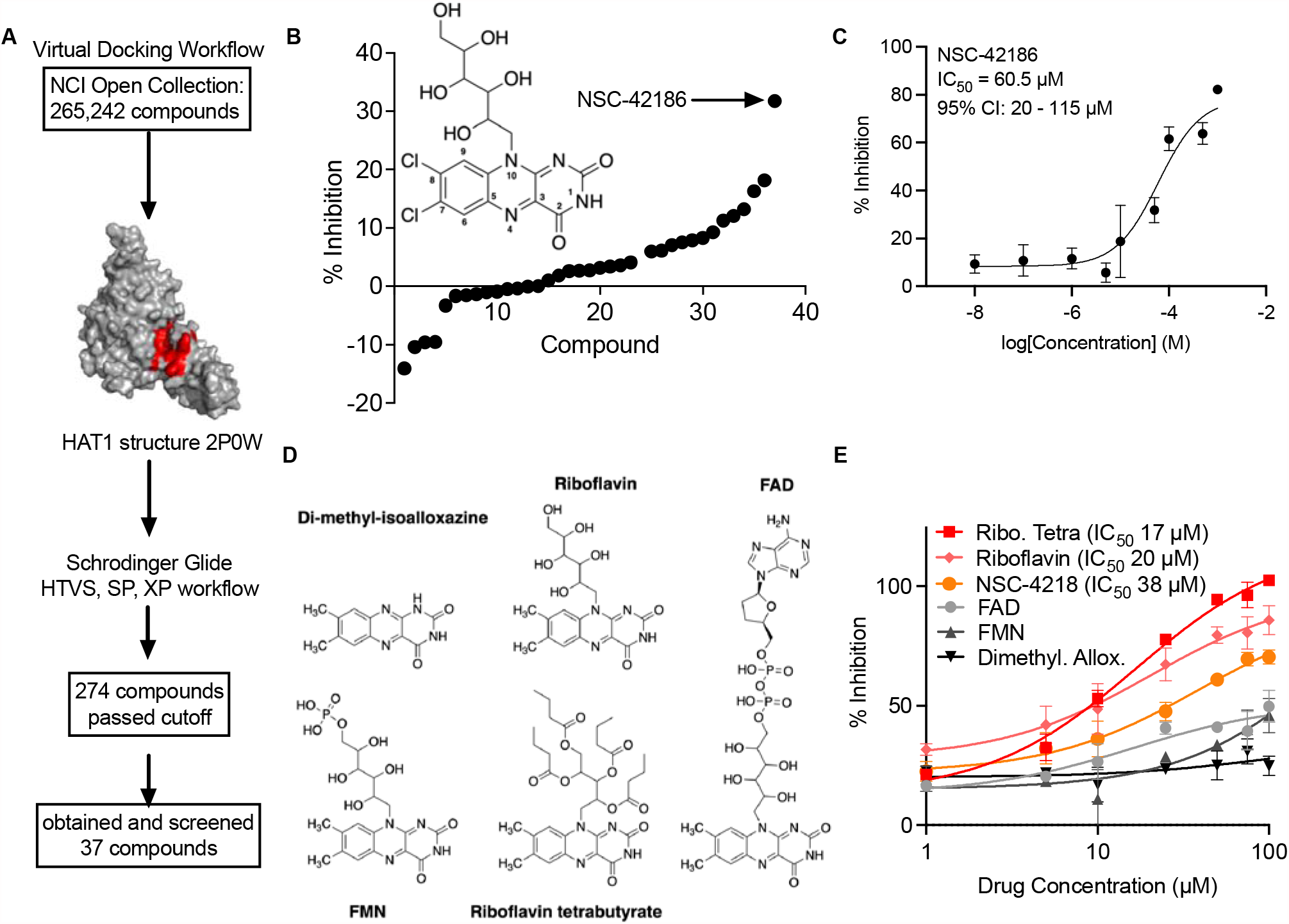
Structural modeling coupled with enzymatic screening to identify HAT1 inhibitors. A. Schematic of virtual docking workflow used to pre-select small molecules capable of binding the the acetyl-CoA cofactor binding site (red) of the HAT1 crystal structure 2P0W. The entire NCI open chemical library was screened with Schrodinger Glide with SP and XP modes to select the top 0.001% of compounds predicted to bind. Compounds were obtained from the NCI/DTP repository. B. Thirty-seven compounds from the NCI/DTP repository screened in the HAT1 acetylation assay. Inset is structure of the best hit NSC-42186. Positions around the isoalloxazine core are numbered for further reference throughout the text. C. Dose-response of NSC-42186 treatment in the HAT1 acetylation assay. IC_50_ was calculated by 3 parameter least-squares regression assuming Hill slope -1.0. D. Chemical structures of riboflavin analogs and derivatives obtained and tested. E. Dose-response of riboflavin analogs in (D) tested in the HAT1 acetylation assay.

NSC-42186 contains 7,8-di-chloro substitutions of tri-cyclic isoalloxazine ring that are the sole features that distinguish it from the 7,8-di-methyl isoalloxazine of riboflavin (ring numbering scheme is shown in Fig. 2B). Riboflavin, commonly known as vitamin B2, is an essential nutrient in metazoans where it is further metabolized to flavin-mononucleotide (FMN) and flavin-adenine-dinucleotide (FAD), which are cofactors for flavin-containing proteins.^33^ To determine if a flavin class effect contributed to HAT1 inhibition the enzymatic assay was repeated with other flavonoids including riboflavin, di-methyl-isoalloxazine, FMN, FAD, and riboflavin tetrabutyrate (Fig. 2D). NSC-42186, riboflavin and riboflavin tetrabutyrate all inhibited HAT1 enzymatic activity with low-micromolar IC_50_s of 38, 20, and 17 μM, respectively (Fig. 2E). However, there was no detectable inhibitory activity for di-methyl-alloxazine, FMN, nor FAD. Because riboflavin, FAD and FMN have similar redox potentials^34^ it is unlikely that redox cycling is a common mechanism for HAT1 inhibition with this class of compounds. Furthermore, we were unable to detect free radical generation by NSC-42186 under the above assay conditions. Therefore, certain riboflavin analogs, but not all, can modulate HAT1 acetylation activity and this effect appears to be independent of redox cycling.

### Screening of focused libraries to develop structure-activity relationships

As various riboflavin-derived analogs affected HAT1 enzymatic activity we assessed an expanded collection of similar compounds. Computational structure searches were performed to identify other compounds in the NCI open library with similarity to the isoalloxazine core of NSC-42186. Of these, 30 additional compounds were experimentally characterized with our assay (Fig. 3A, Supp. Fig. 2A). The top hit (NSC-3064) also contained a 7,8-di-methyl-isoalloxazine core but with a different N_10_ appended sidechain comprised of an acetyl-ethyl (Fig. 3B). Also, the fifth best hit (NSC-275266) also contained a 7,8-dimethyl-isoalloxazine core but lacked the amino group at position 4. Therefore, the 7,8-di-substituted-isoalloxazine structure is repeatedly found in compounds with HAT1-inhibitory activity, although the N_10_-appended sidechain can vary.

**Figure 3:**
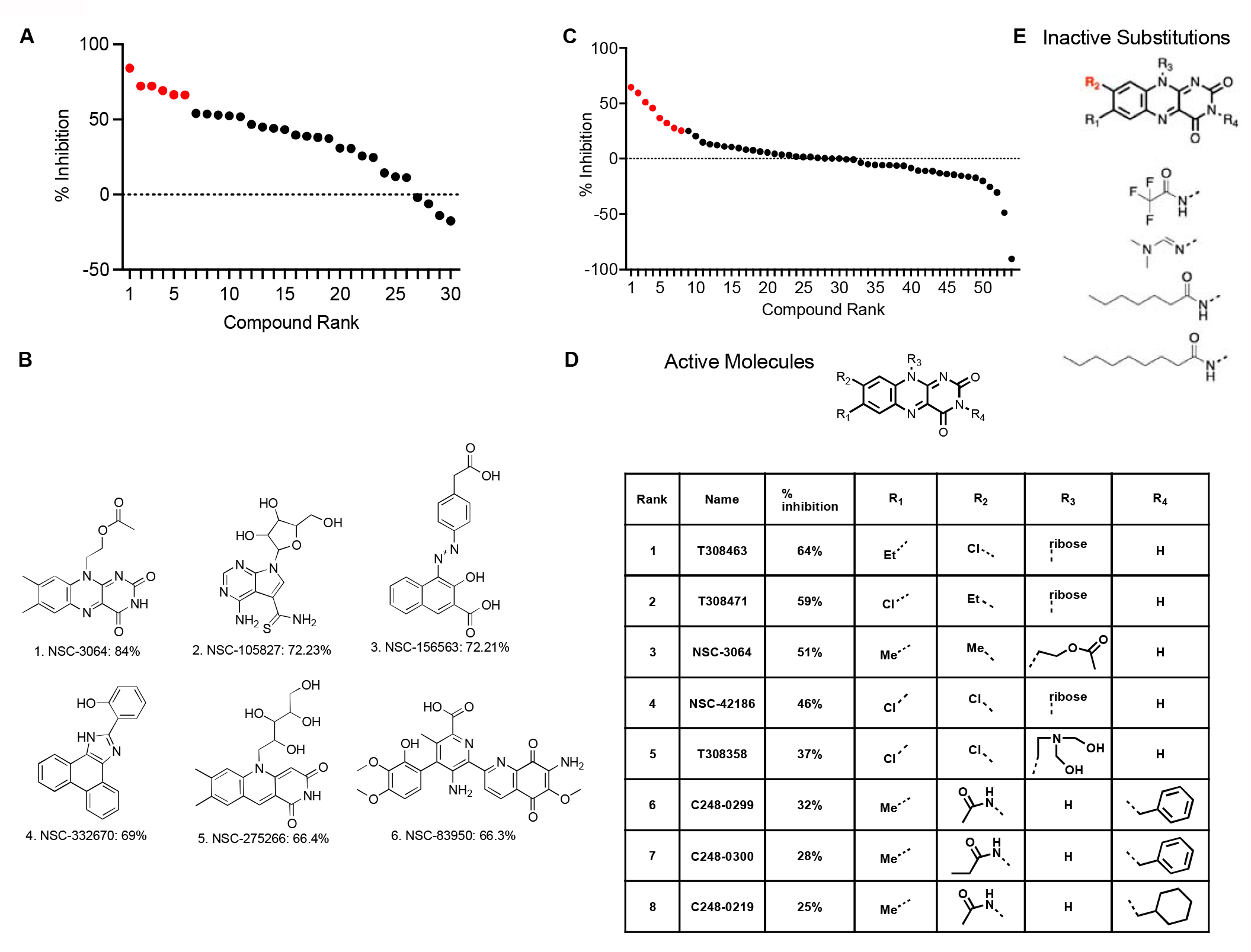
Screening of focused libraries to develop structure-activity relationships. A. Structural similarity screen of the isoalloxazine core was used to select similar multi-cycle cores from the NCI open compounds directory, which were obtained from NCI/DTP and tested for inhibitory properties in the HAT1 acetylation assay. Compounds were screened in duplicate on separate days a mean % inhibition values are plotted (Supp. Fig. 2A). Red indicates the top 6 compounds. B. Chemical structures of the top 6 compounds identified in A. C. A focused library of 54 compounds containing the isoalloxazine core, as well as related structures, was screened by the HAT1 acetylation assay. Compounds were screened in duplicate on separate days and mean % inhibition values are plotted (Supp. Fig. 2B). Red indicates the top 8 compounds. D. Chemical structures of the top 8 compounds from C with greatest inhibition. E. R_2_-groups of compounds from C that were inactive in the assay.

Based on the detection of multiple riboflavin analogs with HAT1 inhibitory properties we next screened a focused library of 54 compounds that all contained a core tri-cyclic ring structure similar to isoalloxazine (Fig. 3C, Supp. Fig. 2B). This library contained molecules with R-group substitutions at four different sites around the isoalloxazine core (Fig. 3D). R-groups from the top 8 most potent compounds are summarized in table format (Fig. 3D). The top two compounds are highly similar with chloro- and ethyl-substitutions at either R_1_ or R_2_ positions and ribityl-groups at position R_3_. This screen also allowed us to identify a number of substitutions that dramatically impaired inhibitory activity, for example, bulky or aliphatic substitutions at R_2_ (Fig. 3E). In contrast, aromatic substitutions at R_4_ were partially tolerated (Fig. 3D). Therefore, this focused library of isoalloxazine derivatives allowed for determination of structural determinants associated with HAT1 inhibitory activity.

### Medicinal chemistry optimization of riboflavin analog 7-chloro-, 8-ethyl-isoalloxazine

Thus far our studies have identified a series of compounds based on the isoalloxazine tricyclic ring system that contains HAT1 inhibitory activity when modified at specific positions. The most potent inhibitors recovered included chloro- or ethyl-substitutions at the 7,8 positions and a sidechain at amino 10. Therefore, we undertook a medicinal chemistry approach to synthesize a library of compounds that incorporated these features. Using a 7-chloro-, 8-ethyl-isoalloxazine core, analogs were synthesized with R-groups at the amino-10 side-chain position. Chemical synthesis of analogs was performed by amination of a nitrosylated di-chloro benzyl ring followed by condensation to yield the isoalloxazine core (see Supp. Fig. 3 and Methods).

Altogether, 73 analogs were synthesized and tested for HAT1 inhibitory activity together with 11 additional compounds with structural similarity (84 compounds total). Of these 84 compounds, 11 (13%) caused at least 50% inhibition of HAT1 activity at 100 μM (Fig. 4A, Supp. Fig. 4). Single-dose inhibition studies identified analog JG-2016 as the most potent HAT1 inhibitor, which caused 69% inhibition at 100 μM (Fig. 4A). This analog has a 1-ethoxy-2-methyl-propane sidechain at the amino-10 position of the isoalloxazine core (Fig. 4B). Dose-response studies showed it to inhibit HAT1 enzymatic activity with IC_50_ 14.8 μM (95% CI: 9.6 – 22.9 μM; Fig. 4C), which is approximately a 2-4 fold improvement over the original hit compound NSC-42186.

**Figure 4:**
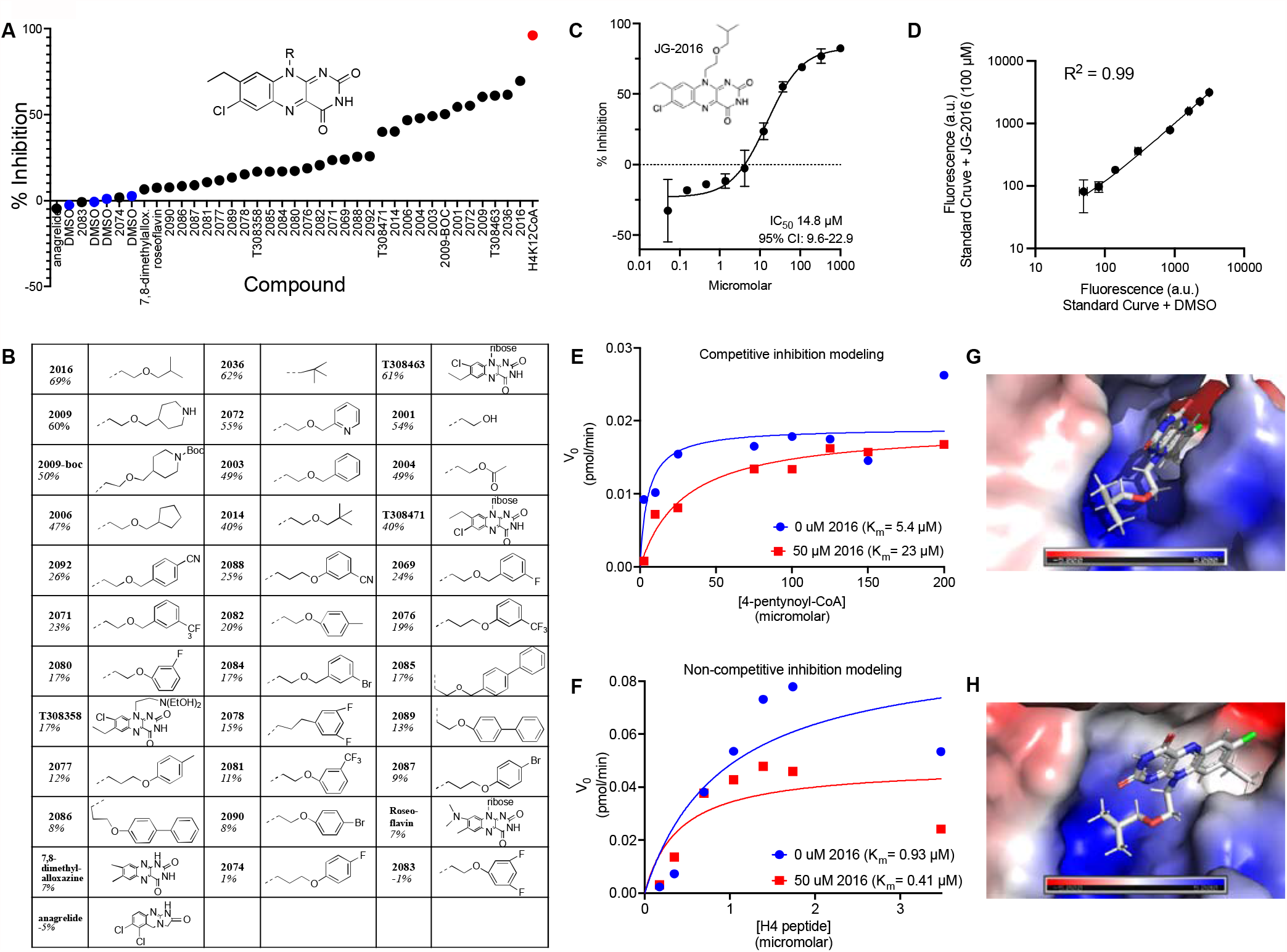
Medicinal chemistry optimization of riboflavin analogs to yield JG-2016. A. Inset shows chemical structure of the core compound used for chemical library generation with R-group at the 10-amino position. Graph shows the mean % inhibition values for each compound performed in duplicate on separate days. See also Supp. Fig. 4. B. Chemical structures of compounds tested in A. Compound name is bolded and below are % inhibition values. Dashed lines indicate bonds joining to the 10-amino position. For compounds that vary from the R-group library the full structure is provided. C. Dose-response of compound JG-2016 in the HAT1 acetylation assay. IC_50_ was calculated by 3 parameter least-squares regression assuming Hill slope -1.0. Mean ± SD of triplicate reactions is plotted. D. Standard curves of positive control H4 peptides were generated in the presence of JG-2016 (100 μM) or DMSO control, followed by binding to neutravidin plates, Cu(I)-catalyzed azide-alkyne cycloaddition of biotin, then streptavidin-HRP binding and amplex red reaction and fluorescence detection (mean ± SD of duplicate samples). E. HAT1 enzyme kinetics for varying concentrations of 4-pentynoyl-CoA were performed in the presence or absence of 50 μM JG-2016 at timepoints 0, 5, 10, 15 minutes of reaction time, quenched in 8 M urea then analyzed jointly by acetylation assay. Datapoints are plotted and curves represent competitive inhibition least squares regression modeling of the data. Michealis-Menton K_m_ were calculated for each dataset. F. HAT1 enzyme kinetics for varying concentrations of H4 peptide were performed in the presence or absence of 50 μM JG-2016 at timepoints 0, 5, 10, 15 minutes of reaction time, quenched in 8 M urea then analyzed jointly by acetylation assay. Datapoints are plotted and curves represent noncompetitive inhibition least squares regression modeling of the data. Michealis-Menton K_m_ were calculated for each dataset. G & H. Poses of JG-2016 docked into the acetyl-CoA cofactor binding pocket. Electrostatic surface potential scale is shown as red (negative) and blue (positive). In G residues 279-281 are removed to improve visualization. See also Supp. Fig. 5.

In addition, this medicinal chemistry approach identified structure-activity relationships. For example, bulky sidechains interfered with enzyme inhibition, most prominently the 1-ethoxymethyl-benzene sidechains substituted at the meta and para positions in compounds 2092, 2088, 2069 and others (Fig. 4B), suggesting a steric effect. In summary, these studies identify active and inactive amino-10 position sidechains with HAT1 enzyme inhibitor properties on an isoalloxazine scaffold.

### Mechanism of HAT1 inhibition by JG-2016

These structure-activity studies nominated the analog JG-2016 for further study as it was the most potent HAT1 inhibitor identified in enzymatic studies. To validate that this compound represented a true enzymatic inhibitor and not an assay-interfering compound, further characterization of assay conditions in the presence or absence of the compound were performed. Standard curves were prepared in the presence of JG-2016 or equivalent amounts of vehicle (DMSO) as a control. The fluorescence activity in the standard curve is generated by titrating a defined amount of biotinylated H4 n-terminal peptide with lysine 5 and 12 substituted for propargylglycines to mimic the HAT1-dependent acylation that occurs in the presence of 4-pentynoyl-CoA. This allowed us to assess the robustness of assay steps in the presence of JG-2016 including binding to neutravidin plates, copper-catalyzed azide-alkyne cycloaddition of biotin-azide, streptavidin-HRP binding and finally amplex red oxidation and fluorescence detection. The standard curves generated in the presence of JG-2016 were indistinguishable from control conditions (Fig. 4D). This indicates that JG-2016 does not interfere with assay conditions, and instead, likely disrupts HAT1-dependent peptide acylation.

Next, enzyme kinetics were performed to further determine the mechanism of action of how JG-2016 inhibited HAT1. Reactions were performed with independent titrations of either the co-factor (4-pentynoyl-CoA) or the substrate (H4 peptide). Curve fitting demonstrated that JG-2016 best fit a model of competitive inhibition with the cofactor (>83% probability compared to noncompetitive model; >89% probability compared to a mixed inhibition model; Fig. 4E). Correspondingly, treatment with JG-2016 best fit a non-competitive inhibition model with the peptide substrate (>99% probability compared to mixed model; >77% probability compared to competitive model; Fig. 4F). Therefore, the JG-2016 compound likely binds HAT1 at the cofactor-binding site and does not compete with the histone substrate.

Using the available human HAT1 crystal structure,^32^ we docked JG-2016 to the cofactor binding site to generate a structural model for inhibition of the enzyme. The flat, rigid isoalloxazine core lay wedged in the pantotheinine-bound canyon with the 8-ethyl substitution in the hydrophobic pocket that accepts the thioester-acetyl moiety of acetyl-CoA (Fig. 4G, H), Supp Fig. 5). The 1-ethoxy-2-methyl-propane amino-substituted sidechain explored the basic cavity typically occupied by the phospho-diester bonded to the adenine nucleotide base of CoA. This model explains aspects of the structure-activity relationship thus far uncovered. For example, the shallow hydrophobic pocket accepting the 8-ethyl substitution would not be capable of accepting longer aliphatic or bulky moieties previously found to be inactive (Fig. 3E). Similarly, bulky substituted aromatic sidechains without hydrogen bond potential at the amino-10 position (Fig. 4B) would be ill-suited in the highly basic pocket. Thus, this structural model recapitulates aspects of the SAR and may be useful to guide further structure-based design.

### Pharmacologic inhibition of HAT1 leads to anti-tumor activity

Development of a HAT1 acetylation assay, coupled to virtual screening and medicinal chemistry prioritized the JG-2016 riboflavin-derived analog for further biological characterization. The triple-negative breast cancer cell line HCC1806 was treated with JG-2016, H4K12-CoA and the riboflavin analog T308463 at varying doses and cell growth was assessed by addition of resazurin, which is reduced to resorufin in cells and emits fluorescence at 590 nm (Fig. 5A). Although H4K12-CoA is a potent enzymatic inhibitor of HAT1 in cell-free assays, it does not cross cell membranes and is susceptible to metabolic degradation and thus had no inhibitory activity against this cell line. In contrast, JG-2016 robustly inhibited cell growth (EC_50_ 10.4 μM; Fig. 5A). Additionally, JG-2016 was a more potent cell growth inhibitor compared to the parent compound T308471 in HCC1937 (triple-negative breast cancer; EC_50_=29.8 μM; Fig. 5B) and A549 (lung cancer; EC_50_=1.9 μM; Fig. 5C) cell lines. The weak inhibitory activity of the T308 compounds with 10-ribityl sidechains mirrors the minimal toxicity of riboflavin which can be tolerated at mega-doses in animals due to urinary excretion.^35^ Thus, the substituted isoalloxazine scaffold derivative JG-2016 has distinct biological properties compared to classical flavonoids.

**Figure 5:**
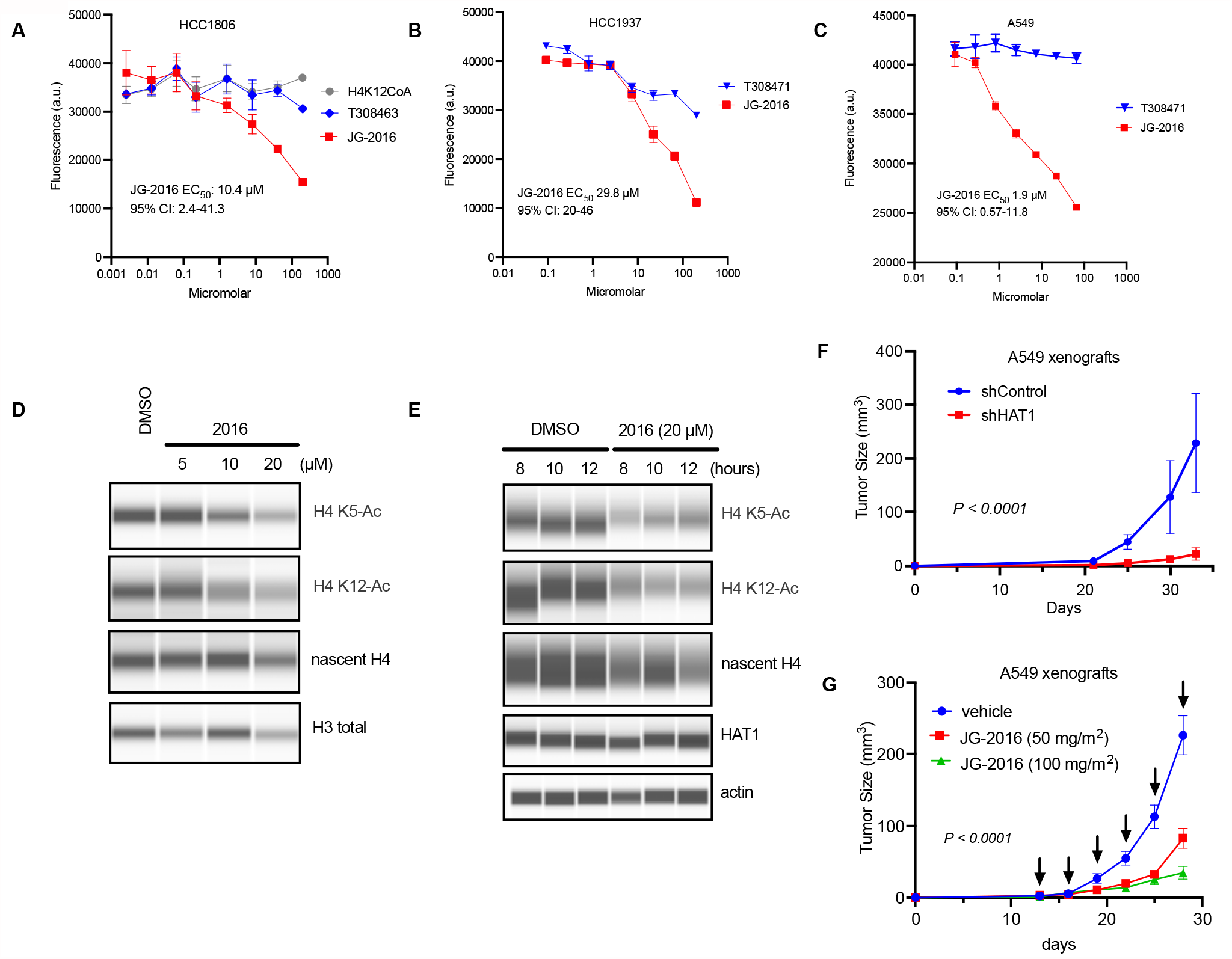
HAT1 inhibition: confirmation of biological activity and anti-tumor effectiveness. A. Dose-response of riboflavin analogs or H4K12-CoA bisubstrate inhibitor in the HCC1806 cell lines with EC_50_ for JG-2016 indicated. B. Dose-response of riboflavin analogs in the HCC1937 cell lines with EC_50_ for JG-2016 indicated. C. Dose-response of riboflavin analogs in the A549 cell line with EC_50_ for JG-2016 indicated. D. hTert-HME1 cells were treated with indicated concentrations of JG-2016 in MEGM media supplemented with 1% dialyzed BSA for 12 hours. Then nascent histone acetylation was assessed by microcapillary immunoassays. E. hTert-HME1 cells were starved of EGF for 16 hours, then pre-treated with JG-2016 for 30 minutes followed by stimulation with EGF for 8, 10, or 12 hours. Then nascent histone acetylation was assessed by microcapillary immunoassays. F. A549 cells were treated with either control lentivirus shRNA or 3 separate lentiviral shRNAs targeting the HAT1 mRNA, then injected into bilateral flanks of NSG mice. N=3 control shRNA mice and 7 HAT1 shRNA mice. * indicates p< 0.0001 by least squares regression and extra sum-of-squares F-test. G. A549 cells were implanted into bilateral flanks of NSG mice, then vehicle control or JG-2016 at doses indicated were injected intraperitoneally every third day starting on day 13 (indicated by arrows). P-value calculated by least squares regression and extra sum-of-squares F-test. N = 5 mice per group.

The EC_50_ for growth inhibition was next assessed in the HCC1806 cell line for 42 flavonoids (selected for a range of HAT1 enzymatic IC_50_s), as well as other isoalloxazine derivatives and control treatments (Supp. Fig. 6A-E). In these experiments, JG-2016 analog was among the most potent inhibitors of cell growth. In contrast, some analogs (eg., roseoflavin) had strong cell growth inhibitory properties but were inactive in the HAT1 enzymatic assay. Among all tested compounds, the correlation of cellular EC_50_ for cell growth versus enzymatic IC_50_ for inhibition demonstrated that JG-2016 had the most optimized combination of these parameters (Supp. Fig. 6F).

Given the properties of JG-2016 as an enzymatic inhibitor of HAT1 and a suppressor of cell growth we next tested if this compound could inhibit HAT1 acetylation *in vivo*. We previously demonstrated that acute suppression of HAT1 protein levels led to decreased H4 lysine 5 and 12 acetylation of nascent histone tetramers, as well as decreased total H4 protein levels in the hTert-HME1 cell line.^2^ When these cells were treated with JG-2016 we observed dose-dependent inhibition of H4 lysine 5 and 12 acetylation by site-specific antibody-based capillary immunoassay (Fig. 5D). Similarly, time-course analysis also showed on-target inhibition as early as 8 hours post-treatment that persisted through 12 hours (Fig. 5E). At some timepoints, decreased H4 protein levels were also observed, consistent with prior studies of the effects of HAT1 depletion.^2^ These experiments indicated that treatment with JG-2016 can impair HAT1 dependent acetylation *in vivo*.

Given the on-target inhibition observed, we next tested whether HAT1 inhibition was sufficient to impair tumor growth in pre-clinical mouse models. We focused on the A549 model because of the low IC_50_ required to impair cell proliferation by JG-2016 treatment (∼1 μM; Fig. 5C). First, the A549 cell line was proven to be HAT1-dependent by depleting HAT1 protein levels. A549 cells treated with shRNAs targeting the HAT1 mRNA evinced suppression of tumor growth in mice compared to cells treated with control shRNAs (p < 0.0001; Fig. 5F). Next, A549 cells were implanted into flanks of mice and allowed to establish for 13 days, then treated with intraperitoneal injections of JG-2016 (Fig. 5G). Dose-finding studies showed that mice could tolerate up to 250 mg/kg or lower as a single intraperitoneal injection with minimal toxicity but signs of distress were observed at doses of 500 mg/kg. Thus, two dose levels of JG-2016 were used: 50 mg/kg and 100 mg/kg, both delivered every third day. These doses are similar to murine dosing of other targeted inhibitors used for cancer.^36-39^ Treatment with JG-2016 at both doses significantly impaired tumor growth compared to vehicle-treated control mice (p < 0.0001; Fig. 5G). A dose-response relationship was observed with the higher dose (100 mg/kg) of JG-2016 causing more profound suppression of tumor growth compared to the lower dose (50 mg/kg). This indicates that HAT1 can be successfully targeted with a small molecule *in vivo* to achieve anti-tumor efficacy.

## DISCUSSION

Multiple reports have suggested that HAT1 may be a cancer therapeutic target based on protein knockdowns or knockouts in various pre-clinical models.^2-5^ These works, which include our own prior studies, motivated the search for small molecules capable of interfering with HAT1 enzymatic activity. We report here the development of a new approach to identify and characterize acetyltransferase activity based on a high-throughput, peptide-based, click-chemistry-enabled enzymatic assay. The advantages of this enzymatic assay include the utilization of the human HAT1/Rbap46 enzyme complex purified from human cells as opposed to a bacterial source. Also, this assay provides a direct readout of enzymatic activity on the peptide substrate without relying on coupled reactions that are prone to nonspecific inhibition. In addition, we have validated its high-throughput characteristics in 96-well plates, which should enable larger chemical screens to be performed. The click chemistry approach based on 4-pentynoyl-CoA as an acetyl-CoA analog should be adaptable to other acetyltransferases as well. Finally, as reaction products are bound prior to functionalization and quantification, it allows for washing to remove potential assay-interfering compounds that commonly cause nonspecific signatures in other assays.^40^

JG-2016 was prioritized based on a workflow that spanned virtual structure-based docking algorithms followed by enzymatic assays, focused library screening, medicinal chemistry and biologic assays. This compound retains the isoalloxazine core common to flavonoids with modifications to the 7,8, and 10 positions that led to significant improvements in enzyme inhibition and cellular growth inhibition. It is notable that its cellular EC_50_ for growth inhibition in the A549 cell line (1.9 μM) is lower than the enzymatic IC_50_ (14 μM). Indeed, prior work has demonstrated that flavonoids are selectively transported into cancer cell lines,^41,42^ indicating that active transport of JG-2016 may play a role in cancer-specific targeting. A family of riboflavin transporters have recently been identified and shown to specifically recognize and transport the isoalloxazine core, rather than the ribityl sidechain.^43-47^ Thus, JG-2016 retains chemical features that may allow for its specific transport into cancer cells. Cancer-specific expression of riboflavin transporters may be a biomarker of sensitivity to this agent. This also raises the possibility of using isoalloxazine as a mechanism to achieve cancer-selective targeting of therapeutic agents, as has been previously demonstrated for another B class vitamin: folate (B9).^48^

HAT1 sits at the intersection of cytoplasmic mitochondrial processes that generate acetyl-CoA and nuclear reactions that consume it to drive transcription of growth programs. HAT1 functions as a cytoplasm-to-nucleus acetyl-shuttle by acetylating nascent histones in the cytoplasm that then become rapidly de-acetylated upon insertion to chromatin leading to a nuclear acetyl pool. This work describes isolation of the first small molecule compounds capable of inhibiting HAT1 enzymatic activity. These compounds may serve as a chemical tools to further our understanding of HAT1 biology, its role in chromatin synthesis and the connection between cellular metabolism, epigenetics and nuclear acetyl flux. JG-2016 or other related analogs may also serve as a chemical scaffolds for more potent HAT1 inhibitors through structure-based design or chemical similarity screens. Finally, this work is the first to suggest that HAT1 may be a therapeutic vulnerability in cancers with an acceptable toxicity profile. Given the importance of HAT1 in the response to EGF stimulation and its role in histone maturation and cell cycle progression, this indicates that further efforts to target HAT1 may be a fruitful for cancer therapy.

## ACKNOWLEDGEMENTS

The authors would like to thank members of the Snyder laboratory for helpful comments and criticisms during the course of this work. We would like to thank Steve Schow for assistance, guidance and oversight of medicinal chemistry synthetic routes and design of chemical libraries. J.J.G. was supported by the Jane Coffin Childs Memorial Fund for Medical Research, a NIH NCI K08 award (1K08CA245024), and a Stanford SPARK Scholars Award. A.M.L was supported by a NIH NIDDK F32 award (F32DK104460) and National Center for Advancing Translational Sciences of the National Institute of Health (UL1TR003142). M.P.S. was supported by grants from the NIH NHGRI (3UM1HG009442) and the NCI (1U2CCA233311). J.J.G., B.S.G., M.P.S., and A.M.L. have filed a patent describing the HAT1 acetylation assay (PCT/US20/29395). The H4K12-CoA bisubstrate inhibitor was a gift from Y. George Zheng at the University of Georgia. Figure 1A was created with BioRender.com. M.P.S. is a cofounder and scientific advisor for Personalis, Qbio, January AI, Filtricine, Protos, Mirvie, NiMo, Oralome, Onza, and Exposomics. He serves on the advisory board of Genapsys.

