## Supplemental Information for "An acetyl-click screening platform identifies a small molecule inhibitor of Histone Acetyltransferase 1 (HAT1) with anti-tumor activity"

### **Supplemental Figures**

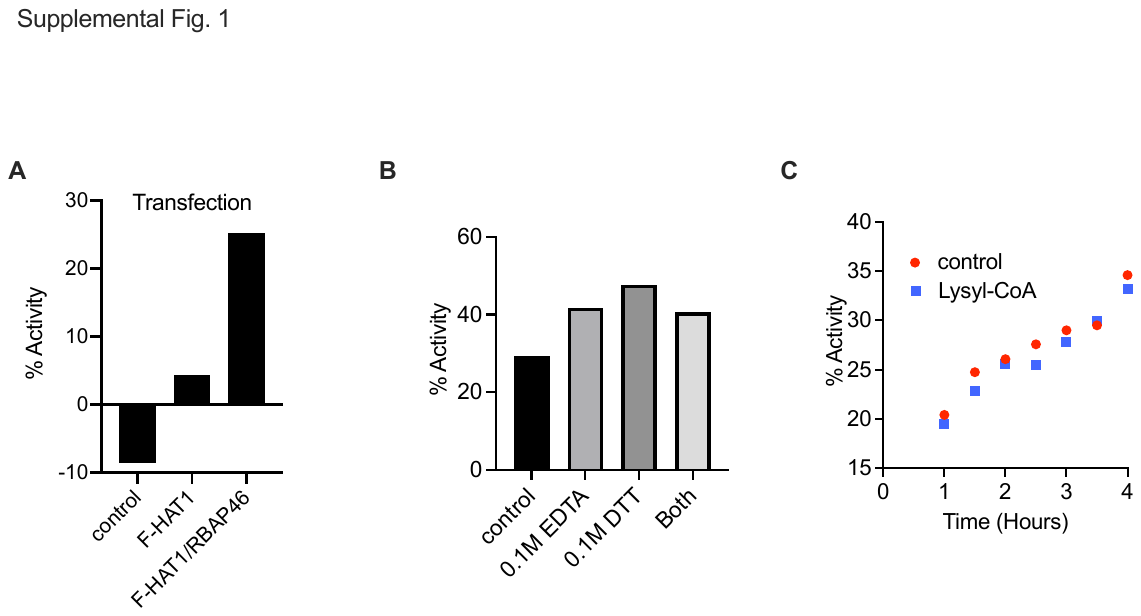

*Supp. Fig. 1: Optimization and characterization of HAT1 acetylation assay. A. 293T cells were transfected with Flag-HAT1 (F-HAT1) alone or F-HAT1 + Rbap46 plasmids, or untransfected (control) then proteins were purified by FLAG immunoprecipitation and HAT1 acetylation assays were performed. B. HAT1 acetylation reactions were performed with 0.1M EDTA, 0.1M DTT or both EDTA + DTT. C. HAT1 acetylation reactions were performed in the presence or absence of Lysyl-CoA (10 μM) over a timecourse of 1-4 hours.*

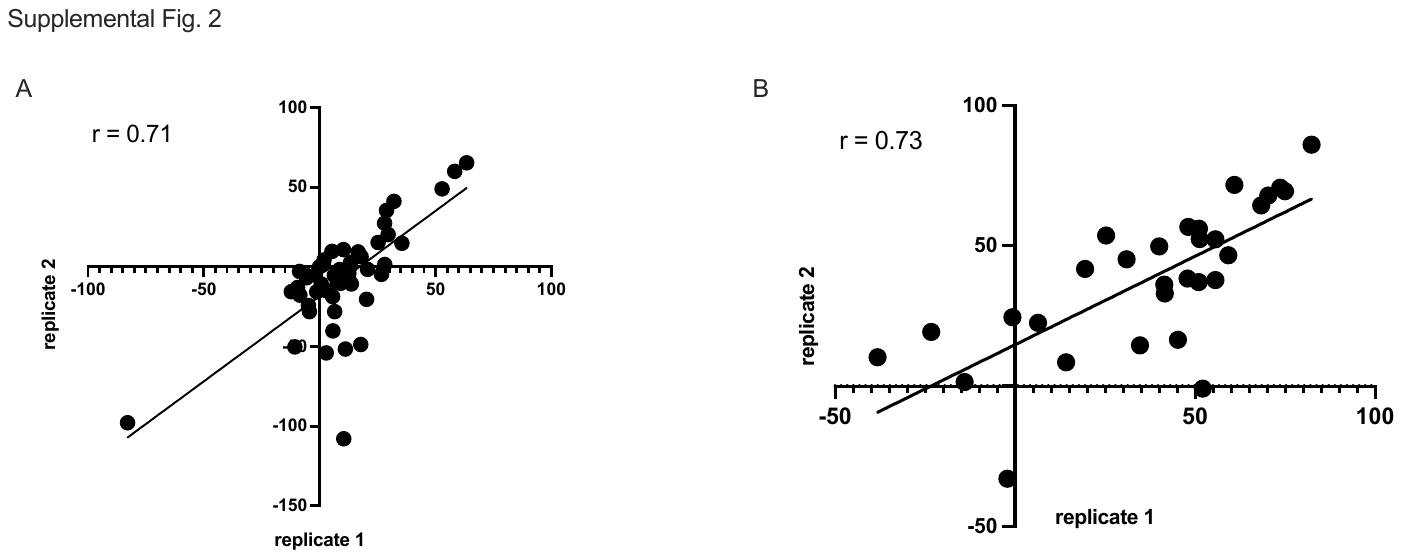

*Supp. Fig. 2: A. Reproducibility scatterplot corresponding to Fig. 3A. B. Reproducibility scatterplot corresponding to Fig. 3C. For both panels linear regression line is plotted and resulting correlation coefficient noted.*

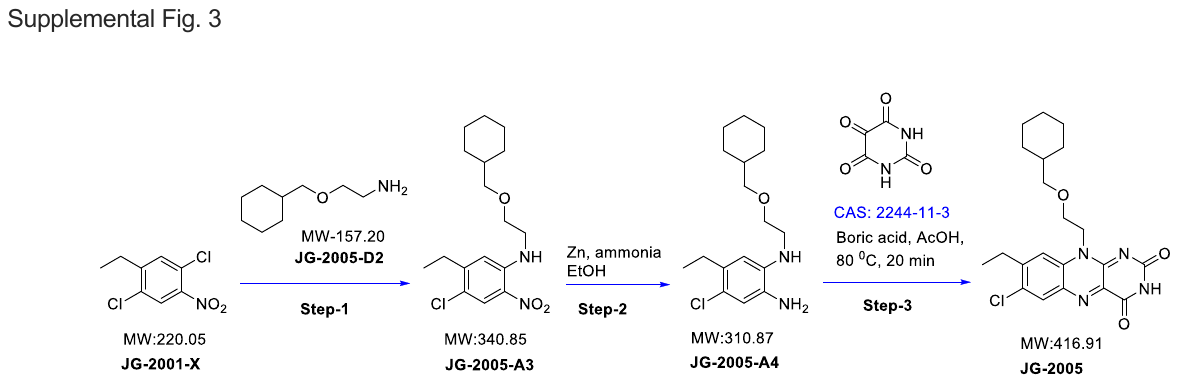

*Supp. Fig. 3: Representative schematic for synthesis of 7-chloro-, 8-ethyl-isoalloxazine core library compounds shown for JG-2005. Additional experimental methods for this synthetic route and other compounds are provided in Supplemental Methods.*

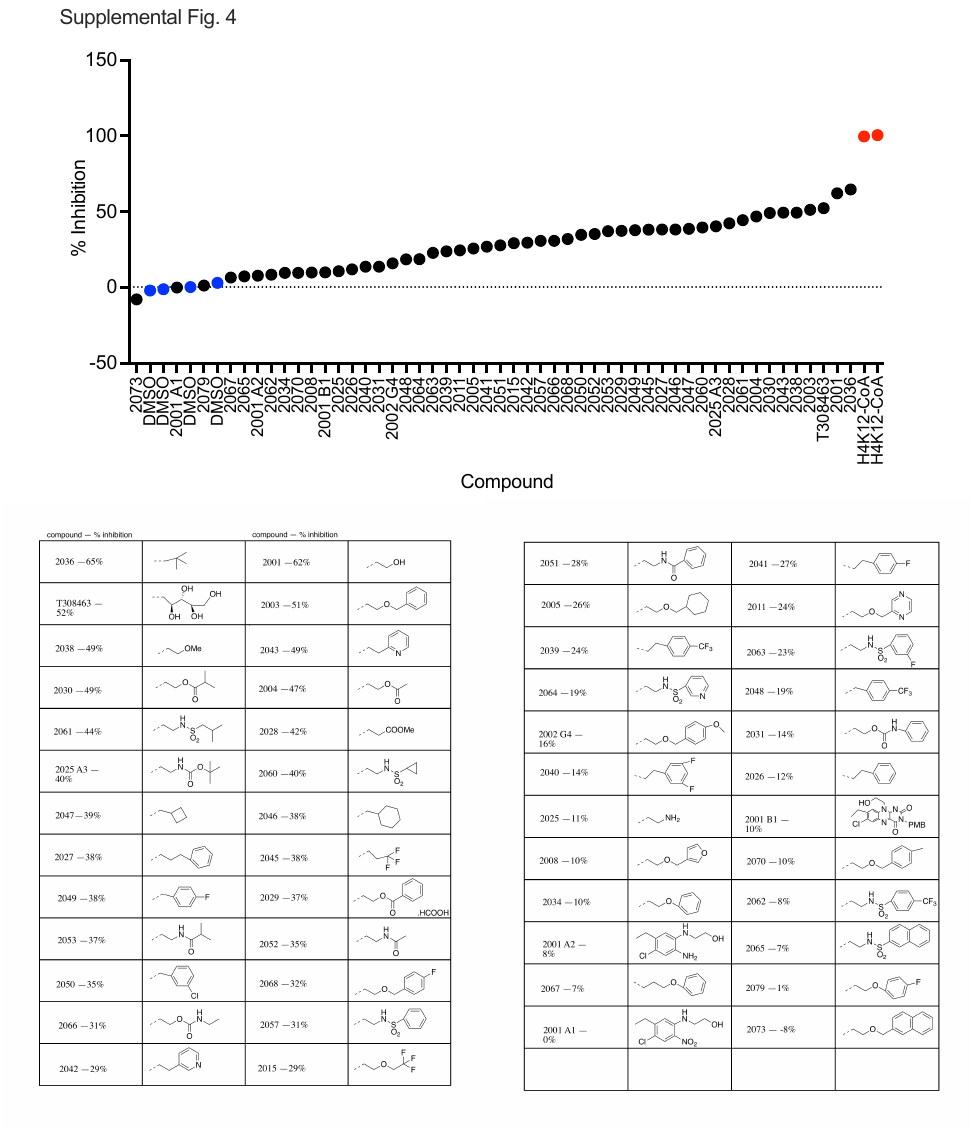

*Supp. Fig. 4: Screening of 7-chloro-, 8-ethyl-isoalloxazine core library. (Top) Assay results of tested compounds (black), negative control DMSO (blue) and positive control bisubstrate inhibitor H4K12-CoA (red). Data is mean of duplicate experiments performed on separate days. (Bottom) Compound name with % inhibition and R-group.*

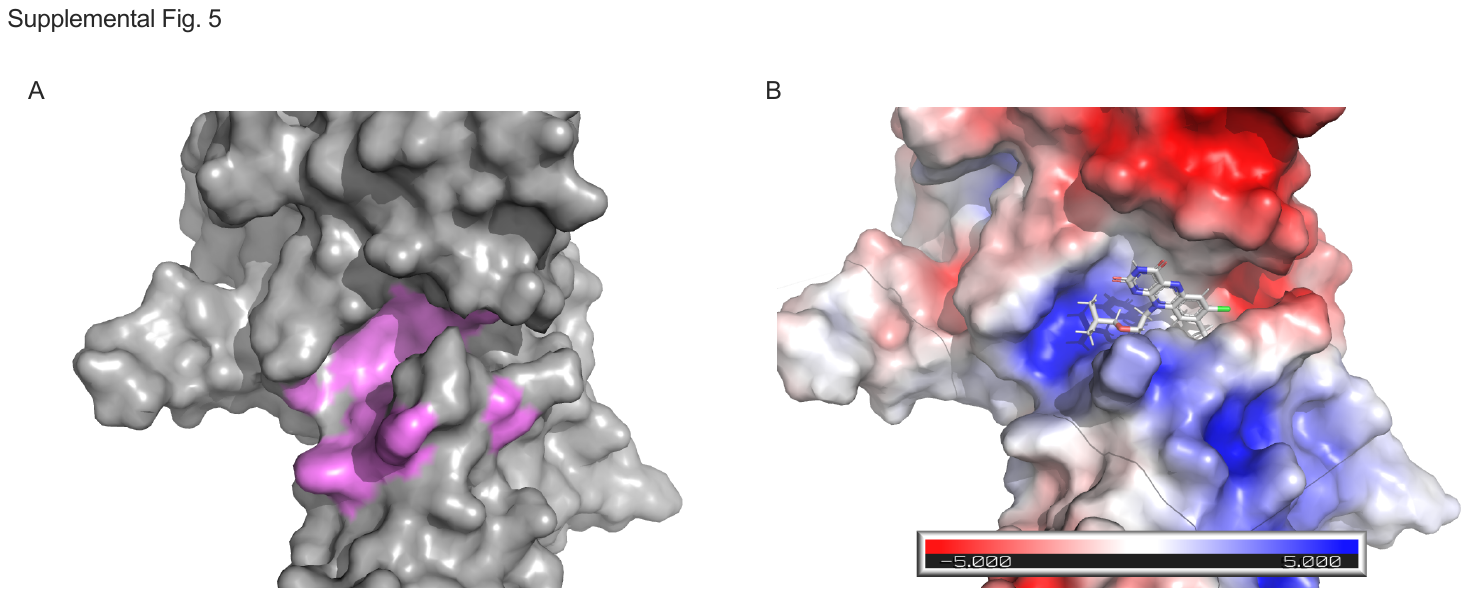

*Supp. Fig. 5: A. Close-up view of the co-factor binding site on HAT1 (2P0W) shaded in violet. B. Comparison view of JG-2016 docked into the HAT1 (2P0W) cofactor-binding site. Electrostatic potential is colored on the protein surface. Residues 279-281 were removed to improve visualization.*

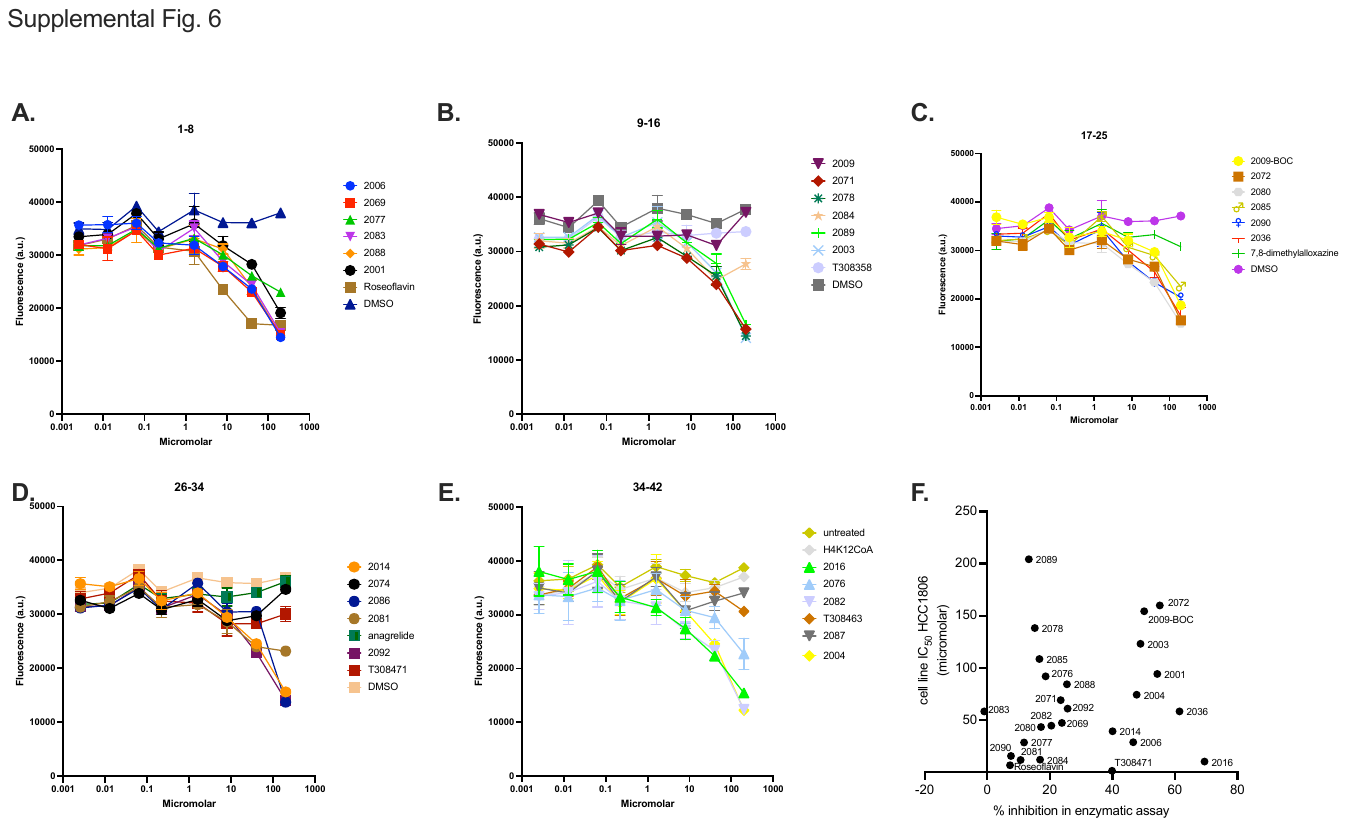

*Supp. Fig. 6: Cell growth inhibition assays of 7-chloro-, 8-ethyl-isoalloxazine core library compounds and related molecules. A-E. HCC1806 cell line was treated with indicated compounds for 48 hours and cell density was measured by CellTiter Blue (fluorescence). F. HCC1806 cell IC_50_ is plotted against corresponding HAT1 enzymatic assay results (% inhibition) at single-dose point (100 μM).*

### Experimental Methods

#### Buffers

- RSB-500: 20 mM Tris/pH7.5, 500 mM NaCl, 25 mM MgCl_2_ – keep at 4degC

- 500 mL = 10 mL 1M Tris, 50 mL 5M NaCl, 1.25 mL 1M MgCl_2_ – H2O to 500 mL

- RSB-100 : 20 mM Tris/pH7.5, 100 mM, NaCl 25 mM, 25 mM MgCl_2_ – keep at 4degC

- 500 mL = 10 mL 1M Tris, 10 mL 5M NaCl, 1.25 mL 1M MgCl_2_ – H2O to 500 mL

- Elution Buffer: (25 mM HEPES pH 8.0, 100mM NaCl, 0.01% Triton X)

- 20x HAT1 Buffer (1M Tris pH 8.5, 0.1% NP40 [IGEPAL])

- PBST: PBS with 0.1% Tween-20

Expression and purification of human HAT1 complex Full-length human HAT1 and Rbap46 were independently cloned into the pHEK293 vector (Takara Bio) with Gibson assembly. HAT1 was appended with a C-terminal FLAG tag, Rbap46 was unmodified. The pHEK293 vector was digested with XbaI then amplified with Phusion polymerase (Thermofisher) with primers (CGAGCCCGGGGAGGCTTAAG, CCTGCAGGCATGCAAGCTTG), then gel purified. HAT and Rbap46 were ordered as gBlocks (IDT) with ~20bp ends overlapping sequence with the pHEK293 vector. Gibson assembly was performed with NEBuilder HiFi DNA Assembly (NEB) and propagated in DH5α cells. 293F cells (Thermo-Fisher) were grown in suspension culture in a humified incubator with 8% CO_2_ at 37degC with constant shaking at 120 RPM. 293F were seeded to a density of 5E5 cells/ml in 300 ml culture volume in a 1L baffle-free plastic Erlenmeyer flask with vented caps. The next day they transfection mix was prepared with 300 μg of pHEK293-HAT1-FLAG and 300 μg of pHEK293-Rbap46 in 30 mls of PBS with 1.2 mL of PEI (0.5 mg/ml), incubated for 15 minutes, then added to 300 mL culture of 293F. After 48 hours, cells were treated with 12.5 μΜ forskolin for 30 minutes at 37degC then collected by centrifugation and snap frozen. Cell pellets were lysed in 40 mL of RSB-500 buffer with 0.1% triton-X-100 with Complete protease inhibitors (Roche). Lysate was sonicated at 10% amplitude for 10 seconds x 3, then centrifuged at 10K RPM for 10 minutes. Supernatant was collected and immunoprecipitated with FLAG M2 agarose (400 μL per 10 mL of extract) for 2 hours, then washed extensively in lysis buffer, then once in RSB-100 buffer + 0.1% triton-X-100. HAT enzyme was eluted with FLAG peptide diluted to 0.5 mg/mL in EB (1.5 mL) for at least one hour at 4degC. Supernatant was collected and combined, washed twice with 15 mL EB and concentrated by centrifugation in Amicon 10K cutoff filters and resuspended in 6 mL EB per original 300 mL culture volume. Agilent protein 230 chip was run to quantitate protein concentration, then snap frozen in 200 μL aliquots and stored at -80degC. HAT1 acetylation assays were performed to validate enzyme activity and then enzyme diluted in EB to yield approximately 1500 fluorescence units (25% full activity on standard curve) and re-frozen. Typically, this was a 1:40 dilution.

Virtual drug screening workflow, docking, similarity searches, visualization Schrodinger software (2018-3) was utilized to prepare the HAT1 crystal structure 2P0W for serving as a target model for virtual screening. The cognate ligand (acetyl-CoA) was used to generate the grid for the virtual screen. SDF file (NCI_Open_2012-05-01.sdf.gz) of the NCI/DTP open compounds was downloaded from <https://cactus.nci.nih.gov/download/nci/>. Using the Schrodinger Virtual Screening Workflow pipeline, the ligands were prepared by desalting, removing duplicates and generating conformers (max 32) at pH 7.0-7.2 using EPIK algorithm. The ligands were docked into the receptor grid in 3 successive steps of increasing precision levels (HTVS, SP, XP) with Schrodinger Glide program. The resulting docked poses were used to estimate the binding affinity of the ligand using MMGBSA method as implemented in Schrodinger. The ligands that had favorable Glide XPScore (< 10.5) and MMGBSA DGBIND (< -60) were prioritized for testing in the experimental assay. The initial hit compound (NSC-42186) was used as a template to find similar compounds from the NCI database using Schrodinger shape screen. The physiochemical properties of the resulting compounds were calculated using qikprop (Schrodinger) then filtered for predicted water solubility (QPlogS < -1.0 & > -5.7) and reactive functional groups (#rtvFG = 0). The python input file for Schrodinger virtual screening workflow has been deposited online: <https://github.com/quantumdolphin/enzyme-inhibition-assay>. For visualization docking was performed with mcule centered on the methionine 241 in the cofactor binding site and visualized in pymol. Electrostatic surface potential was calculated in pymol with the APBS plugin.

HAT1 acetylation assay Acetylation reactions were assembled from the following components: histone H4 peptide (1-23-GGK-biotin; Anaspec #AS65097) resuspended in DMSO to 0.1 mg/mL [34.8 μM], HAT1 enzyme pre-diluted in EB, 20x buffer, 2 mM DTT, 4-pentynoyl-CoA dissolved in water to 1 mg/mL [1 mM]. A 20 μL reaction comprised 10 μL of enzyme, 1 μL of H4 peptide, 1 μL of 20x buffer, 1 μL of DTT pre-mixed and aliquoted to wells of a 96-well PCR plate on ice. Then 1 μL of DMSO or test compound dissolved in DMSO was added per well and mixed by gentle pipetting and allowed to incubate for 10 minutes on ice. Then 2 μL of 4-pentynoyl-CoA and 4 μL of water were pre-mixed and added together to wells, which were then gently mixed by pipetting, centrifuged briefly at 2500 RPM to collect contents, and incubated at 37degC for 1 hour. Contents were then either directly processed for reaction products or quenched with 20 μL of 8M urea and stored at -20degC until processing. Reaction contents were added to BSA pre-blocked black Neutravidin coated 96-well plates (Fisher # 15217) containing 80 μL of PBST per well and bound with gentle orbital shaking for 1 hour at room temperature. Wells were washed 200 μL PBST 3x 15 strokes (180 μL stroke volume) with a Hydra96 (Art Robbins Instruments). Click reactions were assembled as follows:

- During Peptide binding prepare click reagents fresh
  - 100 mM THPTA ligand in aqueous buffer or water (4.35 mg/ 100 uL)
  - 20 mM CuSO4 in water (4.99mg/mL)
  - 300 mM sodium ascorbate in water (29.7 mg/500 uL)
  - 2.5 mM Biotin Azide (stock is 2.5 mM)
- One click reaction contains (scale appropriately to the number of reactions needed):
  - 140 uL PBS
  - 10 uL of THPTA
  - 10 uL of CuSO4
  - 10 uL of Na Ascorbate
  - 20 uL of Biotin Azide

Click reagents were dispensed to each well and incubated at 37degC for 1 hour, then plate washed 3x as before. Dilute Strept-HRP (CST#3999S @ 0.224 mg/m) 1:10 in Streptavidin (0.224 mg/ml; EMD Millipore #189730), then further dilute this mix 1:1000 in PBST. Add 100 uL per well then incubate at room temp for 1 hour with gentle orbital shaking. Wash 3x with PBST, then add amplex red detection reagents:

- 4.45 mL NaHPO4 buffer (1x) (50 mM final Concentration)
  - 50 uL Amplex Red (20 mM diluted in DMSO)
  - 500 uL of dilution H2O2
    - Dilution of H2O2 (30% stock)
      a. 3% H2O2 - Dilute 3 uL of stock H2O2 in 27 uL of 1x NaHPO4 buffer
      b. Add 22.7 uL of that 3% H2O2 to 977 uL of 1x NaHPO4 (this is the H2O2 to add to mix)

Add 100 uL of amplex red reaction mix per well, incubate at room temp for 30 minutes protected from light then read fluorescence excitation/emission 571/585 nm. Calculate % inhibition according to the following formula:$\frac{D-X}{D-BG}*100$, where *D* is the fluorescence value of control reactions treated with DMSO only, *X* is value of reactions treated with test compounds, and *BG* is value of background wells (no enzyme added). Least squares regression was used to fit dose-response and competitive versus non-competitive inhibition curves. Modes of competition were compared by Akaike’s Information Criterion.

HAT1 acetylation standard curve A positive control H4 N-terminal peptide was synthesized (Genscript) with sequence:

SCRG[Pra]GGKGLG[Pra]GGAKRHRKVLRGG[Lys(Biotin)], where [Pra] denotes Propargylglycine. Peptide was resuspended at 0.1 mg/mL in DMSO, then mixed with H4 N-terminal peptide (Anaspec #AS65097) to create a standard curve. These mixtures were bound to neutravidin plates, functionalized by click chemistry, bound with streptavidin-HRP and reacted with amplex red as described above for the HAT1 acetylation assay. LoD was calculated as the assay baseline (negative controls) plus 3x the standard deviation of the baseline measurements. LoQ was calculated as the assay baseline plus 10x the standard deviation of the baseline measurements.

Cell culture, drug treatments, immunoassays hTert-HME1, HCC1806, HCC1937, A549 were maintained in a humified incubator at 37degC with 5% CO_2_. HCC1806 and HCC1937 were grown in RPMI with 10% FBS and 1% penicillin-streptomycin. A549 was grown in F-12 media with 10% FBS and 1% penicillin-streptomycin. hTert-HME1 was grown in Mammary Epithelial Growth Media (PromoCell). Drug treatments were performed in 96-well plates. Cells were seeded at 5000 cells per well, then allowed to attach overnight. Drug dilutions were made in 96-well plates then transferred to plate containing cells and incubated for 48-72 hours. Cell density was then determined by CellTiter-Blue (Promega). For capillary immunoassays hTert-HME1 5E5 cells were seeded in 10 cm plates and cultured for 48 hours, then cells were washed 2x in PBS and EGF-free MEGM was added overnight. The next day, cells were treated in EGF-free media with 1% dialyzed BSA + drug for 30 minutes, then EGF was added to the plate and cells were cultured for 8, 10 and 12 hours before harvesting. Microcapillary immunoassays were performed on a SimpleWestern Wes instrument using H4K12-Ac antibody (Abcam ab46983) at 1:25 dilution, H4K5-Ac antibody (Millipore 07-327) at 1:50 dilution, HAT1 (Abcam ab194296) at 1:50 dilution, nascent H4 (Abcam ab7311) at 1:10 dilution, nascent H3 (Abcam ab18521) at 1:50 dilution, actin (Thermo-Fisher MA5-15452) at 1:100 dilution.

Mouse models All mouse experiments were conducted with prior approval of the Administrative Panel on Laboratory Animal Care. A549 cells were infected with 3 lentiviral shRNAs targeting the human HAT1 mRNA (Origene # TL312517). Six-to-eight-week-old NSG female mice (Jackson Labs #005557) were shaved, then 500,000 A549 tumor cells were injected into bilateral flanks and tumor growth was assessed by tri-dimensional tumor measurements to yield tumor volumes. For drug treatments, analog 2016 was resuspended in a 1:1 mixture of PEG-400 and PBS + 1% dialyzed BSA-0.2 μm-filtered (MilliporeSigma #12-660-910GM). Intraperitoneal injections (200 μL) were performed with a 21-G needle. Tumor measurements were performed with digital calipers.

Chemicals and synthesis Riboflavin analogs from Figure 5C were purchased from Chemdiv. For chemical synthesis of 7-chloro-, 8-ethyl-isoalloxazine compound library, three general schemes were pursued. In the first, R-group amine side chain was either synthesized or purchased and used for nucleophilic addition of the common di-chloro-nitrosyl-ethyl-benzene (JG-2001-X), followed by reduction of the nitrosyl group to amine and then condensation with alloxan to generate the isoalloxazine core. In a second synthetic route (eg. for JG-2031 and JG-2029), the precursors JG-2001 or the PMB-protected analog JG-2001-B1 underwent either (1) esterification of free alcohol in the presence of triethylamine or (2) EDC, HCl + HOBt coupling in the presence of DMF and DMAP. In the third route, the free amine sidechain of compound JG-2025 was coupled to carboxylic acid sidechains in the presence of HATU, DIPEA and DMF (eg. to make JG-2051). Alternatively, JG-2025 was coupled to sulfonyl chloride containing sidechains in the presence of base (TEA or K_2_CO_3_) and DMF (eg. to make JG-2060). See Supplemental Material for detailed reaction conditions for all routes. Compounds were purified by chromatography and validated by proton NMR and mass spectrometry.

NMR, Mass spectrometry, Compound Analysis ^1^H-NMR spectra were measured on a Bruker 400 MHz spectrometer (i- probe 5mm with Topspin 3.2. Software). All 13C -NMR were recorded at 100 MHz. All purified products were determined to be ≥95% pure unless otherwise noted.

LCMS or mass analysis was performed using either of the following LCMS machine.

(a) Waters Acquity Ultra performance LC equipped with PDA and attached with QDA detector with a Waters X-bridge C18, 50*2.1 mm, 2.5-micron column using a binary solvent system [A: 0.1% FA in Water, B= 0.1% FA in H2O: ACN (10:90)].

(b) Water Acquity UPLC- H Class equipped with PDA and Acquity SQ detector with a Waters X-bridge C18, 50*2.1 mm, 2.5-micron column using a Quaternary solvent system [A: 0.1% FA in Water, B= 0.1% FA in H2O: ACN (10:90)].

(c) Waters Acquity Ultraperfomance LC connected with PDA and equipped with SQ detector with a Waters X-bridge C18, 50*4.6 mm, 3.5-micron column using a Binary solvent system [A= 5mM Ammonium Bicarbonate in H2O and B=ACN].

Mass analysis was performed on a Waters Acquity Ultraperfomance LC equipped with SQ detector using a Binary solvent system [A= 5mM Ammonium Acetate and 0.1 % Formic acid in H2O and B= Methanol].

### Primers and DNA sequences

JJG573: Flag-HAT1 gBlock for pHEK293 Ultra Expression I (overlap for Gibson)

5’ – CTTAAGCCTCCCCGGGCTCGGCCAGATATGGCGGGATTTGGTGCTATGGAGAAATTTTTGGTAGAATATAAGAGTGCAGTGGAGAAGAAACTGGCAGAGTACAAATGTAACACCAACACAGCAATTGAACTAAAATTAGTTCGTTTTCCTGAAGATCTTGAAAATGACATTAGAACTTTCTTTCCTGAGTATACCCATCAACTCTTTGGGGATGATGAAACTGCTTTTGGTTACAAGGGTCTAAAGATCCTGTTATACTATATTGCTGGTAGCCTGTCAACAATGTTCCGTGTTGAATATGCATCTAAAGTTGATGAGAACTTTGACTGTGTAGAGGCAGATGATGTTGAGGGCAAAATTAGACAAATCATTCCACCTGGATTTTGCACAAACACGAATGATTTCCTTTCTTTACTGGAAAAGGAAGTTGATTTCAAGCCATTCGGAACCTTACTTCATACCTACTCAGTTCTCAGTCCAACAGGAGGAGAAAACTTTACCTTTCAGATATATAAGGCTGACATGACATGTAGAGGCTTTCGAGAATATCATGAAAGGCTTCAGACCTTTTTGATGTGGTTTATTGAAACTGCTAGCTTTATTGACGTGGATGATGAAAGATGGCACTACTTTCTAGTATTTGAGAAGTATAATAAGGATGGAGCTACGCTCTTTGCGACCGTAGGCTACATGACAGTCTATAATTACTATGTGTACCCAGACAAAACCCGGCCACGTGTAAGTCAGATGCTGATTTTGACTCCATTTCAAGGTCAAGGCCATGGTGCTCAACTTCTTGAAACAGTTCATAGATACTACACTGAATTTCCTACAGTTCTTGATATTACAGCGGAAGATCCATCCAAAAGCTATGTGAAATTACGAGACTTTGTGCTTGTGAAGCTTTGTCAAGATTTGCCCTGTTTTTCCCGGGAAAAATTAATGCAAGGATTCAATGAAGATATGGCGATAGAGGCACAACAGAAGTTCAAAATAAATAAGCAACACGCTAGAAGGGTTTATGAAATTCTTCGACTACTGGTAACTGACATGAGTGATGCCGAACAATACAGAAGCTACAGACTGGATATTAAAAGAAGACTAATTAGCCCATATAAGAAAAAGCAGAGAGATCTTGCTAAGATGAGAAAATGTCTCAGACCAGAAGAACTGACAAACCAGATGAACCAAATAGAAATAAGCATGCAACATGAACAGCTGGAAGAGAGTTTTCAGGAACTAGTGGAAGATTACCGGCGTGTTATTGAACGACTTGCTCAAGAGACGCGTACGCGGCCGCTCGAGCAGAAACTCATCTCAGAAGAGGATCTGGCAGCAAATGATATCCTGGATTACAAGGATGACGACGATAAGGTTTAACCTGCAGGCATGCAAGCTTG

JJG574: Rbap46 gBlock for pHEK293 Ultra Expression I (overlap for Gibson)

5’ – CTTAAGCCTCCCCGGGCTCG**ATG**gctgccgaagcaggagtcgtgggagctggagcttctcctgatggagattggagagaccaggcctgtgggcttctgctacacgtgcatttgtcttcccgactgggtcgcgcagcccctgtacgtacaggtcgtcatcttagaacagtgtttgaagatactgtggaggagcgtgtcatcaatgaagaatataaaatctggaagaagaatacaccgtttctatatgacctggttatgacccatgctcttcagtggcccagtcttaccgttcagtggcttcctgaagtgactaaacctgaaggaaaagattatgcccttcattggctagtgctggggactcatacgtctgatgagcagaatcatctggtggttgctcgagtacatattcccaatgatgatgcacagtttgatgcttcccattgtgacagtgacaagggtgaatttggtggctttggttctgtaacaggaaaaattgaatgtgaaattaaaatcaatcacgaaggagaagtaaaccgtgctcgttacatgccgcagaatcctcacatcattgctacaaaaacaccatcttctgatgtgttggtttttgactatacaaaacaccctgctaaaccagacccaagtggagaatgtaatcctgatctcagattaagaggtcaccagaaggaaggctatggtctctcctggaattcaaatttgagtggacatctcctaagtgcatctgatgaccatactgtttgtctgtgggatataaacgcaggaccaaaagaaggcaaaattgtggatgctaaagccatctttactggccactcagctgttgtagaggatgtggcctggcacctgctgcacgagtcattgtttggatctgttgctgatgatcagaaacttatgatatgggacaccaggtccaataccacctccaagccgagtcacttggtggatgcgcacactgccgaagtcaactgcctctcattcaatccctacagcgaatttattctagccaccggctctgcggataagaccgtagctttatgggatctgcgtaacttaaaattaaaactccataccttcgaatctcataaagatgaaattttccaggtccactggtctccacataatgaaactattctggcttcaagtggtactgaccgccgcctgaatgtgtgggatttaagtaaaattggggaagaacaatcagcagaagatgcagaagatgggcctccagaactcctgtttattcatggaggacacactgctaagatttcagattttagctggaaccccaatgagccttgggtcatttgctcagtgtctgaggataacatcatgcagatatggcaaatggctgaaaatatttacaatgatgaagagtcagatgtcacgacatccgaactggagggacaaggatct**TAA**CCTGCAGGCATGCAAGCTTG

JJG575: Upstream for Gibson PCR to pHEK293 Ultra Expression I (Xba I digest before PCR)

5’-CGAGCCCGGGGAGGCTTAAG

JJG576: Downstream for Gibson PCR to pHEK293 Ultra Expression I (Xba I digest before PCR)

5’- CCTGCAGGCATGCAAGCTTG

### Preparation of 7-chloro-, 8-ethyl-isoalloxazine core library compounds

#### Synthesis of intermediate JG-2001-X

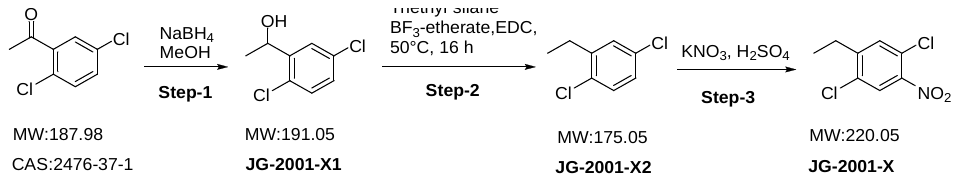

**Step-1: Synthesis of 1-(2,5-dichlorophenyl)ethan-1-ol**

**Batch no. STL7-A-588-JG-2001-X1-163-b**

To a solution of 1-(2,5-dichlorophenyl)ethan-1-one (50.0 g, 264.4 mmol) in MeOH (500 mL), sodium borohydride (15.0g, 396.7 mmol) was added at -10°C and reaction mixture was allowed to stir at 0°C for 1h. After completion of reaction as indicated by TLC, the reaction mixture was concentrated carefully and crude was diluted with water and EtOAc. The reaction mixture was partitioned between water (1000 mL) and EtOAc (1000 mL×2). The organic layer was separated and washed with brine (2 x 1000 mL). The combined organic layer was evaporated under vacuo to obtain title compound as off-white solid (54.0g, 77.63 %). 1H NMR: (400MHz, DMSO) δ 7.59-7.58 (t, J=4Hz, 1H), δ 7.44-7.42 (dd, J= 8Hz, 1H), δ 7.35-7.32 (t, J= 12Hz, 1H), δ 5.55-5.54 (d, 1H), δ 4.99-4.95 (q, 1H), δ 1.31-1.30 (d, J= 4Hz, 3H).

**Step-2:** **Synthesis of 1,4-dichloro-2-ethylbenzene**

**Batch no. STL7-A-588-JG-2001-A2-167-C**

To a solution of 1-(2,5-dichlorophenyl)ethan-1-ol (**JG-2001-X1**) (54.0g, 284.2 mmol) in ethylene dichloride at 0°C, BF3-etherate (67.1 mL, 539.9 mmol) and triethylsilane (92.0 mL, 568.4 mmol) were added at 0°C and stirred for 16 hr at 50°C. After completion of reaction as indicated by TLC, the reaction mixture was cooled at 0°C and quenched with sat. NaHCO3 (1500 ml). The reaction mixture was partitioned between EtOAc (1500mL × 2) and the organic layer was washed with brine solution (2 x 1500 mL). The combined organic layer was dried over Na_2_SO4 and evaporated to obtain title compound as pale-yellow oily liquid (45.0g, quantitative). 1H NMR: (400Mz, DMSO). δ 7.50-7.43 (m, 2H), δ 7.31-7.28 (dd, J= 12Hz, 1H), δ 2.72-2.66 (q, J= 8Hz, 2H), δ 1.24-1.14 (t, 3H).

**Step-3:** **Synthesis of 1,4-dichloro-2-ethyl-5-nitrobenzene**

**Batch no. STL7-A-588-JG-2001-X-178-c**

To a solution of 1,4-dichloro-2-ethylbenzene (**JG-2001-X2**) (45.0 g, 258.6 mmol) in H2SO4 (450 mL) at 0 °C, KNO3 (26.1g, 258.6mmole) was added at 0°C and stirred for 30 min. After completion of reaction as indicated by TLC, the reaction mixture was quenched with ice water (1000 mL) slowly and stirred for 20 min. The product was extracted with EtOAc (500 ml×2) and washed with brine solution (2 x 500 mL). The combined organic layer was dried over Na2SO4 and evaporated under vacuum. The crude product was purified by column chromatography (0-15 % EtOAc/hexane) to obtain title compound as pale-yellow oil (35.0g, 56.89 %). 1HNMR: (400Mz, CDCl_3_) δ 1.20-1.16 (t, 3H), 2.792-2.735 (q, 2H), 7.799 (s, 1H), 8.23 (s, 1H).

#### Synthesis of JG-2001

**7-chloro-8-ethyl-10-(2-hydroxyethyl)benzo[g]pteridine-2,4(3H,10H)-dione**

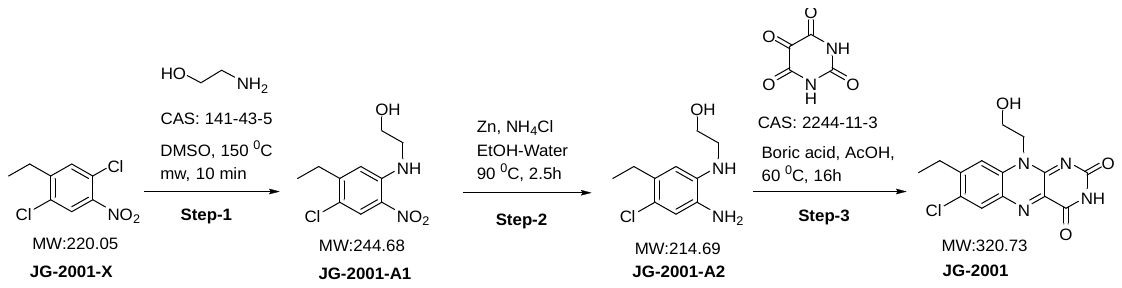

**Step-1: Synthesis of 2-((4-chloro-5-ethyl-2-nitrophenyl)amino)ethan-1-ol**

**Batch ID: STL-7-A-525-JG-2001-A1-028-A**

To a solution of 1,4-dichloro-2-ethyl-5-nitrobenzene (**JG-2001-X)** (0.10 g, 0.456 mmol, 1.0 eq) in DMSO (1.5 mL)**)** 2-aminoethan-1-ol **(CAS: 141-43-5)** (0.064 g, 1.05 mmol, 5.0 eq) was added and the reaction mixture was heated at 150 °C under microwave irradiation for 10 min. After completion of reaction as indicated by TLC, the reaction mixture was poured into ice cold water (50 mL) and extracted with ethyl acetate (3 x 50 mL). The combined organic layer was dried over anhydrous sodium sulphate and concentrated under vacuum. The obtained crude product was further purified by column chromatography (35 % ethyl acetate/hexane) to obtain title compound as orange solid (0.085g, 93.27 %, 76.44%) LCMS m/z 245.0 & 247.0 (M & M+2)

**Step-2: Synthesis of 2-((2-amino-4-chloro-5-ethylphenyl)amino)ethan-1-ol**

**Batch ID: STL-7-A-525-JG-2001-A2-027-A**

To a solution of 2-((4-chloro-5-ethyl-2-nitrophenyl)amino)ethan-1-ol (**JG-2001-A1)** (0.40g, 1.63 mmol, 1.0 eq) in EtOH: water (9:1 mL), Zn Dust (0.86g, 13.1mmol, 8.0 eq) and NH_4_Cl (0.70g, 13.1 mmol, 8.0 eq) were added and stirred at 90 °C for 2.5h. After completion of reaction as indicated by TLC, the reaction mixture was filtered through Celite and filtrate concentrated under reduced pressure the reaction mixture was poured into water (100 mL) and extracted with ethyl acetate (4 x 25 mL). The combined organic layer was dried over anhydrous sodium sulphate and concentrated under vacuum to afford orange solid as crude (0.37g, Quantitative) LCMS m/z 215.1 & 217.1 (M & M+2)

**Step-3: Synthesis of 7-chloro-8-ethyl-10-(2-hydroxyethyl)benzo[g]pteridine-2,4(3H,10H)-dione (JG-2001)**

**Batch ID: STL-7-A-525-JG-2001-031-A**

To a solution of 2-((2-amino-4-chloro-5-ethylphenyl)amino)ethan-1-ol (**JG-2001-A2**) (0.300g, 1.40 mmol, 1.0 eq) in AcOH (6 mL), alloxan monohydrate (**CAS No: 2244-11-3)** (0.224g, 1.40mmol, 1.0 eq) and boric anhydride (0195 g, 2.80 mmol, 2.0 eq) were added and the reaction mixture was stirred at 60°C for 16 hrs. After completion of reaction as indicated by TLC, the reaction mixture was poured into water (50 mL) and extracted with 10 % DCM:MeOH (3x50 mL). The combined organic layer was dried over anhydrous sodium sulphate and concentrated under vacuum.The obtained crude product was further purified by column chromatography (10 % MDC:MeOH) to afford orange solid (0.221g, 49.31%) LCMS m/z 321.2 & 323.2(M & M+2); ^1^H NMR (400 MHz, DMSO) δ 11.46 (s, 1H), 8.22 (s, 1H), 8.05 (s, 1H), 4.97 (t, J = 5.6Hz, 1H), 4.71 (t, J=6,2H), 3.83 (m,2H), 2.95 (q, J=7.6, 2H), 1.30 (t, J = 7.6 Hz, 3H).

#### Synthesis of JG-2004

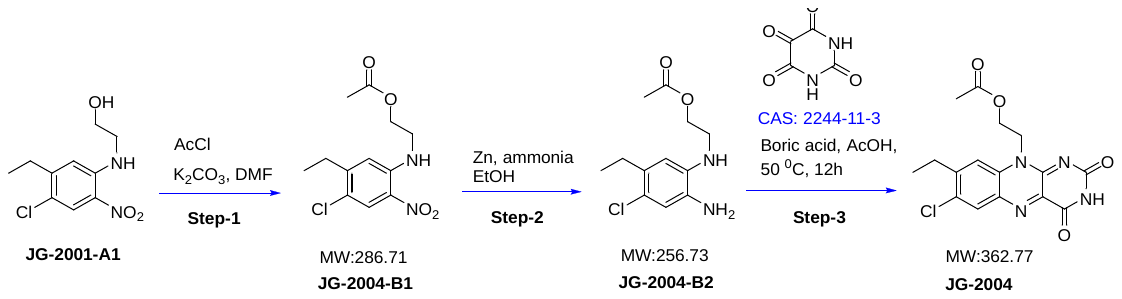

JG-2001-A1 (150 mg, 1 eq) was combined with acetyl chloride (3 eq) and K_2_CO_3_ (2 eq) in DMF, cooled to 0 ^0^C, then allowed to warm to room temperature for 30 minutes. Product formation was confirmed by LCMS and ^1^H-NMR and purified by column chromatography (JG-2004-B1, yield 130 mg). Next, JG-2004-B1 (120 mg, 1 eq) was combined with Zn (3 eq) and NH_4_Cl (5 eq) in a 1:1 mixture of ethanol and water, then incubated at room temperature for 3 hours. Product formation was confirmed by TLC, LCMS and ^1^H-NMR, then purified by column chromatography (JG-2004-B2, yield 50 mg). JG-2004-B2 (30 mg, 1 eq) was mixed with CA 2244-11-3 (1 eq) and CAS 1303-86-1 (1 eq) in acetic acid, heated to 70 ^0^C for 1 hour. Product formation (JG-2004) was confirmed by TLC, LCMS and purified by column chromatography (yield 10 mg).

**2-(7-chloro-8-ethyl-2,4-dioxo-3,4-dihydrobenzo[*g*]pteridin-10(2*H*)-yl)ethyl acetate (JG-2004)-**

1H NMR (400 MHz, DMSO, ppm) δ 11.49 (s, 1H), 8.23 (s, 1H), 8.04 (s, 1H), 4.89 (t, J = 5.6 Hz, 2H), 4.42 (t, J = 5.2 Hz, 2H), 2.95 (q, J = 7.2 Hz, 2H), 1.89 (s, 3H), 1.30 (t, J = 7.6 Hz, 3H). LCMS, calc’d 362.77 m/z; found 363.12 m/z

#### Synthesis of two methylene linker compounds (JG-2042, JG-2039, JG-2040, JG-2041, JG-2043, JG-2048, JG-2049, JG-2050, JG-2026, JG-2027)

**Synthesis of 7-chloro-8-ethyl-10-(2-(pyridin-3-yl)ethyl)benzo[g]pteridine-2,4(3H,10H)-dione**

**(JG-2042)**

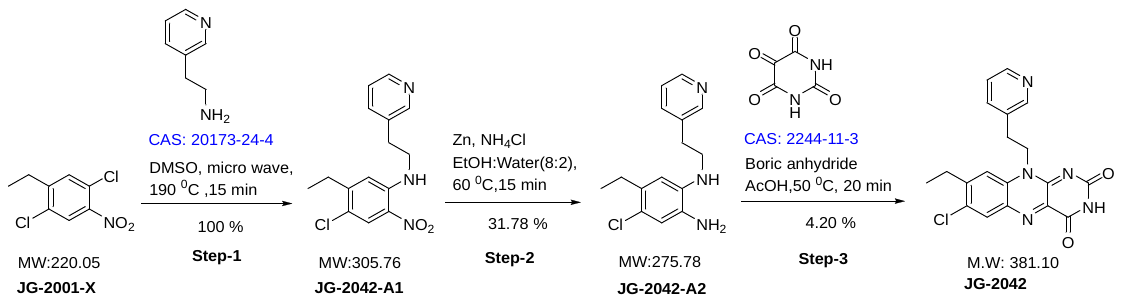

**Step-1: Synthesis of 4-chloro-5-ethyl-2-nitro-N-(2-(pyridin-3-yl)ethyl)aniline (JG-2042-A1)**

**Batch ID: STL-7-A-588-JG-2042-A1-111-a**

To a solution of 1,4-dichloro-2-ethyl-5-nitrobenzene (**JG-2001-X)** (0.200 g, 0.91 mmol, 1 eq) in DMSO (3 mL), 2-(yridine-3-yl)ethan-1-amine (**CAS: 20173-24-4)** (0.445 g, 3.65 mmol, 3.0 eq) was added and the reaction mixture was heated at 190°C under microwave irradiation for 15 min. After completion of reaction as indicated by TLC, the reaction mixture was poured into water (10 mL) and extracted with ethyl acetate (3 x 10 mL). The combined organic layer was dried over anhydrous sodium sulphate and concentrated under vacuum to afford yellow liquid (0.30g, 100 %) LCMS m/z 306.5 & 308.5(M & M+2)

Note: Crude material was directly used in next step without further purification.

**Step-2: Synthesis of 4-chloro-5-ethyl-N1-(2-(yridine-3-yl)ethyl)benzene-1,2-diamine (JG-2042-A2)**

**Batch ID: STL-7-A-588-JG-2042-A2-119-b**

To a solution of 4-chloro-5-ethyl-2-nitro-N-(2-(yridine-3-yl)ethyl)aniline (**JG-2042-A1)** (0.3g, 0.98 mmol, 1.0 eq) in EtOH: water (8:2 mL), Zn Dust (0.513g, 7.80 mmol, 8.0 eq) and NH_4_Cl (0.419g, 7.80 mmol, 8.0 eq) were added and stirred at 60°C for 15 min. After completion of reaction as indicated by TLC, the reaction mixture was fitered through Celite and filtrate was extracted with ethyl acetate (2 x 10 mL). The combined organic layer was dried over anhydrous sodium sulphate and concentrated under reduced pressure. The obtained crude product was further purified by column chromatography (20 % ethyl acetate/hexane) to obtain title compound as yellow solid (0.086g, 31.78 %) LCMS m/z 276.5 & 278.5 (M & M+2)

**Step-3: Synthesis of 7-chloro-8-ethyl-10-(2-(pyridin-3-yl)ethyl)benzo[g]pteridine-2,4(3H,10H)-dione (JG-2042)**

**Batch ID: STL-7-A-588-JG-2042-133-b**

To a solution of 4-chloro-5-ethyl-N1-(2-(pyridin-3-yl)ethyl)benzene-1,2-diamine(**JG-2042-A2**) (0.086g, 0.31 mmol, 1.0 eq) in AcOH (1.5 mL), alloxan monohydrate (**CAS No: 2244-11-3)** (0.049 g, 0.31 mmol, 1.0 eq) and boric anhydride (0.043 g, 0.62 mmol, 2.0 eq) were added and the reaction mixture was stirred at 50°C for 20 min. After completion of reaction as indicated by TLC, The reaction mixture was slowly poured into ice cooled water (10 mL) and extracted with ethyl acetate (4 x 10 mL). The combined organic layer was dried over anhydrous sodium sulphate and concentrated under reduced pressure. The obtained crude product was further purified by column chromatography (8.2 % MeOH/DCM) to obtain title compound as yellow solid (0.003g, 4.20 %). LCMS m/z 382.7 & 384.7(M & M+2); ^1^H NMR (400 MHz, DMSO) δ 11.48 (s, 1H), 8.53 (s, 1H), 8.41 (d, J = 4.0 Hz, 1H), 8.18 (s, 1H), 7.77 – 7.64 (m, 2H), 7.29 (dd, J = 7.5, 4.9 Hz, 1H), 4.85 (t, J = 6.8 Hz, 2H), 3.10 (t, J = 6.9 Hz, 2H), 2.88-2.80 (m, 2H), 1.20 (t, J = 7.4 Hz, 3H).

The following compounds were made according to the procedure described for JG-2042 using JG-2001-X.

| **Target ID** | **Structure** | **Analytical data** |
| --- | --- | --- |
| **JG-2039** | 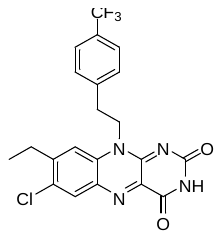 | LCMS m/z 449.8 & 451.8 (M & M+2); ^1^H NMR (400 MHz, DMSO) δ 11.49 (s, 1H), 8.18 (s, 1H), 7.66 (s, 1H), 7.63-7.54 (m, 4H), 4.86 (bs, 2H), 3.17 (bs, 2H), 2.82 (q, 2H), 1.17 (t, J = 6.8 Hz, 3H). |
| **JG-2040** | 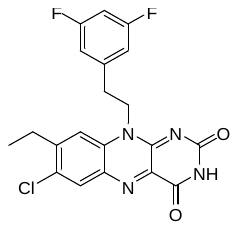 | LCMS m/z 417.1 & 419.0 (M & M+2); 1H NMR (400MHz, DMSO, ppm) δ 11.49 (s, 1H), 8.19 (s, 1H), 7.70 (s, 1H), 7.09 (m, J = 6.4 Hz, 3H), 4.84 (t, 2H), 3.10 (t, J = 6.4 Hz, 2H), 2.84 (q, J = 7.2 Hz, 2H), 1.21 (t, J = 7.2 Hz, 3H) |
| **JG-2041** | 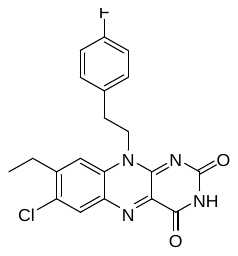 | LCMS m/z 399.2 & 401.0 (M & M+2); 1H NMR (400MHz, DMSO, ppm) δ 11.47 (s, 1H), 8.18 (s, 1H), 7.69 (s, 1H), 7.36 (q, J = 6 Hz, 2H), 7.10 (t, J = 8.8 Hz, 2H), 4.81 (t, J = 7.2 Hz, 2H), 3.05 (t, J = 6.85 Hz, 2H), 2.85 (q, J = 7.2 Hz, 2H), 1.21 (t, J = 7.6 Hz, 3H) |
| **JG-2043** | 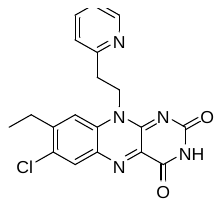 | LCMS m/z 382.6 & 384.7(M & M+2); ^1^H NMR (400 MHz, DMSO) δ 11.47 (s, 1H), 8.53 (d, J=3.6 Hz, 1H), 8.20 (s, 1H), 7.78 (s, 1H), 7.69 (m, 1H), 7.33 (d, J = 7.6Hz, 1H), 7.25(m, 1H), 4.98 (t, 2H), 3.24 (t, 2H), 2.87(q, 2H), 1.24 (t, J = 6.8 Hz, 3H). |
| **JG-2048** | 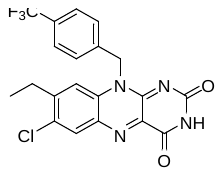 | LCMS m/z 435.1 & 437.1(M & M+2); 1H NMR (400MHz, DMSO, ppm) δ 11.52 (s, 1H), 8.25 (s, 1H), 7.65 (m, J = 8 Hz, 5H), 5.99 (s, 2H), 2.81 (q, J = 7.2 Hz, 2H), 1.13 (t, J = 7.6 Hz, 3H) |
| **JG-2049** | 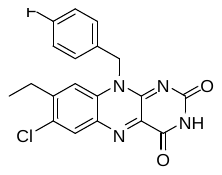 | LCMS m/z 385.6 &387.56(M & M+2); 1H NMR (400MHz, DMSO, ppm) δ 11.50 (s, 1H), 8.23 (s, 1H), 7.6 (s, 1H), 7.46 (d-d, J = 5.6 Hz, 2H), 7.17 (t, J = 8.8 Hz, 2H), 5.89 (s, 2H), 2.81 (q, J = 7.6 Hz, 2H), 1.15 (t, J = 7.6 Hz, 3H) |
| **JG-2050** | 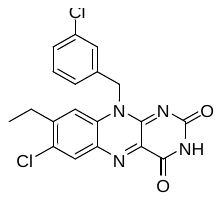 | LCMS m/z 401.2 & 403.1(M & M+2); 1H NMR (400 MHz, DMSO, ppm) δ 11.51 (s, 1H), 8.24 (s, 1H), 7.64 (s, 1H), 7.50 (s, 1H), 7.37 (d, J = 5.2 Hz, 3H), 5.90 (s, 2H), 2.81 (q, J = 7.2 Hz, 2H), 1.47 (t, J = 7.6 Hz, 3H) |
| **JG-2026** | 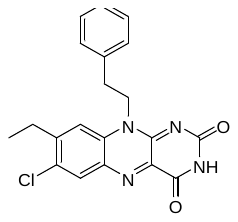 | LCMS m/z 381.1 & 383.1(M & M+2); 1H NMR (400MHz, DMSO, ppm) δ 11.47 (s, 1H), 8.19 (s, 1H), 7.7 (1H, s), 7.34 (d-d, J = 7.2 Hz, 2H), 7.27 (d-d, J = 7.6 Hz, 2H), 7.22 (m, J = 6.82 Hz, 1H), 4.8 (t, 2H), 3.06 (t, J = 6.8 Hz, 2H), 2.85 (q, J = 7.2 Hz, 2H), 1.23 (t, J = 7.2 Hz, 3H) |
| **JG-2027** | 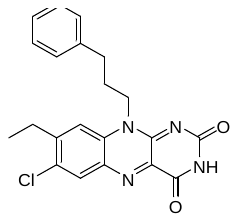 | LCMS m/z 385.1 & 397.3(M & M+2); 1H NMR (400MHz, DMSO, ppm) δ 11.44 (s, 1H), 8.21 (s, 1H), 7.67 (s, 1H), 7.29 (d-d, J = 6.8 Hz, 2H), 7.29 (d-d, J = 6.8 Hz, 2H), 7.21 (m, J = 6.4 Hz, 1H), 4.6 (t, 2H), 3.13 (q, J = 7.7 Hz, 2H), 2.90 (m, J = 7.6 Hz, 2H), 2.81 (t, J = 7.6 Hz, 2H), 1.25 (t, J = 7.6Hz, 3H) |

#### Synthesis methylene sidechain derivatives (JG-2046, JG-2047, JG-2036, JG-2038, JG-2045, JG-2028)

**Synthesis of 7-chloro-10-(cyclohexylmethyl)-8-ethylbenzo[g]pteridine-2,4(3H,10H)-dione**

**(JG-2046)**

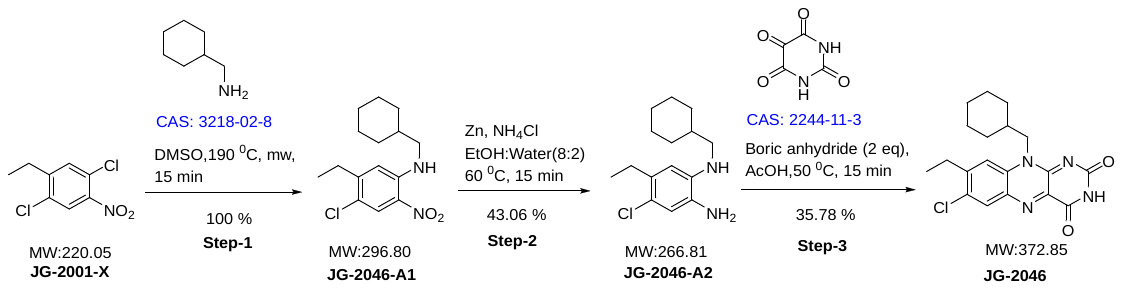

**Step-1: Synthesis of 4-chloro-N-(cyclohexylmethyl)-5-ethyl-2-nitroaniline**

**Batch ID: STL-7-A-588-JG-2046-A1-110-a**

To a solution of 1,4-dichloro-2-ethyl-5-nitrobenzene (**JG-2001-X**) (0.200 g, 0.91 mmol, 1 eq) in DMSO (3 ml), cyclohexylmethanamine (**CAS: 3218-02-8)** (0.413 g, 3.62 mmol, 3.0 eq) was added and the reaction mixture was stirred at 190°C temperature under microwave irradiation for 15 min. After completion of reaction as indicated by TLC, the reaction mixture was poured into water (10 mL) and extracted with ethyl acetate (3 x 10mL). The combined organic layer was dried over anhydrous sodium sulphate and concentrated under vacuum to obtain title compound as yellow solid (0.310 g, 100 %). LCMS m/z 297.6 &299.7 (M & M+2)

Crude material which was directly used in next step without further purification.

**Step-2: Synthesis of 4-chloro-N1-(cyclohexylmethyl)-5-ethylbenzene-1,2-diamine**

**Batch ID: STL-7-A-588-JG-2046-A2-120-b**

To a solution of 4-chloro-N-(cyclohexylmethyl)-5-ethyl-2-nitroaniline (**JG-2046-A1**) (0.310g, 1.04 mmol, 1.0 eq) in EtOH: water (8:2 mL), Zn Dust (0.546g, 8.35 mmol, 8.0 eq) and NH4Cl (0.446g, 8.35 mmol, 8.0 eq) were added and the reaction mixture was stirred at 60°C for 15 min. After completion of reaction as indicated by TLC, the reaction mixture was filtered through celite and extracted with ethyl acetate (2 x 10 mL). The combined organic layer was dried over anhydrous sodium sulphate and concentrated under vacuum. The obtained crude product was further purified by column chromatography (30 % ethyl acetate/hexane) to obtain yellow solid (0.120g, 43.06 %). LCMS m/z 267.6 & 269.6 (M & M+2).

**Step-3: Synthesis of 7-chloro-10-(cyclohexylmethyl)-8-ethylbenzo[g]pteridine-2,4(3H,10H)-dione**

**Batch ID: STL-7-A-588-JG-2046-134-a**

To a solution of 4-chloro-N1-(cyclohexylmethyl)-5-ethylbenzene-1,2-diamine **(JG-2046-A2**) (0.120g, 0.44mmol, 1.0 eq) in AcOH (1.5 mL), alloxan monohydrate (**CAS: 2244-11-3)** (0.072 g, 0.44 mmol, 1.0 eq) and Boric anhydride (0.062 g, 0.89 mmol, 2.0 eq) were added and the reaction mixture was stirred at 50°C for 15 min. After completion of reaction as indicated by TLC, the reaction mixture was poured into water and extracted with ethyl acetate (4 x 10 mL). The reaction mixture was dried over anhydrous sodium sulphate and concentrated under vacuum. The obtained crude product was further purified by column chromatography (5 % MeOH and DCM) to obtain title compound as yellow solid (0.060g, 35.78 %) LCMS m/z 373.8 (M+1); 1H NMR (400 MHz, DMSO) δ 11.42 (s, 1H), δ 8.21 (s, 1H), 7.96 (s, 1H), 4.50 (bs, 2H), 2.95 (q, J=7.2Hz 2H), 2.10-1.90 (m, 1H), 1.70 – 1.52 (m, 4H), 1.30 (t, 3H), 1.30-1.00 (m, 6H).

The following compounds were made according to the procedure described for **JG-2046** using

**JG-2001-X**

| **Target ID** | **Structure** | **Analytical data** |
| --- | --- | --- |
| **JG-2047** | 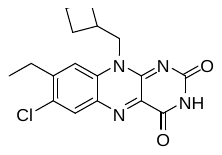 | LCMS m/z 345.5 & 347.6 (M&M+2); 1H NMR (400MHz, DMSO, ppm) δ 11.43 (s, 1H), 8.21 (s, 1H), 8.0 (s, 1H), 4.72 (d, J = 6.4 Hz, 2H), 2.95 (q, J = 7.2 Hz, 2H), 2.89 (m, J = 7.6 Hz, 1H), 1.98 (m, J = 8.8 Hz, 4H), 1.80 (m, J = 7.6 Hz, 2H), 1.29 (t, J = 7.2 Hz, 3H) |
| **JG-2036** | 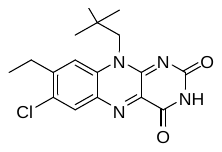 | LCMS m/z 347.5 & 349.9 (M&M+2); 1H NMR (400MHz, DMSO, ppm) δ 11.44 (s, 1H), 8.19 (s, 1H), 8.08 (s, 1H), 4.97 (s, 1H), 4.34 (s, 1H), 2.94 (q, J = 7.2 Hz, 2H), 1.27 (t, J = 7.2 Hz, 3H), 1.00 (s, 9H) |
| **JG-2038** | 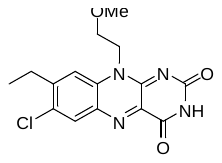 | LCMS m/z 335.6 & 337.6 (M&M+2); 1H NMR (400 MHz, DMSO) δ 11.45 (s, 1H), δ 8.20 (s, 1H), 8.02 (s, 1H), 4.80 (t, 2H), 3.75(t, 2H), 3.26 (s, 3H), 2.94 (q, J=7.2Hz, 2H), 1.29 (t, J=7.2Hz, 3H). |
| **JG-2045** | 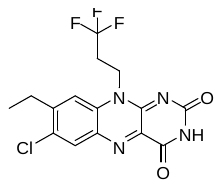 | LCMS m/z 373.2 & 375.2 (M&M+2); 1H NMR (400MHz, DMSO, ppm) δ 11.52 (s, 1H), 8.25 (s, 1H), 7.90 (s, 1H), 4.85 (t, J = 6.8 Hz, 2H), 2.96 (q, J = 7.2 Hz, 2H), 2.83 (q, J = 7.6 Hz, 2H), 1.35 (t, J = 7.2 Hz, 3H) |
| **JG-2028** | 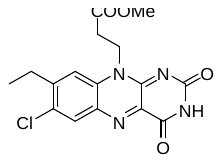 | LCMS m/z 363.5 & 365.7 (M&M+2); 1H NMR (400MHz, DMSO, ppm) δ 11.48 (s, 1H), 8.22 (s, 1H), 8.05 (s, 1H), 4.80 (t, J = 7.6 Hz, 2H), 3.66 (s, 3H), 2.97 (q, J = 7.6 Hz, 2H), 2.82 (t, J = 7.6 Hz, 2H), 1.3 (t, J = 7.6 Hz, 3H) |

#### Synthesis of JG-2003

**(10-(2-(benzyloxy)ethyl)-7-chloro-8-ethylbenzo[g]pteridine-2,4(3H,10H)-dione)**

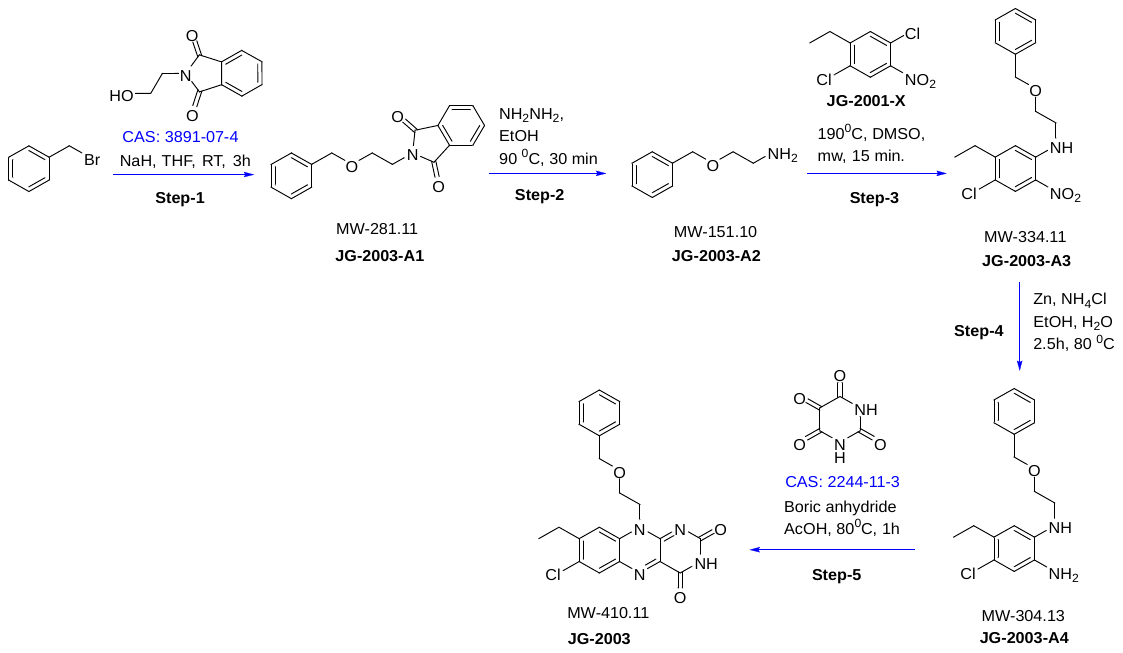

**Step-1: Synthesis of 2-(2-(benzyloxy)ethyl)isoindoline-1,3-dione**

**Batch ID: STL-7-A-525-JG-2003-A1-095-A**

To a solution of (bromomethyl)benzene (**CAS:100-39-0)** (1.0 g, 5.23 mmol, 1 eq) in THF (10 mL), NaH(0.19 g, 7.84 mmol, 1.5 eq) and 2-(2-hydroxyethyl)isoindoline-1,3-dione (0.89g, 5.23 mmol, 1eq ) were added and the reaction mixture was stirred at RT for 3h. After completion of reaction as indicated by TLC, the reaction mixture was poured into water (50 mL) and extracted with ethyl acetate (3 x 50 mL). The combined organic layer was dried over anhydrous sodium sulphate and concentrated under vacuum to obtain yellow liquid (1.19g,72.40 %). The crude was forwarded to next step.

**Step-2: Synthesis of 2-(benzyloxy)ethan-1-amine**

**Batch ID: STL-7-A-525-JG-2003-A2-096-Ao**

To a solution of 2-(2-(benzyloxy)ethyl)isoindoline-1,3-dione **(JG-2003-A1)** (1.19g) in ethanol(17 mL), hydrazine hydrate (1.7 ml ) was added and the reaction mixture was stirred at 90 ^0^C for 30 min. After completion of reaction as indicated by TLC, the reaction mixture was filtered and filtrate was concentrated under vacuum. The filtrate was poured into water (50 mL) and extracted with (9:1 MDC : MeOH) (3 x 50 mL). The combined organic layer was dried over anhydrous sodium sulphate and concentrated under vacuum to afford yellow liquid (0.60g, 93.80 %) LCMS m/z 152.1 (M+1)

**Step-3: Synthesis of N-(2-(benzyloxy)ethyl)-4-chloro-5-ethyl-2-nitroaniline**

**Batch ID: STL-7-A-525-JG-2003-A3-098-A**

To a solution of 1,4-dichloro-2-ethyl-5-nitrobenzene (**JG-2001-X)** (0.29g, 1.31 mmol, 1.0eq) in DMSO (9 mL), 2-(benzyloxy)ethan-1-amine (**JG-2003-A2)** (0.600 g, 3.95 mmol, 3.0 eq) was added and the reaction mixture was heated at 190°C under microwave irradiation for 15 min. After completion of reaction as indicated by TLC, the reaction mixture was poured into ice cold water (50 mL) and extracted with ethyl acetate (3 x 50 mL). The combined organic layer was dried over anhydrous sodium sulphate and concentrated under vacuum to obtain orange solid. The crude was carry forwarded for next step without any purification. LCMS m/z 335.3 (M+1)

**Step-4: Synthesis of N1-(2-(benzyloxy)ethyl)-4-chloro-5-ethylbenzene-1,2-diamine**

**Batch ID: STL-7-A-525-JG-2003-A4-099-A**

To a solution of N-(2-(benzyloxy)ethyl)-4-chloro-5-ethyl-2-nitroaniline (**JG-2003-A3)** (1.03g, 3.08 mmol, 1.0 eq) in EtOH: water (10:2.5 mL), Zn Dust (1.60g, 24.5 mmol, 8.0 eq) and NH_4_Cl (1.31g, 25.52 mmol, 8.0 eq) were added and stirred at 80 °C for 2.5 hr. After completion of reaction as indicated by TLC, the reaction mixture was filtered through Celite and filtrate was concentrated under reduced pressure. The filtrate was poured into ice cold water (50 mL) and extracted with ethyl acetate (3 x 50 mL). The combined organic layer was dried over anhydrous sodium sulphate and concentrated under vacuum. The crude was purified by column chromatography (30 % ethyl acetate/hexane) to obtain title compound (0.16g, 17.15 %, over 2 steps). LCMS m/z 304.6 & 306.9 (M & M+2)

**Step-5: Synthesis of 10-(2-(benzyloxy)ethyl)-7-chloro-8-ethylbenzo[g]pteridine-2,4(3H,10H)-dione (JG-2003)**

**Batch ID: STL-7-A-525-JG-2003-117-B**

To a solution of N1-(2-(benzyloxy)ethyl)-4-chloro-5-ethylbenzene-1,2-diamine (**JG-2003-A4**) (0.15 g, 0.493 mmol, 1.0 eq) in AcOH (3 mL), alloxan monohydrate (**CAS No: 2244-11-3)** (0.078g, 0.49 mmol, 1.0 eq) and boric anhydride (0.068 g, 0.98 mmol, 2.0 eq) were added and the reaction mixture was stirred at 80°C for 1 hr . After completion of reaction as indicated by TLC, The reaction mixture was poured into ice cooled water (10 mL) and filtered the solid. The obtained crude product was further triturated with diethylether to obtain title compound as yellow solid (0.065g, 32.13 %). LCMS m/z 411.6 & 413.6 (M & M+2); ^1^H NMR (400 MHz, DMSO) δ 11.44 (s, 1H), 8.19 (s, 1H), 8.03 (s, 1H), 7.22 - 7.11 (m, 5H), 4.85 (t, 2H), 4.45 (s,2H), 3.91 (t, J=4.8 Hz, 2H), 2.87 (q, J=7.2 Hz, 2H), 1.17 (t, J = 7.2 Hz, 3H).

#### Synthesis of benzyl-oxy-ethyl derivatives (JG-2068, JG-2069, JG-2070, JG-2071, JG-2072, JG-2073, JG-2084, JG-2085, JG-2002-G4)

**Synthesis of 7-chloro-8-ethyl-10-(2-((4-fluorobenzyl)oxy)ethyl)benzo[g]pteridine-2,4(3H,10H)-dione (JG-2068)**

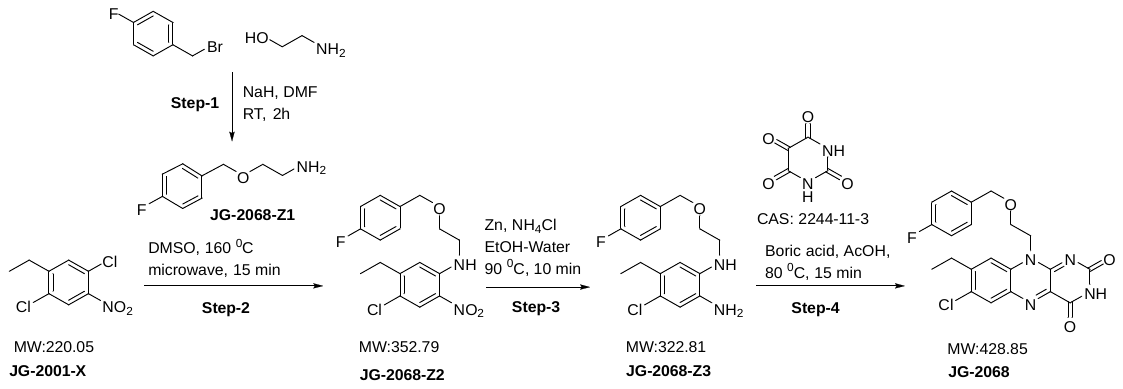

**Step-1: Synthesis of 2-((4-fluorobenzyl)oxy)ethan-1-amine**

**Batch ID: STL-7-A-607-JG-2001-Z1-019a**

To a solution of 1-(bromomethyl)-4-fluorobenzene (**CAS: 459-46-1)** (1.0 g, 5.29 mmol, 1 eq) in DMF (5 mL), 2-aminoethan-1-ol (**CAS: 141-43-5)** (0.645 g, 10.5 mmol, 2.0 eq) and NaH (0.126 g, 5.29 mmol, 1 eq) were added and the reaction mixture was stirred at RT for 2 h. After completion of reaction as indicated by TLC, the reaction mixture was quenched by methanol and poured into water (50 mL) and extracted with ethyl acetate (3 x 50 mL). The combined organic layer was dried over anhydrous sodium sulphate and concentrated under vacuum to afford yellow liquid. The crude was forwarded to next step. LCMS m/z 170 (M+1)

**Step-2: Synthesis of 4-chloro-5-ethyl-N-(2-((4-fluorobenzyl)oxy)ethyl)-2-nitroaniline**

**Batch ID: STL-7-A-588-JG-2001-Z2-183-A**

To a solution of 1,4-dichloro-2-ethyl-5-nitrobenzene (**JG-2001-X)** (0.22g, 0.99 mmol, 1.0 eq) in DMSO (8 mL), 2-((4-fluorobenzyl)oxy)ethan-1-amine(**JG-2001-Z1)** (0.84 g, 4.70 mmol, 5.0 eq) was added and the reaction mixture was heated at 160 °C under microwave irradiation for 15 min. After completion of reaction as indicated by TLC, the reaction mixture was poured into ice cold water (50 mL) and extracted with ethyl acetate (3 x 50 mL). The combined organic layer was dried over anhydrous sodium sulphate and concentrated under vacuum. The obtained crude product was further purified by column chromatography (45 % ethyl acetate/hexane) to obtain title compound as yellow solid (0.045g, 12.93 %) LCMS m/z 353.0 & 355.1 (M & M+2)

**Step-3: Synthesis of 4-chloro-5-ethyl-N1-(2-((4-fluorobenzyl)oxy)ethyl)benzene-1,2-diamine**

**Batch ID: STL-7-A-607-JG-2001-Z3-029-A**

To a solution of 4-chloro-5-ethyl-N-(2-((4-fluorobenzyl)oxy)ethyl)-2-nitroaniline (**JG-2001-Z2)** (0.29g, 0.82 mmol, 1.0 eq) in EtOH: water (4:0.5 mL), Zn Dust (0.43g, 6.57 mmol, 8.0 eq) and NH_4_Cl (0.35 g, 6.57 mmol, 8.0 eq) were added and stirred at 90 °C for 10 min. After completion of reaction as indicated by TLC, the reaction mixture was filtered through Celite and filtrate was concentrated under reduced pressure. The obtained crude product was further purified by column chromatography (50 % ethyl acetate/hexane) to obtain title compound as yellow solid (0.19g, 71.60 %). Mass Ms 323.54 & 325.54 (M+1)

**Step-4: Synthesis of 7-chloro-8-ethyl-10-(2-((4-fluorobenzyl)oxy)ethyl)benzo[g]pteridine-2,4(3H,10H)-dione**

**Batch ID: STL-7-A-607-JG-2068-035-c**

To a solution of 4-chloro-5-ethyl-N1-(2-((4-fluorobenzyl)oxy)ethyl)benzene-1,2-diamine (**JG-2001-Z3**) (0.19g, 0.59 mmol, 1.0 eq) in AcOH (3 mL), alloxan monohydrate (**CAS No: 2244-11-3)** (0.094g, 0.11 mmol, 1.0 eq) and boric anhydride (0.082 g, 0.11 mmol, 2.0 eq) were added and the reaction mixture was stirred at 80°C for 15 min. After completion of reaction as indicated by TLC, The reaction mixture was poured into ice cooled water (10 mL) and filtered. The obtained crude product was further triturated with diethyl ether to obtain title compound as yellow solid (0.020 g, 7.92 %). LCMS m/z 429.0 & 431.1 (M & M+2); ^1^H NMR (400 MHz, DMSO) δ 11.44 (s, 1H), 8.18 (s,1H), 8.00 (s,1H), 7.20-7.12 (m,2H), 7.06-7.02 (m,2H), 4.84 (bs,2H), 4.43 (s,2H), 3.89 (bs,2H), 2.84 (q, J=7.2, 2H), 1.14 (t, J=7.2 Hz, 3H).

The following compounds were made according to the procedure described for **JG-2068** using

**JG-2001-X**

| **Target ID** | **Structure** | **Analytical data** |
| --- | --- | --- |
| **JG-2069** | 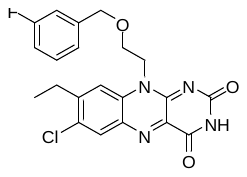 | LCMS m/z 429.0 & 431.0(M & M+2); ^1^H NMR (400 MHz, DMSO) δ 11.44 (s, 1H), 8.19 (s, 1H), 8.04(s, 1H), 7.29-7.23(m, 1H), 7.05-6.95(m, 2H), 6.87 (d,J=10 Hz,1H), 4.88-485(m, 2H), 4.46(s, 2H), 3.93-3.91 (m, 2H), 2.86 (q, J = 7.6 Hz, 2H), 1.16(t J=7.6 Hz 3H). |
| **JG-2070** | 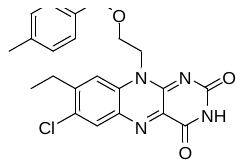 | LCMS m/z 425.2 & 427.1(M & M+2); ^1^H NMR (400 MHz, DMSO) δ 11.43(s, 1H), 8.18 (s, 1H), 7.99(s, 1H), 6.98 (d, J=8 Hz, 2H), 6.94 (d, J=8 Hz, 2H), 4.82(t, J=4.8, 2H), 4.37(s, 2H), 3.87 (t, J=4.8, 2H), 2.86(t, J=7.6, 2H), 2.22(s, 3H), 1.18(t, J=7.2 Hz, 3H). |
| **JG-2071** | 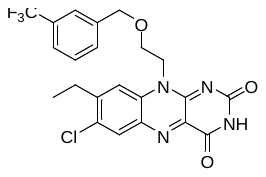 | LCMS m/z 479.1 & 481(M & M+2); ^1^H NMR (400 MHz, DMSO) δ 11.44(s, 1H), 8.18 (s, 1H), 8.03(s, 1H), 7.56 (d, J =7.2 Hz, 1H), 7.48-7.41(m, 2H), 7.36(s, 1H), 4.89(t,J=4.8, 2H), 4.54(s, 2H), 3.96 (t, J=4.8, 2H), 2.81 (q, J=7.2Hz, 2H), 1.09(t, J =7.2 Hz, 3H). |
| **JG-2072** | 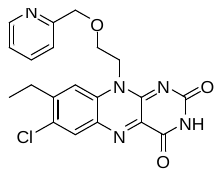 | LCMS m/z 412.0 & 414.0(M & M+2); 1H NMR (400 MHz, DMSO) δ 11.45 (s, 1H) 8.43-8.42 (m, 1H), 8.35(d, 1H), 8.18(s, 1H), 8.02(s, 1H), 7.51 (d, J=7.6 Hz, 1H), 7.25(dd, J=4.8 & 8Hz, 1H), 4.86(t, 2H), 4.49(s, 2H), 3.93 (t, 2H), 2.84 (q, J = 7.2 Hz, 2H), 1.13(t, J=7.2Hz, 3H). |
| **JG-2073** | 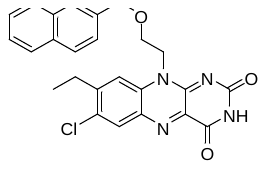 | LCMS m/z 461.1 & 463.1(M & M+2); ^1^H NMR (400 MHz, DMSO) δ 11.43(s, 1H), 8.17 (s, 1H), 8.03(s, 1H), 7.86-7.70 (m, 3H), 7.58(s, 1H), 7.50-7.46(m, 2H), 7.22(d, J=8.4Hz, 1H), 4.89(t, 2H), 4.61(s, 2H), 3.96(t, J=4.8, 2H), 2.79(q, J=7.2Hz, 2H), 1.09(t, J = 7.6 Hz, 3H). |
| **JG-2084** | 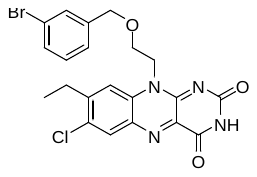 | LCMS m/z 488.9 & 490.9 (M & M+2); ^1^HNMR (400 MHz, DMSO) δ 11.44(s, 1H) ,8.19(s, 1H), 8.03(s, 1H), 7.39 (d, J=7.6Hz, 1H), 7.20-7.10 (m,3H), 4.87 (t, 2H), 4.44(s, 2H), 3.93(t, 2H), 2.87(q, J=6.8, 2H), 1.15(t, J=7.2Hz, 3H). |
| **JG-2085** | 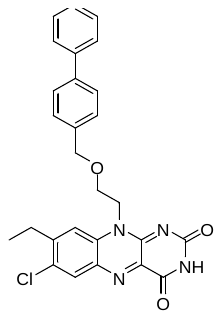 | LCMS m/z 487.1 & 489.3(M & M+2); ^1^H NMR (400 MHz, DMSO) δ 11.44 (s, 1H), 8.18 (s, 1H), 8.03(s, 1H), 7.61 (d, J = 7.2 Hz, 2H), 7.51-7.35(m, 5H), 7.19(d, J= 8.0Hz, 2H), 4.87 (m,2H),4.49(. 2H), 3.93 (m, 2H), 2.85 (q, J=7.2Hz, 2H), 1.15 (t, J = 7.6 Hz, 3H). |
| **JG-2002-G4** | 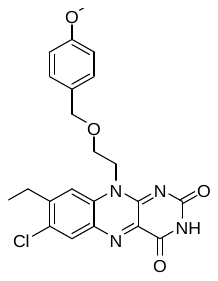 | LCMS m/z 441.8 & 443.9 (M & M+2); ^1^H NMR (400 MHz, DMSO) δ 11.44 (s, 1H), 8.19 (s, 1H), 7.98(s, 1H), 7.02(d, J=8.4Hz, 2H), 6.74(d,J=8.4Hz, 2H), 4.81(m, 2H), 4.36(bs, 2H), 3.86 (bs, 2H), 3.70(s, 3H), 2.85 (q, 2H), 1.17(t, 3H). |

#### Synthesis of alkyl ether analogues (JG-2005, JG-2006, JG-2008, JG-2009, JG-2014, JG-2015, JG-2016, JG-2092)

**Synthesis of 7-chloro-10-(2-(cyclohexylmethoxy)ethyl)-8-ethylbenzo[g]pteridine-2,4(3H,10H)-dione**

**(JG-2005)**

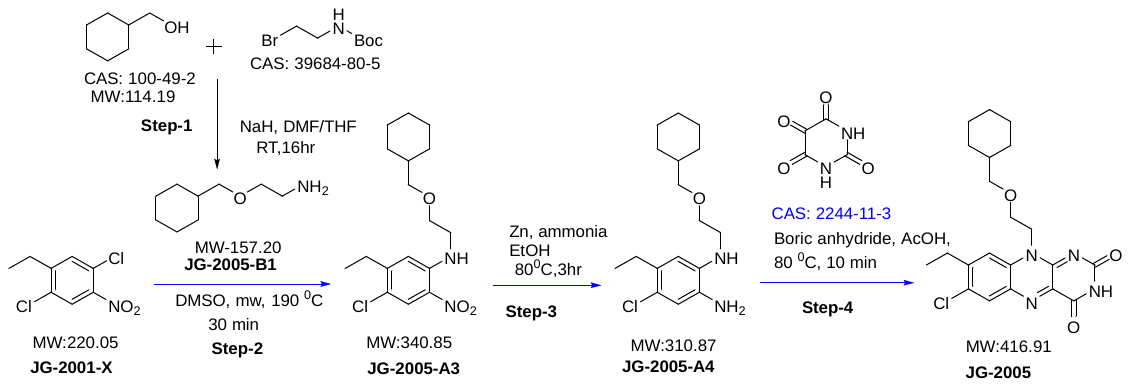

**Step-1: Synthesis of 2-(cyclohexylmethoxy)ethan-1-amine**

**Batch ID: STL-7-A-525-JG-2005-B1-188-A**

To a solution of cyclohexylmethanol (**CAS:100-49-2)** (1.0 g, 8.75 mmol, 1 eq) in THF (10 mL), tert-butyl (2-bromoethyl)carbamate (**CAS:39684-80-5)** (1.96g, 8.75 mmol, 1eq) and NaH(1.05 g, 43.7 mmol, 5 eq) were added and the reaction mixture was stirred at RT for 16 hr. After completion of reaction as indicated by TLC, the reaction mixture was quenched by methanol and concentrated under vacuum the reaction mixture was poured into water (50 mL) and extracted with (9:1 MDC in MeOH) (3 x 50 mL). The combined organic layer was dried over anhydrous sodium sulphate and concentrated under vacuum to to afford yellow liquid (1.68g, crude) 158.4 (M+1).

Note: TLC and analysis shown that Boc group was cleaved in reaction itself.

**Step-2: Synthesis of 4-chloro-N-(2-(cyclohexylmethoxy)ethyl)-5-ethyl-2-nitroaniline**

**Batch ID: STL-7-A-525-JG-2005-A3-192-A**

To a solution of 1,4-dichloro-2-ethyl-5-nitrobenzene (**JG-2001-X)** (0.300 g, 1.36 mmol, 1.0eq) in DMSO (2 mL), 2-(cyclohexylmethoxy)ethan-1-amine (**JG-2005-B1)** (0.645 g, 4.10 mmol, 3.0 eq) was added and the reaction mixture was heated at 190°C under microwave irradiation for 30 min. After completion of reaction as indicated by TLC, the reaction mixture was poured into ice cold water (50 mL) and extracted with ethyl acetate (3 x 50 mL). The combined organic layer was dried over anhydrous sodium sulphate and concentrated under vacuum to The obtained crude product was further purified by column chromatography ( neat hexane) to afford orange solid (0.057 g, 12.27%) LCMS m/z 341.7 & 343.7 & (M&M+2)

**Step-3: Synthesis of 4-chloro-N1-(2-(cyclohexylmethoxy)ethyl)-5-ethylbenzene-1,2-diamine**

**Batch ID: STL-7-A-525-JG-2005-A4-200-A**

To a solution of 4-chloro-N-(2-(cyclohexylmethoxy)ethyl)-5-ethyl-2-nitroaniline (**JG-2005-A3)** (0.215g, 0.63 mmol, 1.0 eq) in EtOH: water (5 : 0.8 mL), Zn Dust (0.330g, 5.05mmol, 8.0 eq) and NH_4_Cl (0.270g, 5.05 mmol, 8.0 eq) were added and stirred at 80 °C for 3 hr. After completion of reaction as indicated by TLC, the reaction mixture was filtered through Celite and filtrate concentrated under reduced pressure. The obtained crude product was further purified by column chromatography (35% ethyl acetate/ hexane) to obtain title compound as gray liquid (0.054g, 27.54%) LCMS m/z 311.6 & 313.6 (M & M+2)

**Step-4: Synthesis of 7-chloro-10-(2-(cyclohexylmethoxy)ethyl)-8-ethylbenzo[g]pteridine-2,4(3H,10H)-dione**

**Batch ID: STL-7-A-588-JG-2005-001-B**

To a solution of 4-chloro-N1-(2-(cyclohexylmethoxy)ethyl)-5-ethylbenzene-1,2-diamine (**JG-2005-A4**) (0.054, 0.17 mmol, 1.0 eq) in AcOH (1.5 mL), alloxan monohydrate (**CAS No: 2244-11-3)** (0.027g, 0.17mmol, 1.0 eq) and boric anhydride (0.024 g, 0.34 mmol, 2.0 eq) were added and the reaction mixture was stirred at 80°C for 10 min. After completion of reaction as indicated by TLC, the reaction mixture was poured into ice cold water (50 mL) and extracted with ethyl acetate (3 x 50 mL). The combined organic layer was dried over anhydrous sodium sulphate and concentrated under vacuum. The obtained crude product was further purified by column chromatography (97% ethyl acetate/hexane) to afford yellow solid (0.007g, 13.81%). LCMS m/z 417.3 & 419.2(M&M+2); 1HNMR (400MHz,DMSO) δ 11.44(s,1H), 8.20(s,1H), 8.04(s,1H), 4.81(t,2H), 3.78(t,2H), 3.15(d,J=6.0,2H), 2.92(q, J=7.6,2H), 1.56-1.46(m, 5H), 1.29-0.98(m, 7H), 0.77-72(m,2H)

**Synthesis of 7-chloro-8-ethyl-10-(2-(neopentyloxy)ethyl)benzo[g]pteridine-2,4(3H,10H)-dione (JG-2014)**

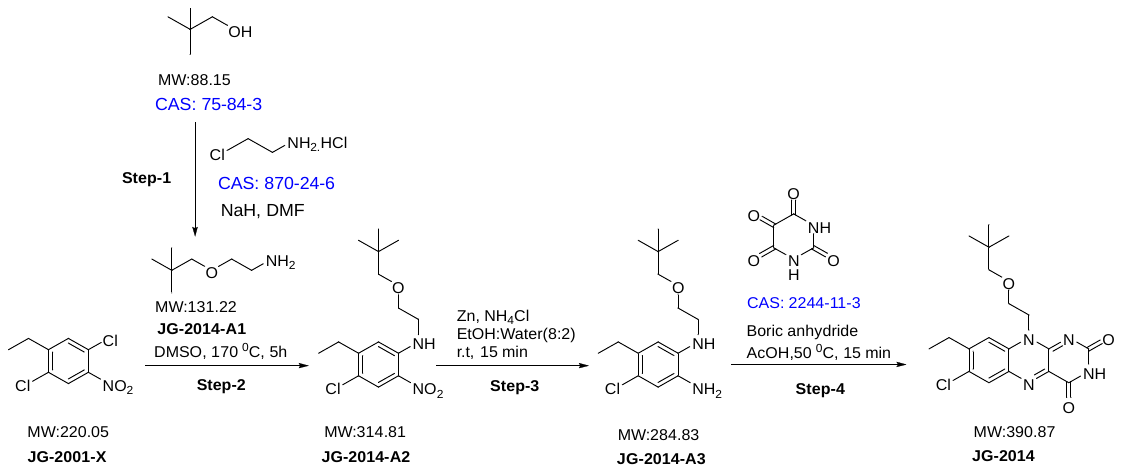

**Step-1: Synthesis of 2-(neopentyloxy)ethan-1-amine (JG-2014-A1)**

**Batch ID: STL-9-A708-JG-2014-A1-036**

To a solution of 2,2-Dimethyl-1-propanol (**CAS: 75-84-3**) (10 g, 113.44 mmol, 1.0 eq) in DMF (100 mL/ 10V), 60% NaH (22.68 g, 567.2 mmol, 5.0 eq) was added portion wise. The reaction mixture was stirred at 0°C temperature for 1.5h under nitrogen atmosphere. 2-Chloroethylamine hydrochloride(**CAS: 870-24-6**) (19.73g, 170.16 mmol, 1.5 eq) was added portion wise to the reaction mixture. The reaction mixture was allowed to stir for 16 hr at room temperature. After completion of reaction as indicated by TLC (mobile phase: 10% MeOH in DCM), the reaction mixture was slowly dumped in to cold aqueous sodium chloride solution (200 mL). The reaction mixture was extracted with ethyl acetate (3 x 1.5L) and combined organic layer was washed with water (2x 500 mL) and dried over anhydrous sodium sulphate. The organic layer was dried over anhydrous sodium sulphate and concentrated under vacuum to afford yellow bi-phasic liquid (12g, 80.61%) which was used in next step without further purification. LCMS: 1.186 min, MS: ES+ 132.34(M+1)

**Step-2: Synthesis of 4-chloro-5-ethyl-N-(2-(neopentyloxy)ethyl)-2-nitroaniline (JG-2014-A2)**

**Batch ID: STL-9-A708-JG-2014-A2-069**

To a solution of 1,4-dichloro-2-ethyl-5-nitrobenzene (**JG-2001-X**) (1.0g, 4.54 mmol, 1.0 eq) in DMSO (10 mL/ 10V) was added 2-(neopentyloxy)ethan-1-amine (**JG-2014-A1**) (2.98 g, 22.70 mmol, 5.0 eq). The reaction mixture was stirred at 190°C temperature under microwave for 30 to 45 min. After completion of reaction as indicated by TLC, the reaction mixture was slowly dumped in ice cooled water (300 mL) and extracted with ethyl acetate (3 x 100 mL). The reaction mixture was dried over anhydrous sodium sulphate and concentrated under vacuum. The obtained crude product was further purified by column chromatography (0.4 % ethyl acetate/hexane) to give title compound as yellow solid (0.25g, 17.47 %). **Note:** Above reaction was repeated in 1g parallel batch under microwave irradiation.

LCMS: 315.3, 317.3(M & M+2); 1H NMR (400 MHz, DMSO) δ 8.25 (t, J = 4.8 Hz, 1H), 8.01 (s, 1H), 7.07 (s, 1H), 3.65-3.63 (m, 2H), 3.56-3.52 (m, 2H), 3.10 (s, 2H), 2.67 (q, J = 7.4 Hz, 2H), 1.18 (t, 3H), 0.85 (s, 9H).

**Step-3: Synthesis of 4-chloro-5-ethyl-N1-(2-(neopentyloxy)ethyl)benzene-1,2-diamine (JG-2014-A3)**

**Batch ID: STL-9-A708-JG-2014-A3-073a**

To a solution of 4-chloro-5-ethyl-N-(2-(neopentyloxy)ethyl)-2-nitroaniline (**JG-2014-A2**) (2.3g, 7.46 mol, 1.0 eq) in ethanol: water (10 mL, 8:2 V) were added Zn Dust (3.90g, 59.7 mmol, 8.0 eq) and NH4Cl (3.19g, 59.7 mmol, 8.0 eq). The reaction mixture was stirred at room temperature for 15 min. After completion of reaction as indicated by TLC, the reaction mixture was filtered through Celite and extracted with ethyl acetate (2 x 50 mL). The filtrate was dried over anhydrous sodium sulphate and concentrated under reduced pressure to afford yellow liquid (2.0g, 96.11 %). It was directly used in next step without further purification. LCMS: 285.1, 286.9(M & M+2).

**Step-4: Synthesis of 7-chloro-8-ethyl-10-(2-(neopentyloxy)ethyl)benzo[g]pteridine-2,4(3H,10H)-dione (JG-2014)**

**Batch ID: STL-9-A708-JG-2014-103-b**

To a solution of 4-chloro-5-ethyl-N1-(2-(neopentyloxy)ethyl)benzene-1,2-diamine (JG-2014-A3) (1.17g, 4.10 mmol, 1.0 eq) in AcOH (6 mL), alloxan monohydrate (**CAS: 2244-11-3**) (0.65 g, 4.10 mmol, 1.0 eq) and boric anhydride (0.571g, 8.20 mmol, 2.0 eq) were added. The reaction mixture was stirred at 50 °C temperature for 15 min. TLC indicated completion of reaction. The reaction mixture was slowly poured into ice cooled water (50 mL) and extracted with ethyl acetate (3 x 50 mL). The combined organic layer was dried over anhydrous sodium sulphate and concentrated under reduce pressure. The crude was purified by column chromatography (70% ethyl acetate/hexane) to afford yellow solid (0.7g, 42.60 %). LCMS: 391.1, 393.1(M& M+2). 1H NMR (400 MHz, DMSO) δ 11.43 (s, 1H), 8.19 (s, 1H), 8.05 (s, 1H), 4.83 (bs, 2H), 3.81 (bs, 2H), 3.00 (s, 2H), 2.92 (q, J=7.6Hz, 2H), 1.29 (t, J = 7.4 Hz, 3H), 0.67 (s, 9H).

**Synthesis of 7-chloro-8-ethyl-10-(2-isobutoxyethyl)benzo[g]pteridine-2,4(3H,10H)-dione**

**(JG-2016)**

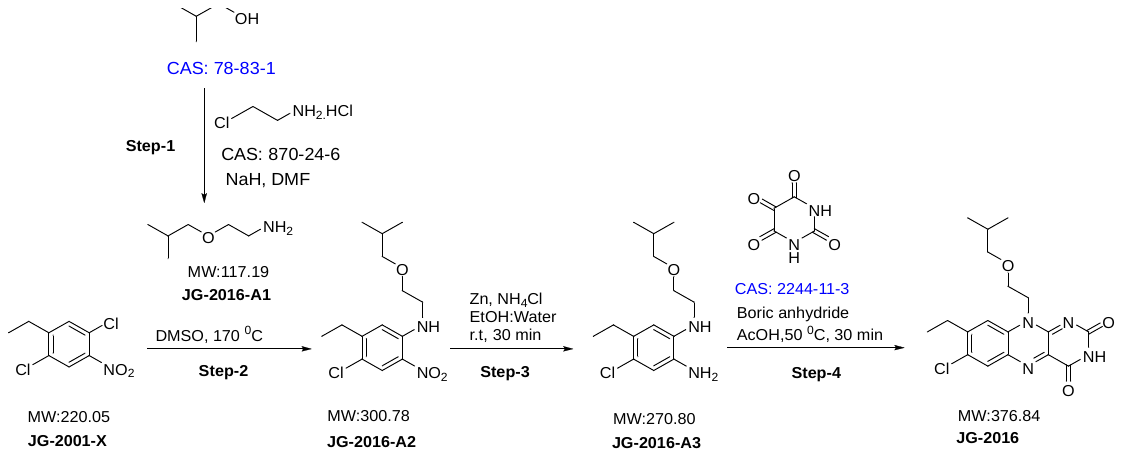

**Step-1: Synthesis of 2-isobutoxyethan-1-amine (JG-2016-A1)**

**Batch ID: STL-9-A708-JG-2016-A1-010**

To a solution of 2-methylpropan-1-ol (**CAS: 78-83-1**) (25g, 337.2 mmol, 1 eq) in DMF (250 mL/ 10V), 60 % NaH (67.45g, 1686.4 mmol, 5.0 eq) was added portion wise and stirred at 0°C temperature for 1h under nitrogen atmosphere. 2-chloroethan-1-amine hydrochloride (**CAS: 870-24-6**) (58.68 g, 505.9 mmol, 1.5 eq) was added portion wise to the reaction mixture and stirred at room temperature for 4h. After completion of reaction as indicated by TLC, the reaction mixture was poured into aqueous sodium chloride solution and extracted with ethyl acetate (3 x 2L). The combined organic layer was dried over anhydrous sodium sulphate and concentrated under vacuum up to 300 ml then washed with water (2 x 500 mL). The organic layer was dried over anhydrous sodium sulphate and concentrated under vacuum to afford yellow biphasic liquid (21.48 g, 54.34 %) crude material which was directly used in next step without further purification. LCMS: 118.2(M+1)

**Step-2: Synthesis of 4-chloro-5-ethyl-N-(2-isobutoxyethyl)-2-nitroaniline (JG-2016-A2)**

**Batch ID: STL-9-A708-JG-2016-A2-026**

To a solution of 1,4-dichloro-2-ethyl-5-nitrobenzene (**JG-2001-X**) (5.0 g, 22.72 mmol, 1.0 eq) in DMSO (5 mL), 2-isobutoxyethan-1-amine **(JG-2016-A1)** (13.31 g, 113.6 mmol, 5.0 eq) was added and stirred at 170 °C temperature for 5h. After completion of reaction as indicated by TLC, the reaction mixture was poured into ice cooled water (300 mL) and extracted with ethyl acetate (3 x 100 mL). The combined organic layer was dried over anhydrous sodium sulphate and concentrated under vacuum. The crude was further purified by column chromatography (4 % ethyl acetate/hexane) to afford yellow liquid (5.0 g, 73.16%) LCMS: 301.3, 303.3(M & M+2).

**Step-3: Synthesis of 4-chloro-5-ethyl-N1-(2-isobutoxyethyl)benzene-1,2-diamine (JG-2016-A3)**

**Batch ID: STL-9-A708-JG-2016-A3-037**

To a solution of 4-chloro-5-ethyl-N-(2-isobutoxyethyl)-2-nitroaniline (**JG-2016-A2**) (6.1 g, 20.28 mmol, 1.0 eq) in ethanol: water (8:2v) Zn Dust (10.60 g, 162.2 mmol, 8.0 eq) and NH4Cl (8.67g, 162.2 mmol, 8.0 eq) were added and stirred the reaction mixture at room temperature for 30 min at room temperature. TLC indicated completion of reaction. The reaction mixture was frittered through celite and filtrate was extracted with ethyl acetate (2 x 50 mL). The organic layer was dried over anhydrous sodium sulphate and concentrated under reduce pressure to afford yellow liquid (5.24g, 95.41 %). The crude product was used in next step without further purification. LCMS: 271.3, 273.2(M&M+2).

**Step-4: Synthesis of 7-chloro-8-ethyl-10-(2-isobutoxyethyl)benzo[g]pteridine-2,4(3H,10H)-dione (JG-2016)**

**Batch ID: STL-9-A708-JG-2016-038-e**

To a solution of 4-chloro-5-ethyl-N1-(2-isobutoxyethyl)benzene-1,2-diamine (**JG-2016-A3**) (5.24 g, 19.35 mmol, 1.0 eq) in AcOH (25 mL) alloxan monohydrate (**CAS: 2244-11-3**) (3.09 g, 19.35 mmol, 1.0 eq) and boric anhydride (2.69 g, 38.70 mmol, 2.0 eq) were added and stirred at 50 °C temperature for 30 min. The reaction mixture was poured into ice cooled water (100 mL) and extracted with ethyl acetate (3 x 50 mL). The reaction mixture was dried over anhydrous sodium sulphate Filtrate was concentrated under reduce pressure. The obtained crude product was further purified by column chromatography (80 % ethyl acetate/hexane) and pure fraction was concentrated under vacuum to afford yellow solid (2.8g, 38.40 %). LCMS: 377.3, 379.3(M&M+2). 1H NMR (400 MHz, DMSO) δ 11.44 (s, 1H), 8.18 (s, 1H), 8.03 (s, 1H), 4.80 (t, 2H), 3.78 (t, J = 4.8 Hz, 2H), 3.12 (d, J = 6.4 Hz, 2H), 2.91 (q, J = 7.2 Hz, 2H), 1.62 (m, 1H), 1.27 (t, J = 7.4 Hz, 3H), 0.70 (d, J = 6.7 Hz, 6H).

**Synthesis of tert-butyl 4-((2-(7-chloro-8-ethyl-2,4-dioxo-3,4-dihydrobenzo [g] pteridin-10 (2H)-yl) ethoxy) methyl) piperidine-1-carboxylate (JG-2009)**

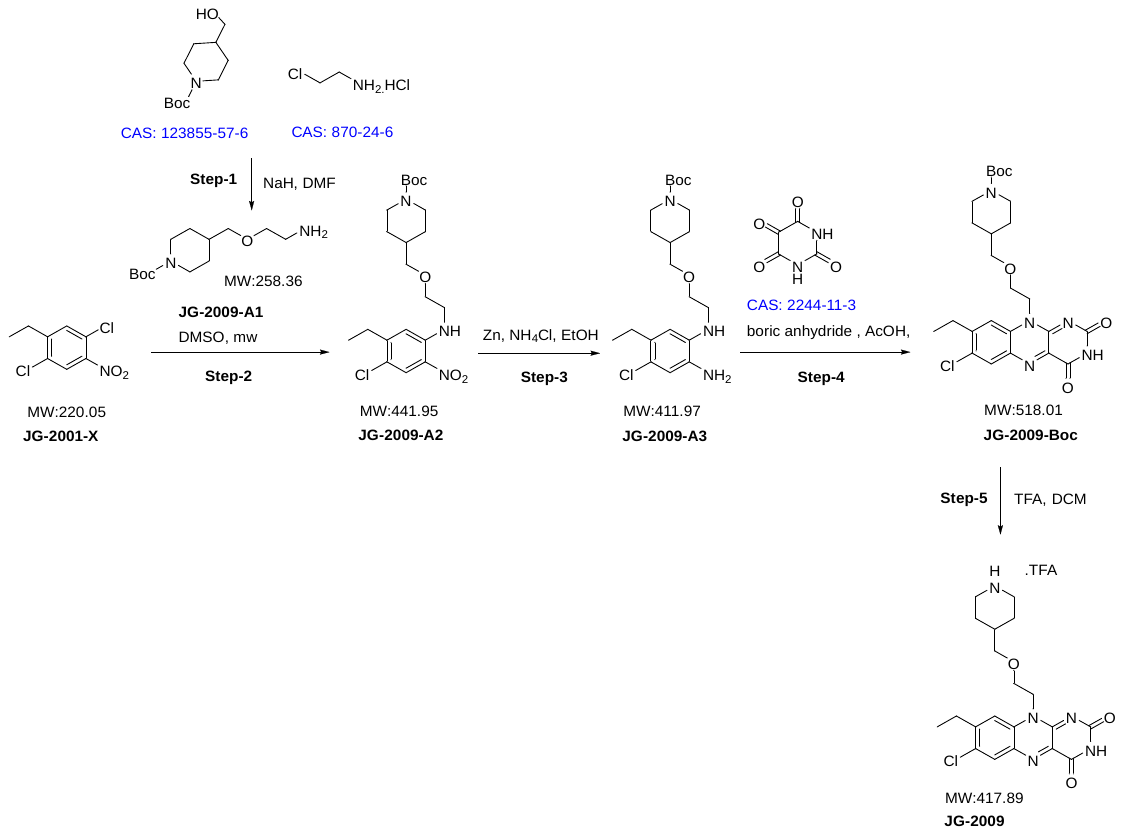

**Step-1: Synthesis of tert-butyl 4-((2-aminoethoxy)methyl)piperidine-1-carboxylate:**

**Batch ID: STL-7-A-574-JG-2009-A1-139**

To a solution of tert-butyl 4-(hydroxymethyl)piperidine-1-carboxylate (**CAS: 123855-57-6**) (0.5 g, 2.30 mmole, 1 eq) in DMF (20 mL), Sodium hydride (0.93 g, 23.20 mmol, 10.0 eq) was added portion wised at 0°C and the reaction mixture was allowed to stir for 1 hr at 0°C. 2-chloroethan-1-amine hydrochloride (**CAS: 870-24-6**) (0.54 g, 4.60 mmole, 2eq) was added slowely at 0°C and reaction mixtiure was further stirred at 0° to room temperature for 16 hr. After completion of reaction as indicated by TLC, the reaction mixture was quinched into ice cold water (20 mL) and extracted with ethyl acetate (3 x 20 mL). The combined organic layer was dried over anhydrous sodium sulphate and concentrated under vacuum to afford crude colourless liquid (0.93g, quantitative). Reaction was only monitored by TLC and used immediately in next step.

**Step-2: Synthesis of tert-butyl 4-((2-((4-chloro-5-ethyl-2-nitrophenyl)amino)ethoxy)methyl)piperidine-1-carboxylate**

**Batch ID: STL-7-A-574-JG-2009-A2-148-B**

To a solution of 1,4-dichloro-2-ethyl-5-nitrobenzene (**JG-2001-X**) (0.25 g, 1.10 mmole, 1 eq) in DMSO (4.0 mL), 4-((2-aminoethoxy)methyl)piperidine-1-carboxylate (**JG-2009-A1**) (0.9 g, 4.50 mmol, 4.0 eq) was added and heated at 190°C under microwave irradiation for 7 min. After completion of reaction as indicated by TLC, the reaction mixture was poured into water (10 mL) and extracted with ethyl acetate (3 x 10 mL). The combined organic layer was dried over anhydrous sodium sulphate and concentrated under vacuum to afford yellow liquid (0.27 g, quantitative), LCMS m/z 442.01

Note: Crude material was directly used in next step without further purification.

**Step-3: Synthesis of tert-butyl 4-((2-((2-amino-4-chloro-5-ethylphenyl) amino) ethoxy) methyl) piperidine-1-carboxylate**

**Batch ID: STL-7-A-574-JG-2009-A3-152-b**

To a solution of tert-butyl 4-((2-((4-chloro-5-ethyl-2-nitrophenyl)amino)ethoxy)methyl)piperidine-1-carboxylate **(JG-2009-A2)** (0.27 g, 6.10 mmole, 1.0 eq) in EtOH: water (8:2 mL), Zn Dust (0.261 g, 4.80 mmole, 8.0 eq) and NH4Cl (0.32 g, 4.80 mmole, 8.0eq) were added and stirred at 50 °C for 1 hr. After completion of reaction as indicated by TLC, the reaction mixture was filtered through Celite and filtrate was extracted with ethyl acetate (2 x 50 mL). The combined organic layer was dried over anhydrous sodium sulphate and concentrated under reduced pressure to obtain crude as sticky solid (0.24 g, quantitative), LCMS m/z 412.2 & 414.2 (M & M+2),

Note: Crude material was directly used in next step without further purification.

**Step-4: Syntheis of tert-butyl 4-((2-(7-chloro-8-ethyl-2,4-dioxo-3,4-dihydrobenzo [g] pteridin-10 (2H)-yl) ethoxy) methyl) piperidine-1-carboxylate:**

**Batch ID: STL-7-A-574-JG-2009-154-Boc**

To a solution of tert-butyl 4-((2-((2-amino-4-chloro-5-ethylphenyl) amino) ethoxy) methyl) piperidine-1-carboxylate (**JG-2009-A3**) (0.24 g, 0.58 mmol, 1.0 eq) in AcOH (5.0 mL), alloxan monohydrate **(CAS: 2244-11-3)** (0.093 g, 0.58 mmol, 1.0 eq) and boric anhydride (0.081 g, 1.10 mmol, 2.0 eq) were added and reaction mixture was stirred at 50 °C for 5 min. After completion of reaction as indicated by TLC the reaction mixture was quenched with ice cold water (30 mL) slowly and stirred for 20 min, the solid was precipitate out. The reaction mass was filtered over micro buckner funnel, solid was washed by ice cold water (5 ml × 2). Then solid was dried under reduced pressure to afford reddish brown solid. The crude was triturated with acetonitrile (10 ml×2) to obtain title compound as yellow solid (0.01 g, 3.0 %). LCMS: m/z 518.1 & 520.1 (M & M+2).

**Step-5: Synthesis of 2,2,2-trifluoroacetaldehyde compound with 7-chloro-8-ethyl-10-(2-(piperidin-4-ylmethoxy) ethyl) benzo [g] pteridine-2,4(3H,10H)-dione (JG-2009)**

**Batch ID: STL-7-A-574-JG-2009-174**

To a solution of tert-butyl 4-((2-(7-chloro-8-ethyl-2,4-dioxo-3,4-dihydrobenzo [g] pteridin-10 (2H)-yl) ethoxy) methyl) piperidine-1-carboxylate (**JG-2009-Boc**) (0.02 g, 0.038 mmol, 1 eq) in DCM (2 mL), Trifluoroacetic acid (0.011 mL, 0.15 mmol, 4.0 eq) was added at 0°C and the reaction mixture was allowed to stir for 2hr at room temperature. After completion of reaction as indicated by TLC, the reaction mixture was concentrated under vaccum and dried under reduced pressure. The crude was triturated with diethyl ether (10ml × 2) to obtained title compound as yellow solid (0.013 g, 81.25%). LCMS: m/z 418.1 & 420.0 (M & M+2); 1H NMR: (400Mz, DMSO) δ 11.49 (s, 1H), 8.40 (bs, 2H), 8.21 (s, 1H), 8.00 (s, 1H), 4.80 (t, 2H), 3.81 (t, 2H), 3.26(d,J=6.4Hz, 2H), 3.10-3.06 (m, 2H), 2.93 (q, J= 7.2Hz, 2H), 2.85-2.60 (m, 2H), 1.75-1.55 (m, 3H), 1.28 (t, J= 7.6Hz, 3H), 1.20-1.10(m, 2H)

The following compounds were made according to the procedure described in above examples using **JG-2001-X** and various amine side chains.

| **Target ID** | **Structure** | **Analytical data** |
| --- | --- | --- |
| **JG-2006** | 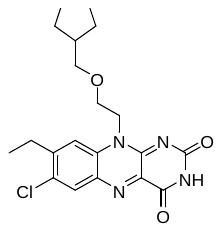 | LCMS m/z 403.1 & 405.1 (M & M+2); 1H NMR (400 MHz, DMSO) δ 11.44 (s, 1H), 8.19 (s, 1H), 8.03 (s, 1H), 4.80 (t, 2H), 3.79 (t, J = 4.8 Hz, 2H), 3.23 (d, J = 6.8 Hz, 2H), 2.91 (q, J = 7.2 Hz, 2H), 1.95-1.88 (m, 1H),1.49-1.38(m, 6H), 1.27 (t, J = 7.6 Hz, 3H), 1.04-0.95 (m, 2H). |
| **JG-2015** | 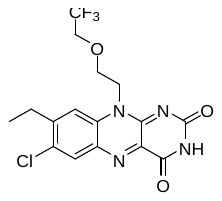 | LCMS m/z 403.0 & 405.1 (M & M+2); 1H NMR (400 MHz, DMSO) δ 11.46 (s, 1H), 8.20 (s, 1H), 8.00 (s, 1H), 4.85 (s, 2H), 4.09-4.00(m, 4H), 2.91 (q, J = 7.2 Hz, 2H), 1.27 (t, J = 7.6 Hz, 3H). |
| **JG-2008** | 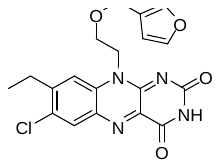 | LCMS m/z 401.0 & 403.0 (M & M+2); 1H NMR (400 MHz, DMSO) δ 11.44 (s, 1H), 8.19 (s, 1H), 7.99 (s, 1H), 7.55-7.53(m, 2H), 6.24(s, 1H),  4.80 (t, 2H), 4.33(s, 2H), 3.83(t, 2H), 2.89 (q, J = 7.2 Hz, 2H), 1.20 (t, J = 7.6 Hz, 3H). |
| **JG-2092** | 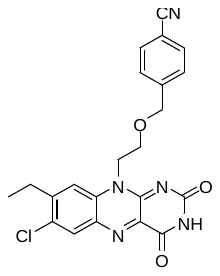 | LCMS m/z 436.0 & 438.0(M & M+2);  1H NMR (400 MHz, DMSO) δ 11.45 (s, 1H), 8.19 (s, 1H), 8.02 (s, 1H), 7.71(d,J=8.0Hz, 2H), 7.31(d,J=8.0Hz, 2H), 4.87 (bs, 2H), 4.56(s, 2H), 3.93(bs, 2H), 2.87 (q, J = 7.2 Hz, 2H), 1.15 (t, J = 7.2Hz, 3H). |

#### Synthesis of phenolic ether derivatives (JG-2082, JG-2067, JG-2074, JG-2076, JG-2077, JG-2078, JG-2086, JG-2087, JG-2088, JG-2034, JG-2079, JG-2080, JG-2081, JG-2083, JG-2089, JG-2090)

**Synthesis of 7-chloro-8-ethyl-10-(2-(p-tolyloxy)ethyl)benzo[g]pteridine-2,4(3H,10H)-dione (JG-2082)**

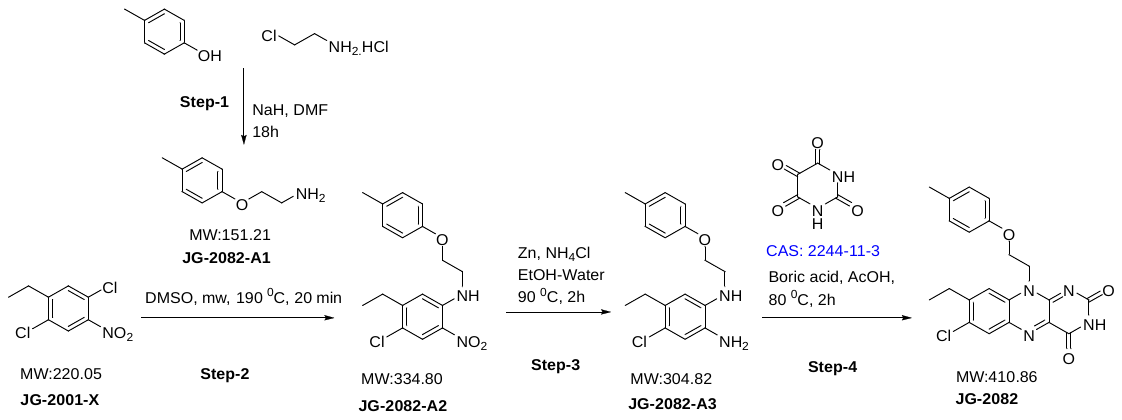

**Step-1 Synthesis of 2-(p-tolyloxy)ethan-1-amine**

**Batch ID: STL7-A574-JG-2082-A1-138**

To a solution of p-cresol (0.20 g, 1.85 mmol, 1 eq) and 2-chloroethylamine hydrochloride (0.21g, 1.85 mmol, 1.0 eq) in DMF (35 mL), sodium hydride (0.46g, 9.25 mmol, 5 eq) was added and the reaction mixture was stirred at room temperature for 18h at room temperature. The reaction mixture was quenched with ice water (25 mL) and extracted with ethyl acetate (3 x 50 mL). The combined organic phase was dried over anhydrous sodium sulphate and concentrated under vacuum. The crude product was purified by column chromatography to obtain title compound as white gummy solid. (0.12g, 42.8%). LCMS m/z 152.02(M+1)

**Step-2 Synthesis of 4-chloro-5-ethyl-2-nitro-N-(2-(p-tolyloxy)ethyl)aniline**

**Batch ID: STL7-A574-JG-2082-A2-141**

A solution of 2-(p-tolyloxy)ethan-1-amine (**JG-2082-A1**) (0.120g, 0.794 mmol, 1 eq) and 1,4-dichloro-2-ethyl-5-nitrobenzene (**JG-2001-X**) (0.21g, 0.794 mmol, 1.0 eq), in DMSO (2 mL) was heated at 190°C under microwave irradiation for 20 min. The reaction mixture was poured into water (25 mL) and extracted with ethyl acetate (2 x 25 mL). The organic phase was dried over anhydrous sodium sulphate and concentrated under vacuum. The crude was purified by column chromatography (20% ethyl acetate/hexane) to obtain title compound as yellow solid. (0.100g, 37.70%). LCMS m/z 335.13(M+1)

**Step-3 Synthesis of 4-chloro-5-ethyl-N1-(2-(p-tolyloxy)ethyl)benzene-1,2-diamine**

**Batch ID: STL7-A574-JG-2082-A3-142**

To a solution of 4-chloro-5-ethyl-2-nitro-N-(2-(p-tolyloxy)ethyl)aniline (**JG-2082-A2**) (0.10g, 0.299 mmol, 1 eq) in ethanol (10 mL) and water (2 mL), Zinc dust (0.156g, 2.39 mmol, 8 eq) and ammonium chloride (0.128 g, 2.39 mmol, 8 eq) were added and the reaction mixture was stirred at 90°C 2h. After completion of reaction as indicated by TLC, the reaction mixture was filtered through celite and washed with ethyl acetate (50ml). The filtrate was washed with water (50 ml) and the organic phase was dried over anhydrous sodium sulphate. The combined organic layer was concentrated under vacuum to afford crude as yellow liquid. (0.080g, 88.01%). LCMS m/z 305.1 & 307.1(M & M+2).

**Step-4 Synthesis of 7-chloro-8-ethyl-10-(2-(p-tolyloxy)ethyl)benzo[g]pteridine-2,4 (3H,10H)-dione**

**Batch ID: STL7-A574-JG-2082-A4-143**

To a solution of 4-chloro-5-ethyl-N1-(2-(p-tolyloxy)ethyl)benzene-1,2-diamine (JG-2082-A3) (0.080g, 0.29 mmol, 1.0 eq) in AcOH (30 mL) (**CAS: 2244-11-3**) (0.042g, 0.296 mmol, 1.0 eq) and Boric anhydride (0.041g, 0.592 mmol, 2.0 eq) were added and the reaction mixture was stirred at 80 °C for 2h. The reaction mixture was quenched with ice cooled water (35 mL) and extracted with ethyl acetate (2 x 50 mL). The combined organic phase was dried over anhydrous sodium sulphate and concentrated under reduce pressure. The obtained crude product was further purified by column chromatography (100 % ethyl acetate) to afford yellow solid (0.010g, 8.24%) LCMS m/z 410.9 & 413.2(M & M+2); 1H NMR (400 MHz, DMSO): δ: 11.45 (s, 1H), 8.21 (s, 1H), 8.10(s, 1H), 7.05(d, J=8.4Hz, 2H), 6.75(d, J=8.0Hz, 2H), 4.99 (t, 2H), 4.37 (t, 2H), 2.92 (q , 2H), 2.19(s, 3H), 1.28 (t, J= 7.6Hz, 3H).

The following compounds were made according to the procedure described for JG-2082 using

**JG-2001-X** and the respective amine.

| **Target ID** | **Structure** | **Analytical data** |
| --- | --- | --- |
| **JG-2067** | 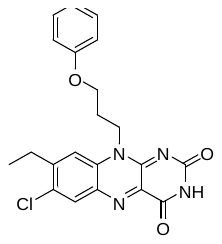 | LCMS m/z 411.0 & 413.0(M &M+2); 1H NMR (400 MHz, DMSO): δ: 11.42 (s, 1H), 8.22 (s, 1H), 7.90 (s, 1H), 7.29-7.25 (m, 2H), 6.94-6.85 (m, 3H), 4.76 (bs, 2H), 4.11 (t, 2H), 2.82 (q, J=7.6Hz, 2H), 2.24 (bs, 2H), 1.16 (t, J=7.2Hz, 3H). |
| **JG-2074** | 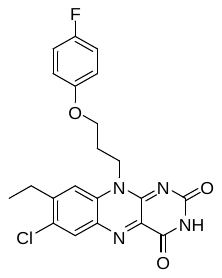 | LCMS m/z 429.0 & 431.0(M &M+2); 1H NMR (400MHz, DMSO, ppm) δ 11.43 (s, 1H), 8.20 (s, 1H), 7.86 (s, 1H), 7.08 (t, J = 8.8 Hz, 2H), 6.84 (m, J = 4 Hz, 2H), 4.75 (t, 2H), 4.07 (t, J = 5.6 Hz, 2H), 2.81 (q, J = 7.6 Hz, 2H), 2.21 (m, J = 6.4 Hz, 2H), 1.14 (t, J = 7.2 Hz, 3H) |
| **JG-2076** | 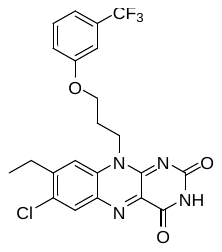 | LCMS m/z 479.0 & 481.0(M &M+2); 1H NMR (400 MHz, DMSO): δ: 11.43 (s, 1H), 8.21 (s, 1H), 7.91 (s, 1H), 7.52-7.48 (m, 1H), 7.28-7.11 (m, 3H), 4.77 (t, 2H), 4.20(bs, 2H), 2.80 (q ,J= 7.6Hz, 2H), 2.30-2.20 (m, 2H), 1.12 (t, J= 7.2Hz, 3H). |
| **JG-2077** | 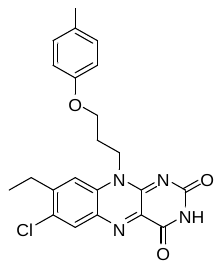 | LCMS m/z 425.0 & 427.0(M &M+2); 1H NMR (400 MHz, DMSO): δ: 11.42 (s, 1H), 8.21 (s, 1H), 7.88 (s, 1H), 7.06 (d,J= 8.0Hz, 2H), 6.76 (d, J= 8.4 Hz, 2H), 4.74 (t, 2H), 4.08 (t, J=5.6Hz, 2H), 2.82 (q ,J= 7.6Hz, 2H),2.21(s, 3H), 2.20-2.15(m, 2H), 1.17 (t, J= 7.2Hz, 3H). |
| **JG-2078** | 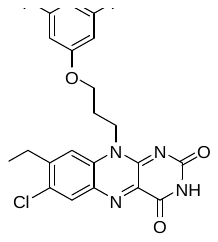 | LCMS m/z 447.0 & 449.0(M &M+2); 1H NMR (400 MHz, DMSO): δ: 11.44 (s, 1H), 8.23 (s, 1H), 7.89 (s, 1H), 6.80-6.75 (m, 1H), 6.64-6.61 (m, 2H), 4.74 (t, 2H), 4.15 (t, J=5.6Hz, 2H), 2.85 (q, J= 7.6Hz, 2H), 2.24-2.21(m, 2H), 1.18 (t, J= 7.6Hz, 3H). |
| **JG-2086** | 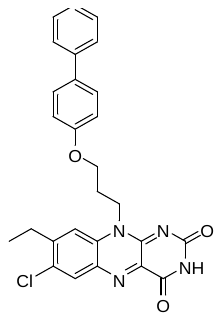 | LCMS m/z 487.0 & 489.0(M &M+2); 1H NMR (400 MHz, DMSO): δ: 11.44 (s, 1H), 8.21 (s, 1H), 7.90 (s, 1H), 7.61-7.56(m, 4H), 7.41 (dd, J=7.6Hz, 2H), 7.30 (t, J= 7.2Hz, 1H), 6.95(d, J= 8.4Hz, 2H), 4.75 (t, 2H), 4.17 (t, J=5.6Hz, 2H), 2.82 (q ,J= 7.2Hz, 2H), 2.28-2.25(m, 2H), 1.16 (t, J= 7.2Hz, 3H). |
| **JG-2087** | 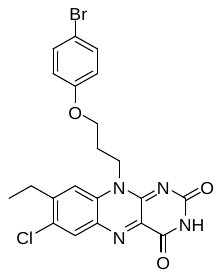 | LCMS m/z 490.9 & 492.9(M &M+2); 1H NMR (400 MHz, DMSO): δ: 11.43 (s, 1H), 8.21 (s, 1H), 7.88 (s, 1H), 7.43 (d,J= 8.4Hz, 2H), 6.84 (d, J= 8.8 Hz, 2H), 4.73 (bs, 2H), 4.11 (t, 2H), 2.82 (q ,J= 7.2Hz, 2H), 2.30-2.25(m, 2H), 1.17 (t, J= 7.2Hz, 3H). |
| **JG-2088** | 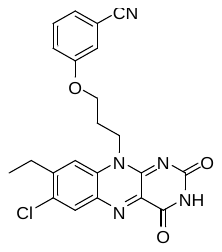 | LCMS m/z 436.1 & 438.1(M &M+2); 1H NMR (400 MHz, DMSO): δ: 11.44 (s, 1H), 8.22 (s, 1H), 7.90 (s, 1H), 7.49-7.38 (m, 2H), 7.33 (s, 1H), 7.20(dd,J=8.0 & 1.6Hz, 1H), 4.75 (t, 2H), 4.19(t, J=5.6Hz, 2H), 2.83 (q ,J= 7.6Hz, 2H), 2.26-2.23 (m, 2H), 1.16 (t, J= 7.2Hz, 3H). |
| **JG-2034** | 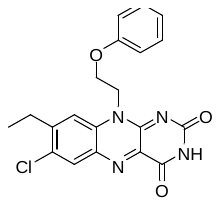 | LCMS m/z 397.50 & 399.62(M &M+2); 1H NMR (400 MHz, DMSO): δ: 11.46 (s, 1H), 8.22 (s, 1H), 8.13 (s, 1H), 7.26(t, J= 8Hz, 2H), 6.94-6.85 (m, 3H), 5.02 (t, 2H), 4.43(bs, 2H), 2.93 (q ,J= 7.2Hz, 2H), 1.29 (t, J= 7.6Hz, 3H). |
| **JG-2079** | 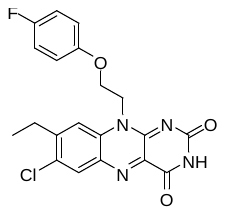 | LCMS m/z 415.0 & 417.0(M &M+2); 1H NMR (400 MHz, DMSO): δ: 11.45 (s, 1H), 8.21 (s, 1H), 8.11(s, 1H), 7.10-7.06(m, 2H), 6.88-6.84(m, 2H), 5.00 (t, 2H), 4.39 (t, 2H), 2.92 (q ,J= 7.2Hz, 2H), 1.26 (t, J= 7.6Hz, 3H). |
| **JG-2080** | 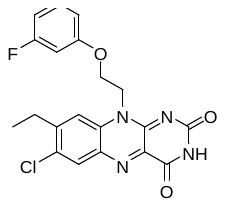 | LCMS m/z 415.0 & 417.1(M &M+2); 1H NMR (400 MHz, DMSO): δ: 11.46 (s, 1H), 8.21 (s, 1H), 8.09(s, 1H), 7.28-7.25(m, 1H), 6.76-6.67(m, 3H), 5.01 (t, 2H), 4.44 (t, 2H), 2.92 (q ,J= 7.2Hz, 2H), 1.26 (t, J= 7.6Hz, 3H). |
| **JG-2081** | 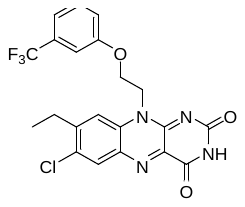 | LCMS m/z 465.0 & 467.0(M &M+2); 1H NMR (400 MHz, DMSO): δ: 11.46 (s, 1H), 8.21 (s, 1H), 8.13(s, 1H), 7.48(t,J=7.6Hz, 1H), 7.27(d, J=7.2Hz,, 1H), 7.17(d, J=8Hz, 1H), 7.09(s, 1H), 5.04 (t, 2H), 4.51 (t, 2H), 2.91 (q ,J= 7.2Hz, 2H), 1.28 (t, 3H). |
| **JG-2083** | 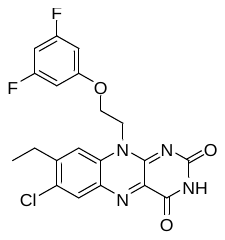 | LCMS m/z 433.0 & 435.0(M &M+2); 1H NMR (400 MHz, DMSO): δ: 11.46 (s, 1H), 8.21 (s, 1H), 8.05(s, 1H), 6.79-6.77(m,1H), 6.65-6.63(m, 2H), 5.01 (t, 2H), 4.46 (t, 2H), 2.92 (q, J=7.2Hz, 2H), 1.27 (t, J= 7.6Hz, 3H). |
| **JG-2089** | 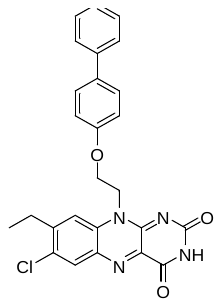 | LCMS m/z 473.2 & 475.1(M &M+2); 1H NMR (400 MHz, DMSO): δ: 11.45 (s, 1H), 8.21 (s, 1H), 8.13(s, 1H), 7.57-7.55(m,4H), 7.42-7.27(m, 3H), 6.95(d, J= , 2H), 5.04 (t, 2H), 4.48 (t, 2H), 2.93 (q, J=7.2Hz, 2H), 1.29 (t, J= 7.2Hz, 3H). |
| **JG-2090** | 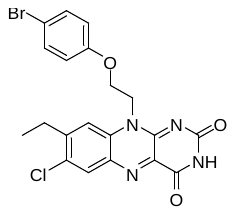 | LCMS m/z 474.8 & 476.9 (M& M+2); 1H NMR (400 MHz, DMSO): δ: 11.46 (s, 1H), 8.21 (s, 1H), 8.11 (s, 1H), 7.42 (d, J=8.8Hz, 2H), 6.83 (d, J=8.8Hz, 2H), 5.00 (t, 2H), 4.41 (t, 2H), 2.93 (q, 2H), 1.28 (t, J=7.6Hz, 3H). |

#### Synthesis of amine sidechain derivatives (JG-2025, JG-2052, JG-2051, JG-2053)

**Synthesis of 10-(2-aminoethyl)-7-chloro-8-ethylbenzo[g]pteridine-2,4(3H,10H)-dione 2,2,2-trifluoroacetate (JG-2025)**

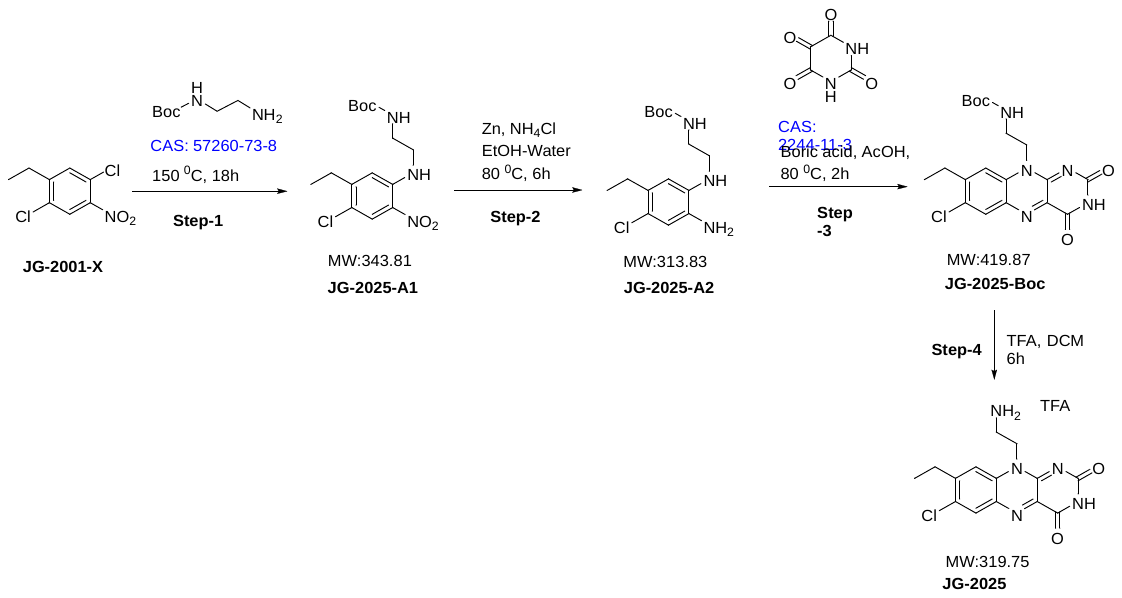

**Step-1 Synthesis of tert-butyl (2-((4-chloro-5-ethyl-2-nitrophenyl)amino)ethyl)carbamate**

**Batch ID: STL7-A574-JG-2025-A1-060**

1,4-dichloro-2-ethyl-5-nitrobenzene (**JG-2001-X**) (3.0 g, 13.63 mmol, 1 eq) and tert-butyl (2-aminoethyl)carbamate (**CAS: 57260-73-8**) (2.18 g, 13.63 mmol, 1.0 eq) were stirred neat at 150°C temperature. After 18h of stirring TLC analysis confirms consumption of SM, the reaction mixture was slowly poured into water (100 mL) and extracted with ethyl acetate (2 x 75 mL). The organic phase was dried over anhydrous sodium sulphate and concentrated under vacuum. The crude material was purified by column chromatography (30 % ethyl acetate/hexane) as yellow solid (3.01g, 64.3 %). LCMS: MS: m/z 344.6 & 346.6(M+1)

**Step-2 Synthesis of tert-butyl (2-((2-amino-4-chloro-5-ethylphenyl)amino)ethyl)carbamate**

**Batch ID: STL7-A574-JG-2025-A2-066**

To a solution of tert-butyl (2-((4-chloro-5-ethyl-2-nitrophenyl)amino)ethyl)carbamate(**JG-2025-A1**) (3.0g, 8.72 mmol) in ethanol: water (8:2V, 40 mL), Zn Dust (4.56g, 69.8 mmol, 8.0 eq) and NH4Cl (3.73 g, 69.8 mmol, 8.0 eq) were added and the reaction mixture was stirred at 80°C. Reaction progress was further monitored using TLC (mobile phase: 20% ethyl acetate in hexane), after 6h of stirring TLC analysis confirms consumption of SM. The reaction mixture was filtered through celite and filtrate was poured into water (75 mL) and extracted with ethyl acetate (2 x 100 mL). The combined organic phase was dried over anhydrous sodium sulphate and concentrated under reduce pressure. The obtained crude product material was directly used in next step (2.8g, Quantitative). LCMS: m/z 314.6 & 316.6(M & M+2).

**Step-3 Synthesis of tert-butyl (2-(7-chloro-8-ethyl-2,4-dioxo-3,4-dihydrobenzo[g]pteridin-10(2H)-yl)ethyl)carbamate**

**Batch ID: STL7-A574-JG-2025-A4-067**

To a solution of tert-butyl (2-((2-amino-4-chloro-5-ethylphenyl)amino)ethyl)carbamate (**JG-2025-A2**) (2.75g, 8.70 mmol, 1.0 eq) in AcOH (30 mL), alloxan monohydrate(**CAS: 2244-11-3**) (1.54 g, 8.70 mmol, 1.0 eq) and boric anhydride (1.36 g, 17.53mmol, 2.0 eq) were added. And the reaction mixture was stirred at 80 °C temperature. Reaction progress was further monitored using TLC (mobile phase: 100 % ethyl acetate in hexane), after 2h of stirring TLC analysis confirms consumption of SM. The reaction mixture was slowly poured in to ice cooled water (100 mL) extracted with ethyl acetate (2 x 100 mL). The organic phase was dried over anhydrous sodium sulphate and concentrated under reduced pressure. The obtained crude product was further purified by column chromatography (100 % ethyl acetate) to obtain yellow solid (2.15g, 58.9 %) LCMS: m/z 420.80 & 422.8 (M& M+2)

**Step-4 Synthesis of 10-(2-aminoethyl)-7-chloro-8-ethylbenzo[g]pteridine-2,4(3H,10H)-dione 2,2,2-trifluoroacetate**

**Batch ID: STL7-A574-JG-2025-A4-069**

To a solution of tert-butyl (2-(7-chloro-8-ethyl-2,4-dioxo-3,4-dihydrobenzo[g]pteridin-10(2H)-yl)ethyl)carbamate (**JG-2025-Boc**) (2.0g, 4.77 mmol, 1.0 eq) in DCM (30 mL), TFA (10.86 g, 95.2mmol, 20.0 eq) was added and the reaction mixture was stirred at room temperature for 6h. Reaction progress was further monitored using TLC (100% ethyl acetate/ hexane). The reaction mixture was concentrated under vacuum, triturated with MTBE and diethyl ether and again dried under vacuum to afford product as yellow solid (1.85g, 90.2 %). LCMS: m/z 320.51 (M+1); 1H NMR (400 MHz, DMSO): δ: 11.58 (s, 1H), 8.29 (s, 1H), 7.95 (bs, 4H), 4.87 (t, 2H), 3.23 (bs, 2H), 2.96 (q, 2H), 1.31 (t, 3H)

**Synthesis of N-(2-(7-chloro-8-ethyl-2,4-dioxo-3,4-dihydrobenzo[g]pteridin-10(2H)-yl)ethyl)acetamide (JG-2052)**

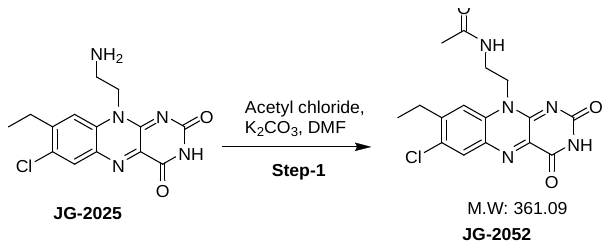

**Batch ID: STL7-A574-JG-2025-A5-036**

To a solution of 10-(2-aminoethyl)-7-chloro-8-ethylbenzo[g]pteridine-2,4(3H,10H)-dione **(JG-2025)** (0.050g, 0.156 mmol, 1 eq) in DMF (2 mL), potassium carbonate (0.064g, 0.468 mmol, 3 eq) and acetyl chloride (0.014 mg, 0.188 mmol, 1.2 eq) were added and the reaction mixture was stirred at room temperature for 18h. The reaction mixture was poured into water (25 mL) and extracted with ethyl acetate (75 mL). The organic phase was dried over anhydrous sodium sulphate and concentrated. The crude material was purified by column chromatography (5 % methanol/dichloromethane) to obtain title compound as yellow solid (2 mg, 3.55 %). LCMS: m/z 362.1 & 364.3 (M& M+2); 1H NMR (400 MHz, DMSO): δ: 11.49 (s, 1H), 8.24 (s, 1H), 8.16 (t, 1H), 8.06 (s, 1H), 4.62 (t, 2H), 3.43 (t, 2H), 2.94 (q, 2H), 1.71 (s, 3H), 1.32 (t, 3H)

**Synthesis of N-(2-(7-chloro-8-ethyl-2,4-dioxo-3,4-dihydrobenzo[g]pteridin-10(2H)-yl)ethyl) benzamide (JG-2051)**

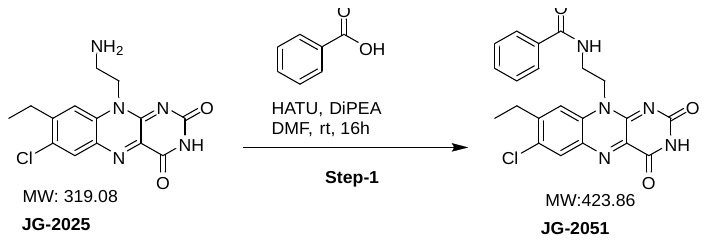

**Batch ID: STL7-A574-JG-2025-A5-040**

To a solution of 10-(2-aminoethyl)-7-chloro-8-ethylbenzo[g]pteridine-2,4(3H,10H)-dione(**JG-2025**) (0.076g, 0.63 mmol, 1 eq) in DMF (2 mL), DIPEA (0.30 mL, 1.88 mmol, 3eq), HATU (0.36g, 0.94 mmol, 1.5eq) and benzoic acid (0.20 g, 0.63 mmol, 1eq) were added. The reaction mixture was stirred at room temperature for 18h. The reaction mixture was poured into water (25 mL) and extracted with ethyl acetate (2 x 25 mL). The combined organic phase was dried over anhydrous sodium sulphate and concentrated under vacuum. The crude material was purified by column chromatography (100 % ethyl acetate) to obtain title compound as yellow solid (0.030g, 11.2 %). LCMS m/z 424.6 & 426.6(M & M+2); 1H NMR (400 MHz, DMSO): δ: 11.53 (s, 1H), 8.65 (t, 1H), 8.22 (s, 1H), 8.01 (s, 1H), 7.61 (d, J=7.2Hz, 2H), 7.51-7.37 (m, 3H), 4.79 (t, 2H), 3.79-3.72 (m, 2H), 2.76 (q, 2H), 1.14 (t, 3H).

The following compound was made according to the procedure described for **JG-2051** using

**JG-2025** and isobutyric acid.

| **Target ID** | **Structure** | **Analytical data** |
| --- | --- | --- |
| **JG-2053** | 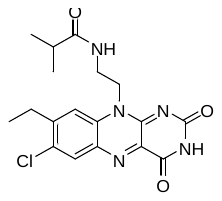 | LCMS m/z 390.7 & 392.7(M & M+2); 1H NMR (400 MHz, DMSO): δ: 11.50 (s, 1H), 8.24 (s, 1H), 8.03 (s, 1H), 7.97 (t, 1H), 4.67 (t, 2H), 3.55 (m, 2H), 2.94 (q, J=7.6Hz, 2H), 2.14(hept, 1H), 1.34 (t,J=7.2Hz, 3H), 0.81 (d,J=6.8Hz, 6H). |

#### Synthesis of sulfonamide sidechain derivatives (JG-2057, JG-2061, JG-2060, JG-2062, JG-2063, JG-2064, JG-2065)

**Synthesis of N-(2-(7-chloro-8-ethyl-2,4-dioxo-3,4-dihydrobenzo[g]pteridin-10(2H)-yl)ethyl) benzene sulfonamide (JG-2057)**

**Batch ID: STL7-A574-JG-2025-A5-055**

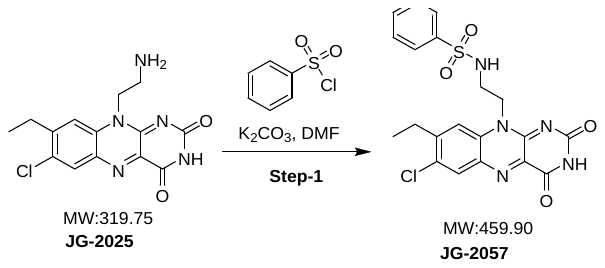

**Batch ID: STL7-A574-JG-2025-A5-055**

To a solution of 10-(2-aminoethyl)-7-chloro-8-ethylbenzo[g]pteridine-2,4(3H,10H)-dione (**JG-2025**) (0.100g, 0.313 mmol, 1 eq) in DMF (2 mL), potassium carbonate (0.13g, 0.939 mmol, 3 eq) and benzenesulfonyl chloride (0.083g, 0.188 mmol, 1.2 eq) were added and stirred for 18h at room temperature. The reaction mixture was poured into water (50 mL) and extracted with ethyl acetate (2 x 25 mL). The combined organic phase was dried over anhydrous sodium sulphate and concentrated under vacuum. The crude material was purified by column chromatography (100 % ethyl acetate) as yellow solid (0.012g, 8.35 %). LCMS: m/z 460.78 (M+1); 1H NMR (400 MHz, DMSO): δ: 11.50 (s, 1H), 8.22 (s, 1H), 7.97 (t, 1H), 7.91 (s, 1H), 7.72 (d, J= 7.6Hz, 2H), 7.62-7.51(m, 3H), 4.66 (t, 2H), 3.23-21 (m, 2H), 2.93 (q, 2H), 1.32 (t, 3H).

The following compounds were made according to the procedure described for **JG-2057** using **JG-2025** and respective sulfonyl chloride.

| **Target ID** | **Structrure** | **Analytical data** |
| --- | --- | --- |
| **JG-2062** | 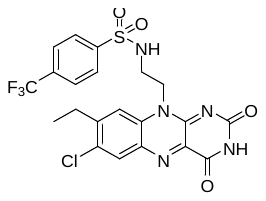 | LCMS: m/z 428.2 & 430.1 (M & M+2); 1H NMR (400 MHz, DMSO): δ: 11.51 (s, 1H),8.20(bs, 1H), 8.18 (s, 1H), 7.95-7.89 (m, 5H), 4.66 (t, 2H), 3.29 (t, 2H), 2.90 (q, J=7.2Hz, 2H), 1.30 (t, J=7.6Hz, 3H). |
| **JG-2063** | 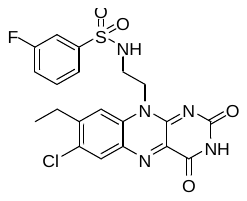 | LCMS: m/z 478.1 & 480.1 (M & M+2); 1H NMR (400 MHz, DMSO): δ: 11.50 (s, 1H),8.20(s, 1H), 8.10 (bs, 1H), 7.90 (s, 1H), 7.58-7.44(m, 4H), 4.66 (t, 2H), 3.27 (t, 2H), 2.92 (q, J=7.6Hz, 2H), 1.31 (t, J=7.6Hz, 3H). |
| **JG-2064** | 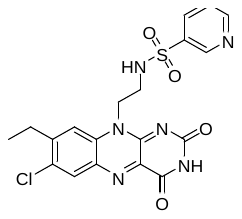 | LCMS: m/z 461.1 & 463.2 (M & M+2); 1H NMR (400 MHz, DMSO): δ: 11.50 (s, 1H), 8.86(s, 1H), 8.77 (d,J=3.6Hz, 1H), 8.30 (bs, 1H), 8.21(s, 1H), 8.13-8.11(m, 1H), 7.96(s, 1H), 7.58-7.54(m, 1H), 4.66 (t, 2H), 3.52-3.40 (m, 2H), 2.92 (q, J=7.2Hz, 2H), 1.31 (t, J=7.2Hz, 3H). |

**Synthesis of N-(2-(7-chloro-8-ethyl-2,4-dioxo-3,4-dihydrobenzo[g]pteridin-10(2H)-yl)ethyl)-2-methyl propane-1-sulfonamide (JG-2061)**

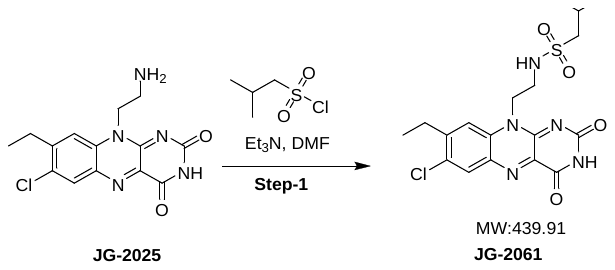

**Batch ID: STL7-A574-JG-2025-A5-087**

To a solution of 10-(2-aminoethyl)-7-chloro-8-ethylbenzo[g]pteridine-2,4(3H,10H)-dione (**JG-2025**) (0.050g, 0.156 mmol, 1 eq) in DMF (3 mL), triethylamine (0.06 mL, 0.468 mmol, 3eq) and isobutylsulfonyl chloride (0.025g, 0.156 mmol, 1eq) were added and the reaction mixture was stirred at room temperature for 18h. The reaction mixture was poured into water (50 mL) and extracted with ethyl acetate (2 x 25 mL). The combined organic phase was dried over anhydrous sodium sulphate and concentrated under vacuum. The crude material was purified by column chromatography (100 % ethyl acetate) to obtain title compound as yellow solid (0.012 g, 8.35 %). LCMS: m/z 440.3 & 442.3 (M & M+2); 1H NMR (400 MHz, DMSO): δ: 11.50 (s, 1H), 8.24 (s, 1H), 7.97 (s, 1H), 4.68 (t, 2H), 3.39 (t, 2H), 2.95-2.90 (m, 4H), 2.00 (hept, 1H), 1.32 (t, J=7.6Hz, 3H), 0.97 (d, J=6.4Hz, 6H).

The following compounds were made according to the procedure described for **JG-2061** using **JG-2025** and respective sulfonyl chloride.

| **Target ID** | **Structure** | **Analytical data** |
| --- | --- | --- |
| **JG-2060** | 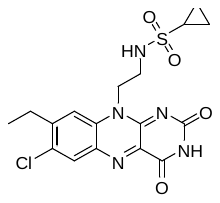 | LCMS: m/z 424.0 & 426.0 (M & M+2); 1H NMR (400 MHz, DMSO): δ: 11.49(s, 1H), 8.23 (s, 1H), 7.95 (s, 1H), 7.33 (t, 1H), 4.70 (t, 2H), 3.50-3.40 (m, 2H), 2.92 (q, 2H), 2.60-2.40(m, 1H), 1.30 (t, 3H), 0.91-0.86(m, 4H). |
| **JG-2065** | 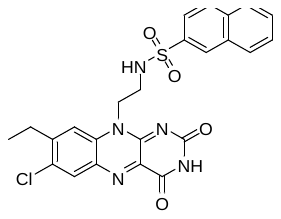 | LCMS: m/z 510.2 & 512.2 (M& M+2); 1H NMR (400 MHz, DMSO): δ: 11.47 (s, 1H), 8.34 (s, 1H), 8.06-7.97(m, 5H) 7.97 (s, 1H), 7.68-7.63(m, 3H), 4.64 (t, 2H), 3.39-3.30 (m, 2H), 2.88 (q, J=7.2Hz, 2H), 1.27 (t, J=7.2Hz, 3H). |

#### Synthesis of 2001-B1

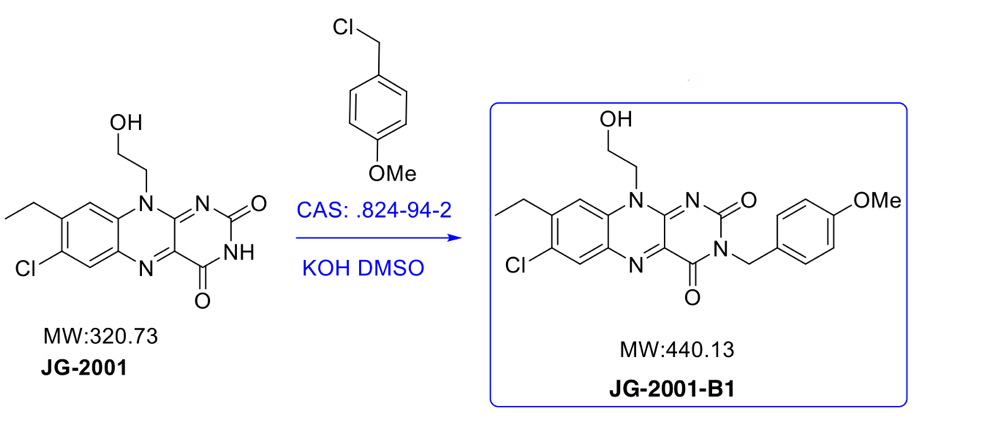

JG-2001 (100 mg, 1 eq) was mixed with CAS 824-94-2 (3 eq) and KOH (4 eq) in DMSO and reacted at room temperature for 2 hours. Product (JG-2001-B1) was purified by column chromatography (yield 25 mg).

**7-chloro-8-ethyl-10-(2-hydroxyethyl)-3-(4-methoxybenzyl)benzo[*g*]pteridine-2,4(3*H*,10*H*)-dione (JG-2001-B1)-**

^1^H-NMR (400 MHz, DMSO, ppm) δ 8.25 (s, 1H), 8.08 (s, 1H), 7.32 (d, J = 8.4 Hz, 2H), 6.86 (d, J = 8.4 Hz, 2H), 5.02 (s, 2H), 4.96 (t, J = 6.0 Hz, 1H), 4.73 (s, 2H), 3.82 (d, J = 5.6 Hz, 2H), 3.71 (s, 3H), 2.94 (d, J = 7.6 Hz, 2H), 1.29 (t, J = 7.6 Hz, 3H)

LCMS, calc’d 440.13 m/z; found 441.79 m/z

#### Synthesis of JG-2011

**Part-1: Amine sidechain synthesis**

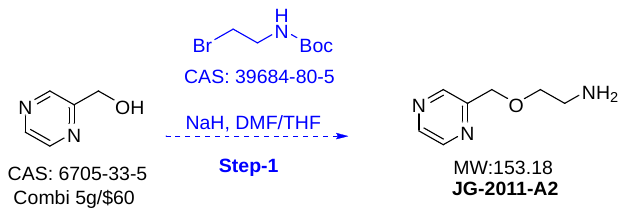

CAS 6705-33-5 (100 mg, 1 eq) was mixed with NaH (5 eq) and CAS 39684-80-5 (2 eq) in THF, then chilled to 0 ^0^C on ice and allowed to warm to room temperature for 3 hours. Product formation (JG-2011-A2) was confirmed by TLC and LCMS, purified by reverse phase chromatography (yield 590 mg).

**Part-2: Synthesis of JG-2011 from amine sidechain**

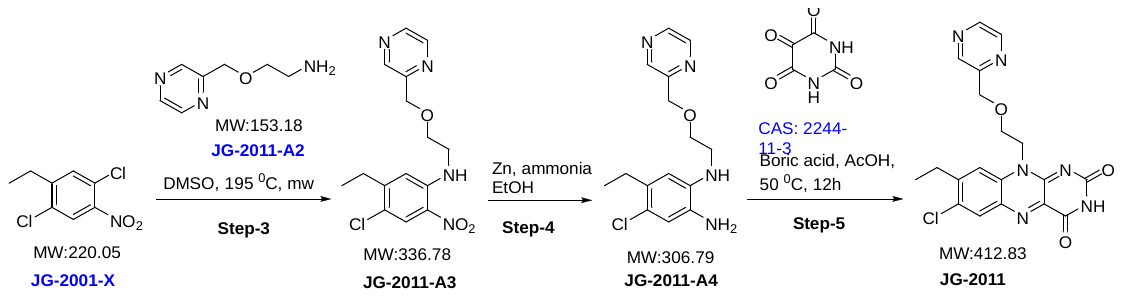

JG-2001-X (238 mg, 1 eq) was combined with JG-2011-A2 (3 eq) in DMSO, then heated to 190 ^0^C for 15 minutes in a microwave. Product formation (JG-2011-A3) was observed by TLC and LCMS, and purified by column chromatography (yield 63 mg). Then, JG-2011-A3 (63 mg, 1 eq) was combined with Zn (8 eq) and NH_4_Cl (8 eq) in a 8:2 mixture of ethanol:water, reacted for 30 minutes at 80 ^0^C and product formation (JG-2011-A4) was confirmed by TLC, LCMS, purified by column chromatography (yield 54 mg). Next, JG-2011-A4 (54 mg, 1 eq) was mixed with alloxan, boric acid and acetic acid, heated to 80 ^0^C for 15 and product formation (JG-2011) was observed, confirmed by TLC, purified by column chromatography (yield 7 mg).

**7-chloro-8-ethyl-10-(2-(pyrazin-2-ylmethoxy)ethyl)benzo[*g*]pteridine-2,4(3*H*,10*H*)-dione**

**(JG-2011)-**

1H NMR (400 MHz, DMSO, ppm) δ 11.46 (s, 1H), 8.52 (s, 2H), 8.42 (s, 1H), 8.20 (s, 1H), 8.08 (s, 1H), 4.90 (s, 2H), 4.63 (s, 2H), 4.02 (s, 2H), 2.86 (s, 2H), 1.16 (s, 3H)

LCMS, calc’d 412.83 m/z; found 413.64 m/z

#### Synthesis of JG-2031

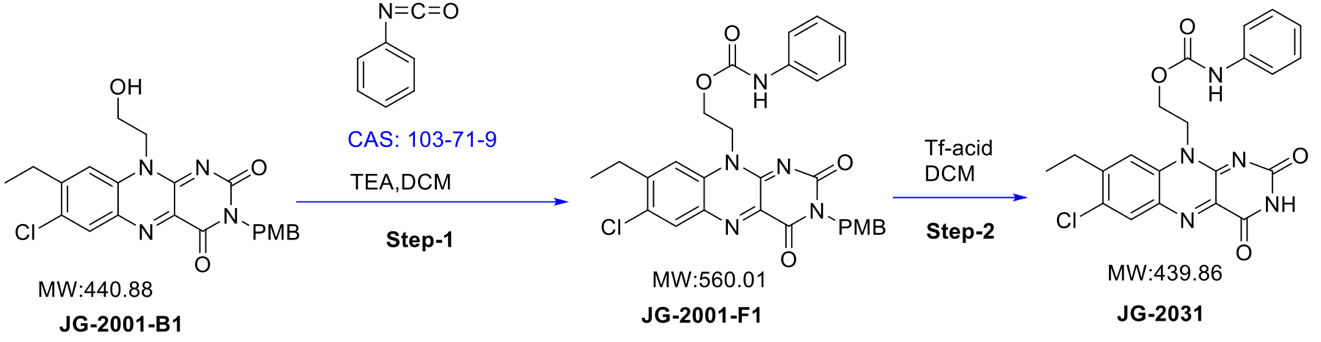

JG-2001-B1 (80, 1 eq) was mixed with DCM + TEA (3 eq) and CAS 103-71-0 (1.5 eq) for 1 hour at room temperature. Product formation (JG-2001-F1) was observed by TLC and LCMS, and purified by column chromatography (yield 50 mg). JG-2001-F1 was mixed with DCM and triflic acid (0.1 ml) and cooled on ice for 30 minutes. Product formation (JG-2031) was observed by TLC and LCMS (yield 20 mg).

**2-(7-chloro-8-ethyl-2,4-dioxo-3,4-dihydrobenzo[g]pteridin-10(2H)-yl)ethyl phenylcarbamate**

**(JG-2031**)-

1H NMR (400MHz, DMSO, ppm)  δ 11.51 (s, 1H), 9.5 (s, 1H), 8.22 (s, 1H), 7.9 (s, 1H), 7.27 (d, 4H), 6.97 (m, 1H), 4.9 (s, 2H), 4.5 (s, 2H), 2.85 (d, J = 7.2 Hz, 2H), 1.21 (t, J = 6.8 Hz, 3H)

LCMS, calc’d 439.86 m/z; found 440.74 m/z

#### Synthesis of JG-2029

**2-(7-chloro-8-ethyl-2,4-dioxo-3,4-dihydrobenzo[g]pteridin-10(2H)yl)ethylbenzoate (JG-2029)**

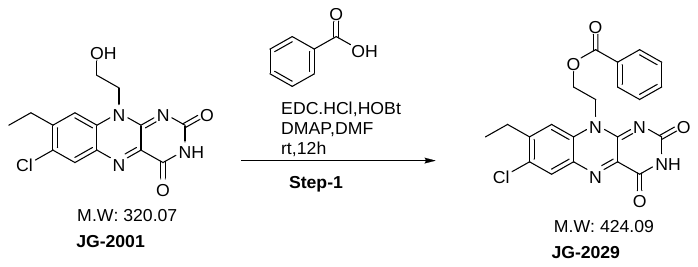

To a solution of benzoic acid (0.15 g, 0.12 mmol) in DMF (3 mL), EDC.HCl (0.35 g, 0.18 mmol), DMAP (0.45 g, 0.17 mmol) and HOBT (0.24 g, 0.17 mmol) were added. 7-chloro-8-ethyl-10-(2-hydroxyethyl)benzo[g]pteridine-2,4(3H,10H)-dione (**JG-2001**) (0.39 g, 0.12 mmol) in DMF (2 mL) was added to the reaction mixture and stirred at rt for 12 min. The reaction mixture was slowly poured into water (10 mL) and extracted with ethyl acetate (3 x 10 mL). The combined organic layer was dried over anhydrous sodium sulphate and concentrated under vacuum. The crude was purified with prep-HPLC to obtained title compound as pale yellow solid (0.036 g, 18.18%). LCMS: 425.19 (M+1); 1H NMR (400 MHz, DMSO-d6) δ 11.49(s, 1H), 8.25 (s, 1H), 8.22 (s, 1H), 7.73-7.59(m, 3H), 7.43-7.39(m, 2H), 5.06(t, 2H), 4.71 (bs, 2H), 2.87(q, J=7.2 Hz, 2H), 1.21(t, J=7.2 Hz, 3H).

#### Synthesis of JG-2030

**2-(7-chloro-8-ethyl-2,4-dioxo-3,4-dihydrobenzo[g]pteridin-10(2H)-yl)ethylisobutyrate (JG-2030)**

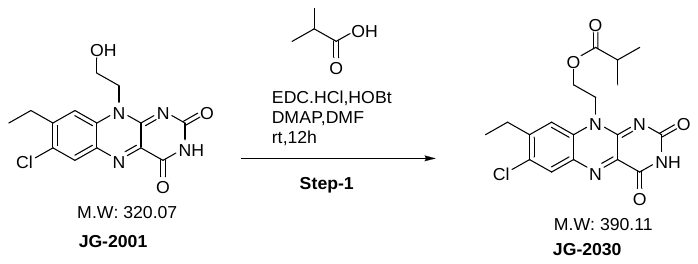

To a solution of Isobutyric acid **(** (0.15 g, 0.17 mmol) in DMF (3 mL), EDC.HCl (0.48 g, 0.25 mmol), DMAP (0.34 g, 0.25 mmol) and HOBT (0.34 g, 0.25 mmol) were added and stirred at room temperature. 7-chloro-8-ethyl-10-(2-hydroxyethyl)benzo[g]pteridine-2,4(3H,10H)-dione (**JG-2001)** (0.54 g, 0.16 mmol) in DMF (3 mL) was added and stirred for 10 min ar room temperature. The reaction mixture was slowly poured into water (10 mL) and extracted with ethyl acetate (3 x 10 mL). The combined organic layer was dried over anhydrous sodium sulphate and concentrated under vacuum. The crude was purified by prep-HPLC to obtained title compound as pale yellow solid (0.066g, 36.76 %). LCMS: 391.8, 393.8 (M& M+2); ^1^H NMR (400 MHz, DMSO-d_6_) δ 11.49 (s,1H), 8.23 (s, 1H), 8.08 (s, 1H), 4.93(t, 2H), 4.44(s, 2H), 2.95(q, J=7.6Hz, 2H), 2.36(hept, 1H), 1.31(t,J=7.6Hz, 3H), 0.90(d,J=6.8Hz, 6H).

#### Synthesis of JG-2066

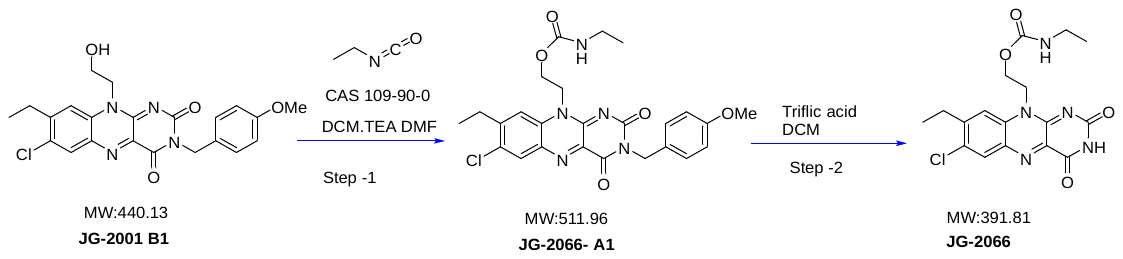

JG-2001-B1 (30 mg, 1 eq) was mixed with CAS 109-90-4 (4 eq), DCM, TEA and DMR (0.2 ml) for 2 hours at room temperature. Product (JG-2066-A1) was observed by TLC (yield 20 mg). The product was mixed with ice cold DCM and triflic acid for 1 hour at 0^0^C. Product (JG-2066) was observed by TLC and purified by column chromatography (yield 13 mg).

**2-(7-chloro-8-ethyl-2,4-dioxo-3,4-dihydrobenzo[*g*]pteridin-10(2*H*)-yl)ethylethylcarbamate**

**(JG-2066)-**

1H NMR (400 MHz, DMSO, ppm) δ 11.49 (s, 1H), 8.33 (s, 1H), 7.95 (s, 1H), 7.11 (t, J = 5.6 Hz, 1H), 4.85 (t, 2H), 4.36 (t, J = 5.2 Hz, 3H), 2.90 (m, 4H), 1.31 (t, J = 7.6 Hz, 3H), 0.85 (t, J = 7.2 Hz, 3H)

LCMS, calc’d 391.81 m/z; found 392.07 m/z

#### Synthesis of JG-2016 (scale-up)

**7-chloro-8-ethyl-10-(2-isobutoxyethyl)benzo[g]pteridine-2,4(3H,10H)-dione**

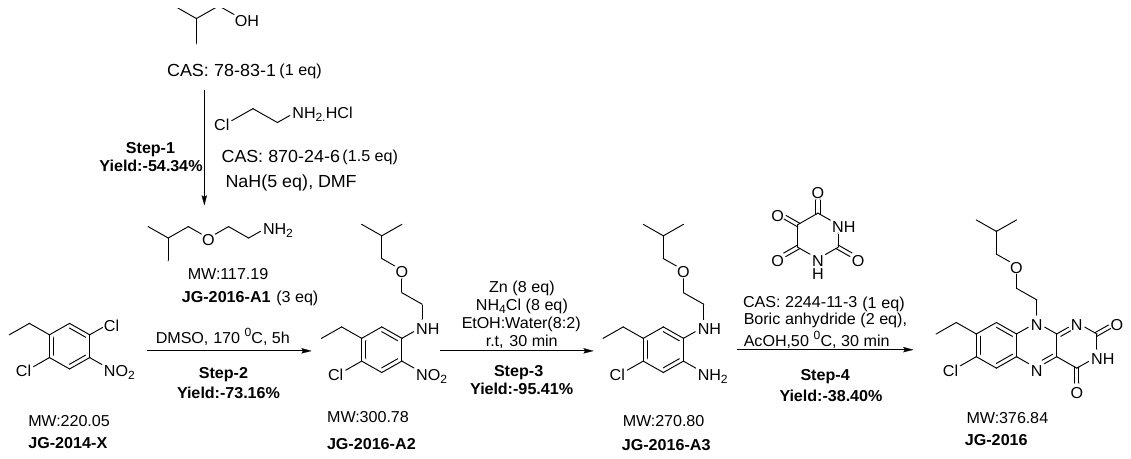

**Step-1 (JG-2016-A1)**

To a solution of 2-methylpropan-1-ol (**CAS: 78-83-1)** (25g, 337.2 mmol, 1 eq) in DMF (250 mL/ 10V), 60 % NaH (67.45g, 1686.4 mmol, 5.0 eq) was added portion wise and stirred at 0°C temperature for 1h under nitrogen atmosphere. 2-chloroethan-1-amine hydrochloride (**CAS: 870-24-6)** (58.68 g, 505.9 mmol, 1.5 eq) was added portion wise to the reaction mixture and stirred at room temperature for 4h. After completion of reaction as indicated by TLC, the reaction mixture was poured into aqueous sodium chloride solution and extracted with ethyl acetate (3 x 2L). The combined organic layer was dried over anhydrous sodium sulphate and concentrated under vacuum up to 300 ml then washed with water (2 x 500 mL). The organic layer was dried over anhydrous sodium sulphate and concentrated under vacuum to afford yellow biphasic liquid (21.48 g, 54.34 %) crude material which was directly used in next step without further purification. LCMS: 118.2(M+1)

**Step-2 (JG-2016-A2)**

To a solution of 1,4-dichloro-2-ethyl-5-nitrobenzene (**JG-2014-X)** (5.0 g, 22.72 mmol, 1.0 eq) in DMSO (5 mL), **JG-2016-A1** (13.31 g, 113.6 mmol, 5.0 eq) was added and stirred at 170 °C temperature for 5h. After completion of reaction as indicated by TLC, the reaction mixture was poured into ice cooled water (300 mL) and extracted with ethyl acetate (3 x 100 mL). The combined organic layer was dried over anhydrous sodium sulphate and concentrated under vacuum. The crude was further purified by column chromatography (4 % ethyl acetate/hexane) to afford yellow liquid (5.0 g, 73.16%) LCMS: 301.3, 303.3(M & M+2).

**Step-3 (JG-2016-A3)**

To a solution of 4-chloro-5-ethyl-N-(2-isobutoxyethyl)-2-nitroaniline (**JG-2016-A2)** (6.1 g, 20.28 mmol, 1.0 eq) in ethanol: water (8:2V) Zn Dust (10.60 g, 162.2 mmol, 8.0 eq) and NH_4_Cl (8.67g, 162.2 mmol, 8.0 eq) were added and stirred the reaction mixture at room temperature for 30 min at room temperature. TLC indicated completion of reaction. The reaction mixture was frittered through celite and filtrate was extracted with ethyl acetate (2 x 50 mL). The organic layer was dried over anhydrous sodium sulphate and concentrated under reduce pressure to afford yellow liquid (5.24g, 95.41 %). The crude product was used in next step without further purification.

LCMS: 271.3, 273.2(M&M+2).

**Step-4 (JG-2016)**

To a solution of 4-chloro-5-ethyl-N1-(2-isobutoxyethyl)benzene-1,2-diamine (**JG-2016-A3)** (5.24 g, 19.35 mmol, 1.0 eq) in AcOH (25 mL) alloxan monohydrate (**CAS: 2244-11-3)** (3.09 g, 19.35 mmol, 1.0 eq) and boric anhydride (2.69 g, 38.70 mmol, 2.0 eq) were added and stirred at 50 °C temperature for 30 min. The reaction mixture was poured into ice cooled water (100 mL) and extracted with ethyl acetate (3 x 50 mL). The reaction mixture was dried over anhydrous sodium sulphate Filtrate was concentrated under reduce pressure. The obtained crude product was further purified by column chromatography (80 % ethyl acetate/hexane) and pure fraction was concentrated under vacuum to afford yellow solid (2.8g, 38.40 %). LCMS: 377.3, 379.3(M&M+2). ^1^H NMR (400 MHz, DMSO) δ 11.44 (s, 1H), 8.18 (s, 1H), 8.03 (s, 1H), 4.80 (t, 2H), 3.78 (t, *J* = 4.8 Hz, 2H), 3.12 (d, *J* = 6.4 Hz, 2H), 2.91 (q, *J* = 7.3 Hz, 2H), 1.62 (m, 1H), 1.27 (t, *J* = 7.4 Hz, 3H), 0.70 (d, *J* = 6.7 Hz, 6H).

### Proton NMR Spectra

JG-2001-A1

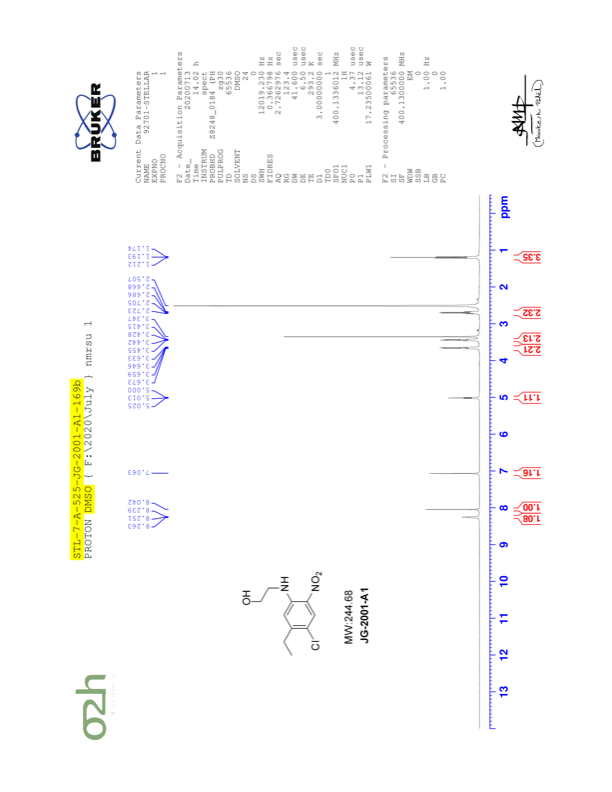

JG-2001-A2

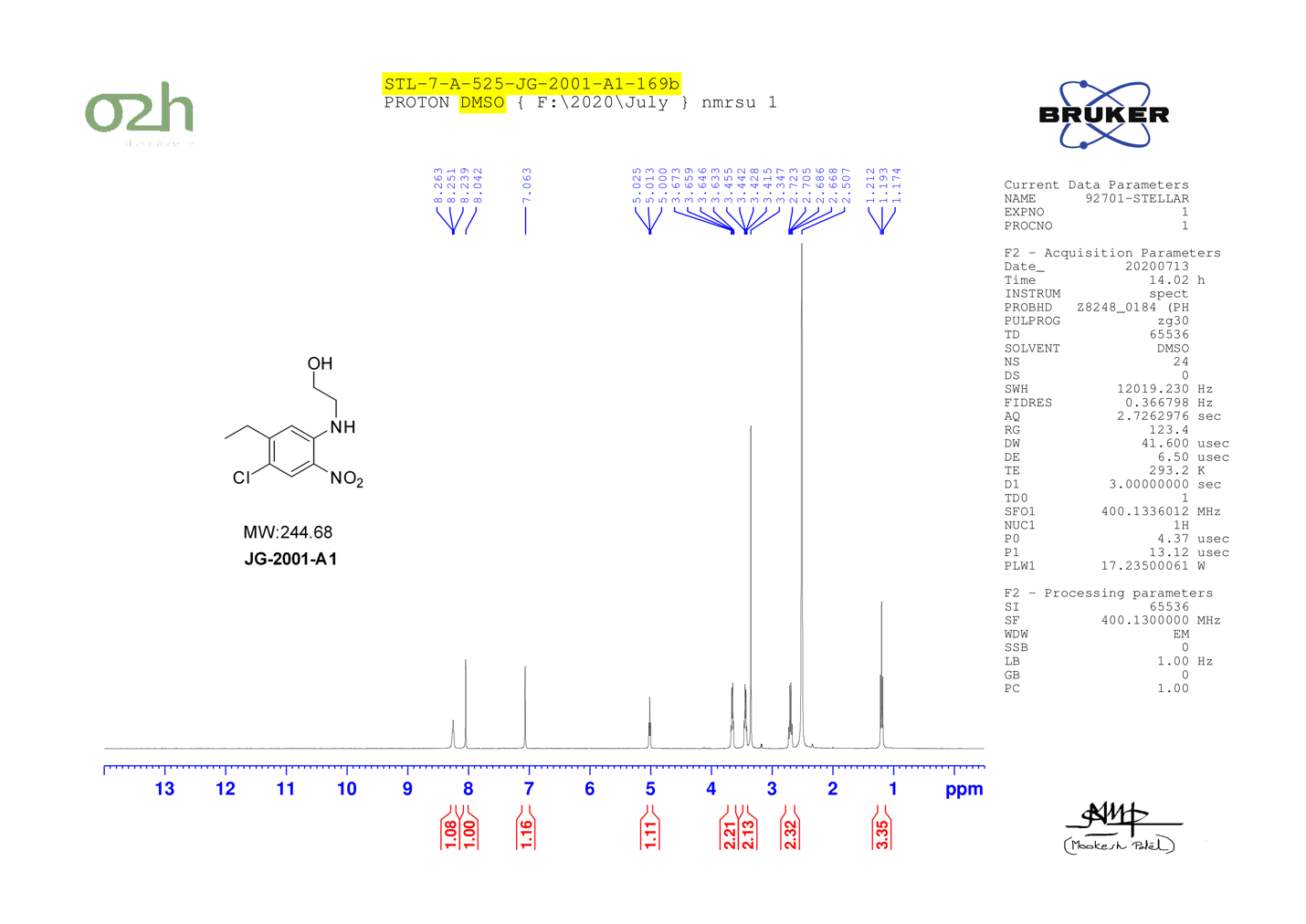

JG-2001-B1

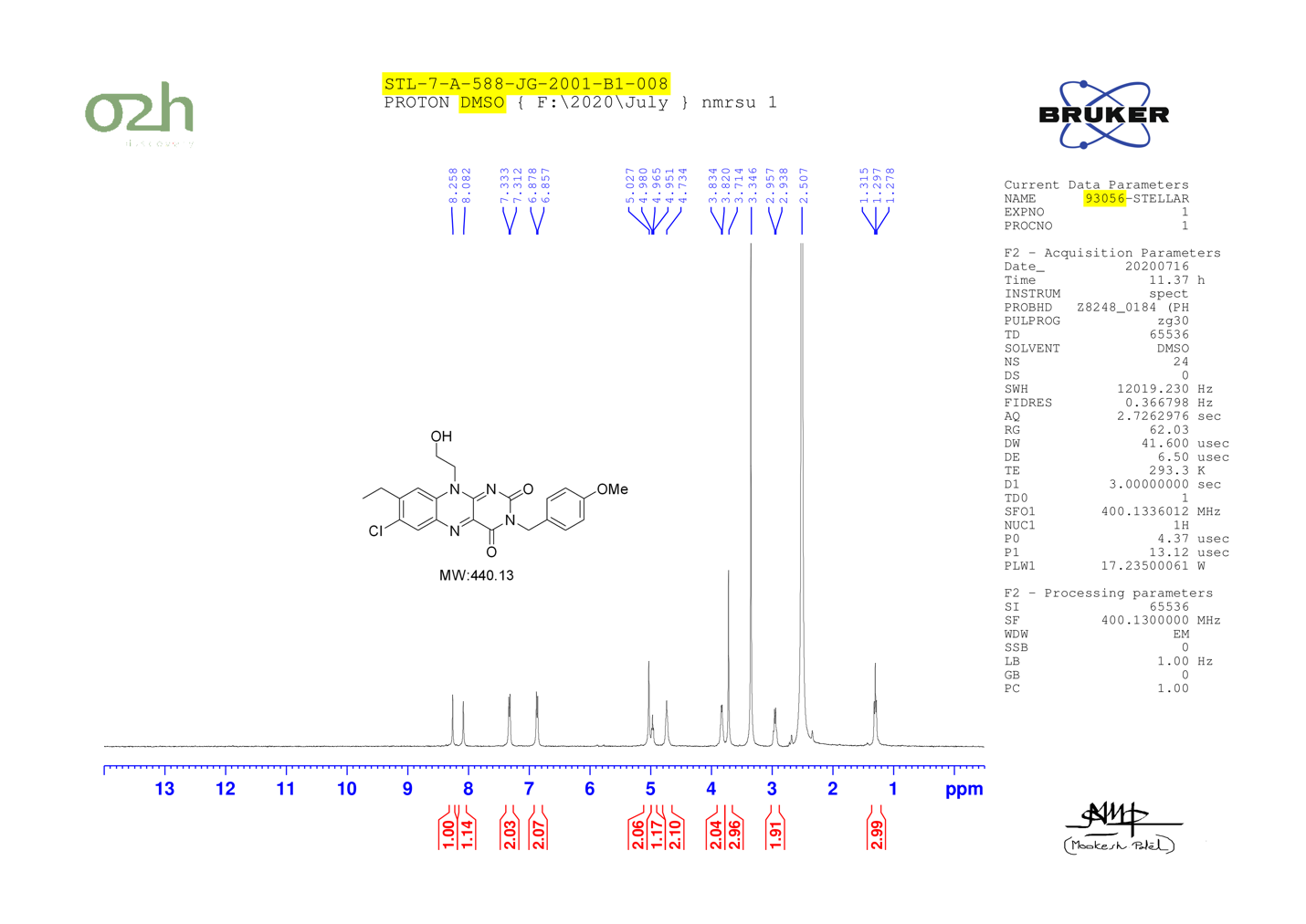

JG-2005

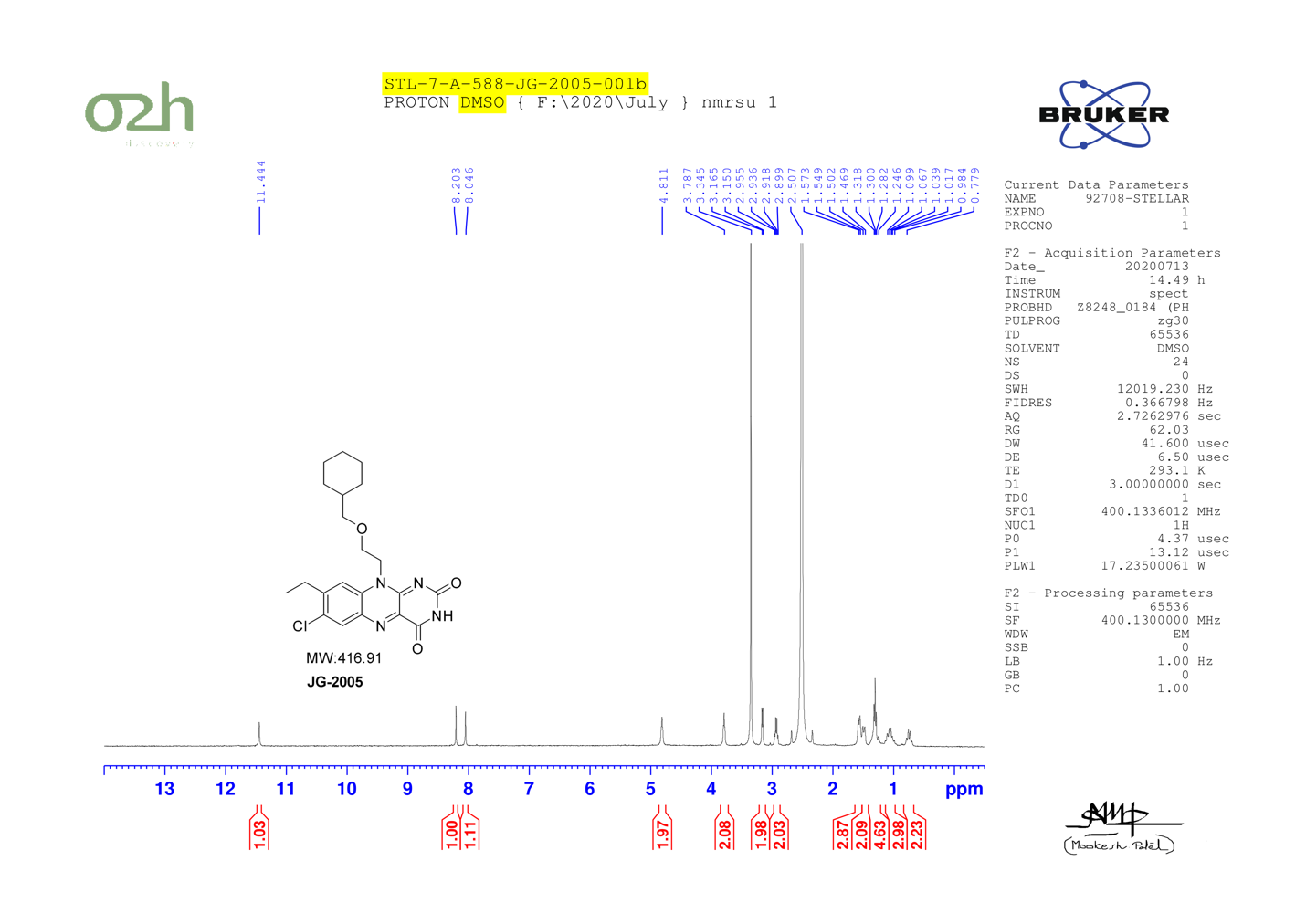

JG-2011

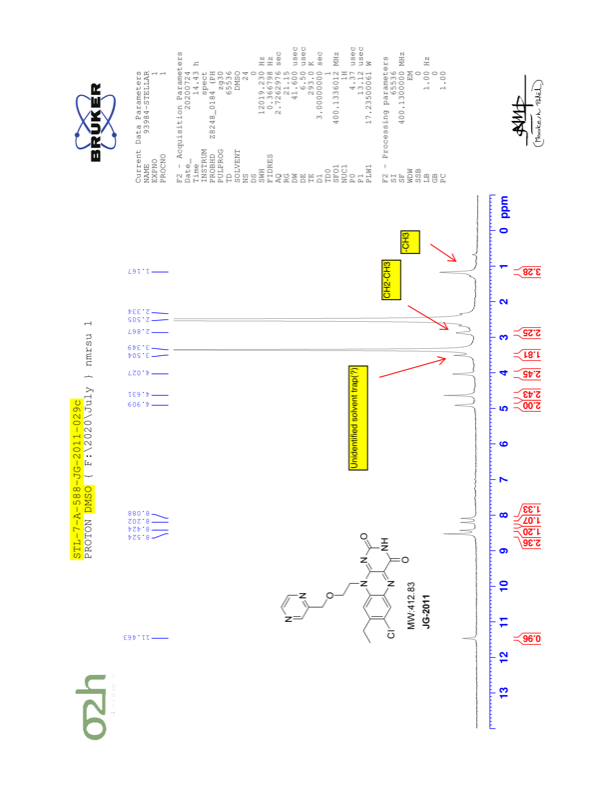

JG-2026

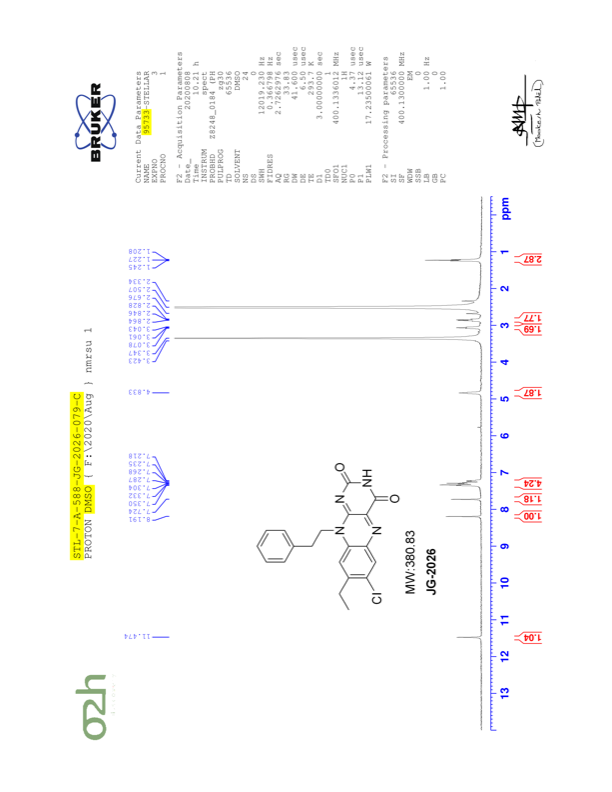

JG-2027

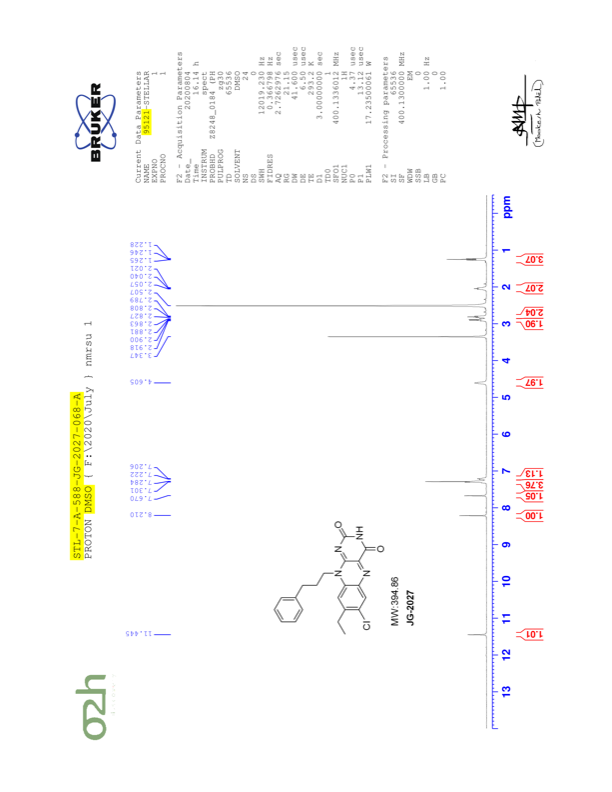

JG-2029

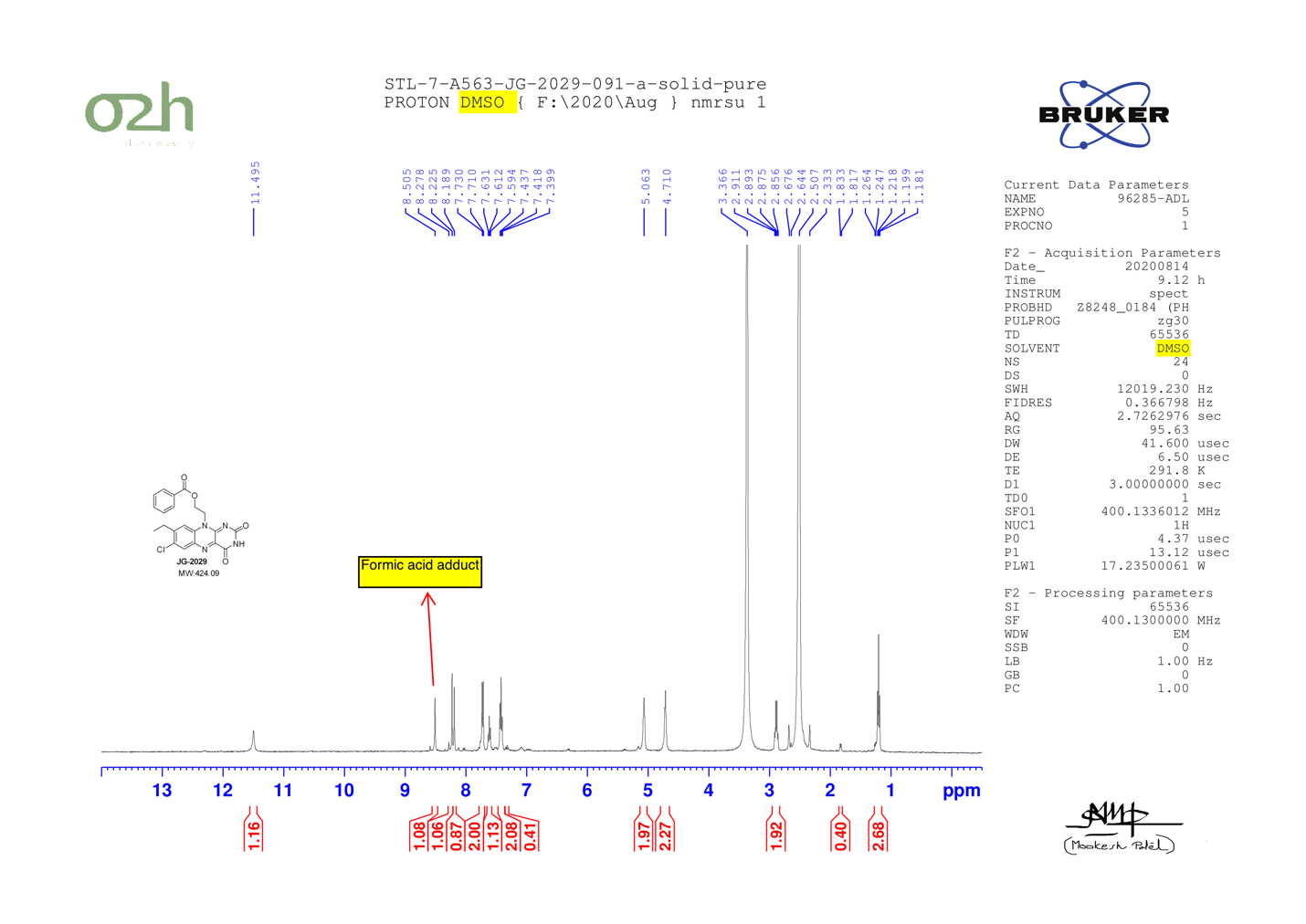

JG-2030

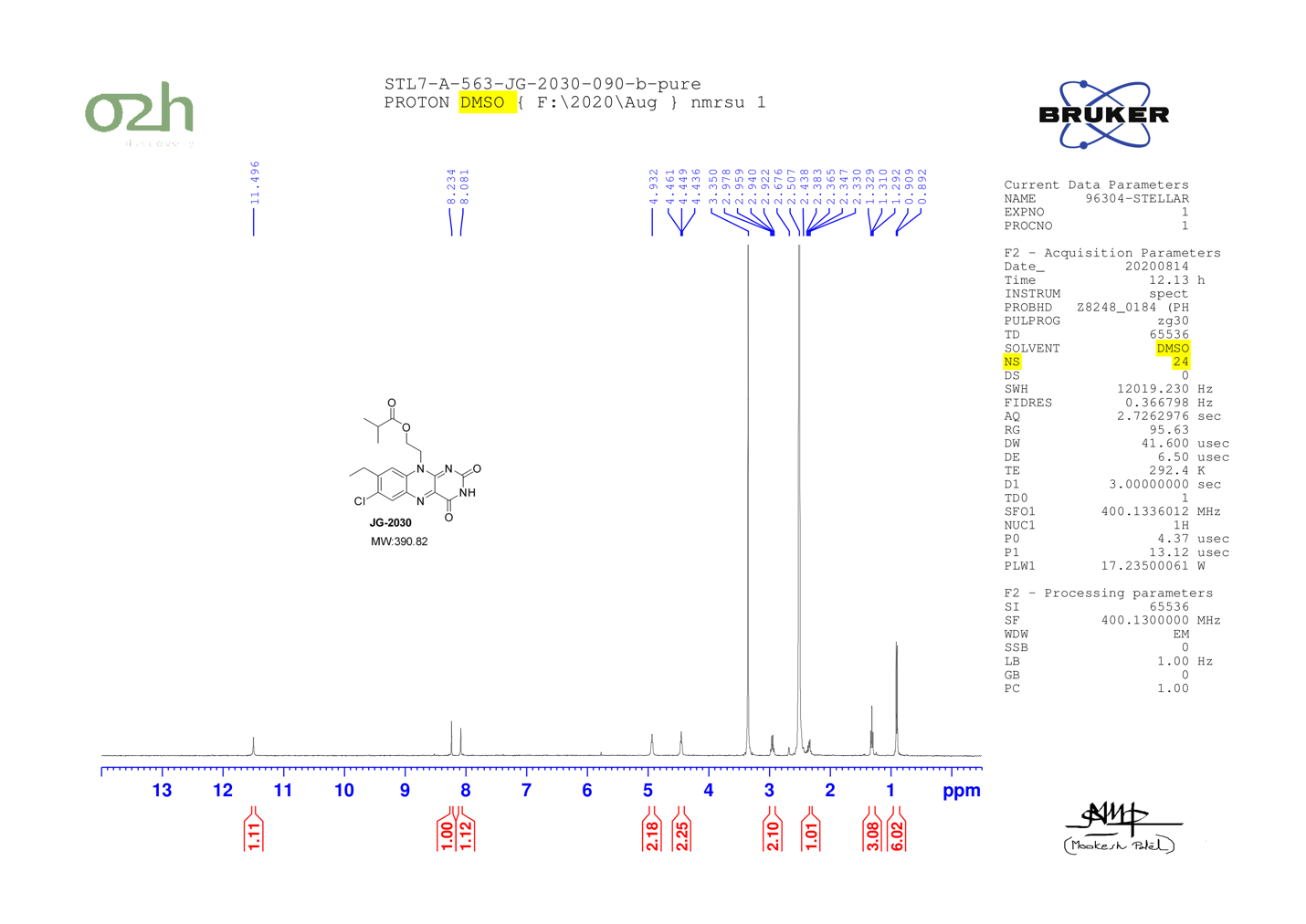

JG-2031

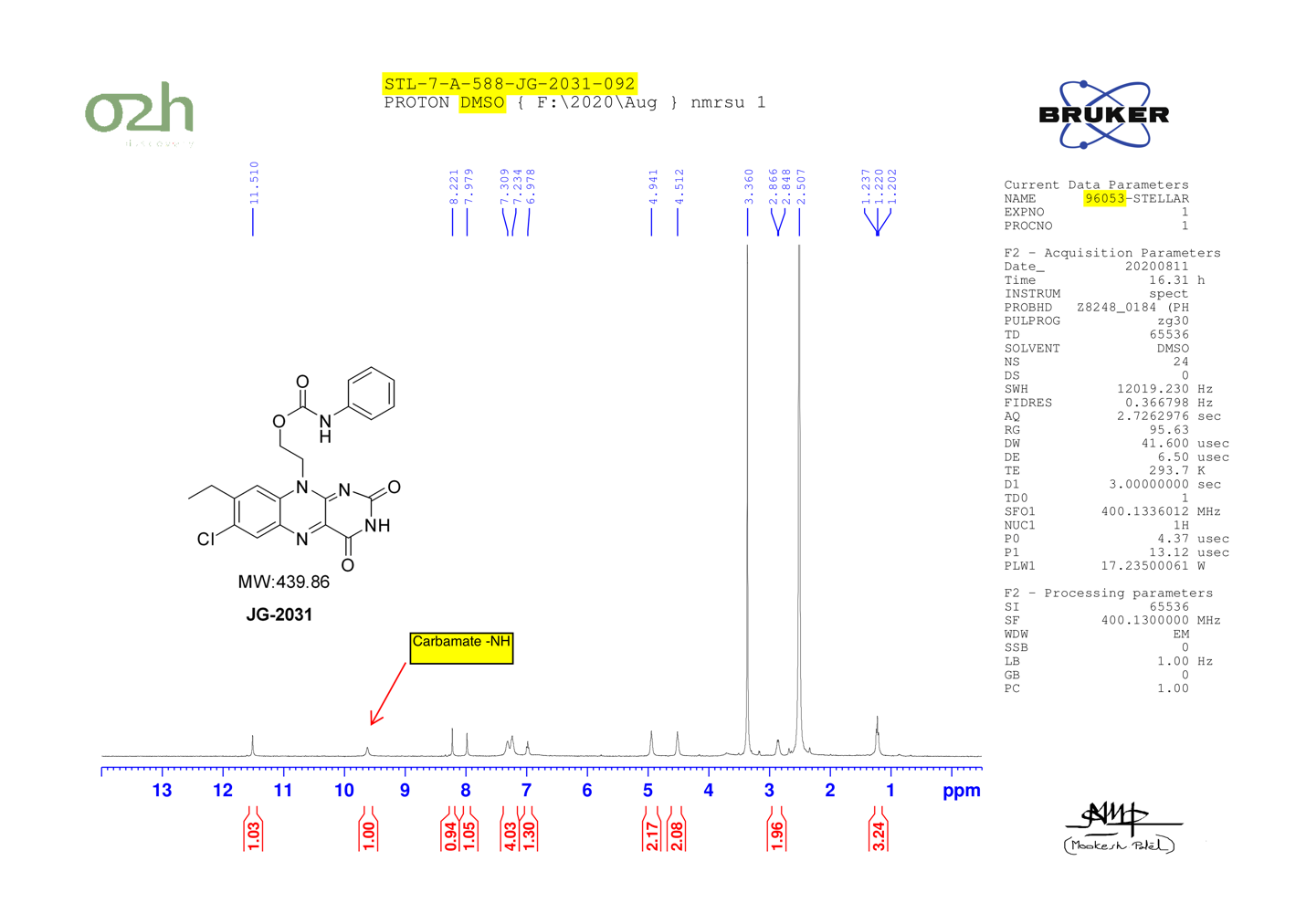

JG-2002-G4

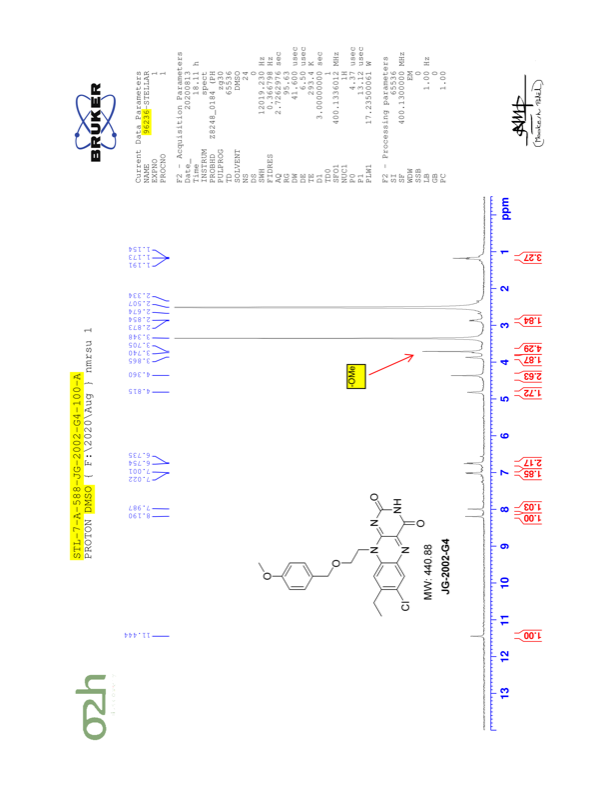

JG-2028

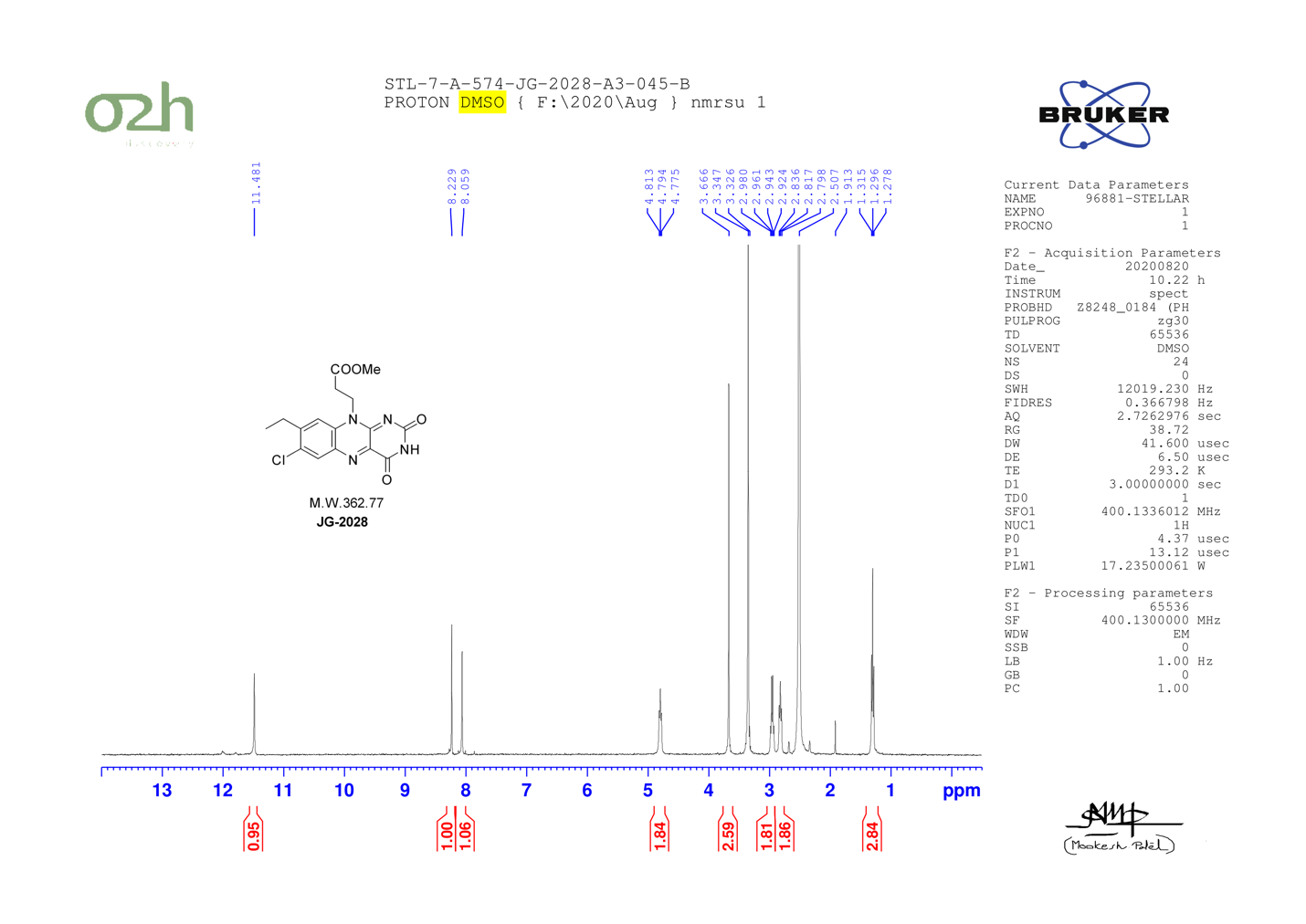

JG-2038

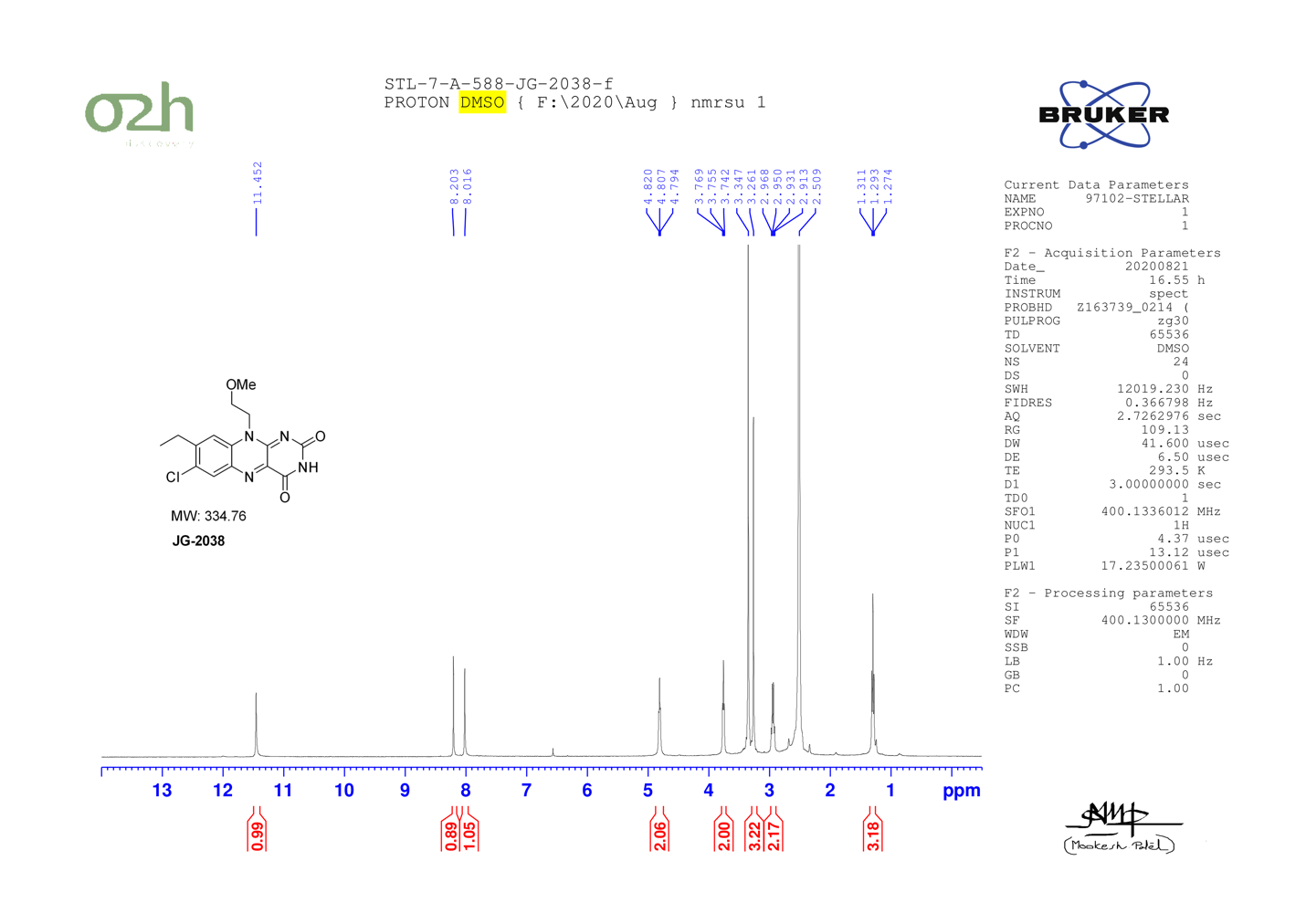

JG-2039
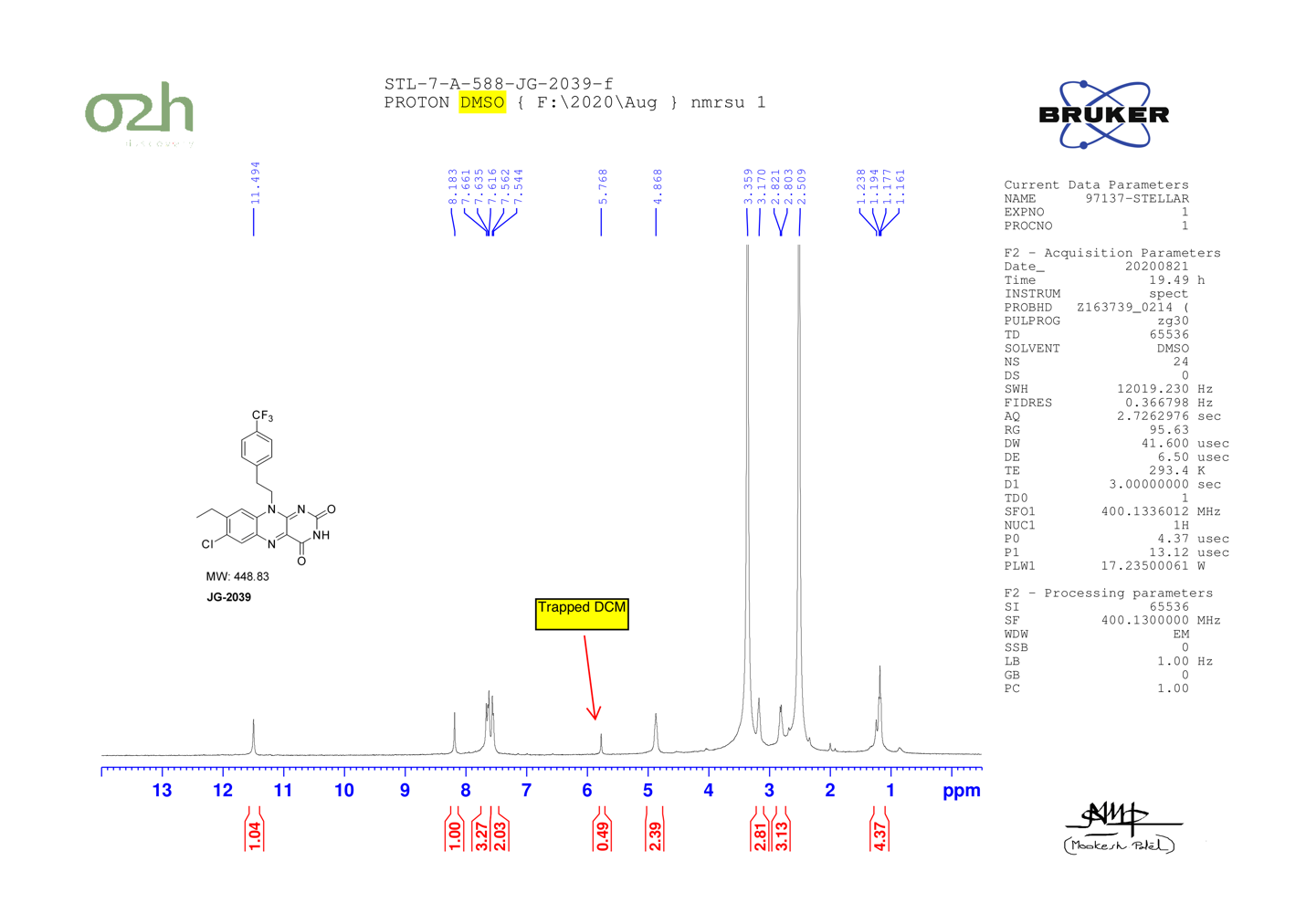

JG-2040
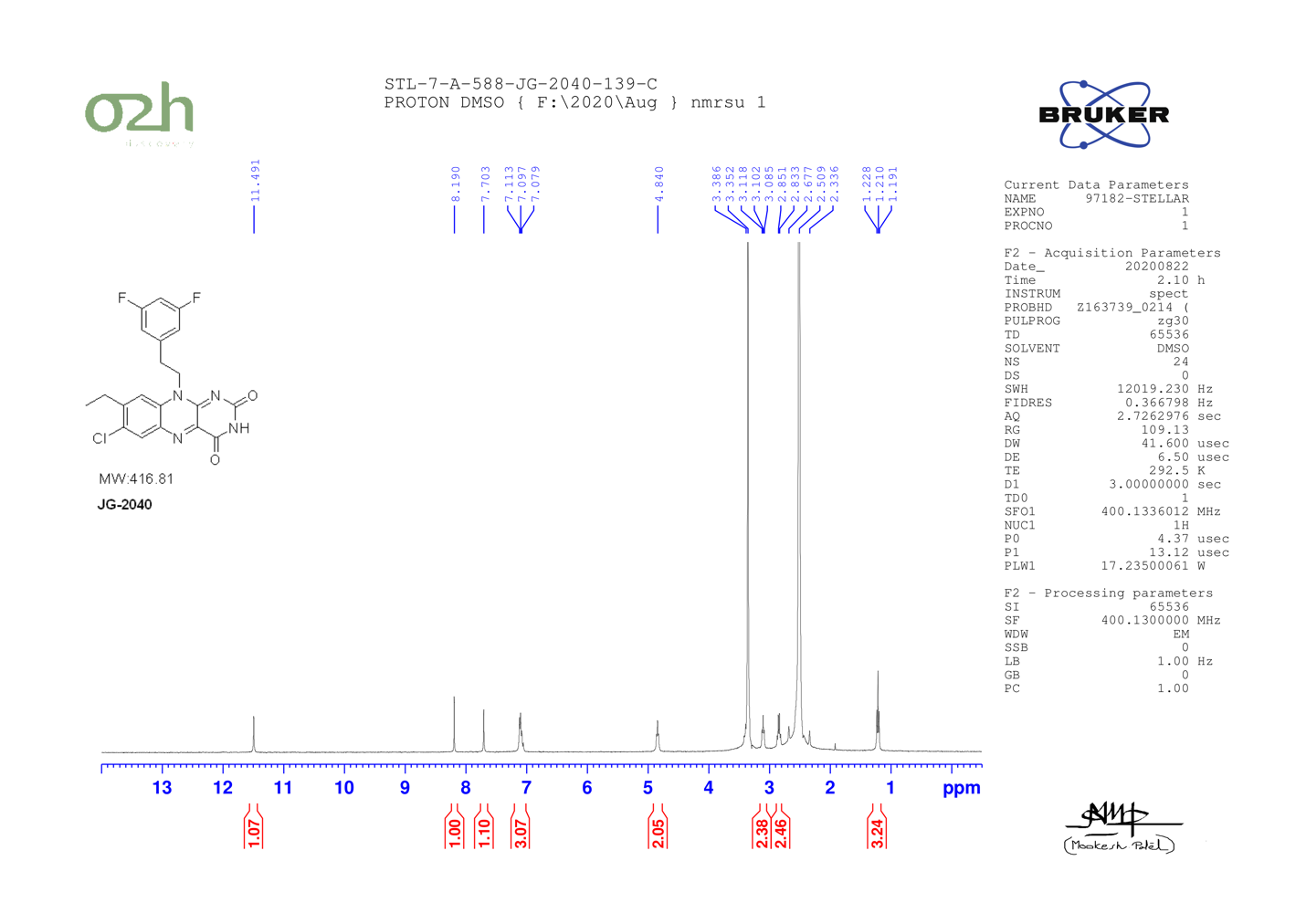

JG-2041
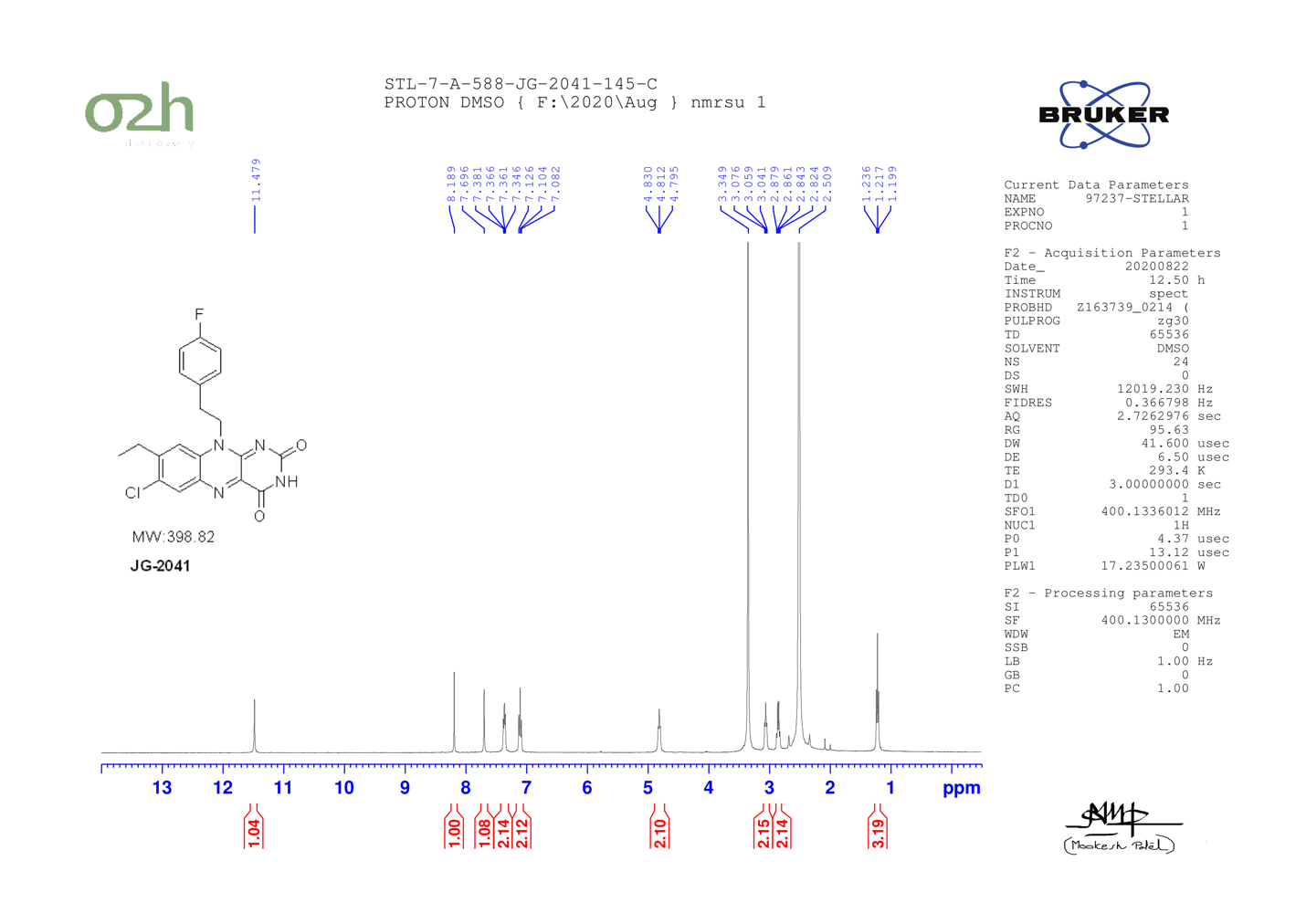

JG-2042
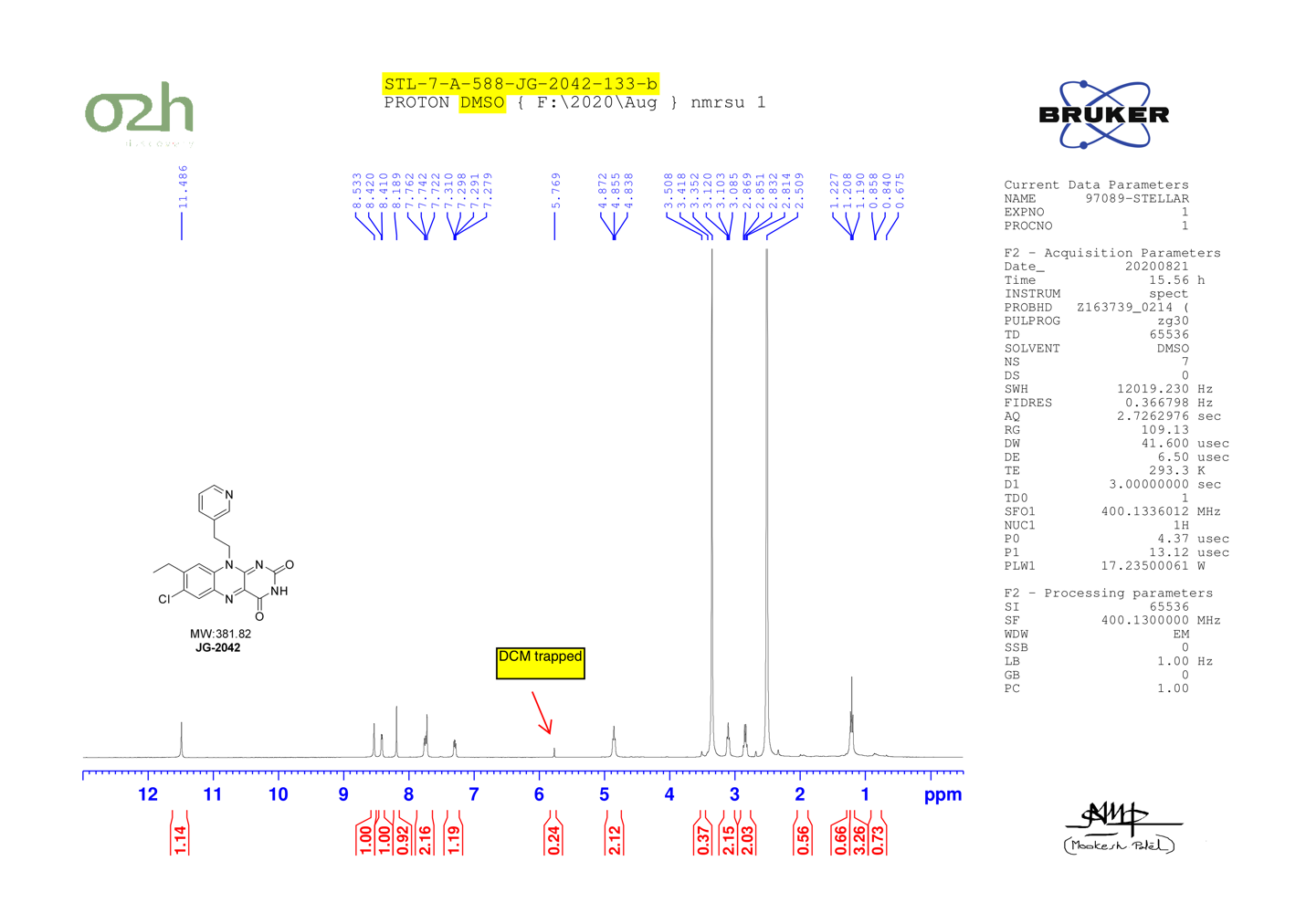

JG-2043

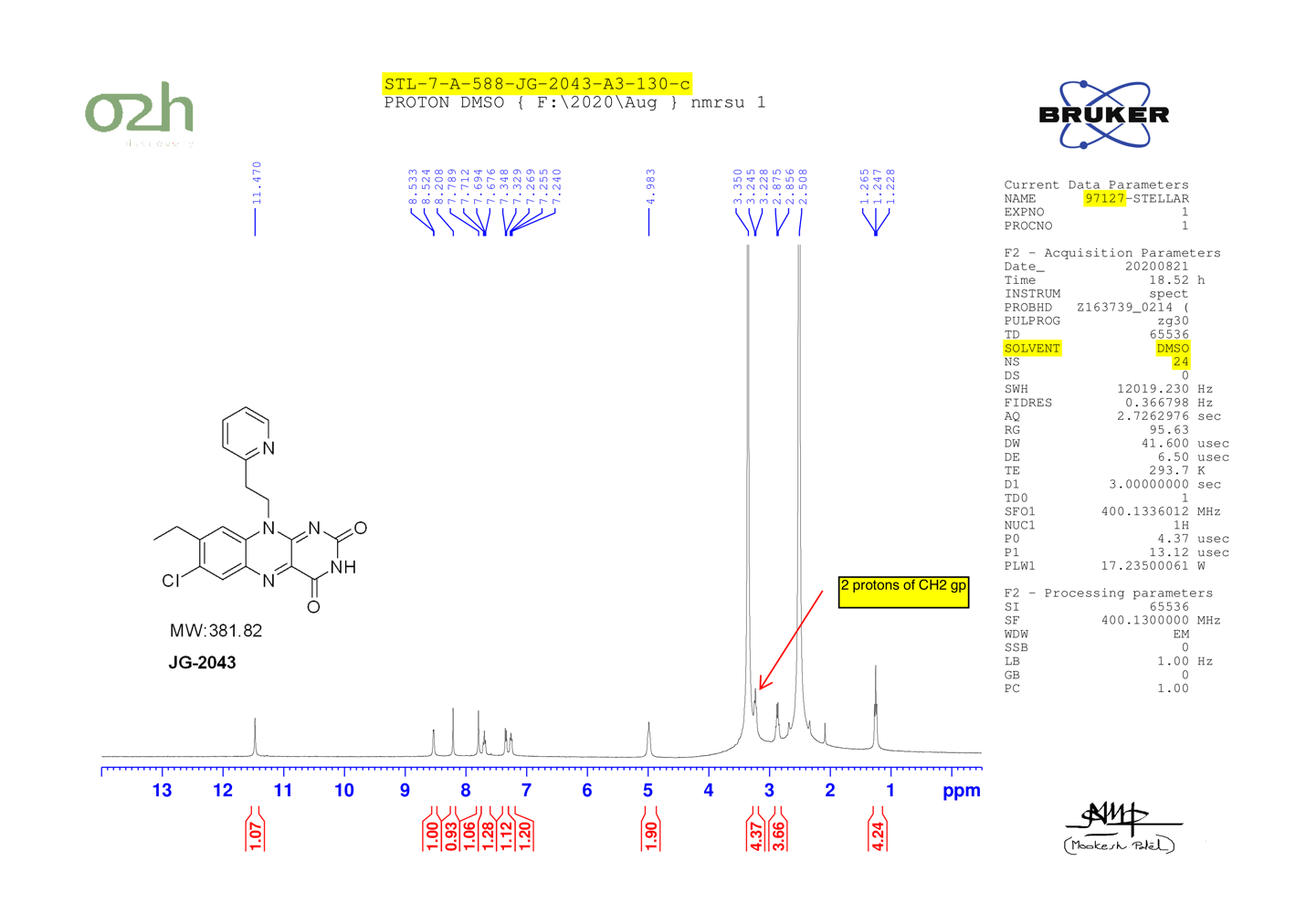

JG-2046
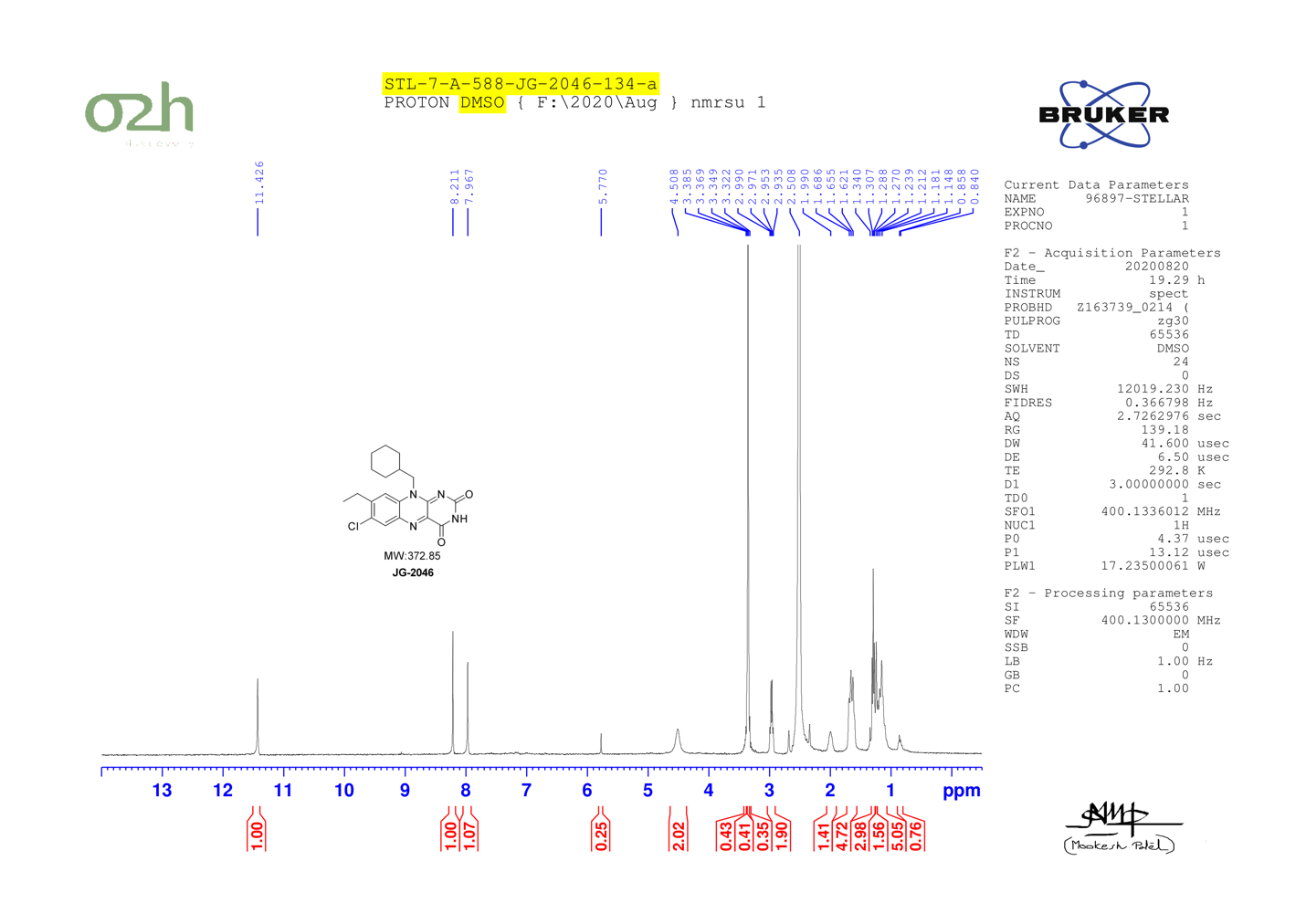

JG-2047

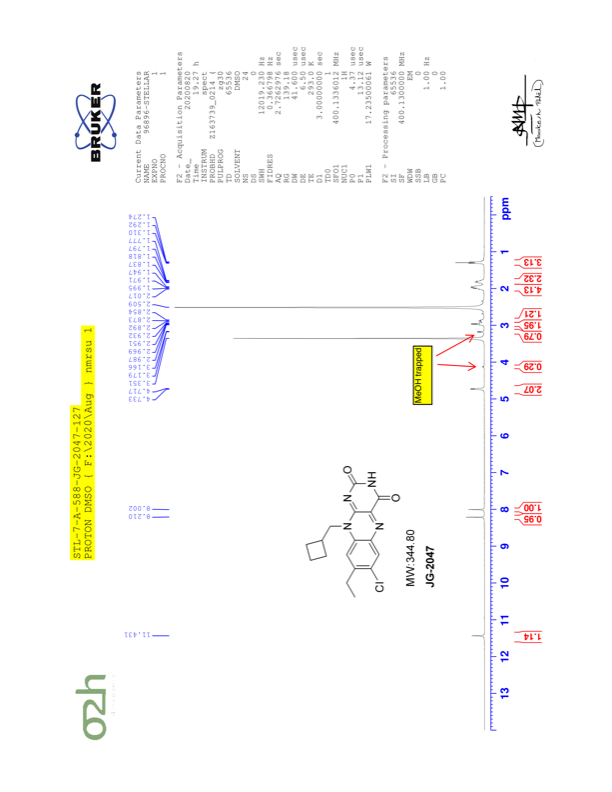

JG-2048

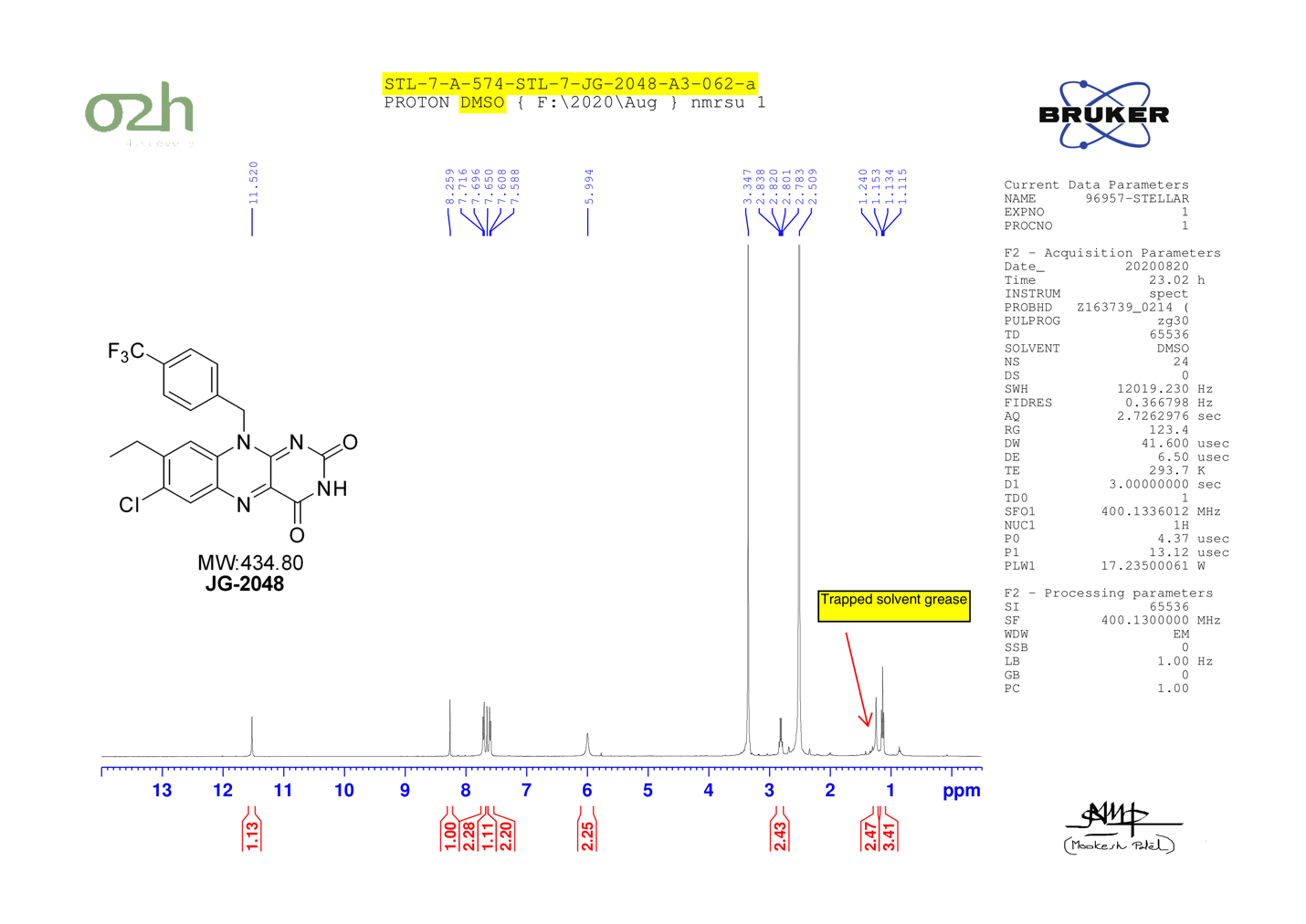

JG-2057
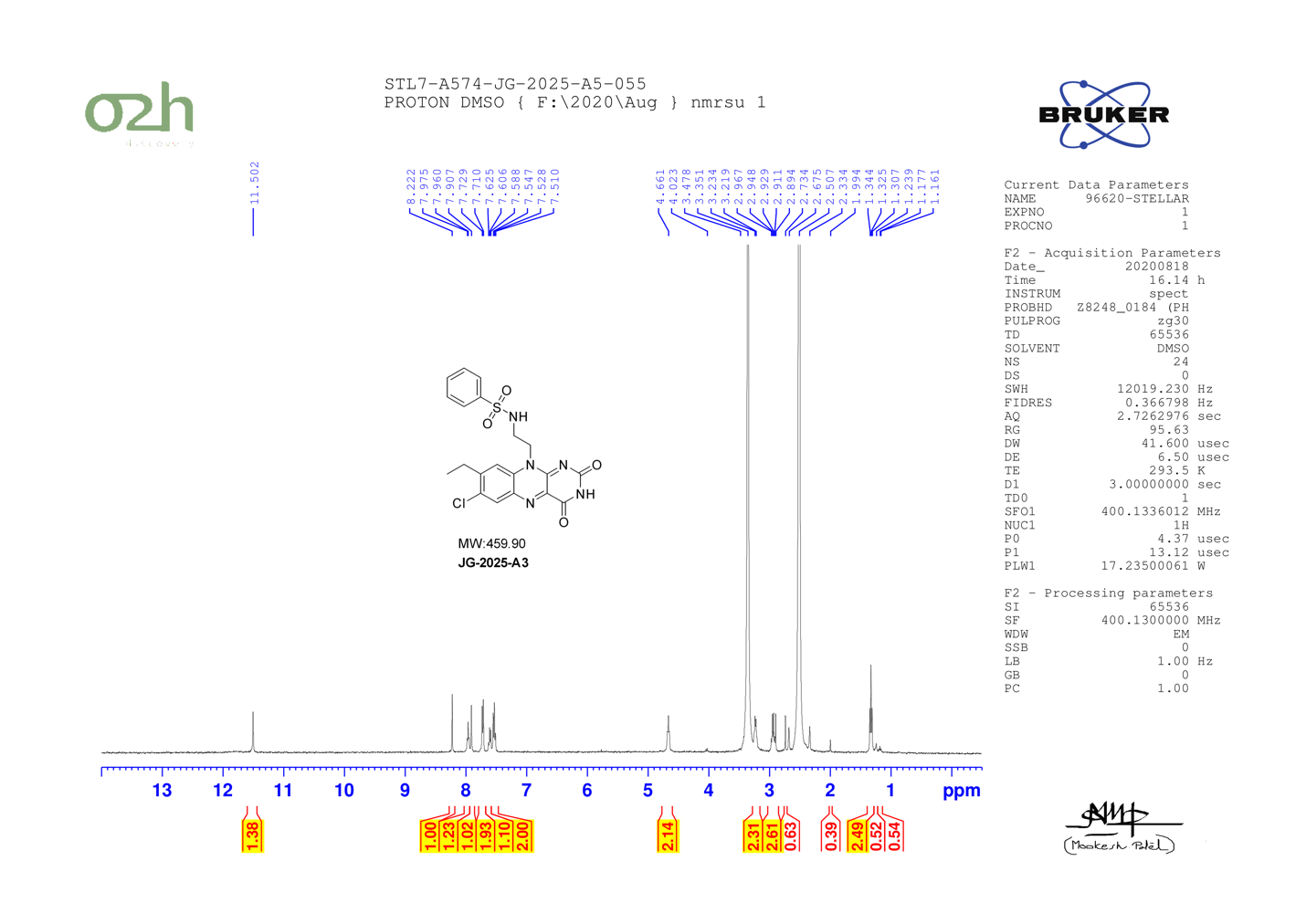

JG-2025-Boc

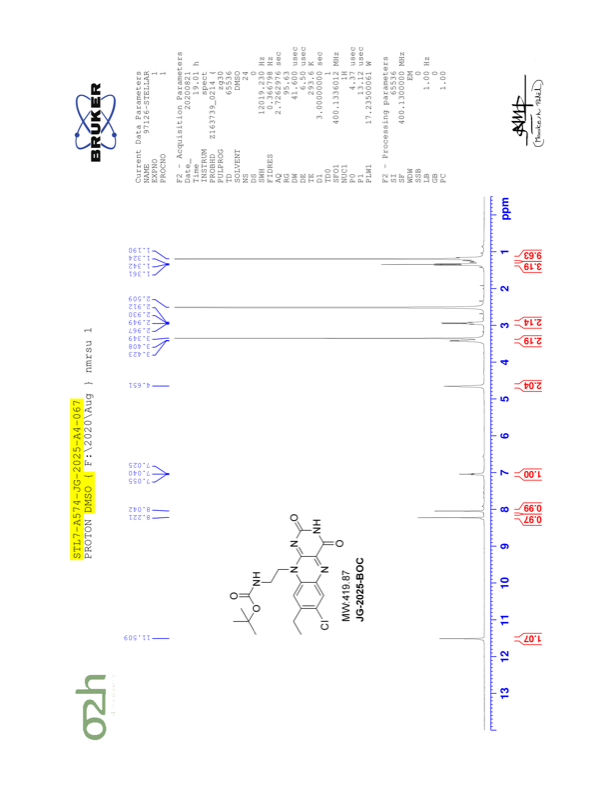

JG-2034

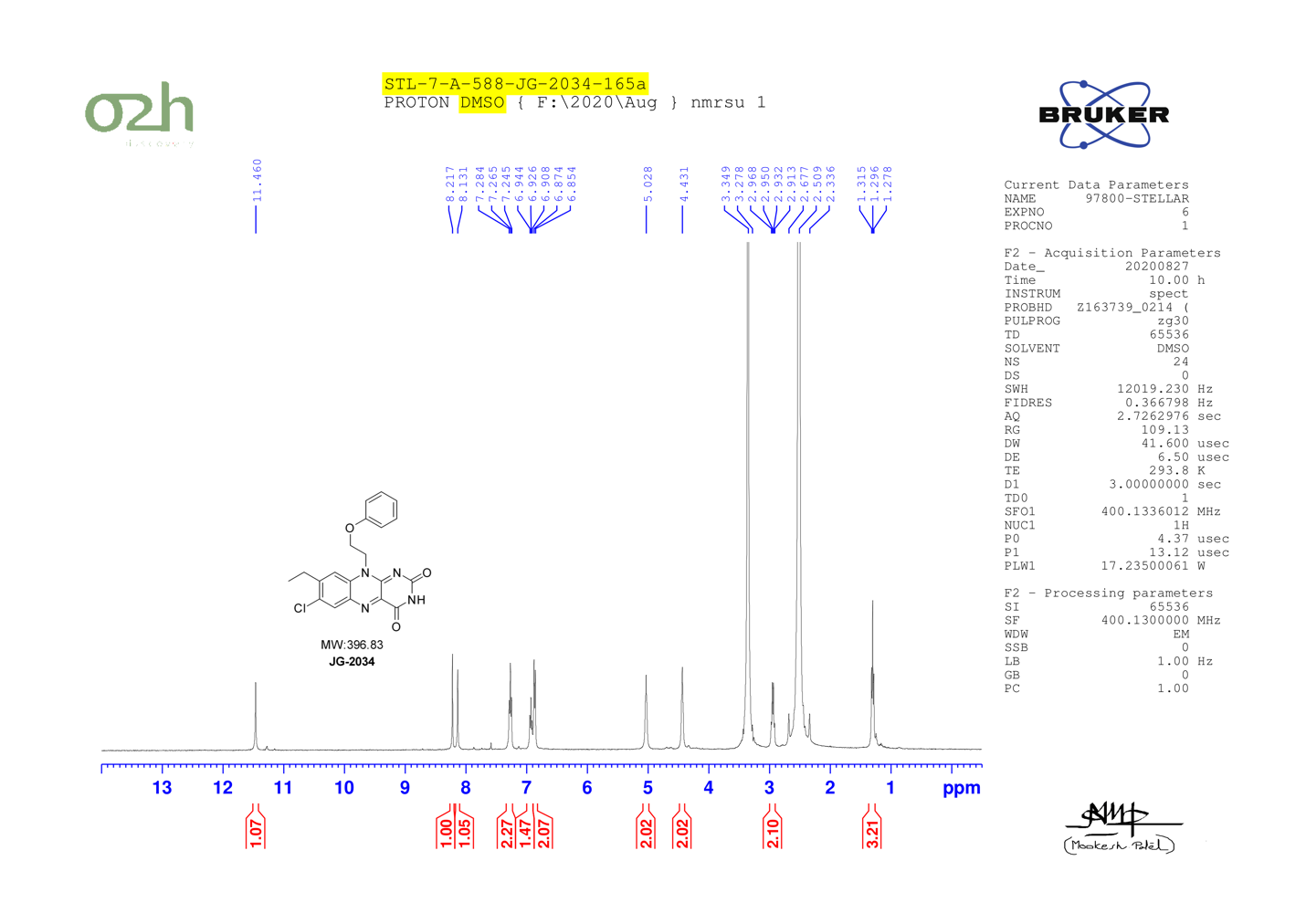

JG-2036

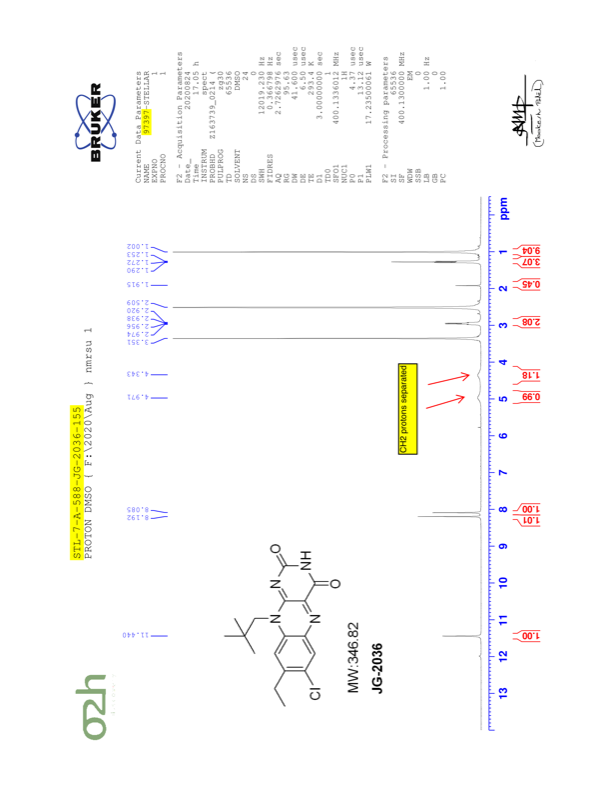

JG-2045

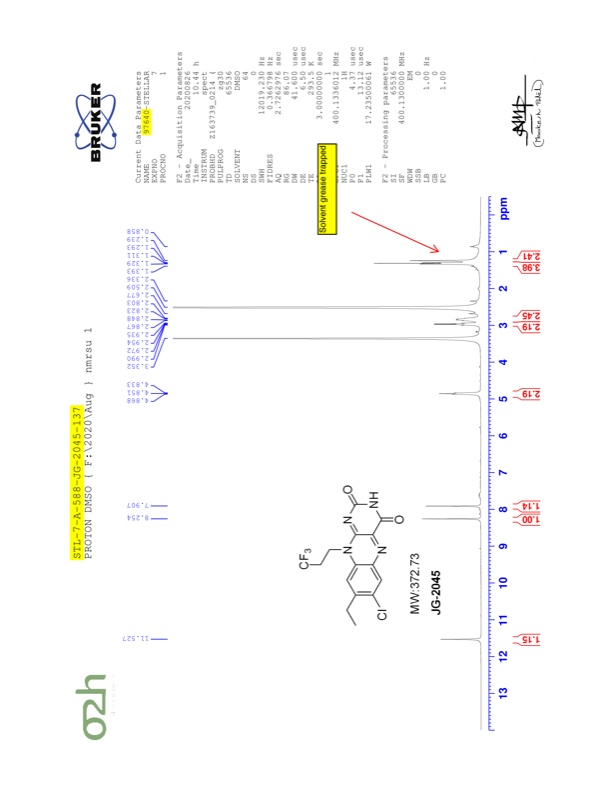

JG-2049

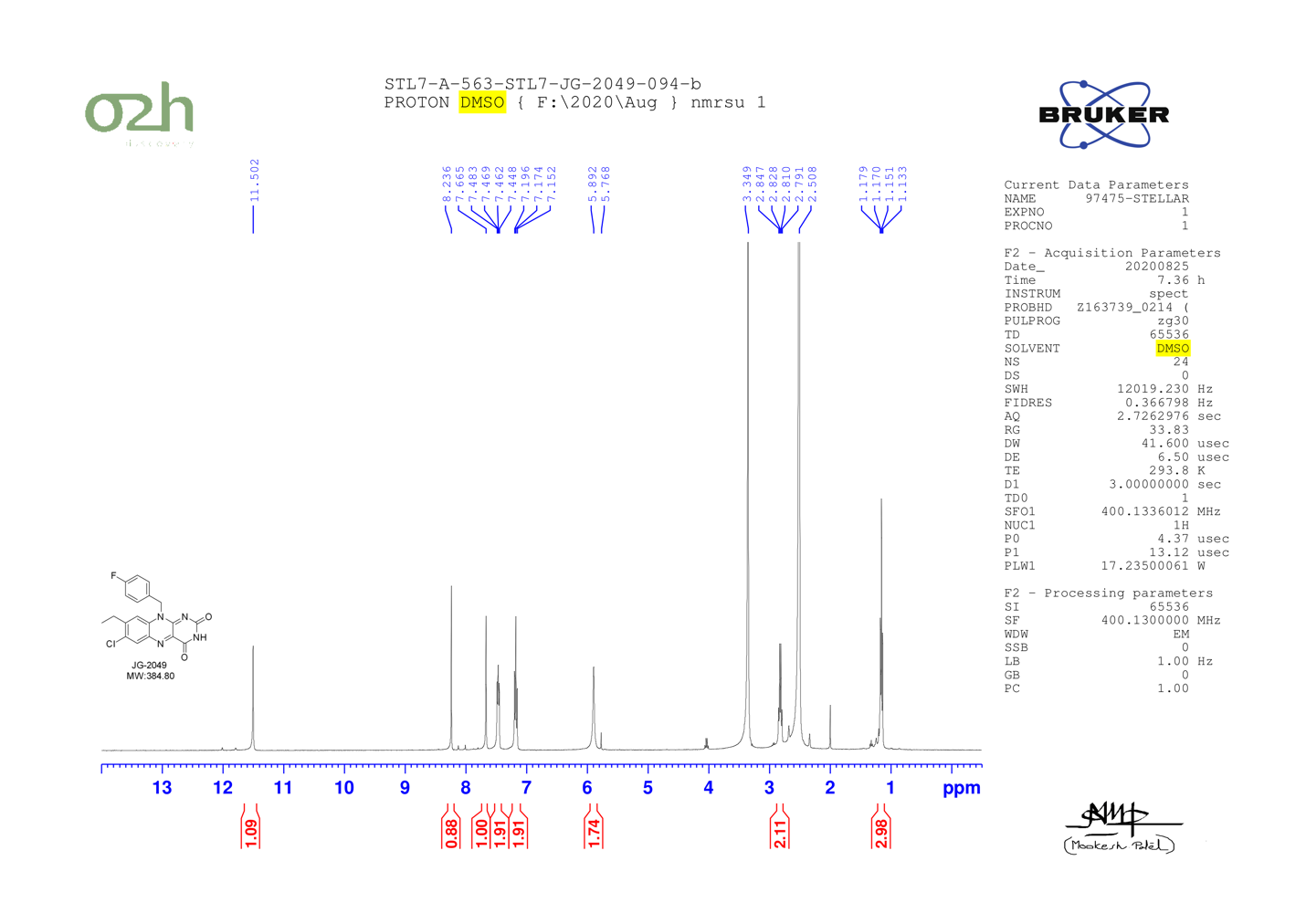

JG-2050

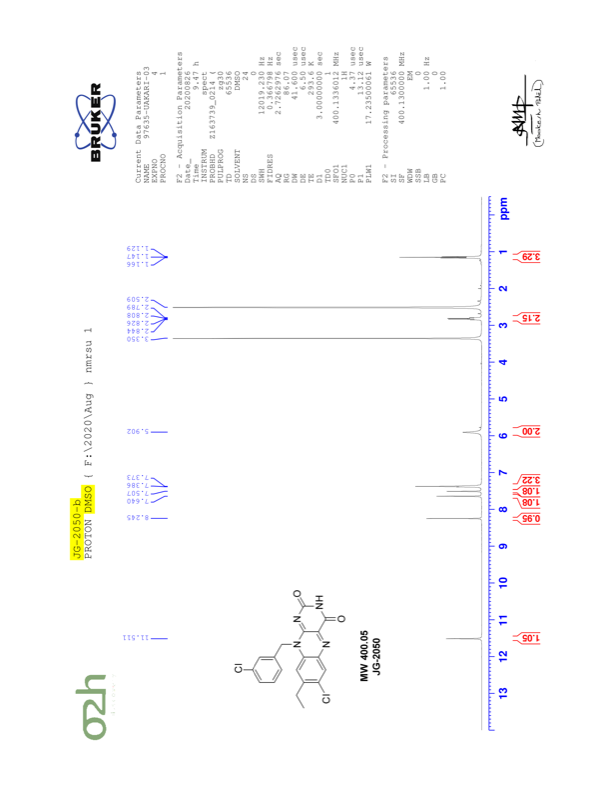

JG-2053

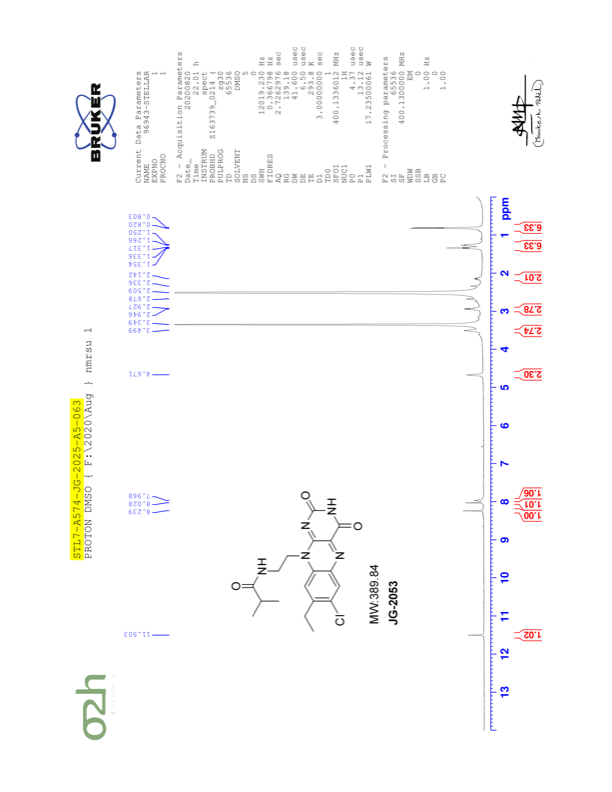

JG-2066

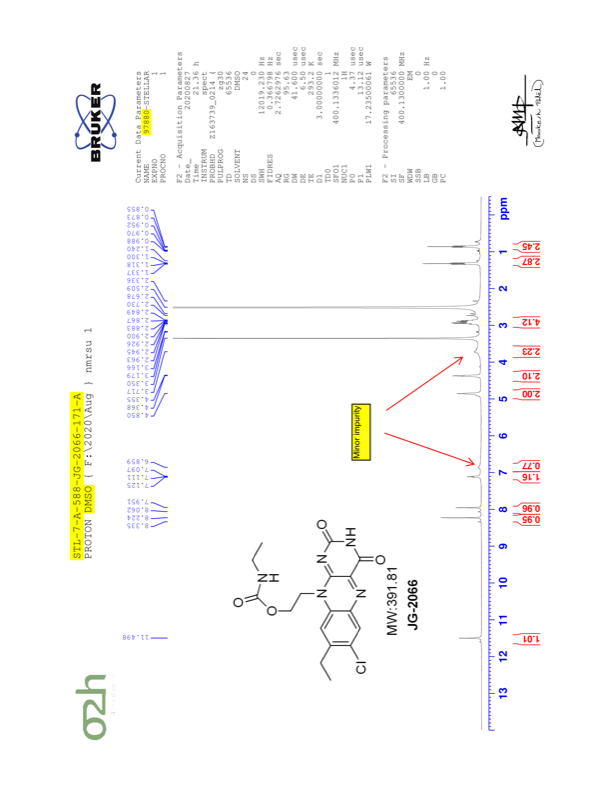

JG-2061

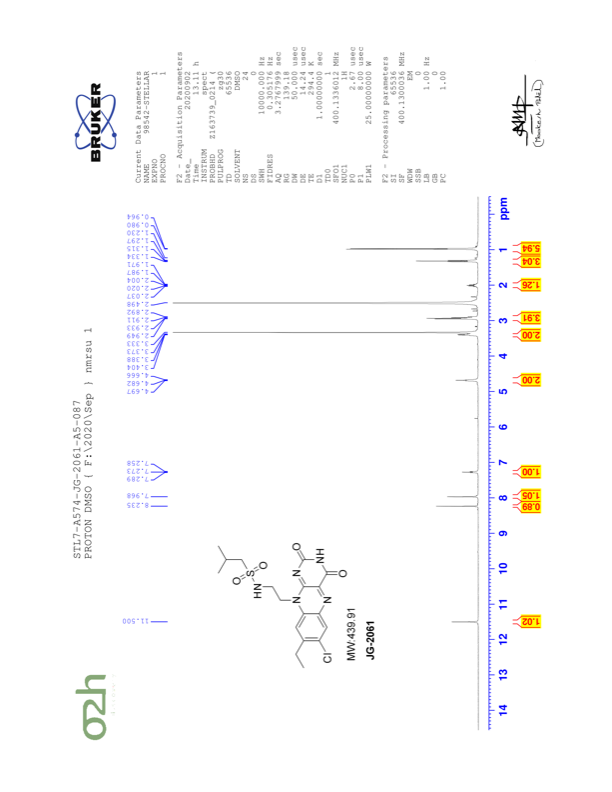

JG-2062

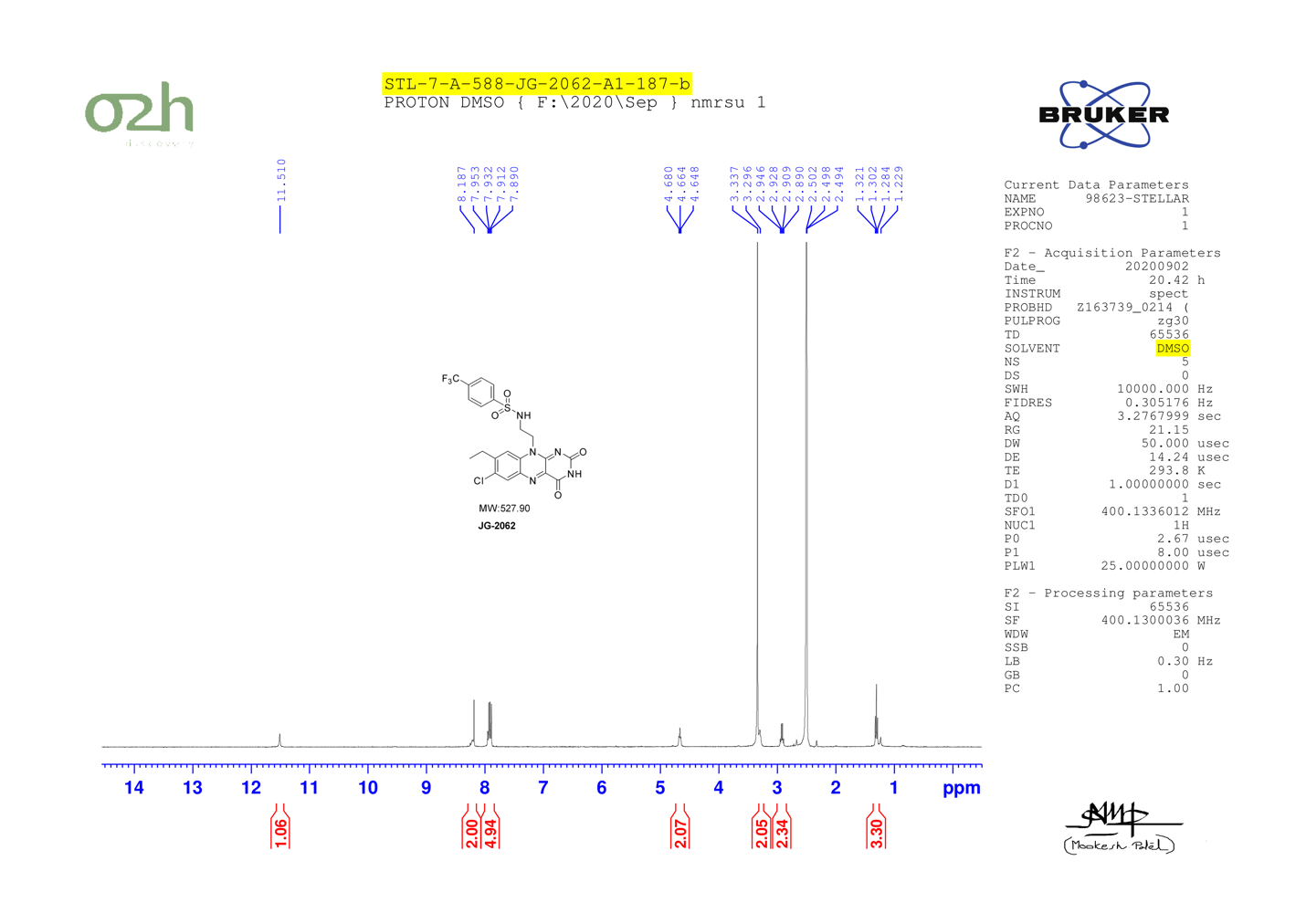

JG-2063

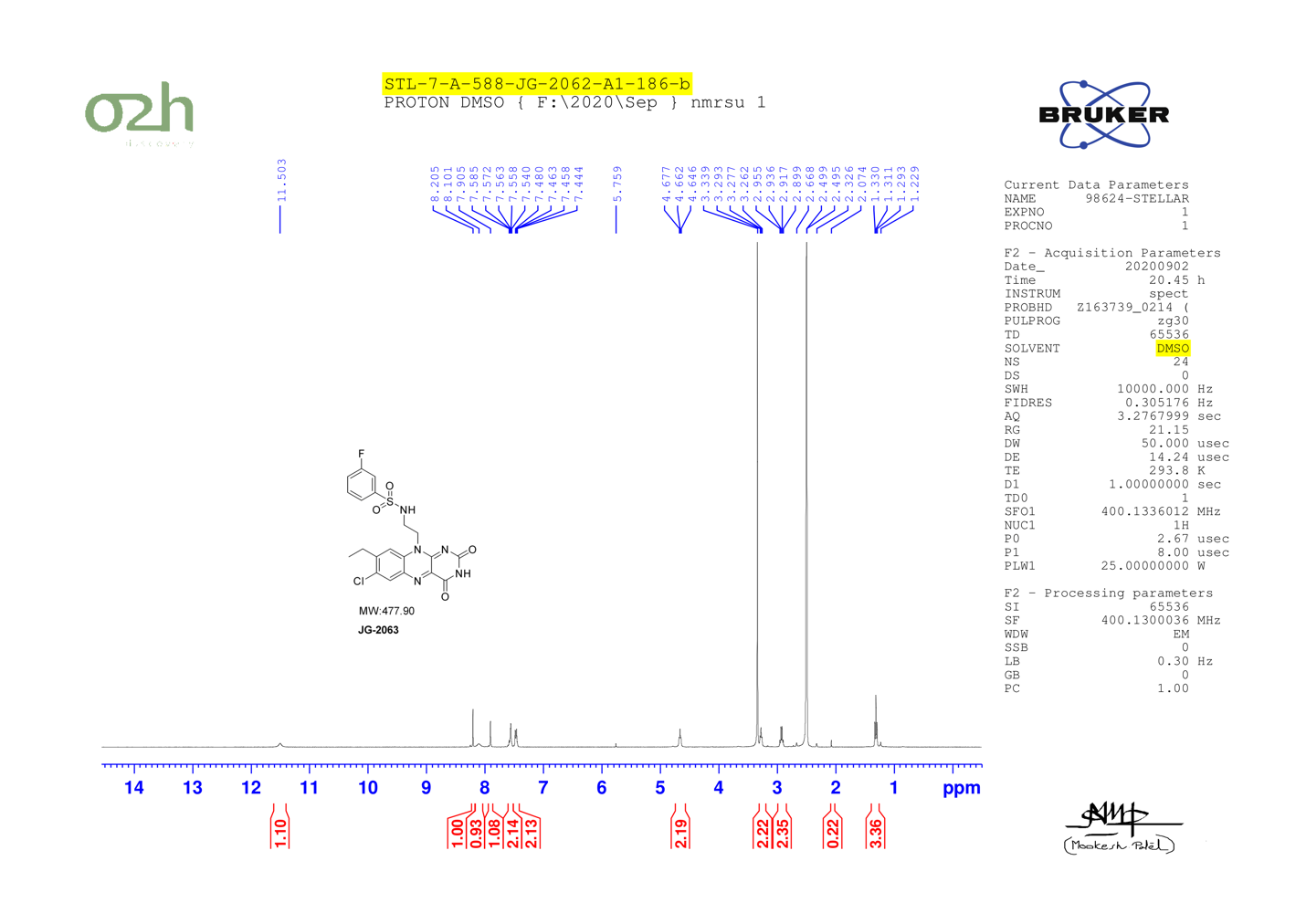

JG-2064

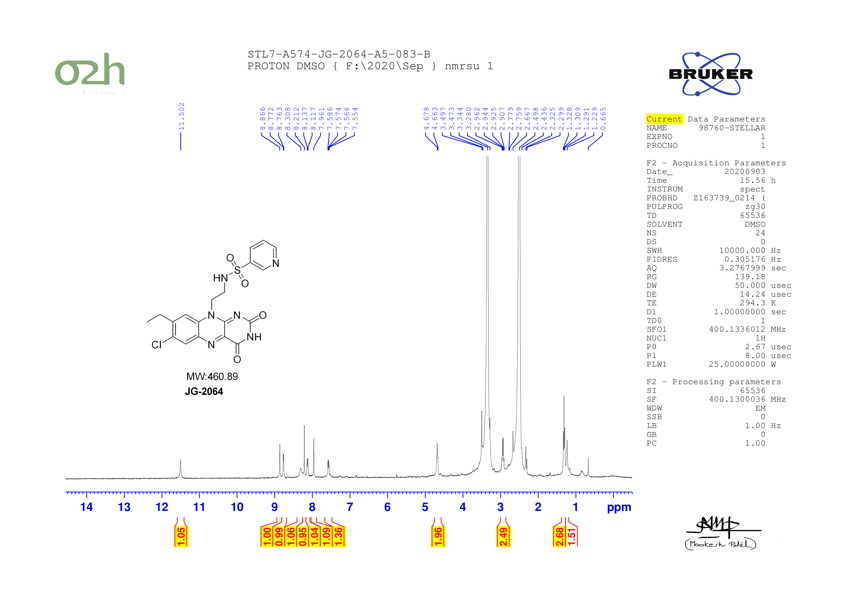

JG-2065

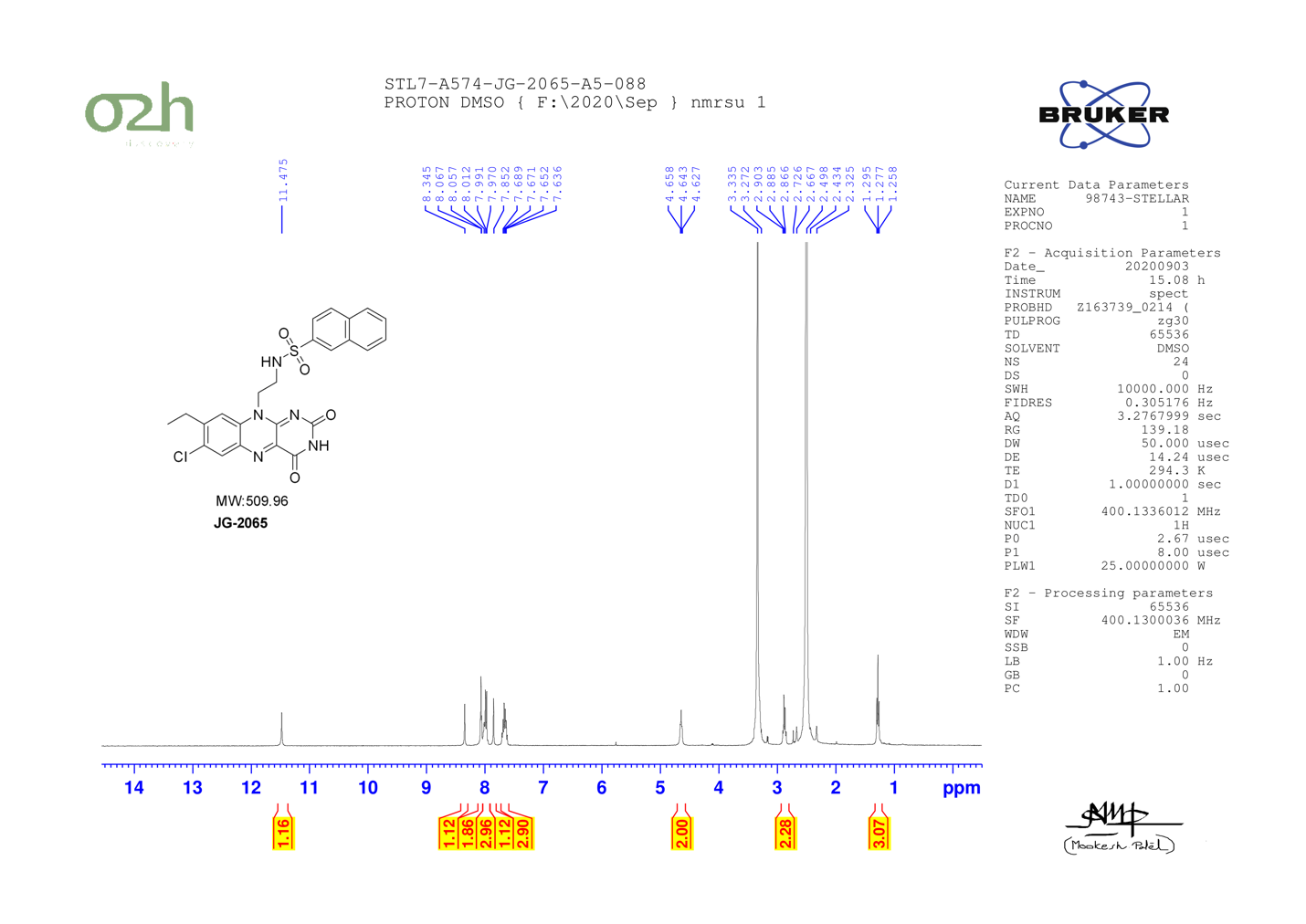

JG-2060

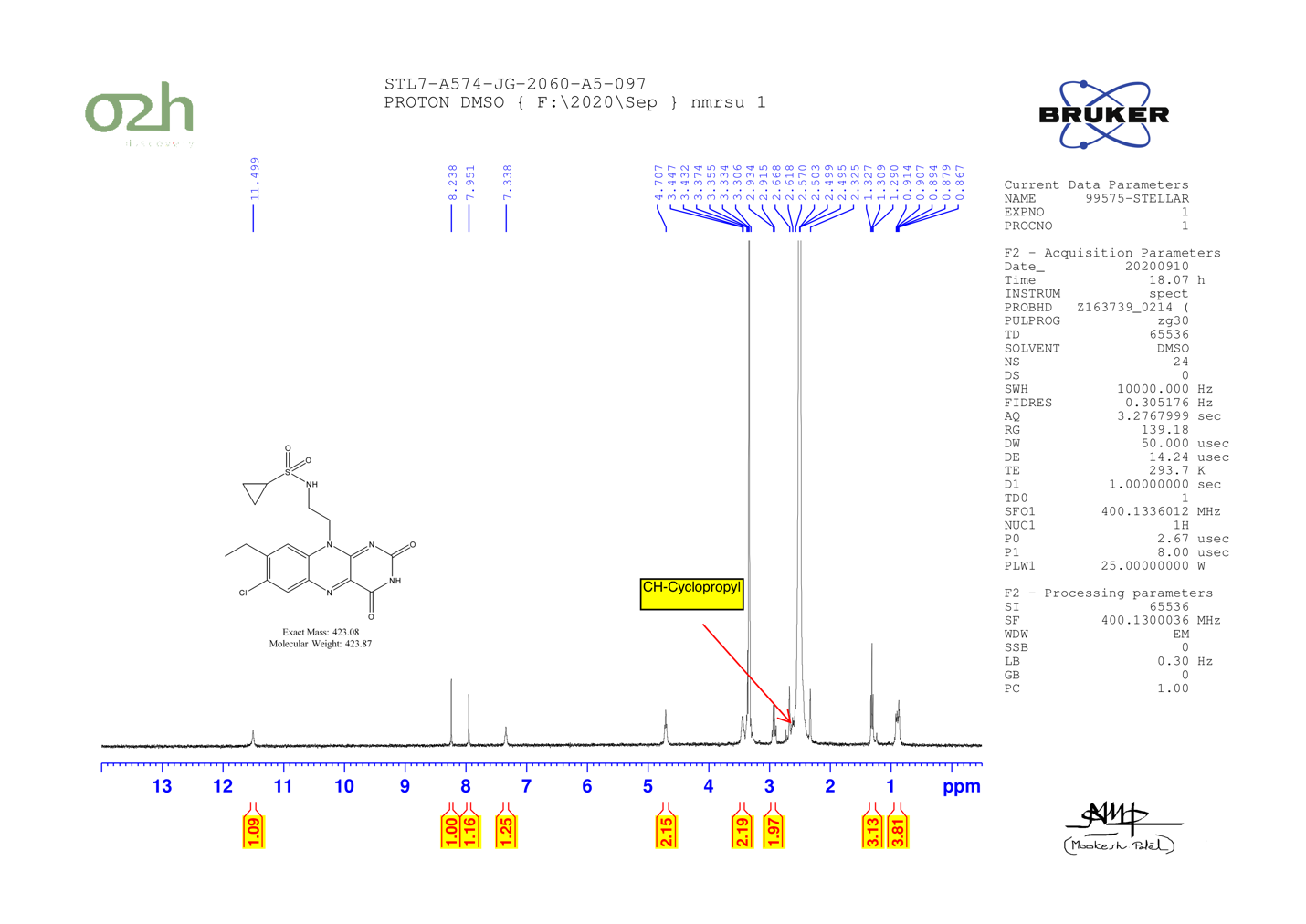

JG-2067

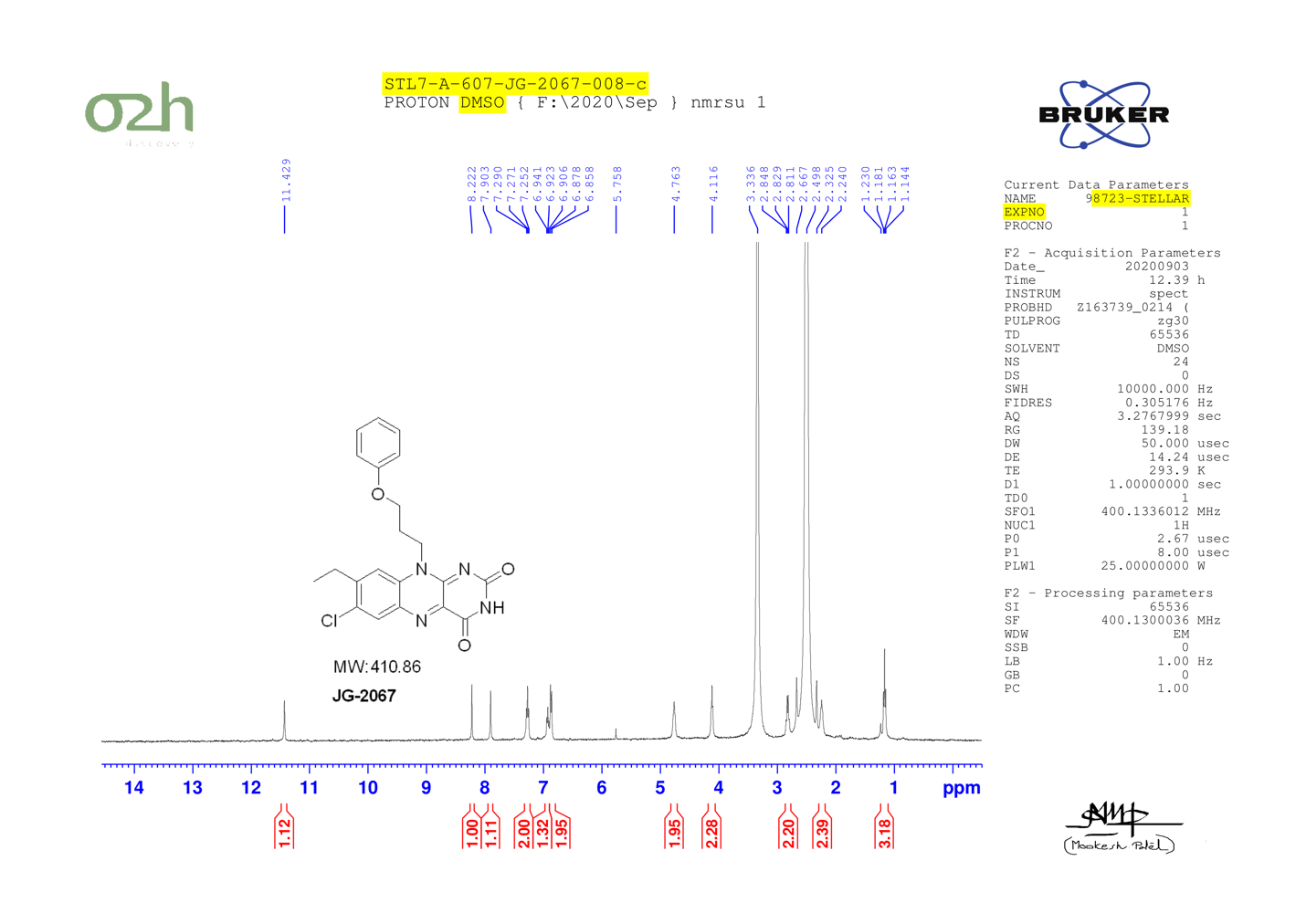

JG-2068

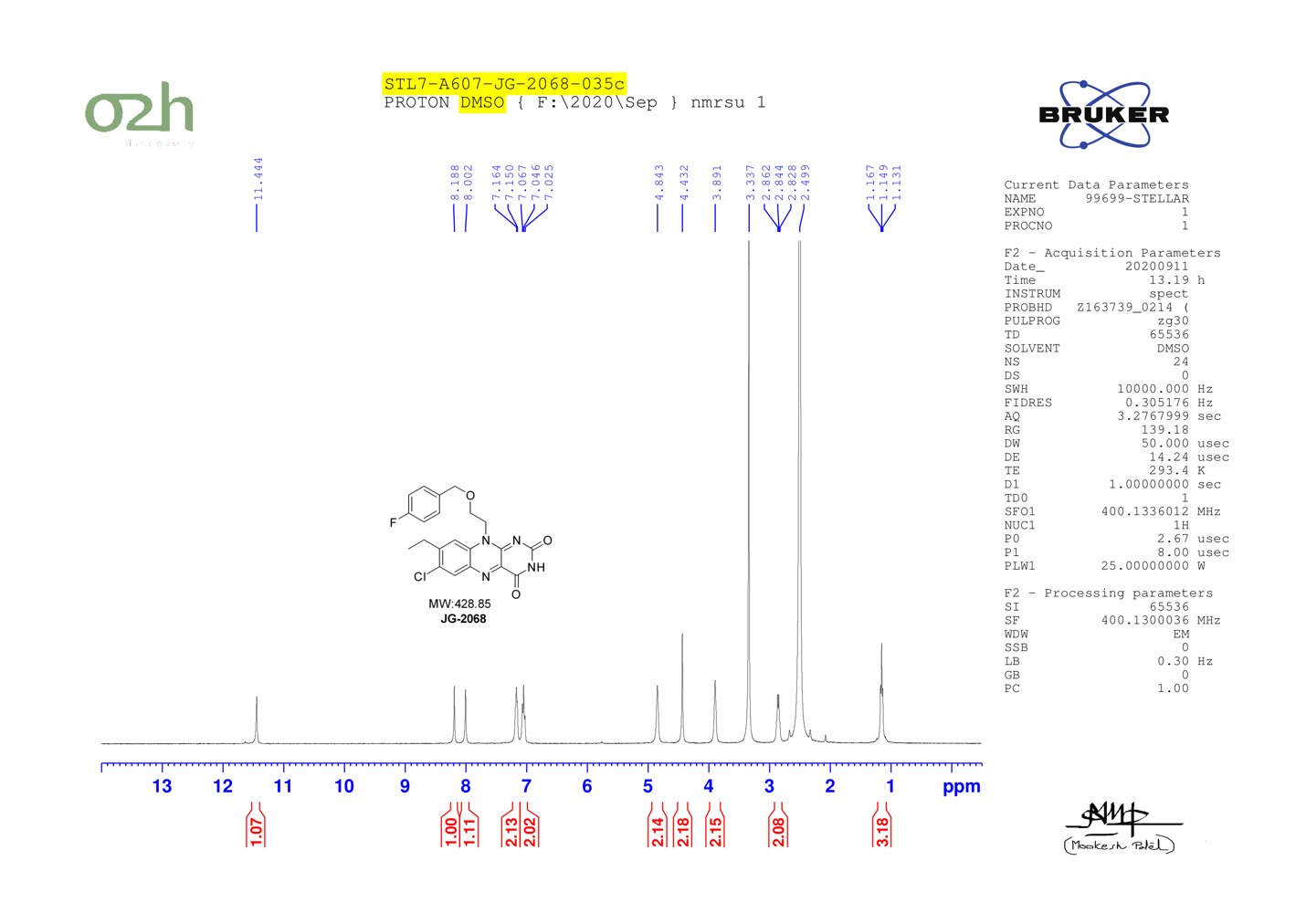

JG-2008

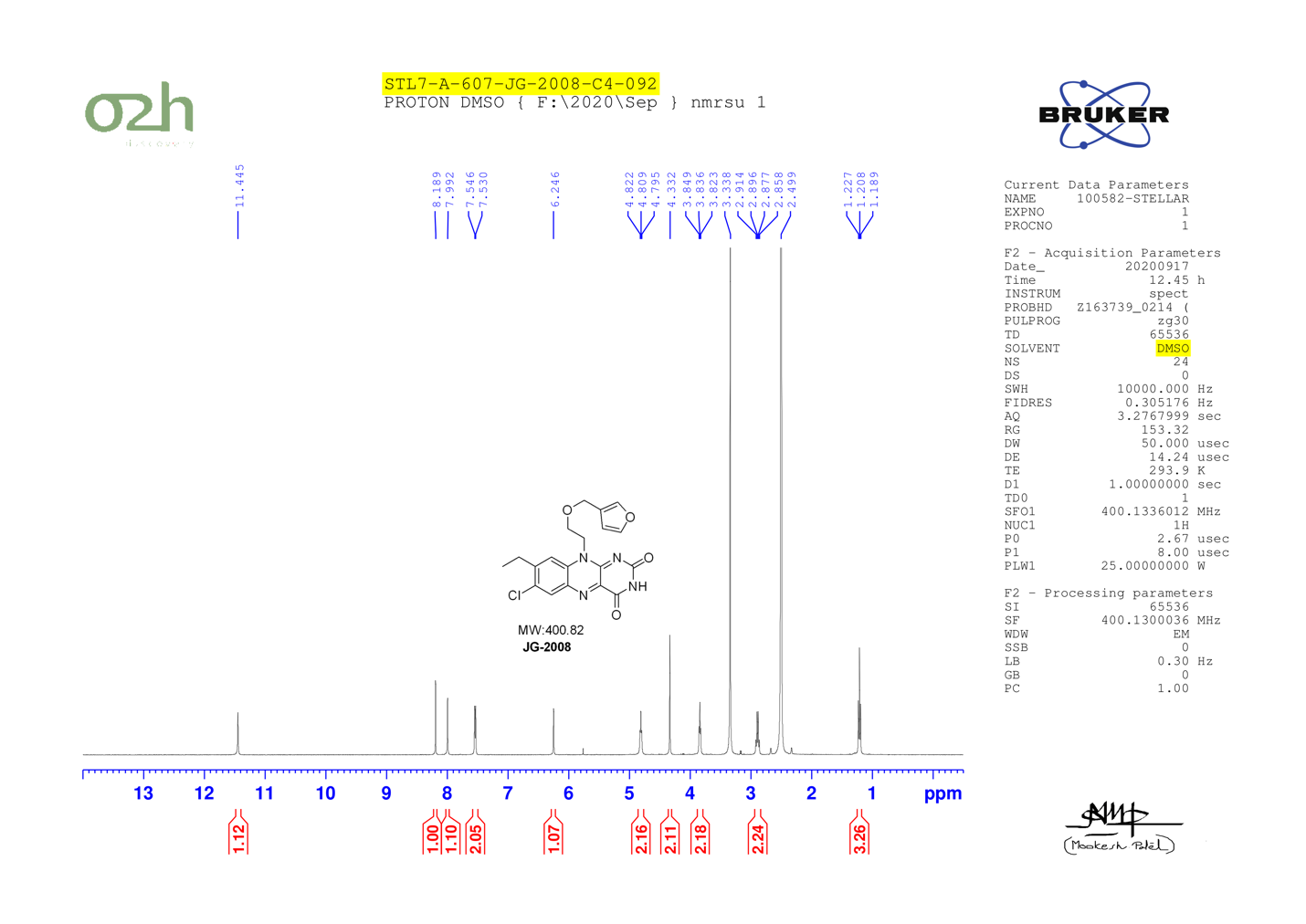

JG-2015

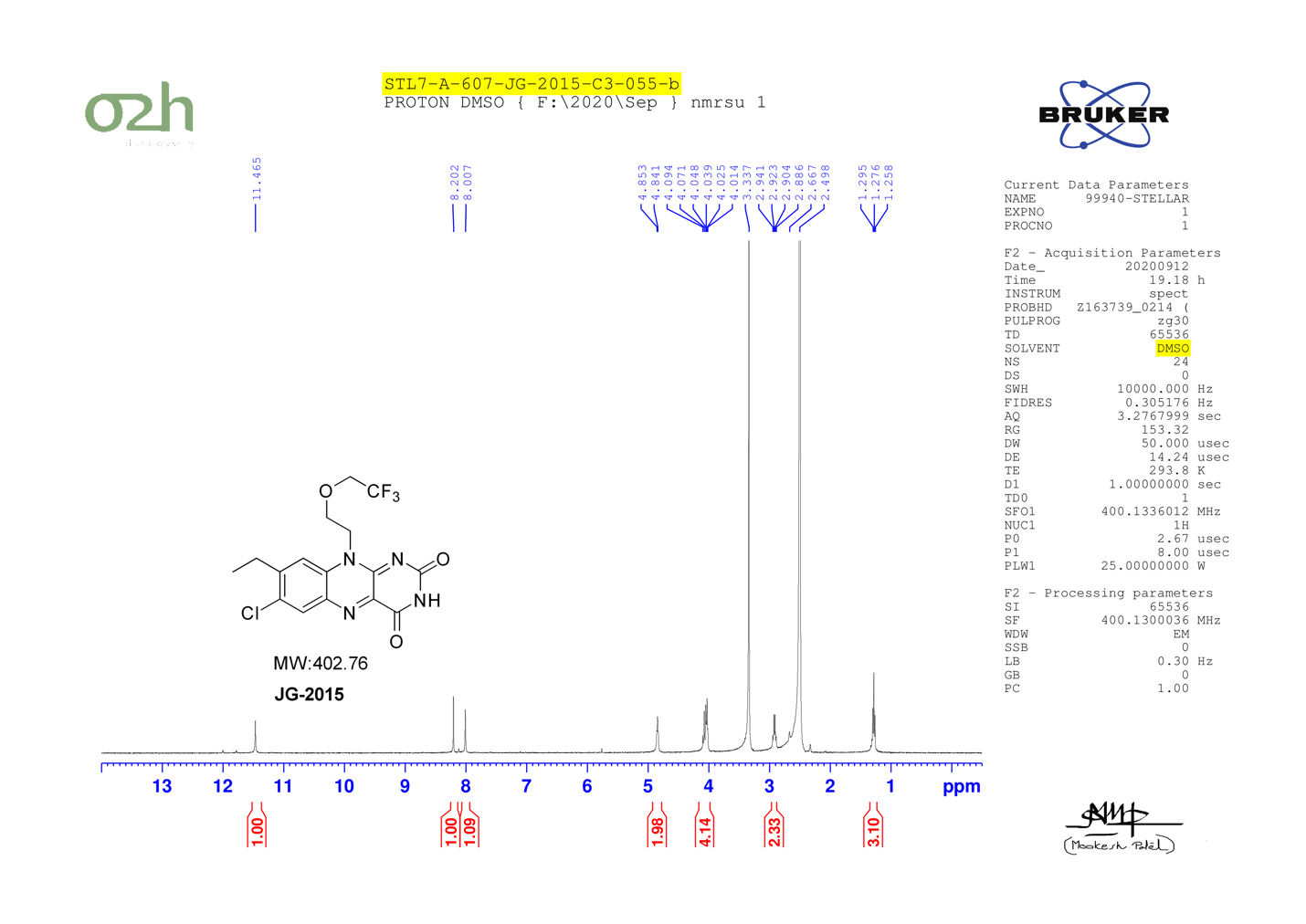

JG-2070

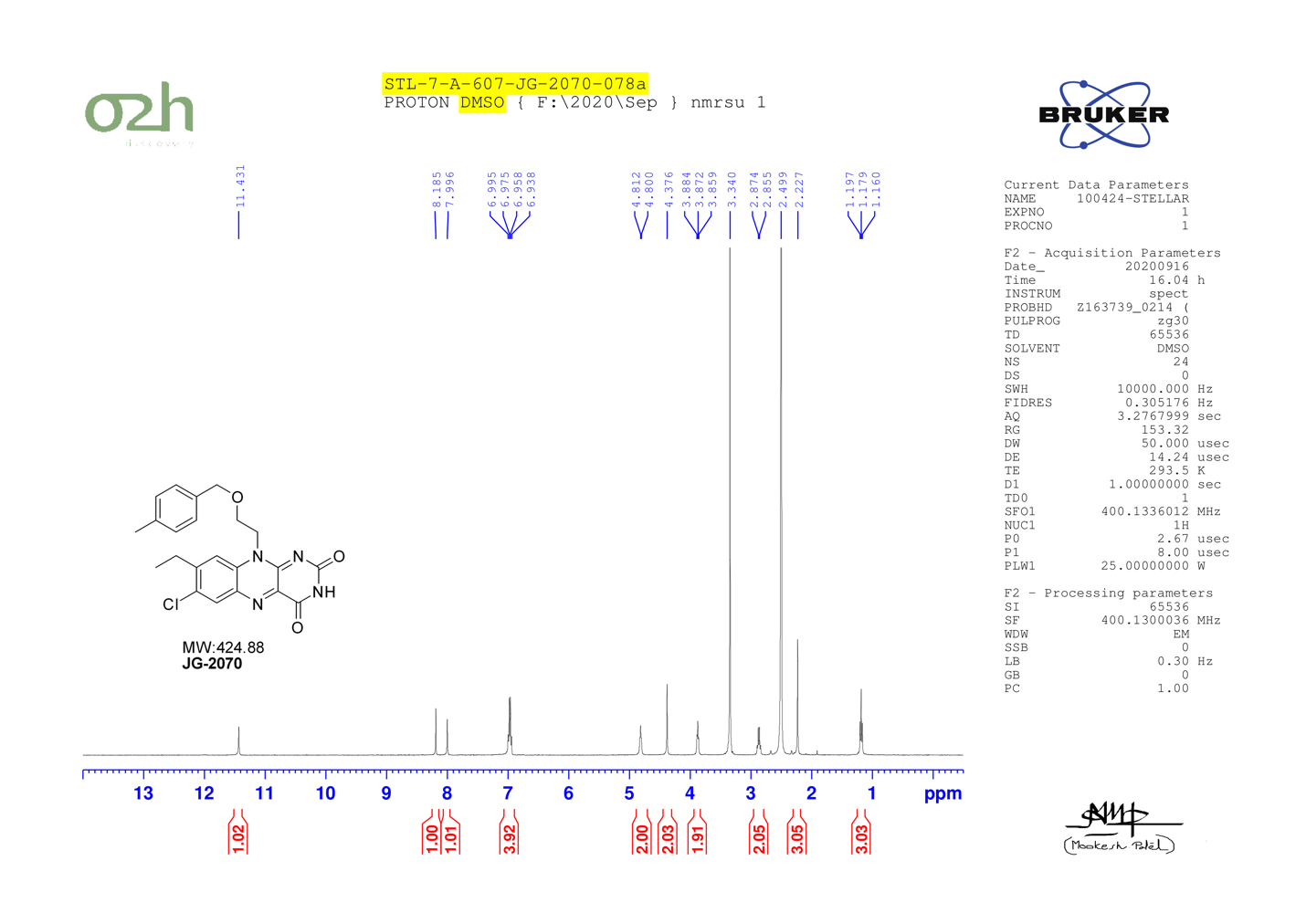

JG-2073

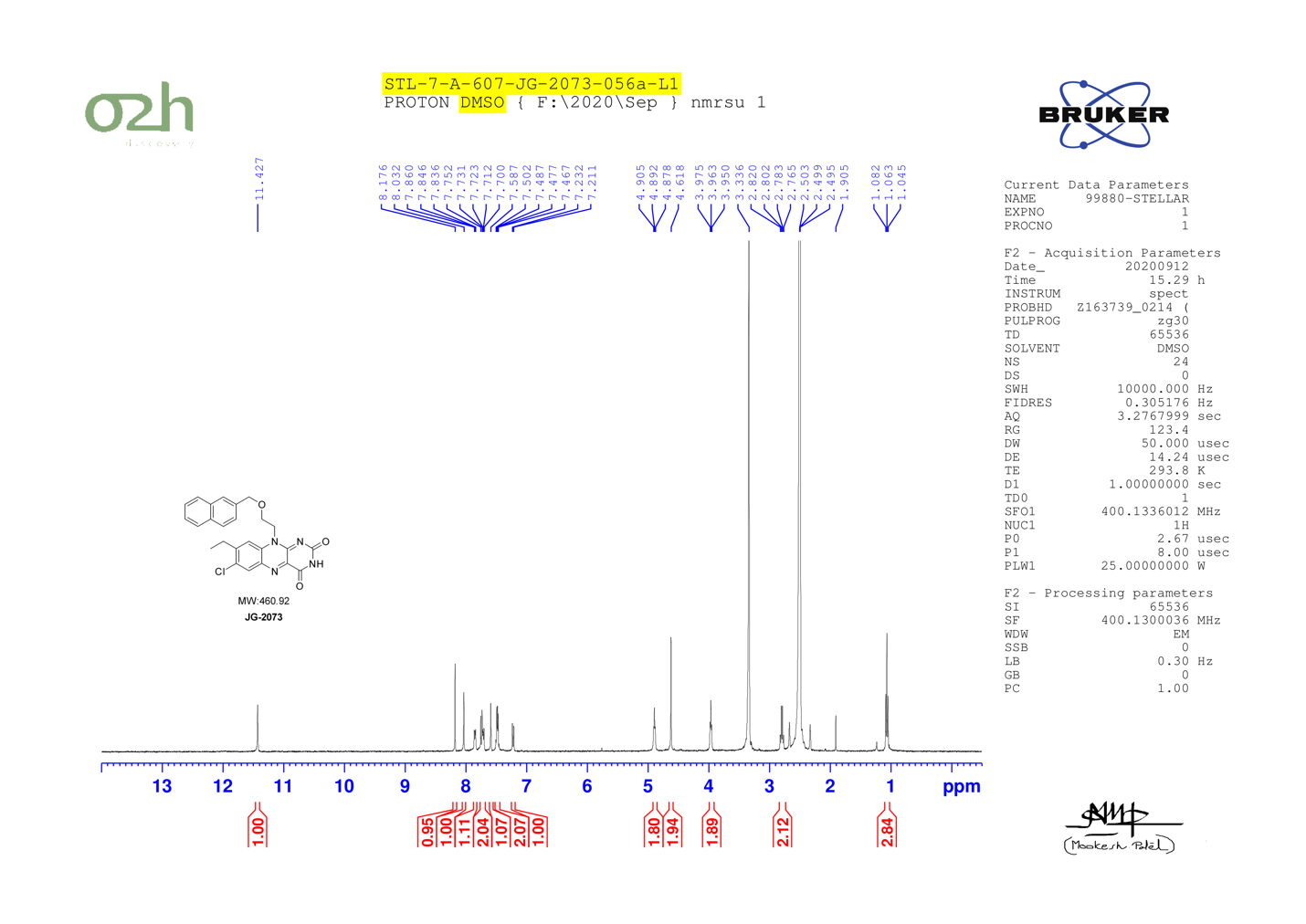

JG-2079

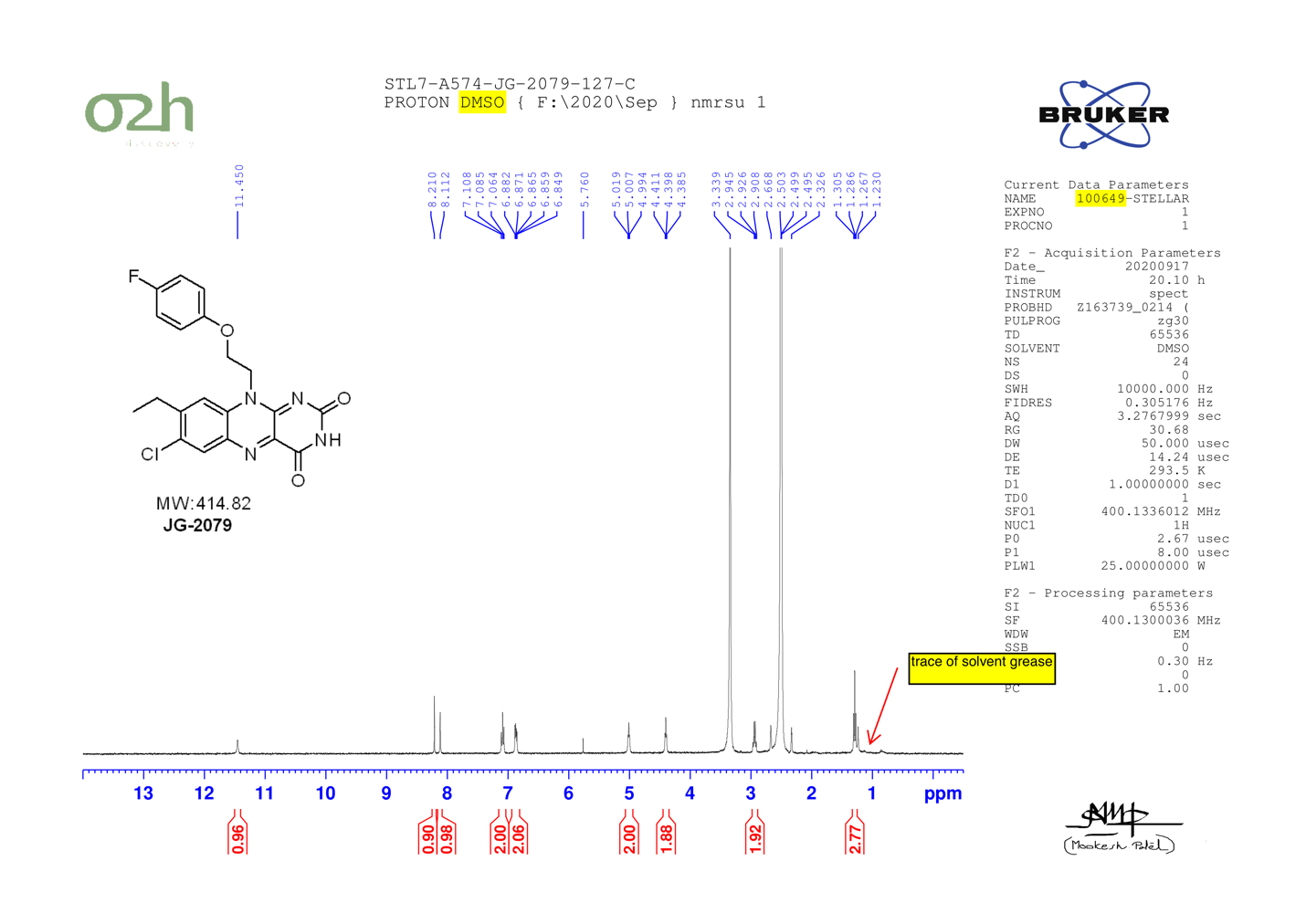

JG-2014

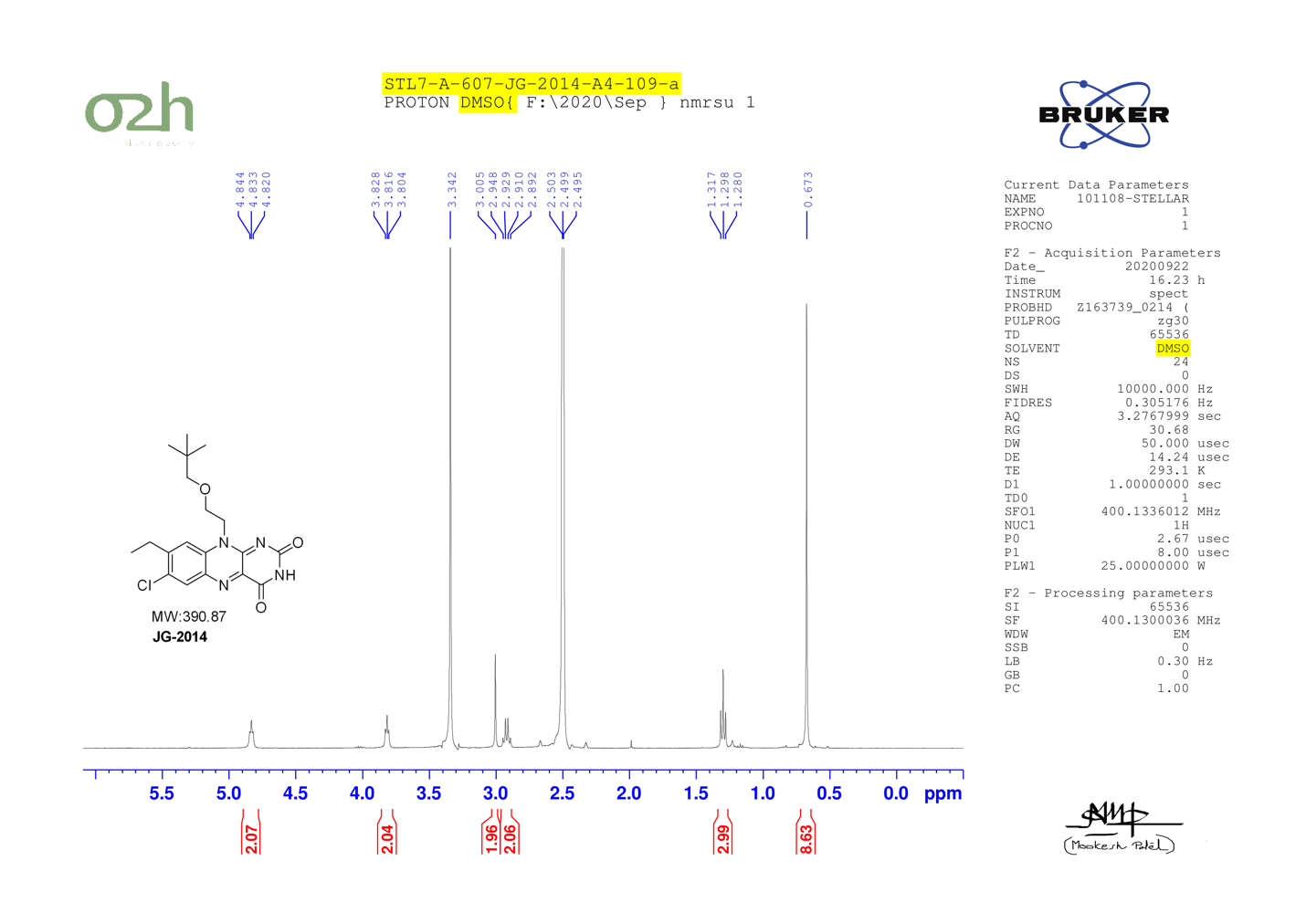

JG-2071

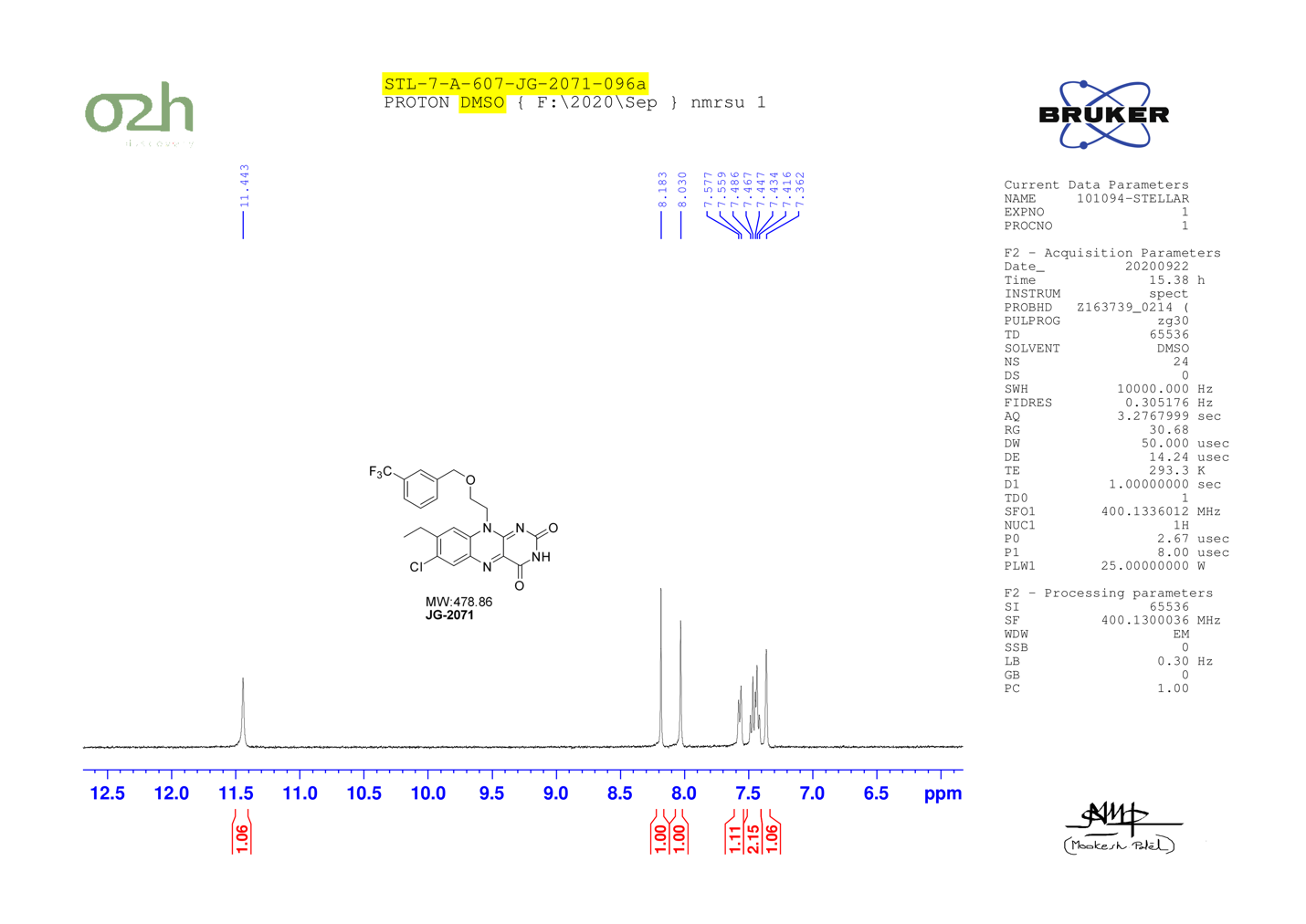

JG-2074

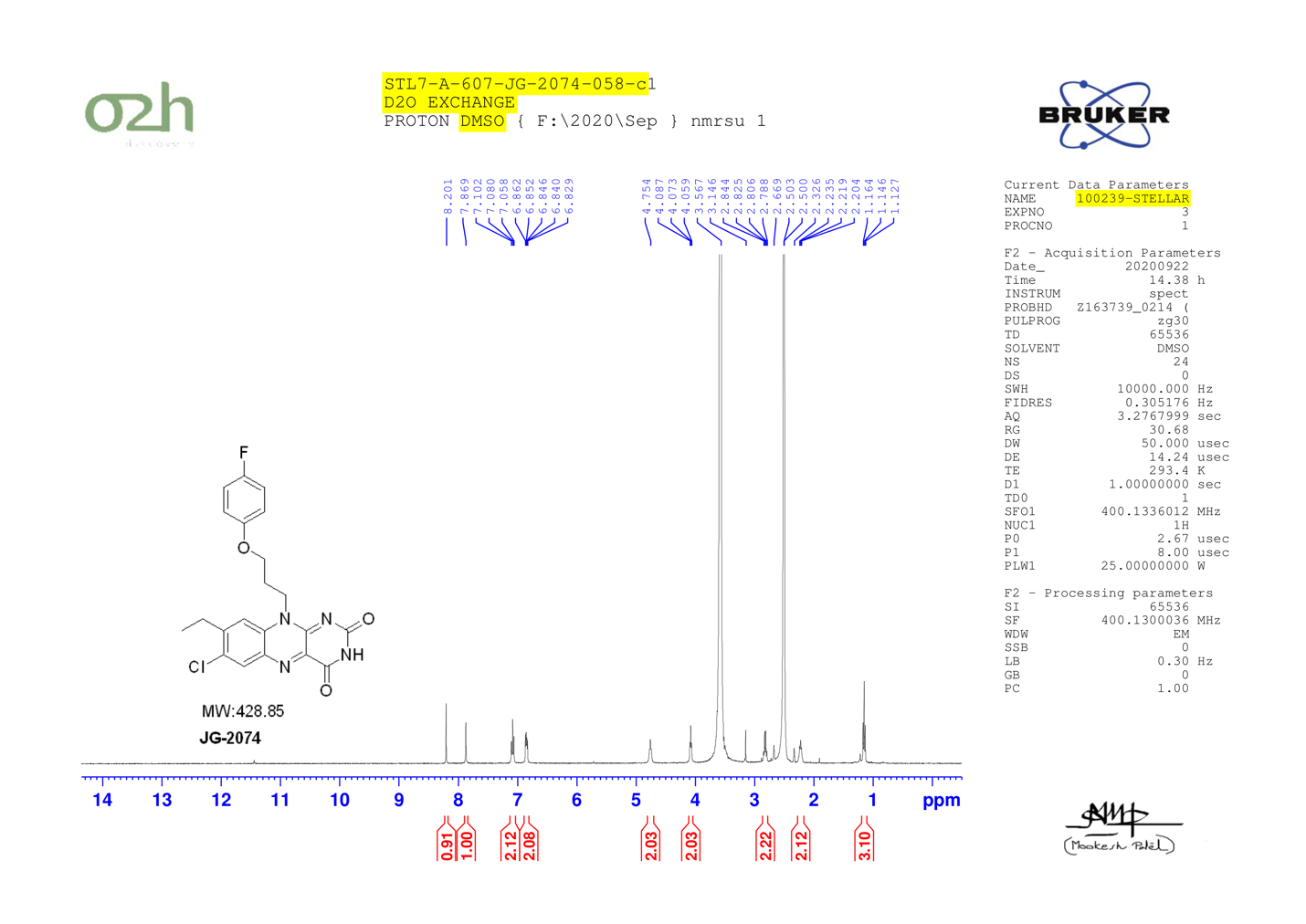

JG-2082

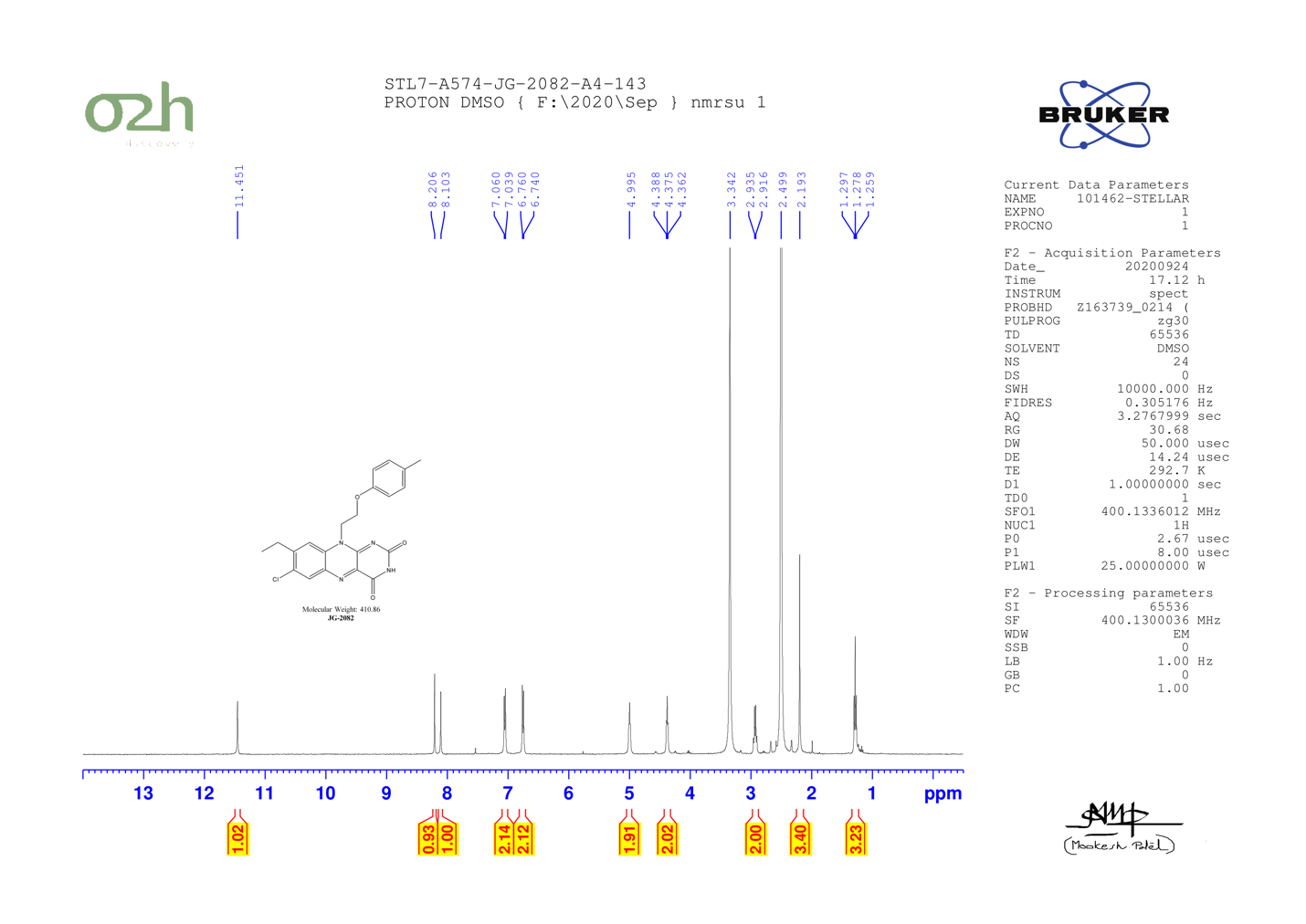

JG-2016

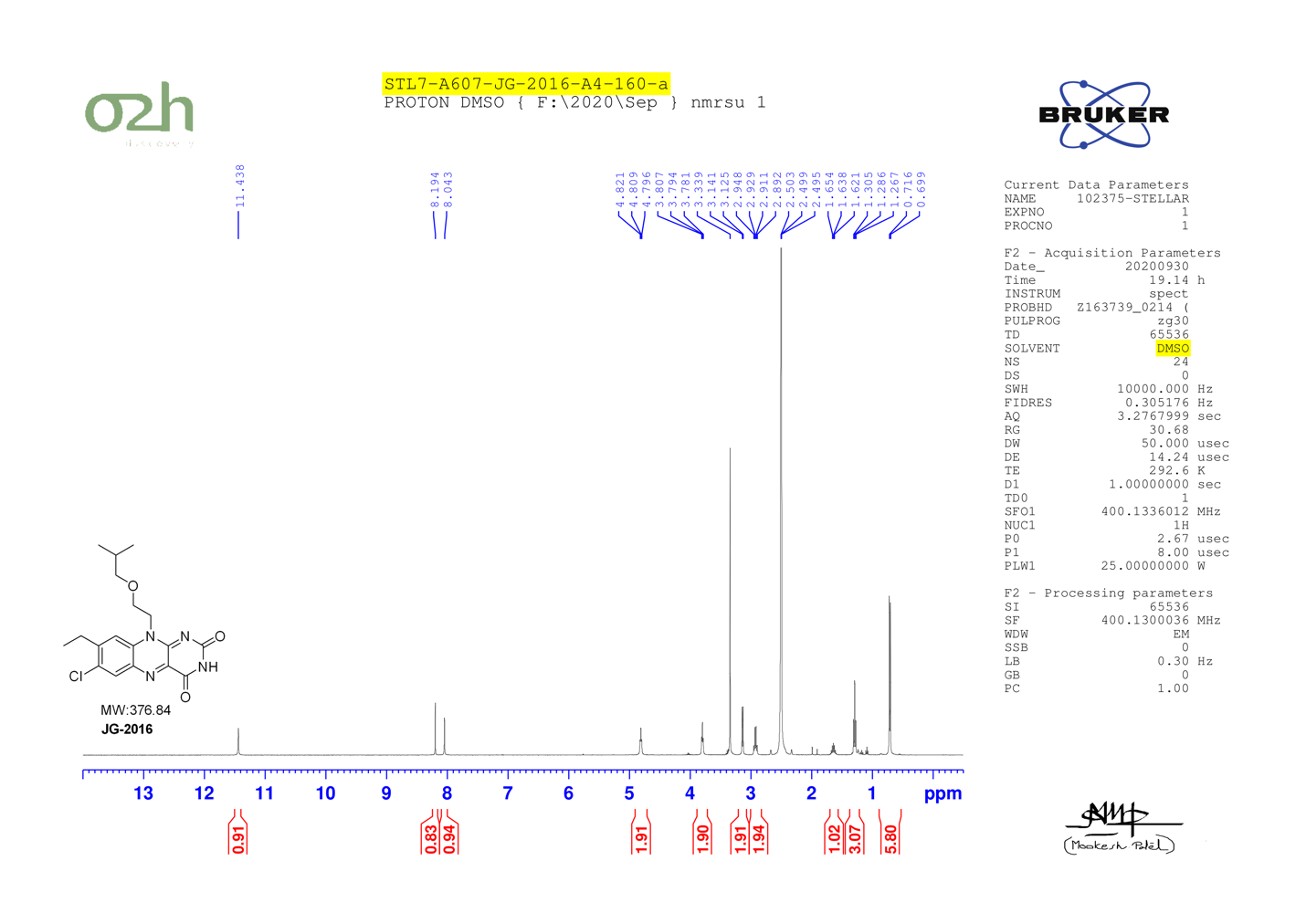

JG-2076

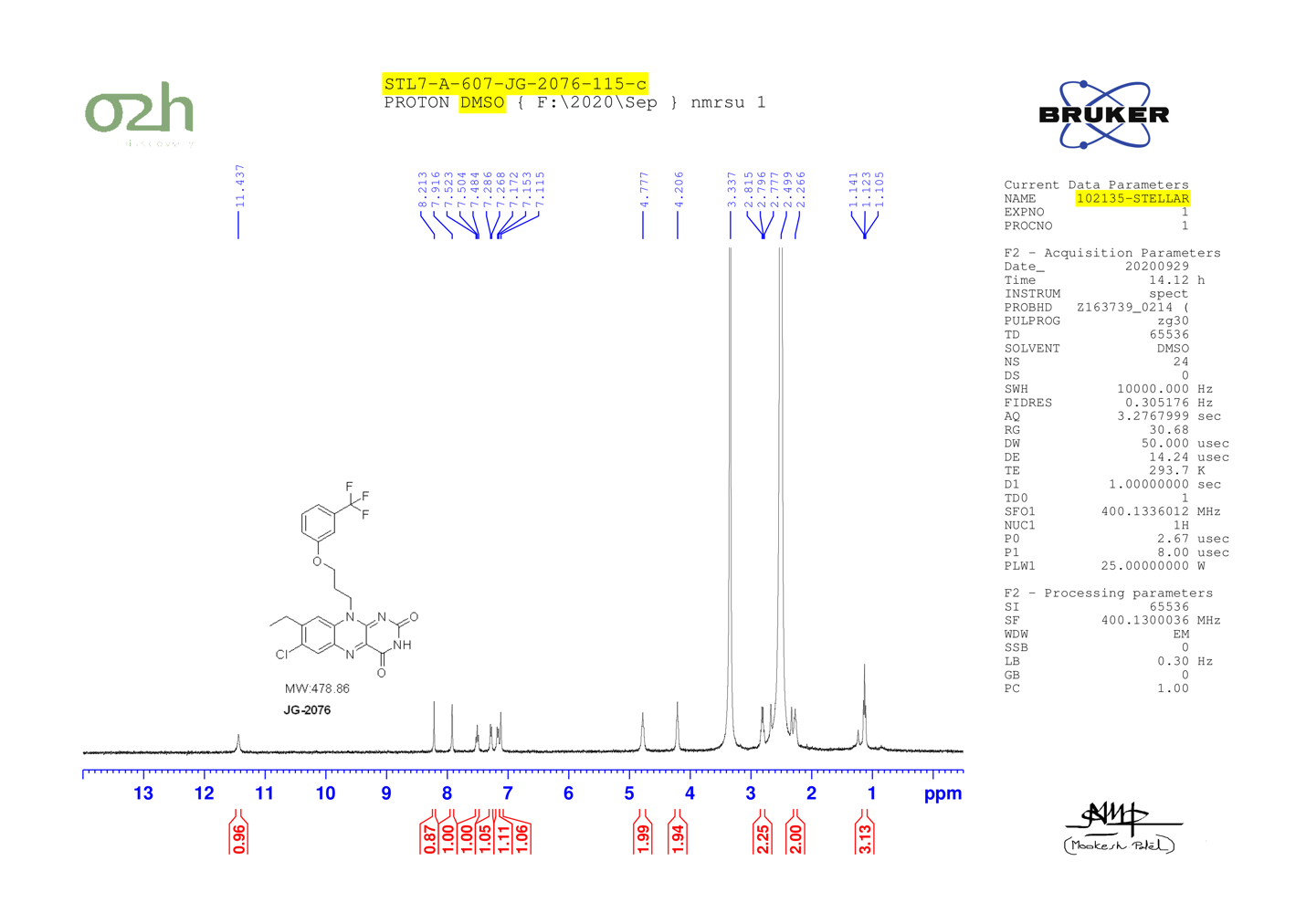

JG-2084

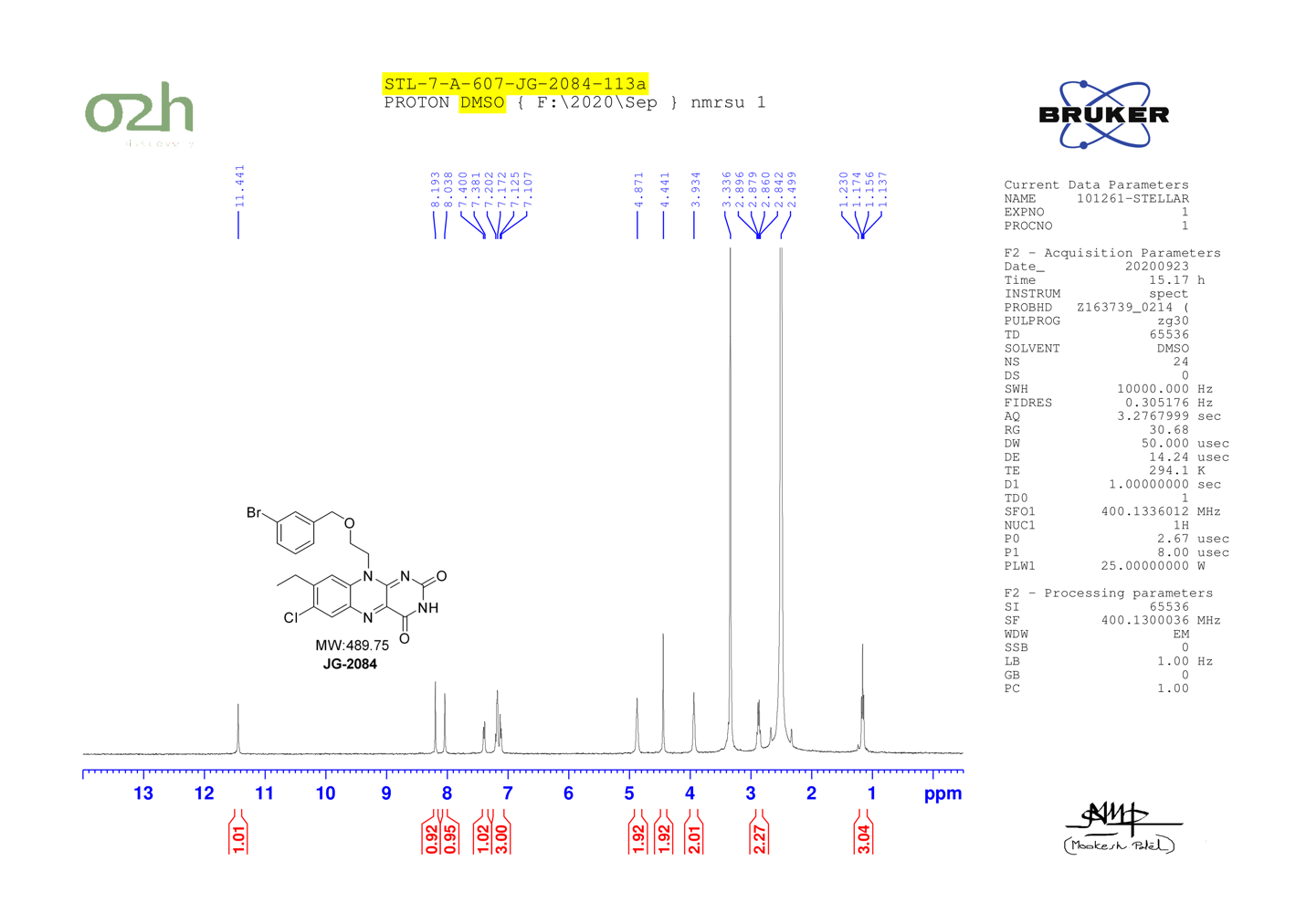

JG-2006

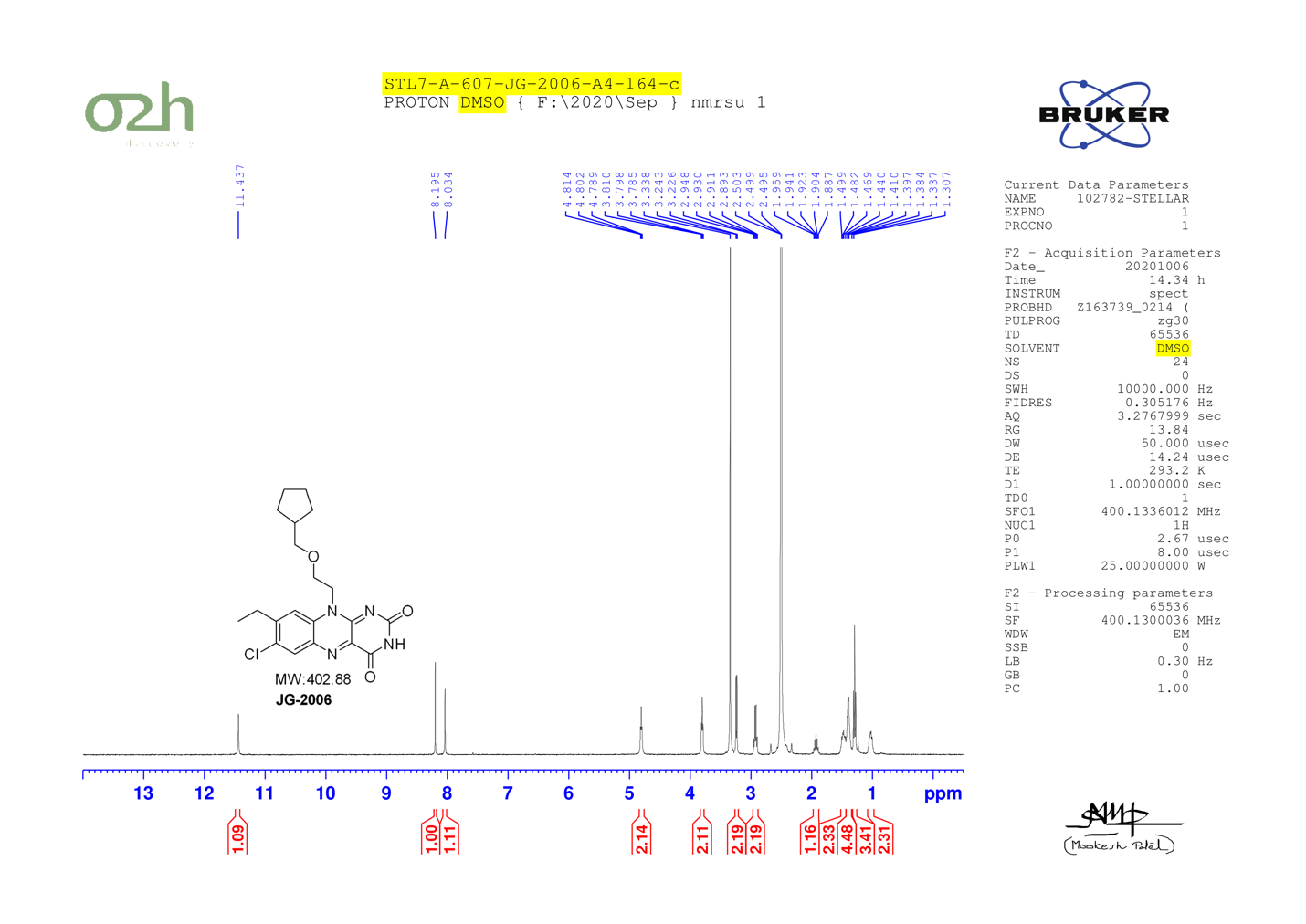

JG-2009

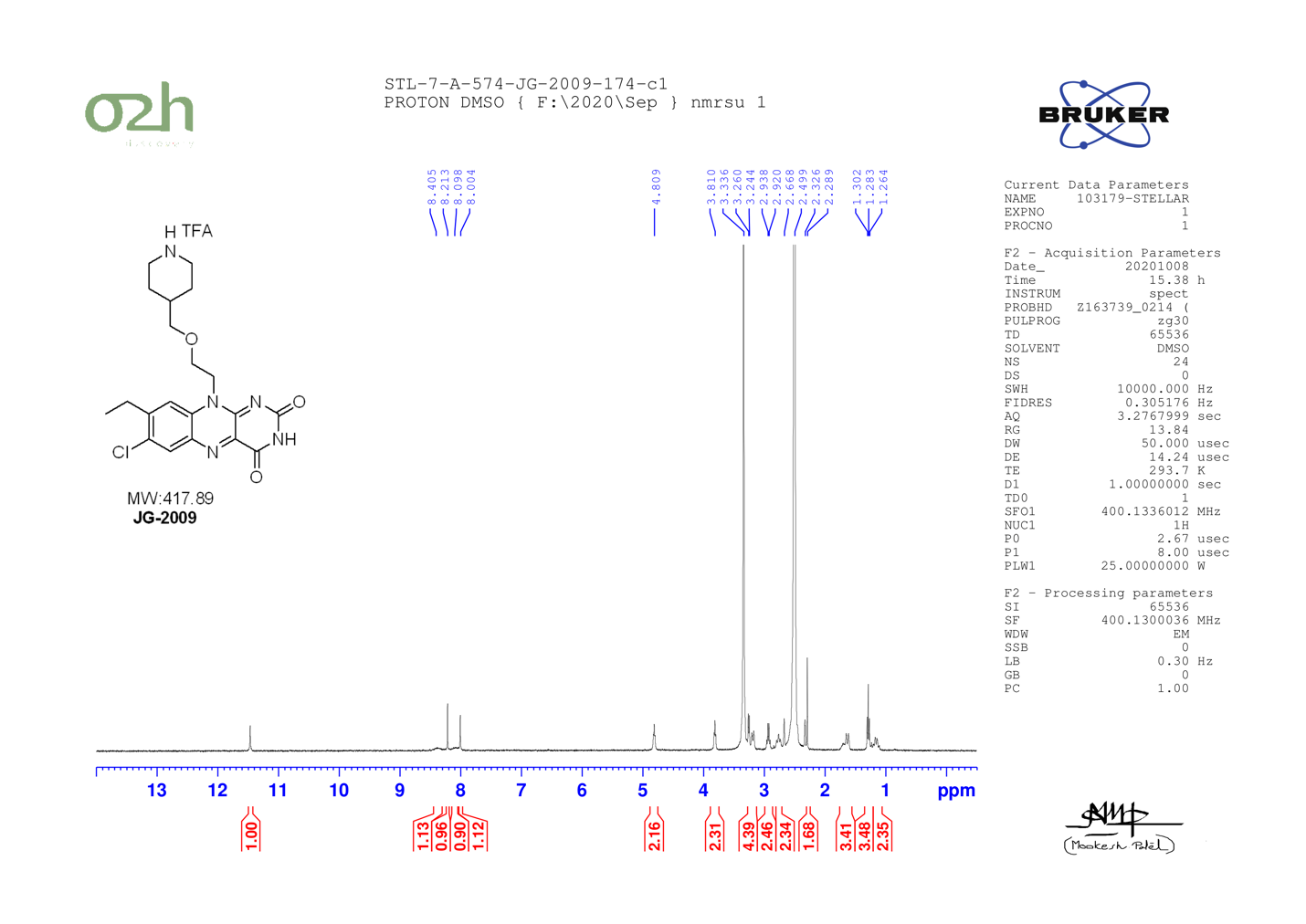

JG-2072

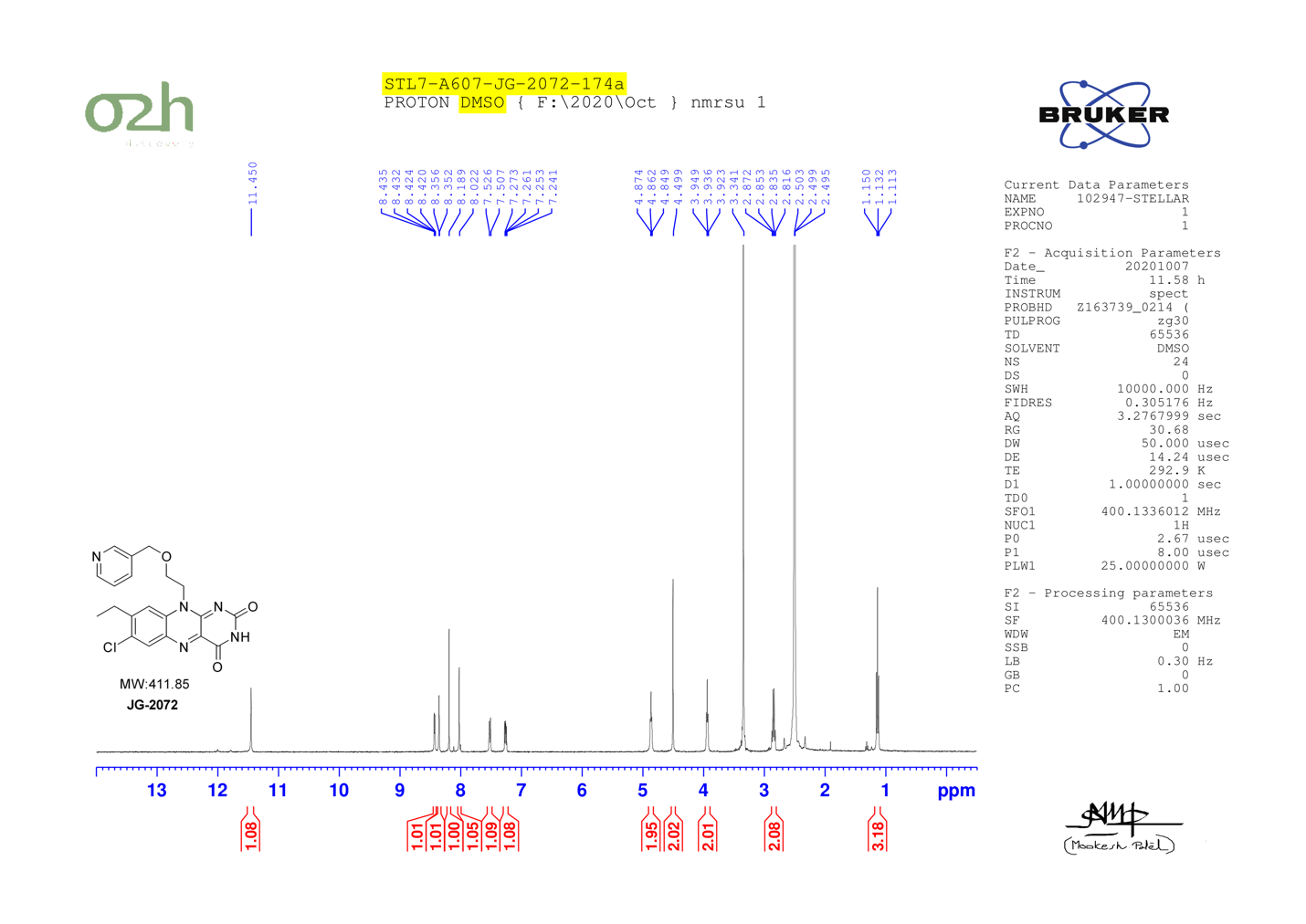

JG-2078

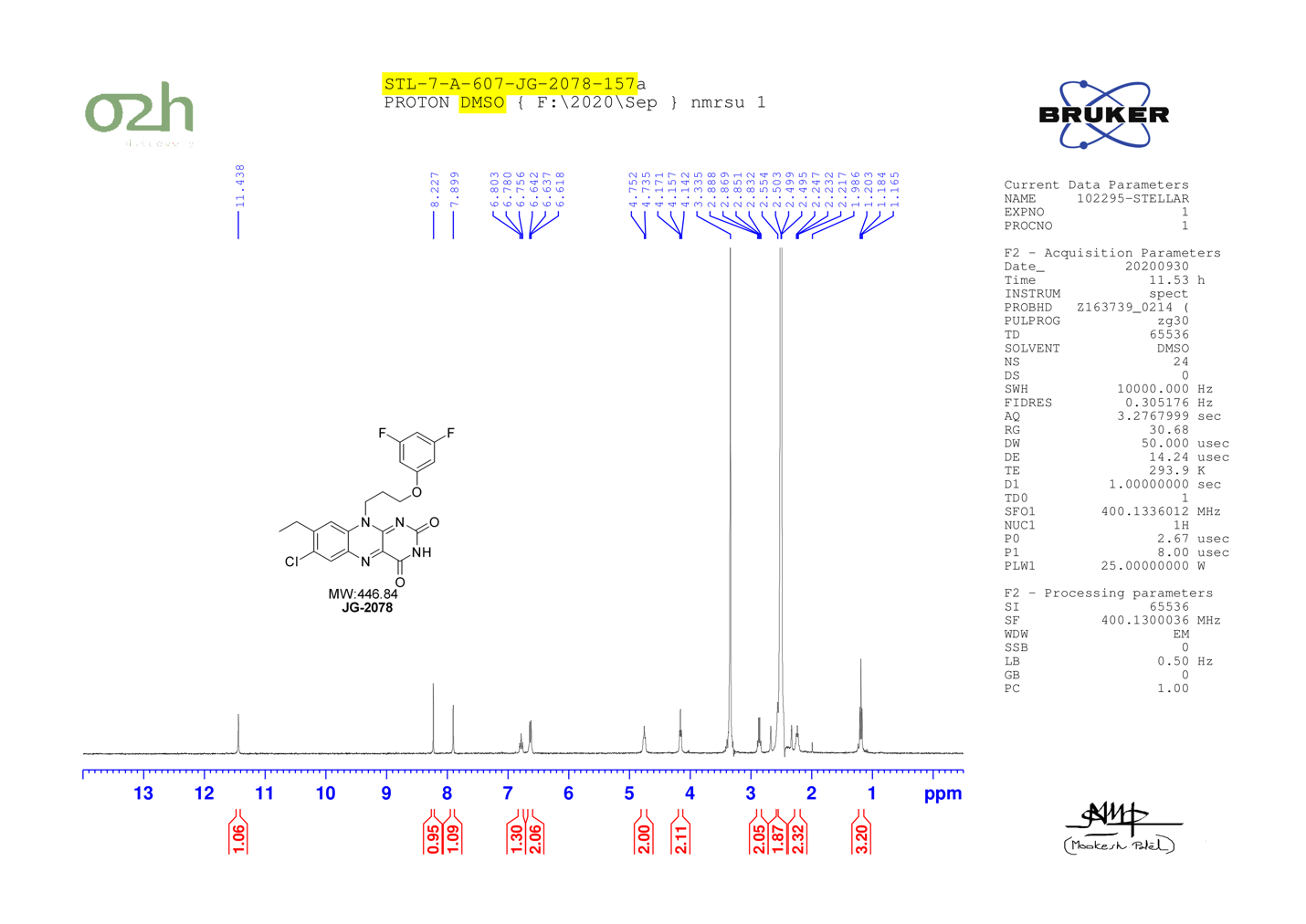

JG-2088

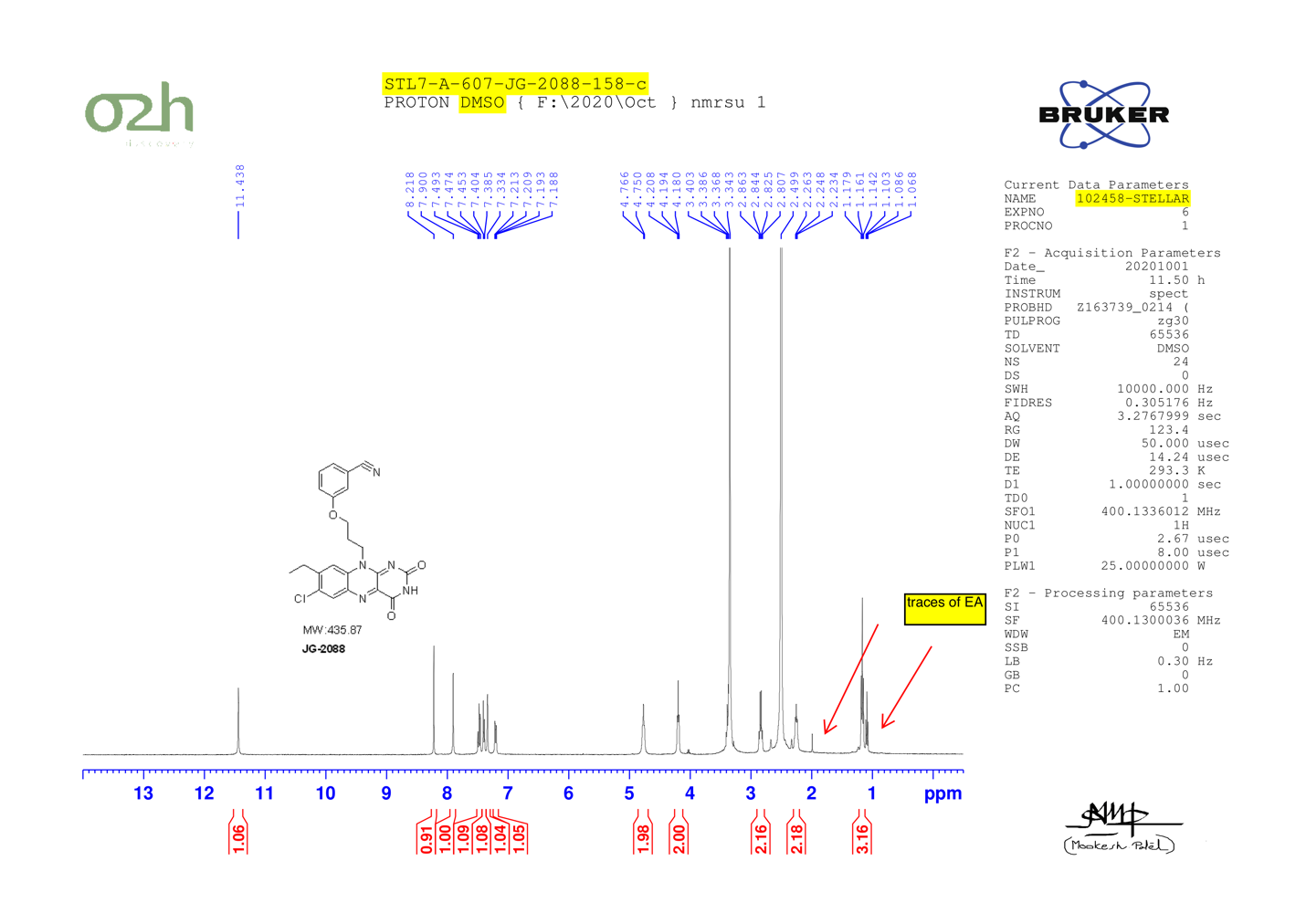

JG-2092

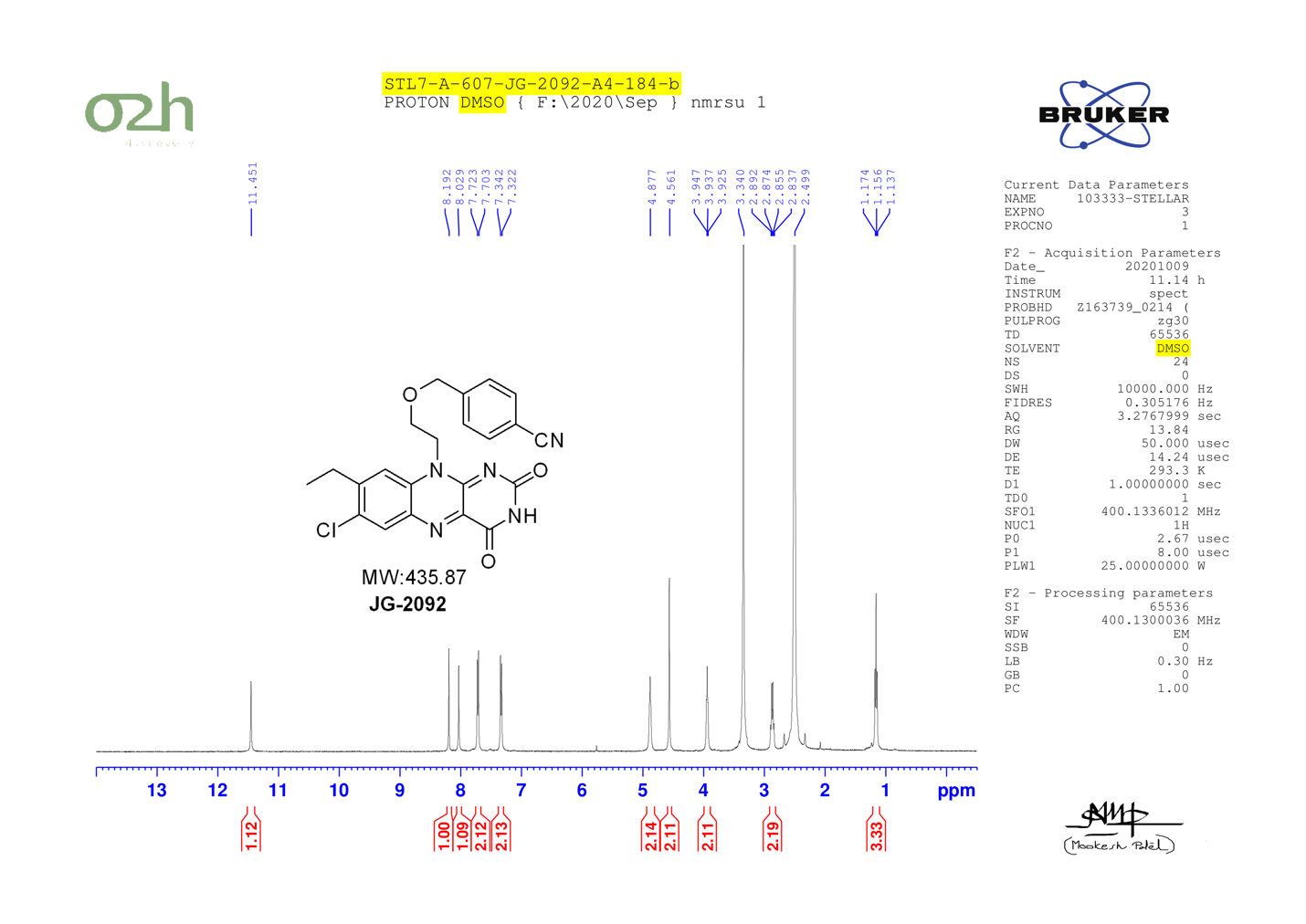

JG-2069

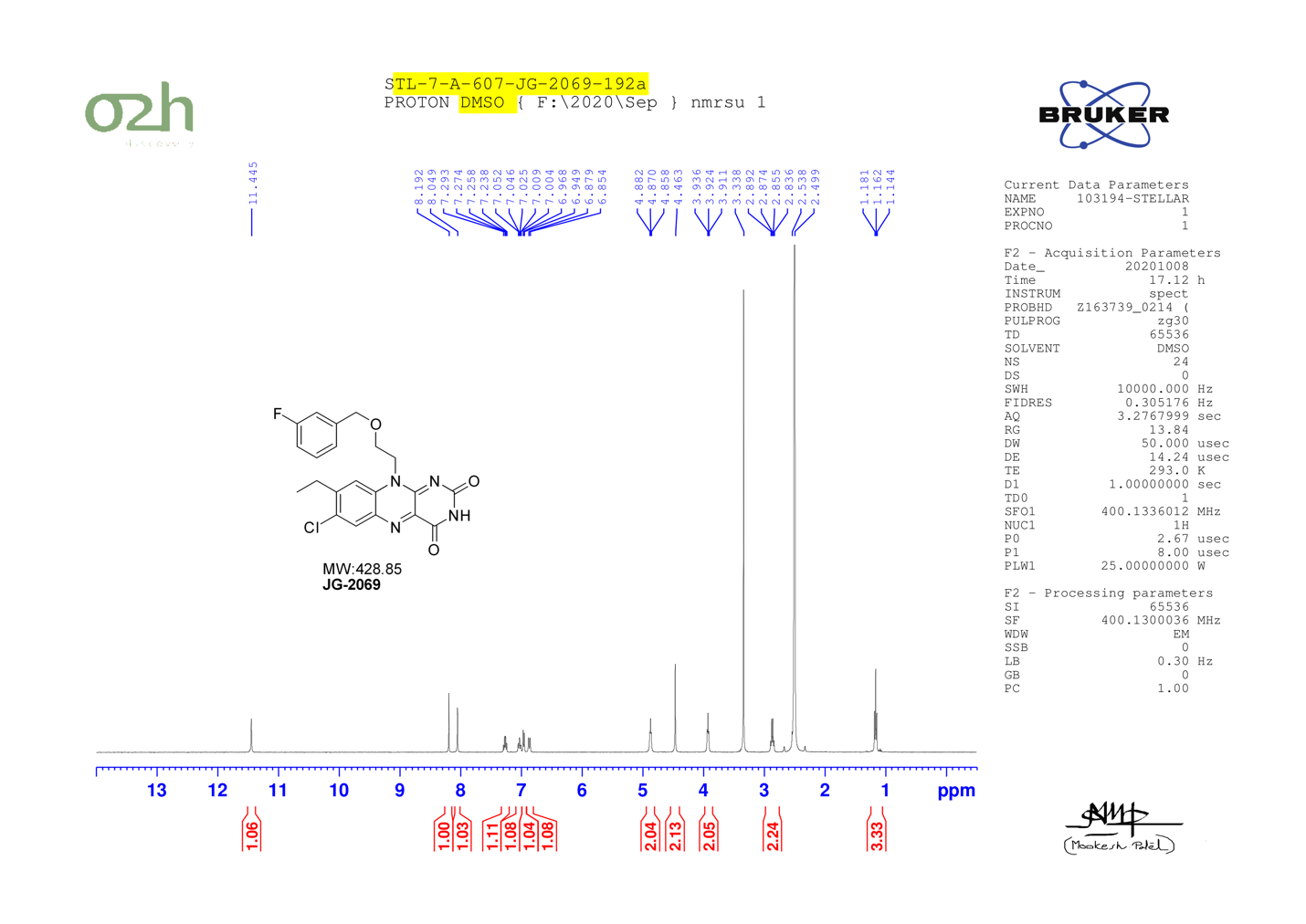

JG-2077

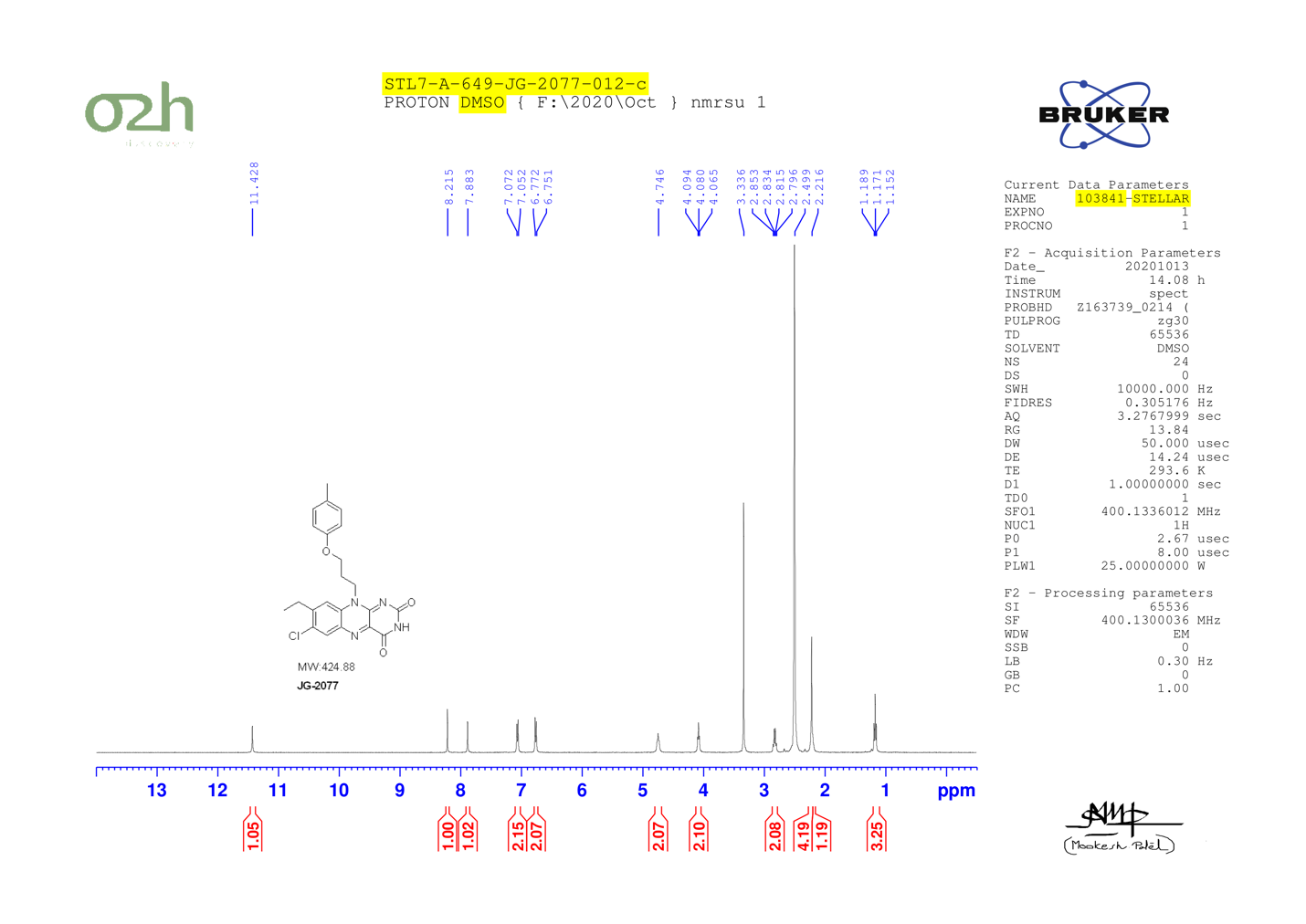

JG-2080

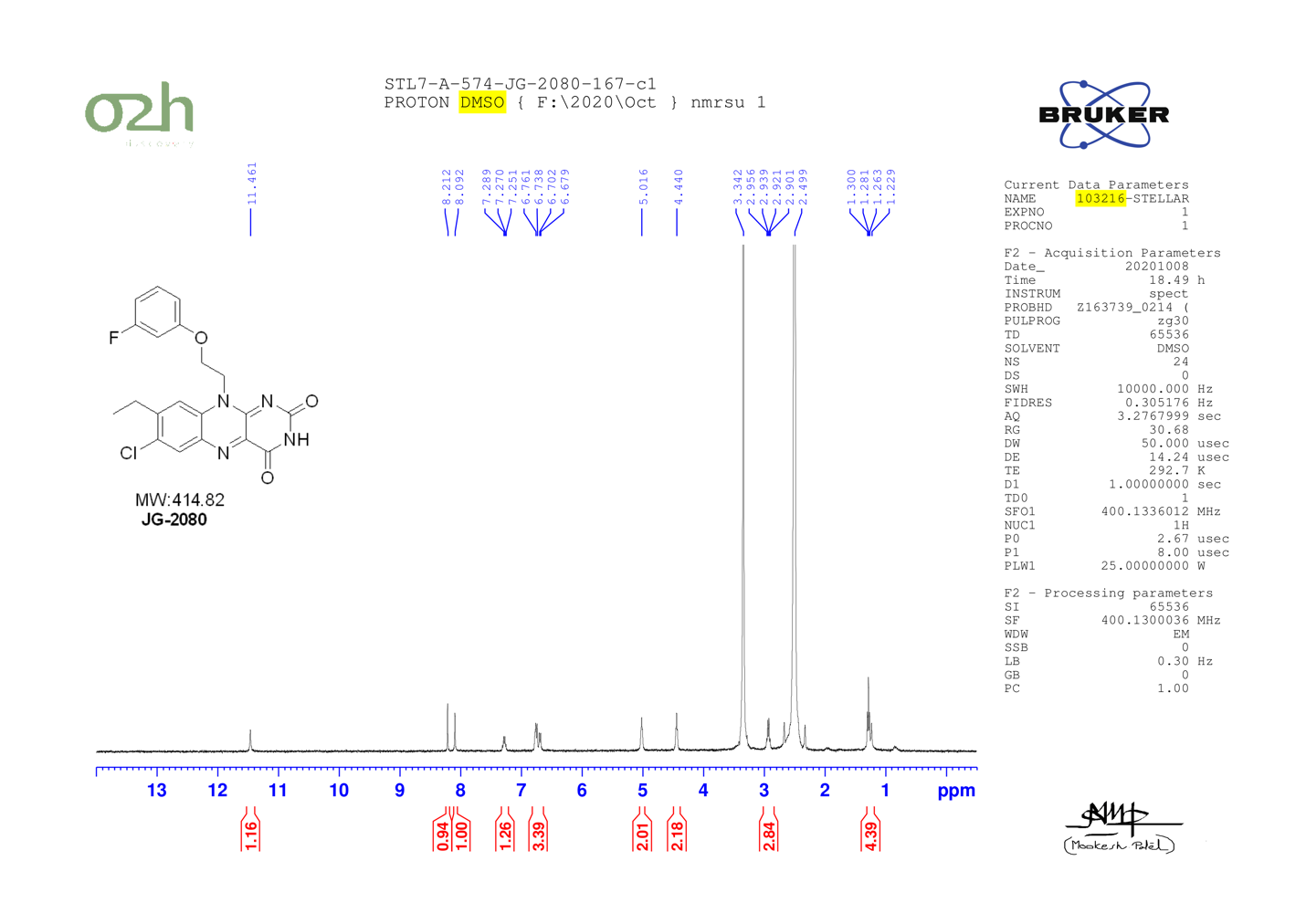

JG-2081

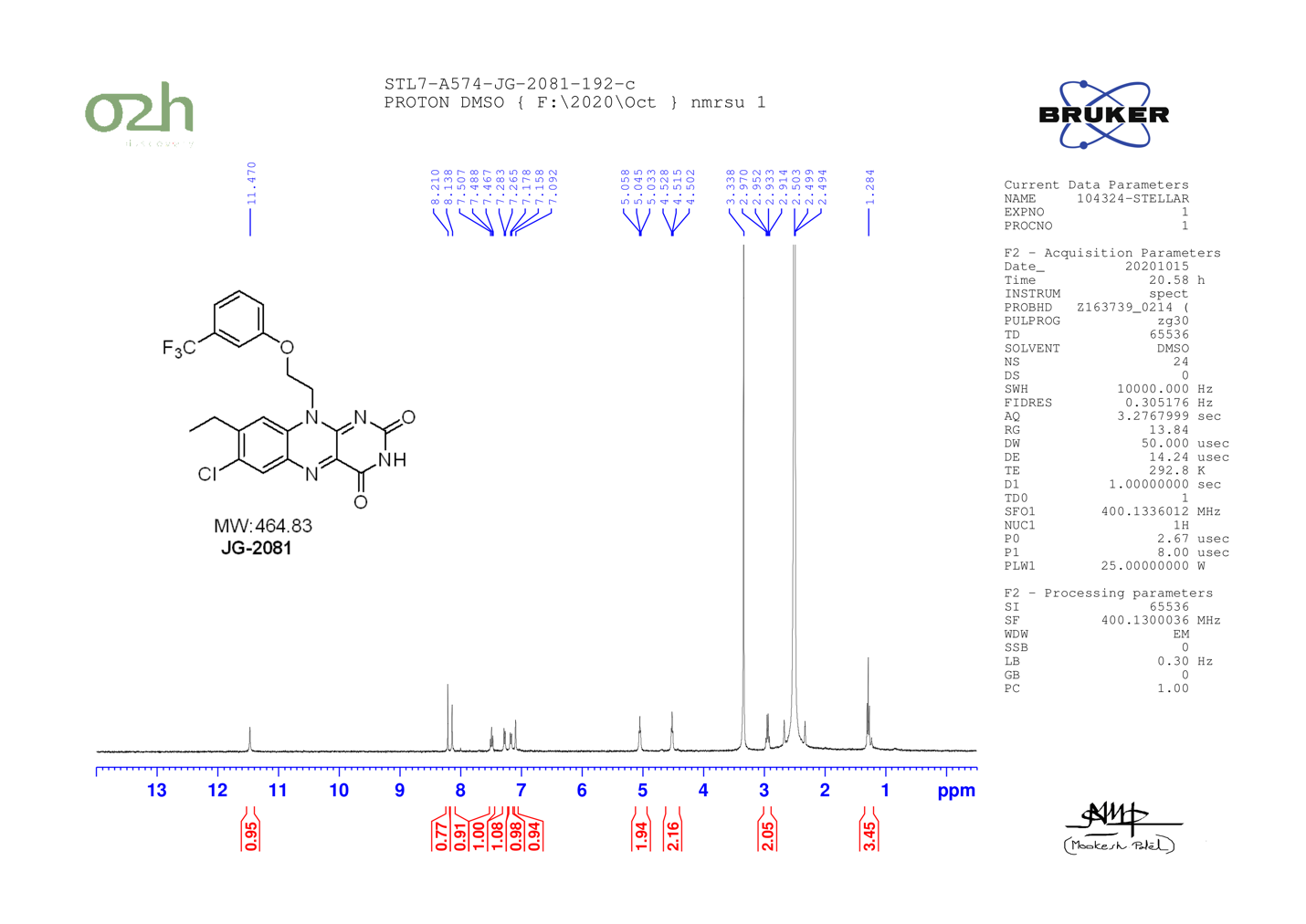

JG-2083

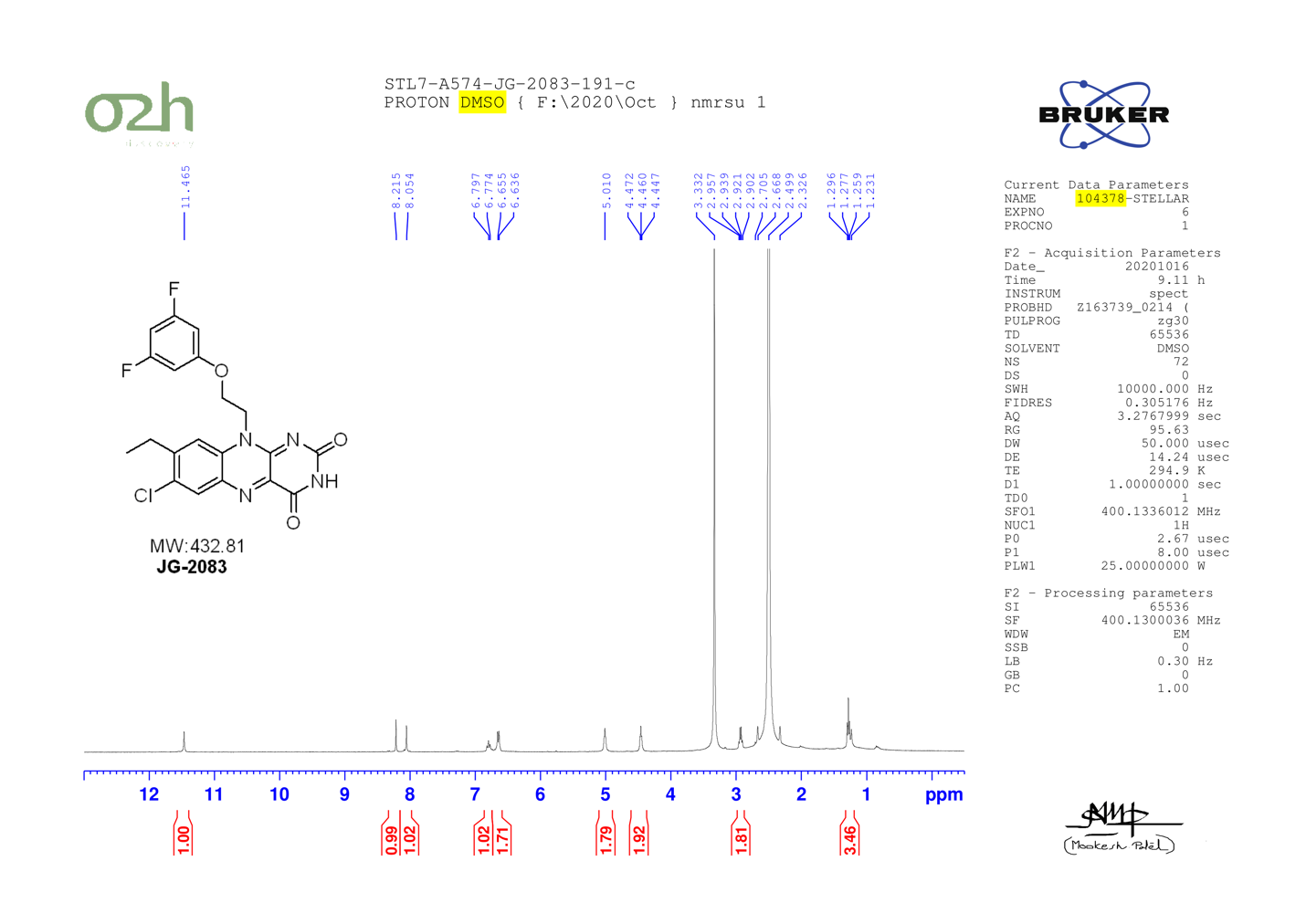

JG-2085

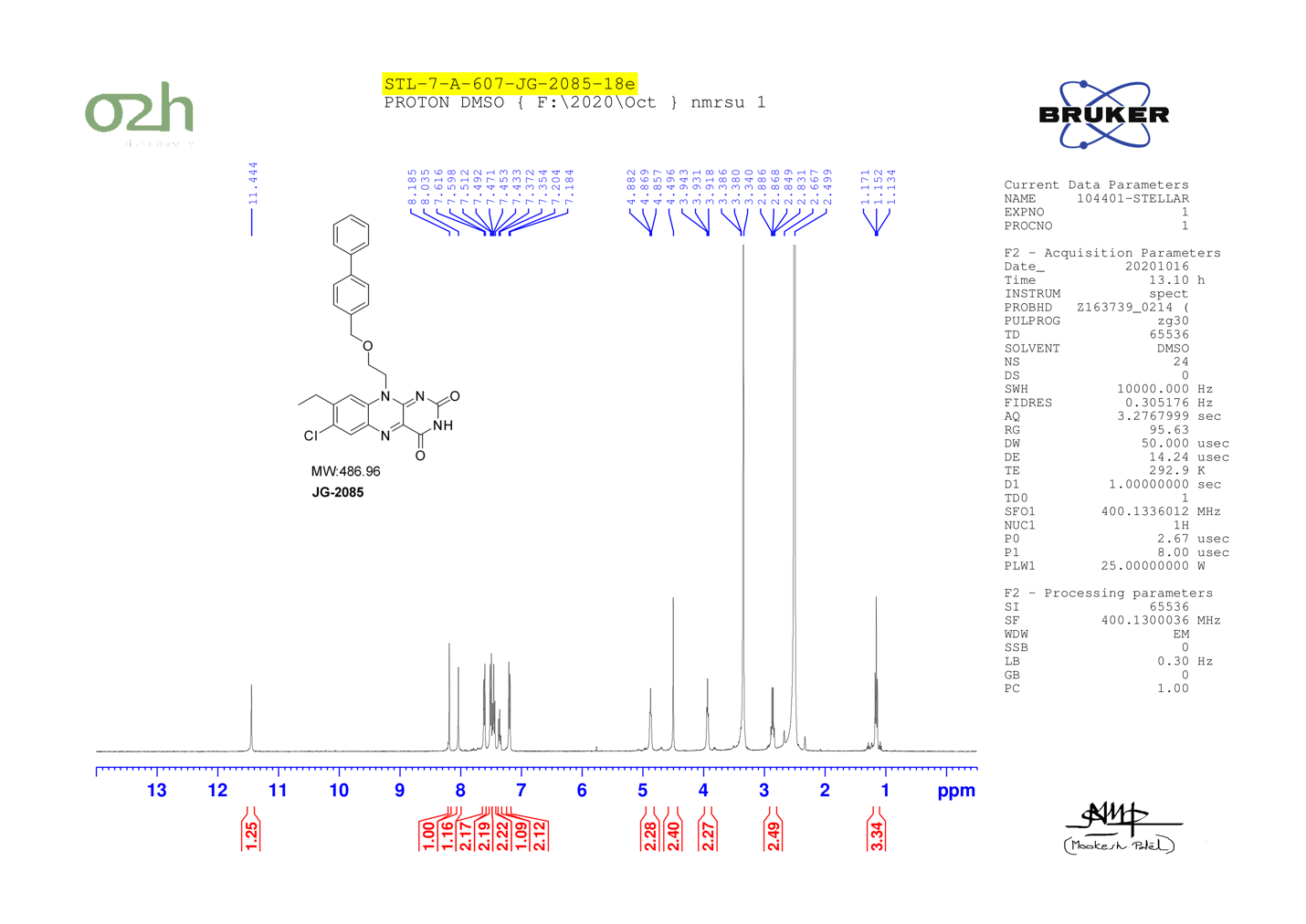

JG-2086

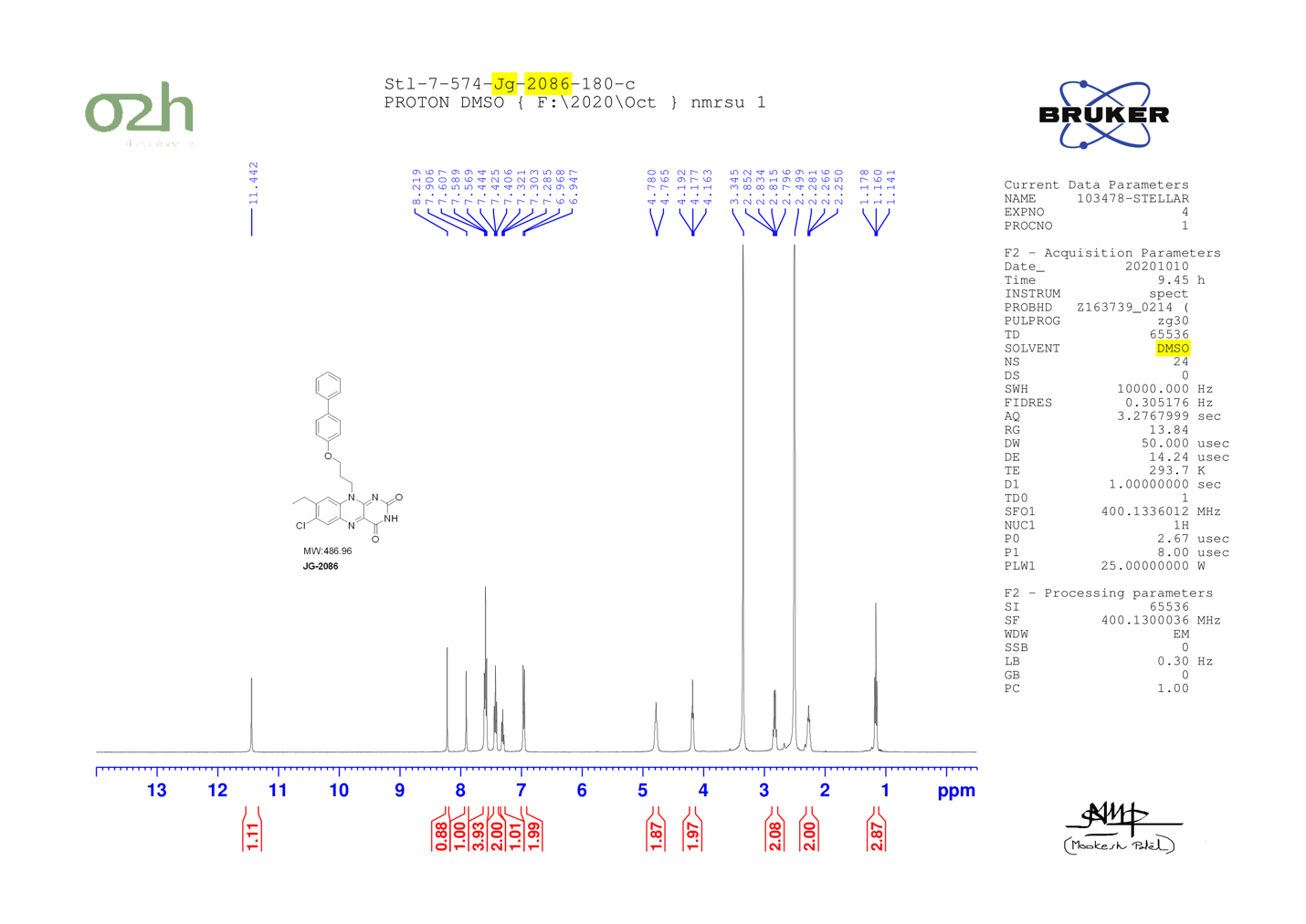

JG-2087

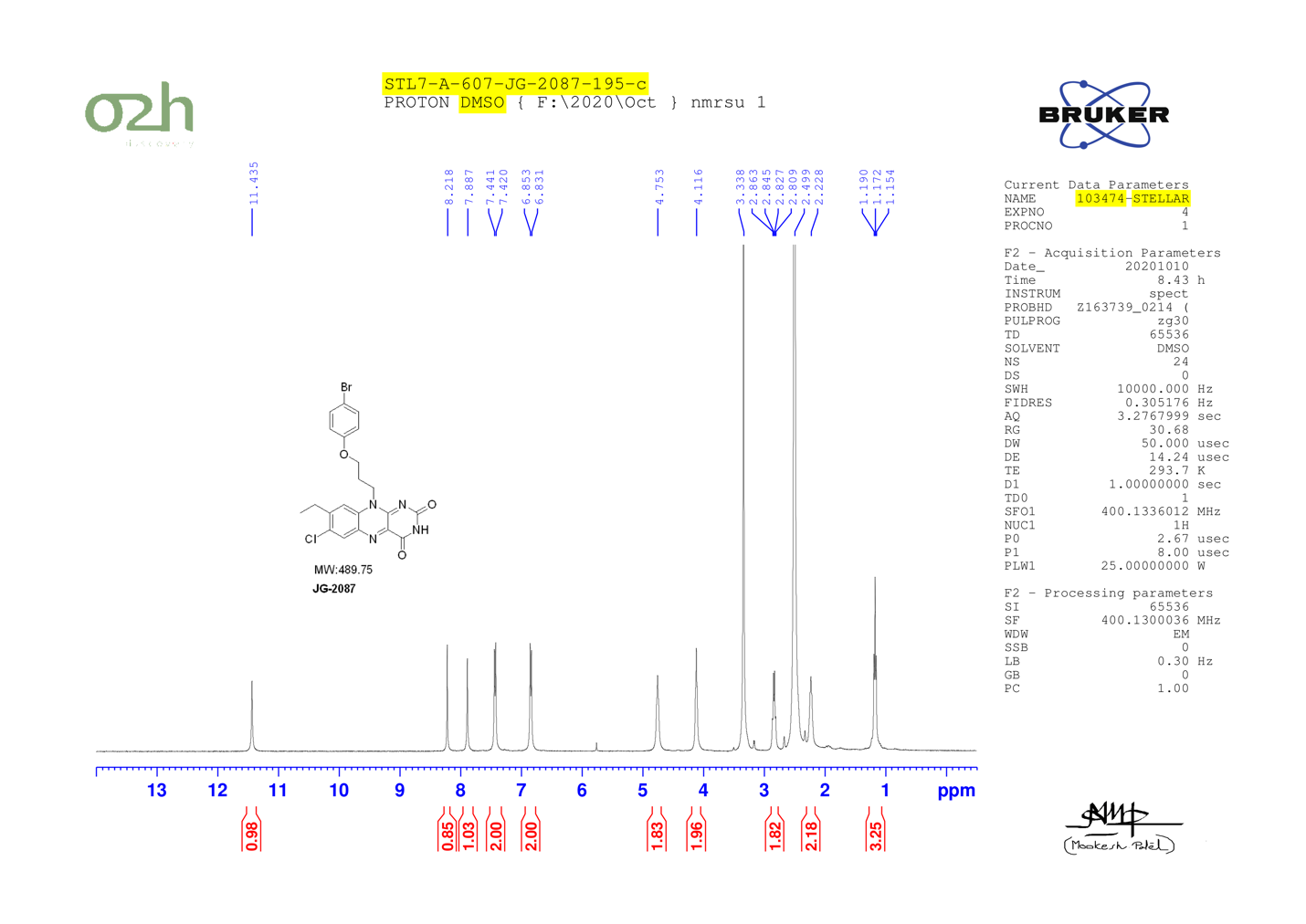

JG-2089

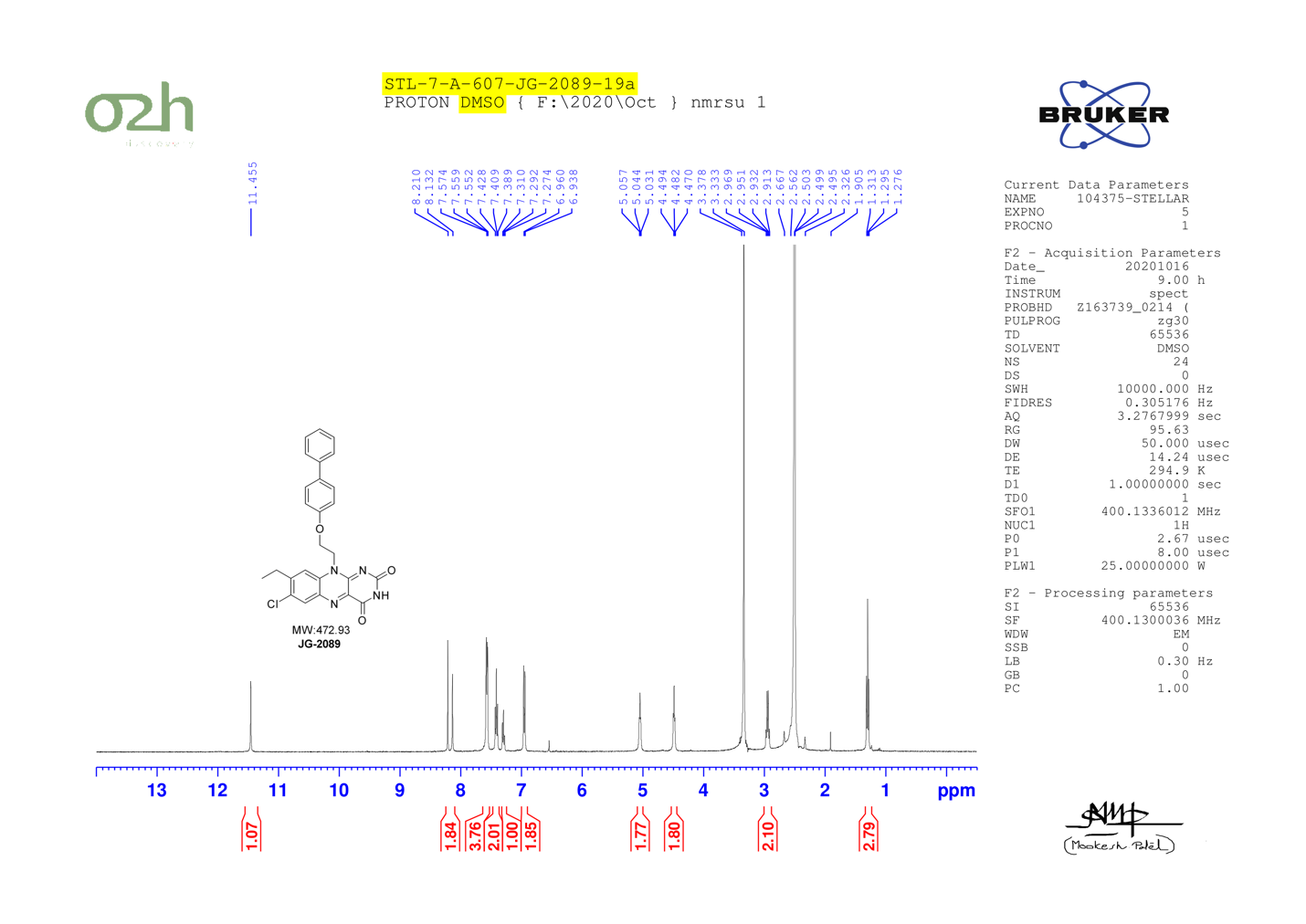

JG-2090

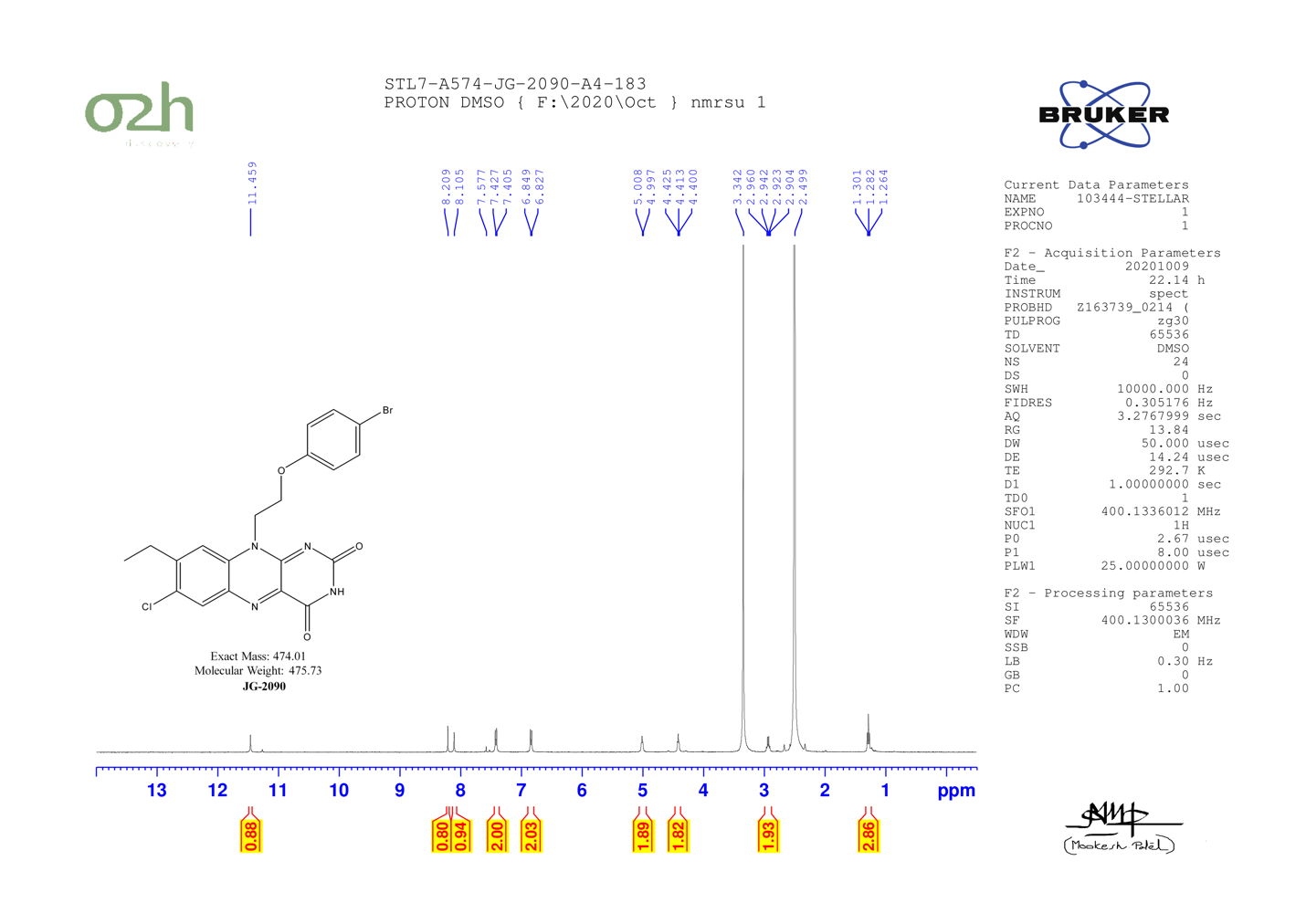

JG-2016 (scale-up full spectrum)

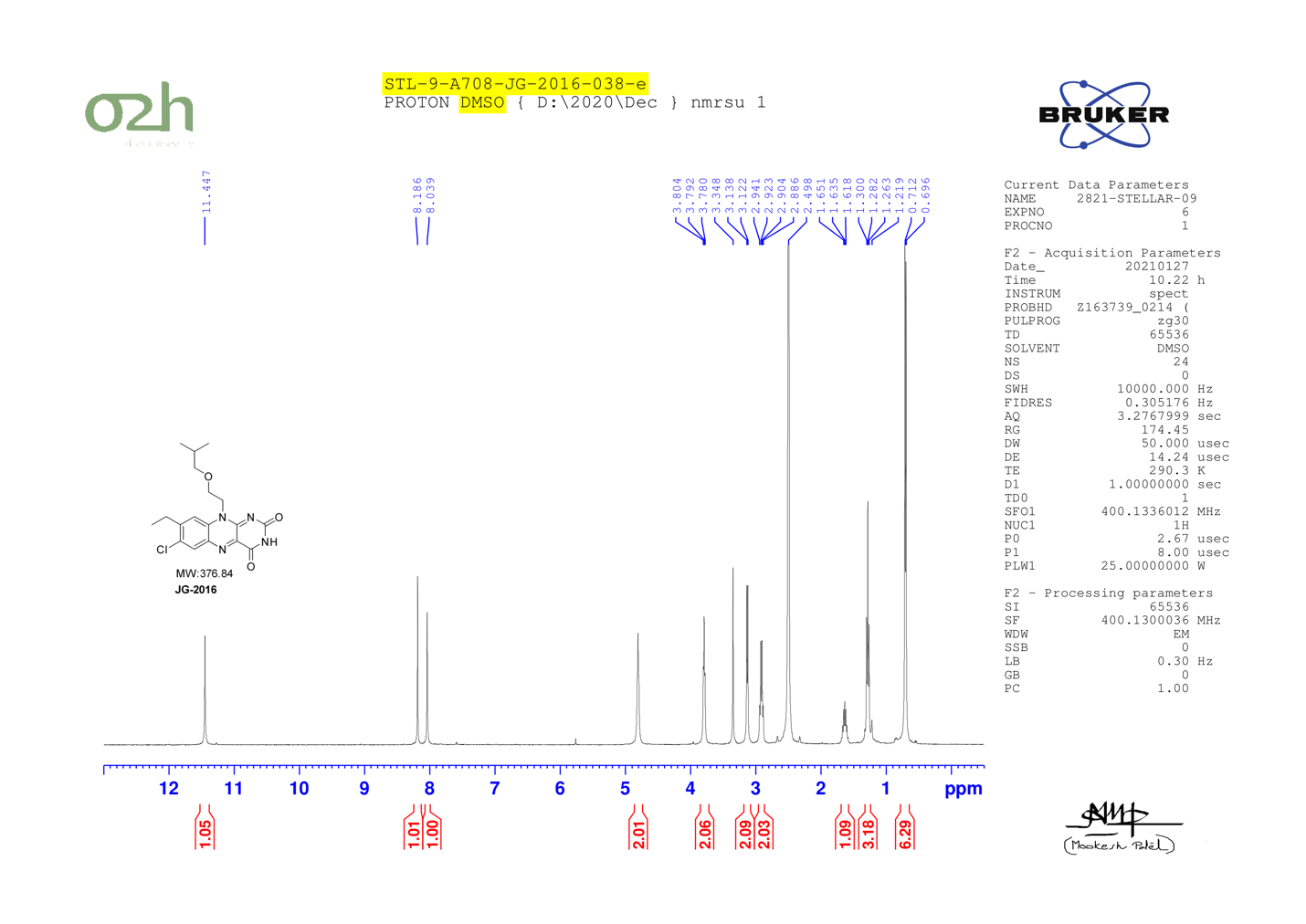

JG-2016 (scale-up zoom: 0.5 - 6.0 ppm)

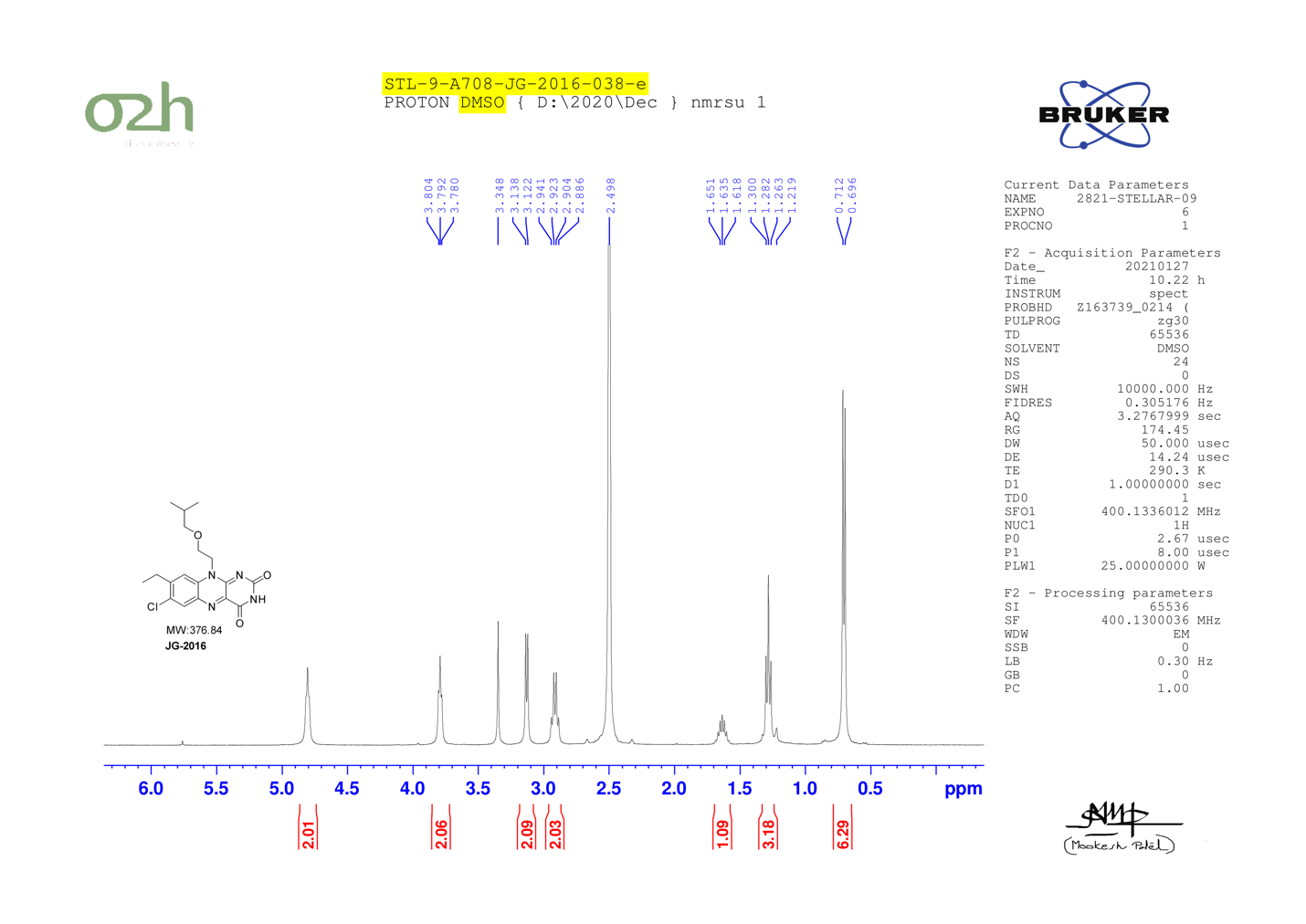

JG-2016 (scale-up, zoom: 6.5 – 12.0 ppm)

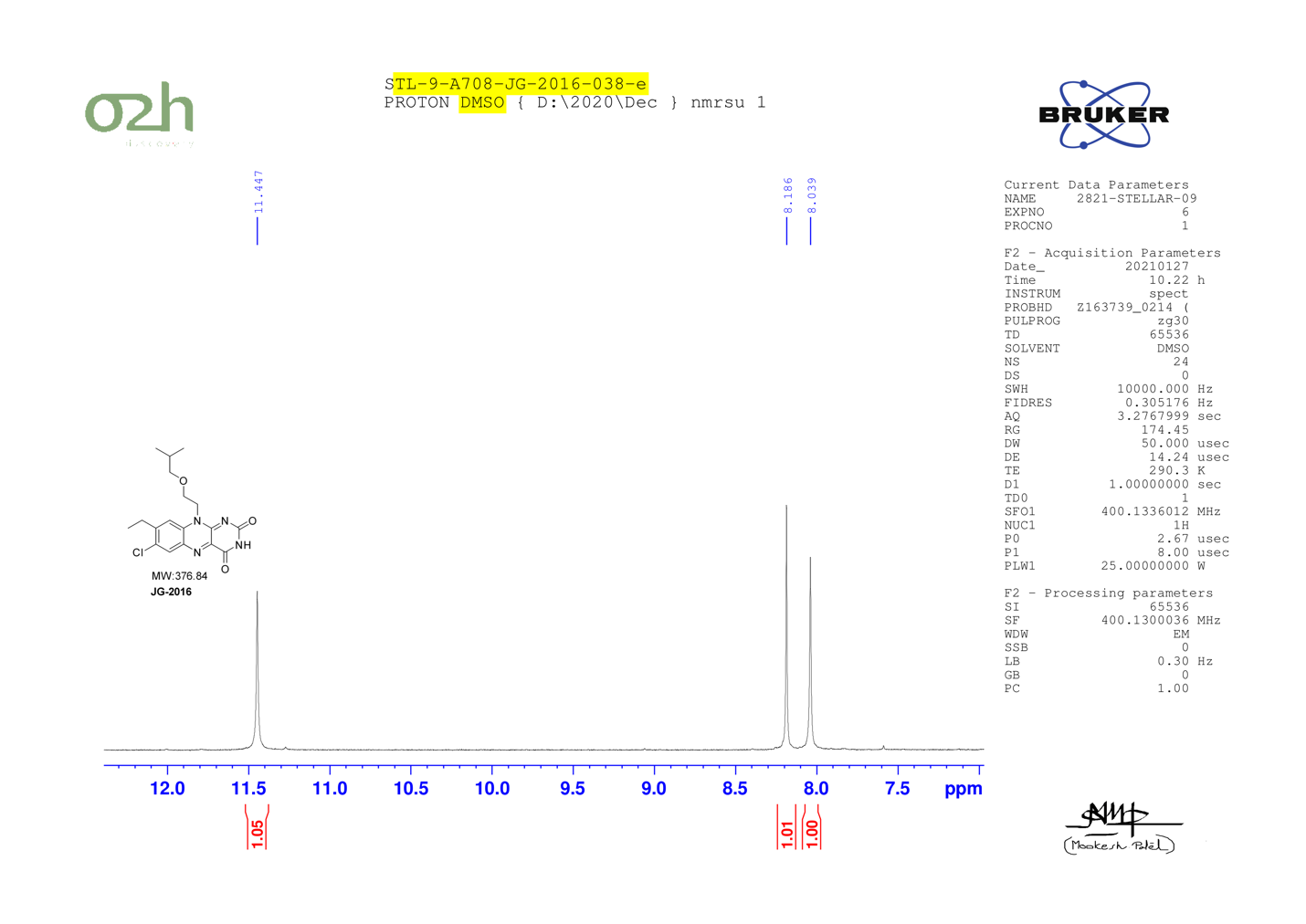
